# Ancestral reconstructions decipher major adaptations of ammonia oxidizing archaea upon radiation into moderate terrestrial and marine environments

**DOI:** 10.1101/2020.06.28.176255

**Authors:** Sophie S. Abby, Melina Kerou, Christa Schleper

## Abstract

Unlike all other archaeal lineages, ammonia oxidizing archaea (AOA) of the phylum Thaumarchaea are widespread and abundant in all moderate and oxic environments on Earth. The evolutionary adaptations that led to such unprecedented ecological success of a microbial clade characterized by highly conserved energy and carbon metabolisms have, however, remained underexplored. Here we reconstructed the genomic content and growth temperature of the ancestor of all AOA as well as the ancestors of the marine and soil lineages based on 39 available complete or nearly complete genomes of AOA. Our evolutionary scenario depicts an extremely thermophilic autotrophic, aerobic ancestor from which three independent lineages of a marine and two terrestrial groups radiated into moderate environments. Their emergence was paralleled by (I) a continuous acquisition of an extensive collection of stress tolerance genes mostly involved in redox maintenance and oxygen detoxification, (II) an expansion of regulatory capacities in transcription and central metabolic functions and (III) an extended repertoire of cell appendages and modifications related to adherence and interactions with the environment. Our analysis provides insights into the evolutionary transitions and key processes that enabled the conquest of the diverse environments in which contemporary AOA are found.

## Introduction

Ammonia oxidizing Archaea of the phylum Thaumarchaeota have a very broad ecological distribution on Earth and are key players in the global Nitrogen cycle. They are generally found as a stable part of the microbial community in studies of aerobic environments, including soils, ocean waters, marine sediments, freshwater environments, hot springs, plants and animals, including humans ^1–3^. This distribution is particularly impressive when considering that even though the domain Archaea includes a large variety of energy and central carbon metabolisms ^4^, no other lineage is known so far to be represented this widely in oxic environments. Other widespread functional guilds of archaea are rather confined to anoxic environments and include methanogenic archaea (of the Classes I & II) and potentially also the so far uncultivated archaea of the novel lineages Bathyarchaeota and Verstraetearchaeota, as well as possibly some lineages of the Asgard and DPANN superphyla ^5–9^.

In contrast to methanogens, AOA form a monophyletic lineage now classified as *Nitrososphaeria* within the phylum Thaumarchaeota ^10^. Based on environmental studies and genomic data, they comprise thermophilic species growing around 70°C ^11–13^, but most cultivated organisms stem from ocean waters and soils ^1,3,14,15^. Phylogenetic and phylogenomic studies have shown that the thermophilic group of *Ca*. Nitrosocaldales from hot springs form a sister group to all AOA adapted to lower temperatures ^11,13,16^. The latter are split into two major lineages, one encompassing the order *Nitrososphaerales* with cultivated (and non-cultivated) representatives residing mostly in soils ^17^. The second major clade is further divided into the orders *Ca*. Nitrosotaleales that are mostly found in acidic soils, as well as the large group of Nitrosopumilales that are mostly found in marine environments ^18,19^. Several cultivated species of AOA have greatly contributed to a better understanding of their physiology and have revealed that AOA as opposed to their bacterial counterparts are adapted to far lower ammonia concentrations^20^ which might explain their ecological success in so many oligotrophic environments. However, a recent meta-analysis based on more than 30,000 *amo*A genes deposited in the public databases revealed that the cultivated strains and enrichments represent only 7 out of 19 total AOA clades, indicating that the (eco-)physiological potential of AOA could be far greater than assumed ^14^. Nevertheless, genomes of cultivated species from diverse environments and a large number of fully or partially assembled (meta-)genomes of AOA are now available and reveal a very well-conserved core of energy and carbon metabolism ^21,22^, with ammonia oxidation catalysed by an ammonia monooxygenase (AMO) and CO_2_ fixed via an extremely efficient version of the 3-hydroxypropionate/4-hydroxybutyrate pathway ^23^. This finding confirms the general assumption that AOA play an important role in the global nitrogen cycle, as ammonia oxidation represents the first step of nitrification, which is an essential process to eventually enable the conversion of reactive nitrogen species via denitrification into the inert gaseous compound N_2_, which is then released to the atmosphere. Their broad occurrence, well-conserved central metabolism as well as their monophyly render AOA an excellent model for elucidating the evolution and adaptations of a microbial lineage that is very likely to have originated in hot springs ^11,12,24–27^ and from there successfully radiated, probably between 1 and 2.1 billion years ago ^28,29^, into most if not all moderate oxic ecosystems on Earth.

In this work we have investigated in a robust phylogenetic framework, 39 complete (or nearly complete) genomes from cultivated or environmental strains of AOA that stem from ocean waters, soils, estuarine, sediments, wastewater, a marine sponge and hot springs. This enabled us to reconstruct the optimal growth temperatures and genome contents of the last common ancestor of all ammonia oxidizing archaea, as well as the ancestors of major AOA lineages to reveal the paths of adaptations that gave rise to radiations into oxic marine and terrestrial environments, respectively.

## Results

### A molecular thermometer suggests a thermophilic ancestor for AOA and subsequent parallel adaptations to mesophily

We compiled a 76 genomes dataset comprising publicly available complete genomes, and high-quality metagenome-assembled or single-cell assembled genomes (see Materials and Methods for selection criteria) from 39 AOA, 13 non-ammonia-oxidizing Thaumarchaeota, 8 Aigarchaeota, 5 Bathyarchaeota and 11 representative genomes of orders within Crenarchaeota to serve as outgroups. We built a species tree from the concatenation of a selection of 33 protein families that were found in at least 74 out of the 76 genomes to serve as a backbone tree for our ancestral reconstructions of growth temperature and genome content (see below) (see Fig. 1, Table S1 and Materials and Methods).

**Figure 1.**
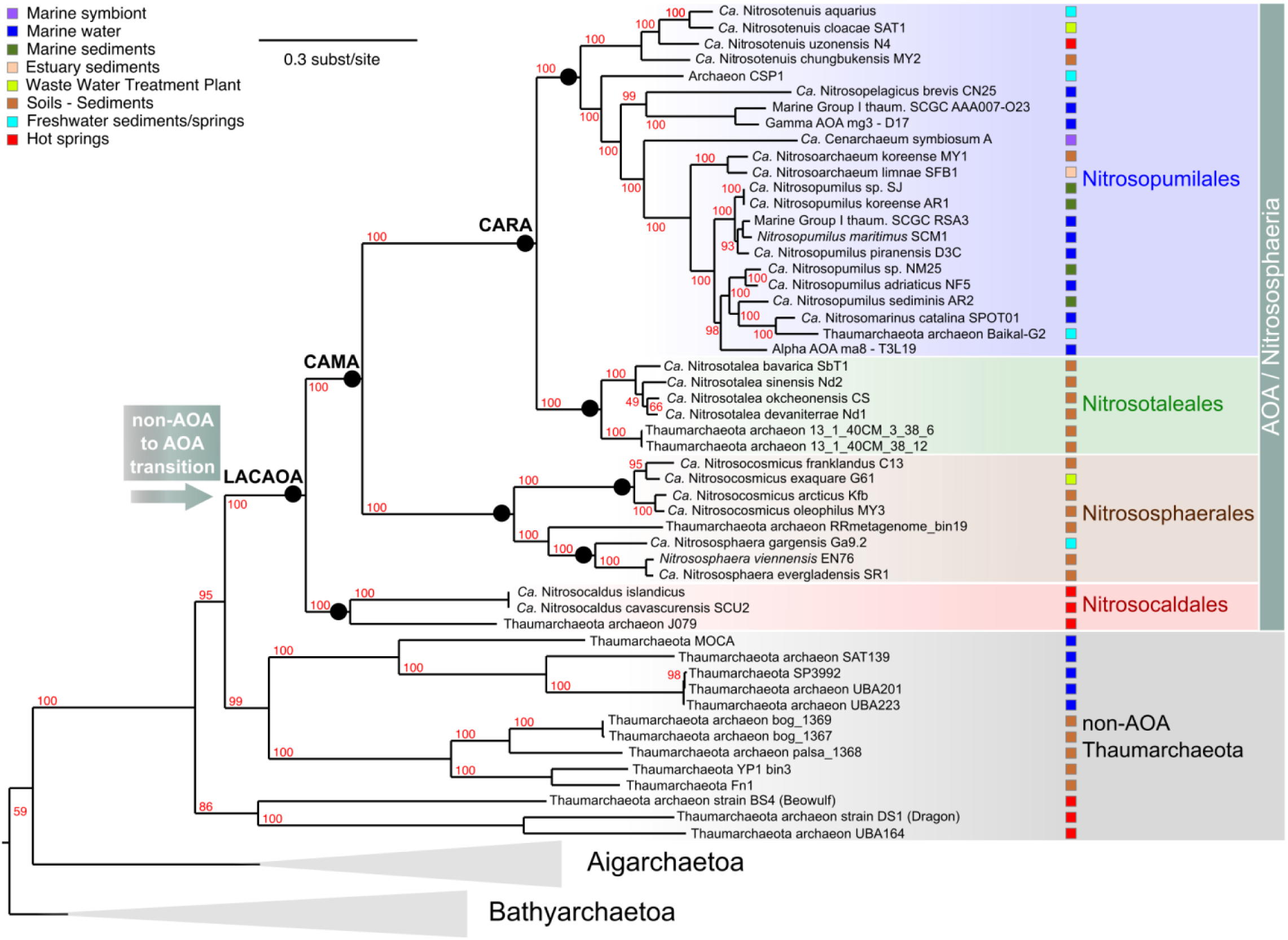
Phylogenetic tree (maximum likelihood) of the 76 genomes analysed, from the concatenation of 33 conserved protein families that were present in 74 out of our 76 genomes dataset (see Table S1). The Crenarchaeota outgroup (11 genomes) is not displayed here, and Bathyarchaeota (5 genomes) and Aigarchaeota (8 genomes) are collapsed. Ammonia oxidizing archaea (AOA) lineages and non-AOA Thaumarchaeota are represented by different colored shades, and their isolation sources are displayed with colored boxes. The tree was built with IQ-tree, using the LG+C60+F model. Supports at nodes are Ultrafast bootstrap supports ^133^.

Our extended tree recovered the major clades of AOA as described earlier with a smaller dataset ^30^, separating all AOA into four orders: *Ca*. Nitrosopumilales from mostly marine environments and sediments and with rather smaller genomes (< 2 Mbp, see Fig. 2), *Ca*. Nitrosotaleales from acidic soils, *Nitrososphaerales* with representatives exclusively from soils and sediments exhibiting genomes of double the size than all other groups (around 3 Mbp) and *Ca*. Nitrosocaldales with organisms exclusively from hot environments. (Fig. 1).

**Figure 2.**
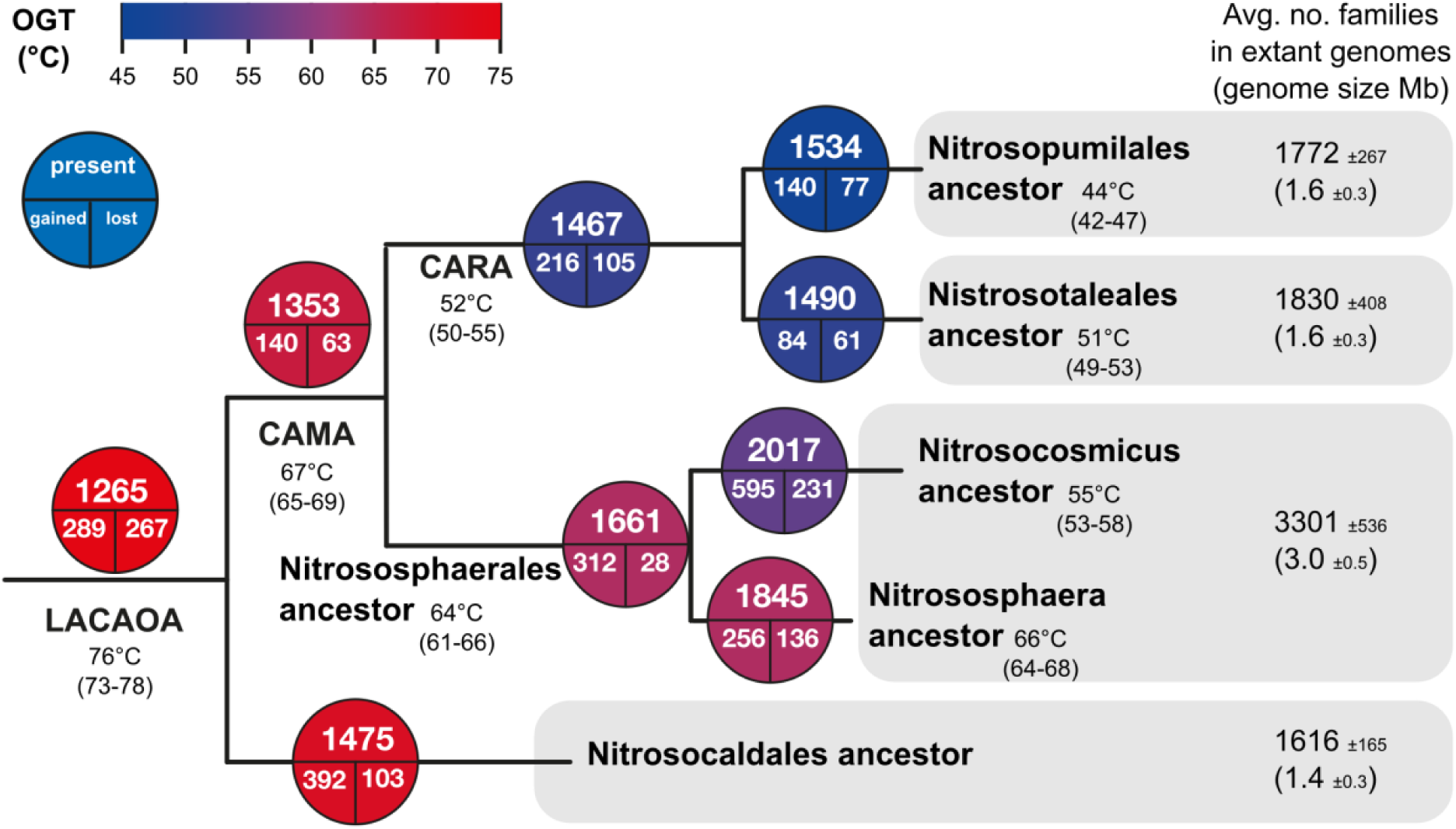
Reconstruction of optimal growth temperatures (OGT) and genome repertoires of AOA ancestors. Numbers of families inferred as having a probability higher than 0.5 to be present, gained or lost are displayed in circles at each node symbolizing the corresponding ancestral genomes and their dynamics (see main text). Ancestral OGT are displayed at the corresponding nodes (top number) together with their confidence interval (number range below). As there was no 16S rRNA sequences available for the J079 genomic bin, we could not infer OGT for the ancestor of Nitrosocaldales. The color of the bubbles represents the transition from hot (red) to colder (blue) growth temperature. For each candidate order of AOA, the number of protein families found on average in their respective genomes is indicated, as well as the average genome size (along with standard deviations). LACAOA stands for “Last Common Ancestor of AOA”, CAMA for “Common Ancestor of Mesophilic AOA” and CARA for “Common Ancestor of Rod-shaped AOA”.

In order to approach the optimal growth temperatures (OGTs) of the ancestor of all AOA and of ancestors of terrestrial and marine lineages, we used a “molecular thermometer” to perform a linear regression that enabled to predict the OGTs from the predicted ancestral 16S rRNA G+C stem compositions as in ^31,32^ (see Table S2, Fig. S1, Materials and Methods and Supplementary Information). Not all genomes could be represented as 20 of them were missing assigned 16S rRNA genes (Table S1). The OGTs of AOA decreased along time, from around 76°C for the <u>LA</u>st <u>C</u>ommon Ancestor of <u>AOA</u> (LACAOA), to 67°C for the <u>C</u>ommon <u>A</u>ncestor of the <u>M</u>esophilic clades of <u>A</u>OA (CAMA), to 64°C and 52°C for the ancestors of *Nitrososphaerales*, and *Ca*. Nitrosopumilales plus *Ca*. Nitrosotaleales (the ancestor of the latter two hereafter called “CARA”, for <u>C</u>ommon <u>A</u>ncestor of <u>R</u>od-shaped <u>A</u>OA including marine and acid soil AOA), respectively (Fig. 2). It thus seems that the ancestors of the three different major orders of AOA were all thermophilic, and parallel adaptations to lower temperatures occurred along time in these lineages. Interestingly, the order *Nitrososphaerales*, containing moderately thermophilic species like *Ca*. Nitrososphaera gargensis (OGT = 46°C) and *N. viennensis* (OGT = 42°C), seemed to have preserved a relatively high G+C content in their 16S rRNA, especially in the genus *Nitrososphaera* (predicted ancestral OGT of 64 - 68°C), which might result from a recent or still ongoing adaptation to lower temperatures from thermophilic ancestors. Based on this analysis, *Ca*. Nitrosopumilales have most drastically adapted to lower temperatures, which is in line with their adaptation to colder, marine environments. Since we infer an ancestor for all AOA with an OGT around 76°C (73°C-78°C), we conclude that the ability to oxidize ammonia for energy acquisition in Archaea was most probably acquired in thermophilic or even hyperthermophilic organisms, as has been suggested since their discovery, and also through subsequent findings of deeply branching thermophilic lineages within Thaumarchaeota and AOA ^11–13,24–27^.

### Genome dynamics and reconstruction of ancestral AOA genomes reveal distinct evolutionary paths from thermophilic to moderate AOA lineages

In order to investigate the evolutionary path of metabolic and adaptive features of AOA, we reconstructed the gene repertoire of LACAOA and of subsequent ancestors by analyzing the distribution of families of homologous proteins from our 76 genomes dataset, and inferring their gains and losses along the species tree with a birth-and-death model for gene family size evolution implemented in the Count program ^33^ (Table S3). Our analysis enabled us to reconstruct key ancestral states on the evolutionary path of AOA (Fig. 1, 2), such as that of LACAOA, that of CAMA, the Common Ancestor of Mesophilic AOA (after the divergence of *Ca*. Nitrosocaldales), that of CARA, the Common Ancestor of Rod-shaped AOA (the common ancestor of *Ca*. Nitrosotaleales and the mostly marine Nitrosopumilales), and the ancestor of *Nitrososphaerales* (encompassing mostly terrestrial lineages), as well as the ancestors of the genera *Ca*. Nitrosocosmicus and *Nitrososphaera*.

We infer that at least 1265 protein families were present in LACAOA, the ancestor of AOA. Along the transition from non-AOA to AOA, 289 families were inferred to be gained, and 267 lost (Fig. 1, 2 and Tables S3, S4). The huge metabolic transition provoked by the adoption of an ammonia-oxidizing metabolism corresponded to a slightly increased gains’ rate for LACAOA (i.e. number of gains events divided by branch length) when compared to other Thaumarchaeota ancestors (Fig. S2, Wilcoxon signed rank test, p-value ∼ 0.001), yet the general gains’ rate trend was consistent with an overall continuous flux of genes along AOA evolution rather than a massive influx of genes that would have marked the onset of this lineage (Sup. Info., Fig. S2).

Despite the streamlined genomes of contemporary Nitrosopumilales (1.6 Mbp on average) compared to *Nitrososphaerales* (3.0 Mbp on average), their ancestors had very similar predicted family content (1534 and 1661, respectively) (Fig. 2 and Table S3). Therefore, most of the differences in genome sizes can be attributed to the massive gains that occurred rather continuously within *Nitrososphaerales*, with 312 gains by the ancestor of *Nitrososphaerales* and after the split between *Ca*. Nitrosocosmicus and *Nitrososphaera*, respectively (595 and 256 gains in the ancestors of the *Ca*. Nitrosocosmicus clade and *Nitrososphaera* clade, respectively). This parallel increase in genome sizes of the two ‘soil’ clades led to distinct genomic repertoires and physiologies (see following sections). Our inferences of ancestral genome sizes differed from those recently obtained in ^24^, resulting in apparent distinct genome dynamics with larger ancestral genomes and more downstream losses in the latter. The Dollo parsimony used in that study to infer gains and losses with Count was earlier deemed inappropriate for studying prokaryotic genomes where lateral gene transfers are common ^34^. Therefore, this difference in Count settings is likely to account for the differences observed between the studies. Furthermore, the probabilistic setting we used, where gains and losses rates were estimated for each family separately in this study, is likely to be more realistic.

In the following sections, we detail the evolutionary path followed by AOA towards adaptations to moderate environments, starting by describing the ancestor of all AOA. In order to confirm the evolutionary scenarios inferred by the Count program we used comprehensive phylogenetic reconstructions for approximately 80 crucial families discussed in the rest of this manuscript (see Table S5). This allowed us to correct - where needed - the evolutionary scenarios inferred by the program Count, and in some cases to make assumptions on the donating lineage (see Materials and Methods, Data S1 and Table S5).

### The last common ancestor of AOA was a thermophilic, autotrophic aerobic organism

#### Metabolic acquisitions by LACAOA

As expected, LACAOA acquired the three genes for the ammonia monooxygenase complex (amoABC) and the fourth candidate subunit (amoX) ^35^, the urease gene set and two urea transporters (SSS and UT types)^12,24,36^ and expanded its set of pre-existing ammonia (Amt) transporters (from one to two, as in extant AOA) (Fig. 3 and Table S4). Although the evolutionary history of the archaeal AMO is not trivial to resolve, recent phylogenetic analyses indicated that it is more closely related to actinobacterial hydrocarbon monooxygenases than the bacterial AMO_14_. It should also be noted that none of the non-AOA Thaumarchaeal genomes to date encode any genes related to ammonia oxidation or urea utilization. However, one has to note that the pathway has not been fully elucidated yet. Meanwhile, a number of cupredoxin-domain families were acquired, equipping it with the unique copper-reliant biochemistry encountered in contemporary AOA. The heavy reliance on copper necessitated the acquisition of copper uptake systems, such as the CopC/CopD family proteins ^37^. Curiously, glutamate synthase (GOGAT), which in most Archaea is participating in the high affinity ammonia assimilation pathway, is inferred to be lost at this stage, leaving the glutamate dehydrogenase (usually exhibiting lower affinity ^38^) as the primary ammonia assimilation route in contemporary AOA. This could be regarded as an adaptation to an energy and carbon-limited lifestyle as less carbon and ATP are consumed during NH_3_ assimilation. It is also the preferred strategy under these conditions in some bacteria ^39^.

**Figure 3.**
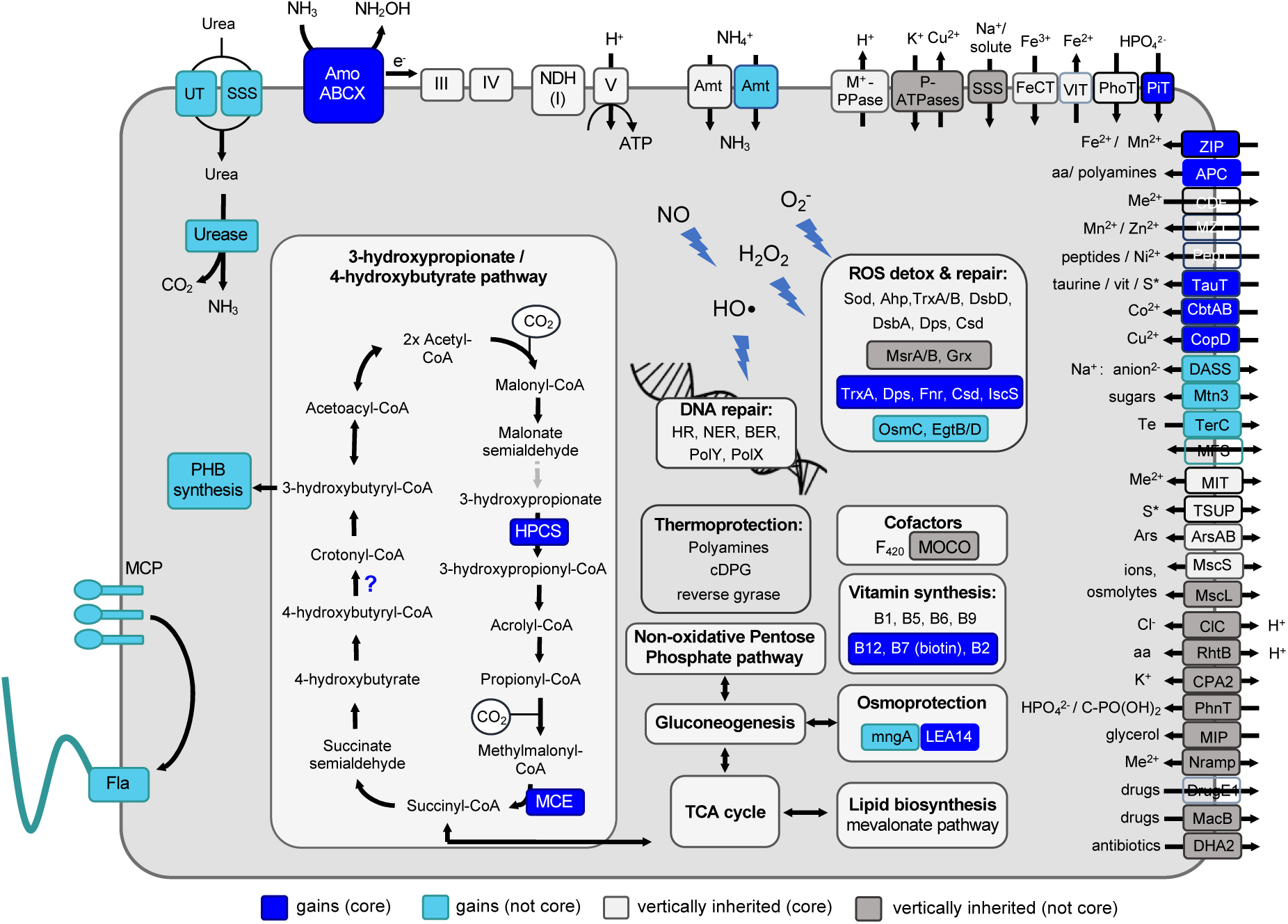
Reconstruction of main metabolic and physiological features and transport capabilities of the <u>LA</u>st <u>C</u>ommon ancestor of <u>AOA</u> (LACAOA). Metabolic modules inferred to be vertically inherited are in light gray, if present in all currently known extant AOA, or light blue if lost in certain lineages (not core). Newly acquired capacities that became core components for all extant AOA are depicted in blue, or in turquoise if they were more scattered throughout extant AOA genomes (not core). Gray lines indicate reactions for which a candidate enzyme is not identified. Due to the inherent difficulty in annotation and substrate prediction, transporters were considered as part of the core genome if found in all ancestors examined in this study, or if a functionally equivalent family exchange took place in the ancestors, but their distribution in extant genomes was not taken into account. Gradient boxes are used in the case of multiple families belonging to different categories. Transporters are named according to the TCDB classification (Table S4 ^134^), with indicative substrates and directionality of transport. Question mark indicates an unclear family history, see text for details. Abbreviations (in addition to the main text): S*, organo-sulfur compounds/sulfite/sulfate; Me^2+^, divalent cations; aa, aminoacids; AcnA, AcnA/IRP group aconitase; MCP, methyl-accepting chemotaxis proteins; Fla, archaeal flagellum (or “archaellum”).

Similarly, at least two genes for crucial steps in the 3-hydroxypropionate/4-hydroxybutyrate (HP/HB) CO_2_ fixation pathway conserved in all AOA were acquired by LACAOA. These were an ADP-forming 3-hydroxypropionyl-CoA synthetase (hpcs), responsible for the energy efficiency of the cycle ^23^, possibly acquired from Euryarchaeota (Data S1), and methylmalonyl-CoA epimerase (mce). A third component of the pathway, the ADP-forming 4-hydroxybutyryl-CoA synthetase, was replaced at the stage of mesophilic AOA (i.e. in CAMA). LACAOA might also have newly acquired the O_2_-tolerant enzyme 4-hydroxybutyryl-CoA dehydratase, (see also Fig. S2 in ^12^). Overall, the carbon fixation pathway found in extant AOA seems to be the result of the integration of mostly vertically inherited, and some more recently acquired enzymes.

The key enzyme for production of polyhydroxyalcanoates (PHAs), PHA synthase, was also acquired by LACAOA (Data S1), equipping AOA with the ability to produce carbon storage compounds typically synthesized in response to unbalanced growth conditions ^40^. Additionally, LACAOA acquired the families associated with the biosynthesis of the vitamin B cofactors cobalamin (B12), biotin (B7) and riboflavin (B2) as pointed out in earlier studies ^29,41^.

#### Genes for aerobic autotrophic growth were already present in LACAOA

The gene families inferred to be present already, and not gained by LACAOA (labeled light and dark grey in Fig. 3), illustrate the genetic background of the archaeon which first acquired the metabolism of ammonia oxidation (Tables S3, S4). The ancestor of LACAOA was capable of using O_2_ as an electron acceptor as suggested by the presence of a heme-copper oxidase (routinely used to infer an aerobic metabolism ^24,42^) and the absence of an alternative electron acceptor. The inference of an aerobic ancestor stands in contrast to a recent study ^29^ that concluded LACAOA arose from an anaerobic ancestor, based on the assumption that all lineages basal to AOA were anaerobic (i.e. non-AOA Thaumarchaeota, Aigarchaeota and Bathyarchaeota). However, most Aigarchaeota and at least two non-AOA Thaumarchaeota (BS4 and pSL12) included in our dataset were predicted to be facultative aerobes – the latter being recently further supported by a study reporting metagenomic-assembled genomes from the pSL12 lineage with potential for aerobic respiration ^24,43–46^. In our ancestral reconstructions, genes involved in anaerobic metabolisms were inferred as gains in the respective lineages, such as the codH subunits in the ancestor of Bathyarchaeota, or nitrate reductase and adenylylsulfate reductase families in Thaumarchaea BS4 and DS1. Additionally, the presence of most enzyme families of the HP/HB carbon fixation pathway (including the key enzymes acetyl-CoA/ propionyl-CoA carboxylase and 4-hydroxybutyryl-CoA dehydratase) indicate the potential for autotrophic growth. They are present in all known AOA and shared with a number of aerobic Crenarchaeota and Aigarchaeota, albeit with distinct differences (see paragraph above and ^23,47,48^).

Concurrently, we observe a loss of gene families associated with heterotrophic growth, in particular those involved in glycolysis and connecting pools of C3 and C4 compounds. Among them are two phosphofructokinases, the 2-oxoacid dehydrogenase complex (OADHC), and an (abcd)^2^-type 2-oxoacid:ferredoxin oxidoreductase (OFOR, while an (ab)^2^-type OFOR exclusively found in aerobes is retained in LACAOA and present in all extant AOA (see Supplementary Information and Data S1). Losses and gene families’ contractions (loss of gene copies in an ancestrally multi-copy family) were also observed in families involved in amino-acid catabolism such as the glycine cleavage system and amino-acid and sugar transporter families (see Supplementary Information for details).

Overall, our analysis depicts an autotrophic aerobic ancestor, which accompanied the switch to ammonia oxidation with an apparent minimization of the ability to grow heterotrophically. Whether this ability was completely lost in the emerging AOA lineages remains an open question, as many genomic studies have indicated the potential for mixotrophic growth and many isolates benefit from the addition of organics (although this effect was later attributed to ROS detoxification^49^). Nevertheless, assimilation of organic carbon has so far not been demonstrated in AOA.

### Radiations to moderate environments were paralleled by independent adaptations in three major functional categories

In accordance with an inferred adaptation to lower temperatures, several traits involved in the thermophilic lifestyle of LACAOA were lost in subsequent evolutionary steps: reverse gyrase, the hallmark enzyme of organisms living at extremely high temperatures ^50,51^ and cyclic 2, 3-diphosphoglycerate (cDPG, thermoprotectant ^52^) synthetase were lost in CAMA, while the thermoprotectant mannosyl-3-phosphoglycerate synthase (MPGS) was lost in CARA (Tables S3, S4, Fig. 4). In contrast, a homologue of the “cold-shock protein” CspA ^53^ was found to be acquired from bacteria in an ancestor of *Ca*. Nitrosopumilales (to the exception of the soil-residing genus *Ca*. Nitrosotenuis) (Fig. 4, Fig. S3, Table S3), corroborating the generally lower OGT of the marine lineages, as inferred by our analysis and existing pure cultures.

**Figure 4.**
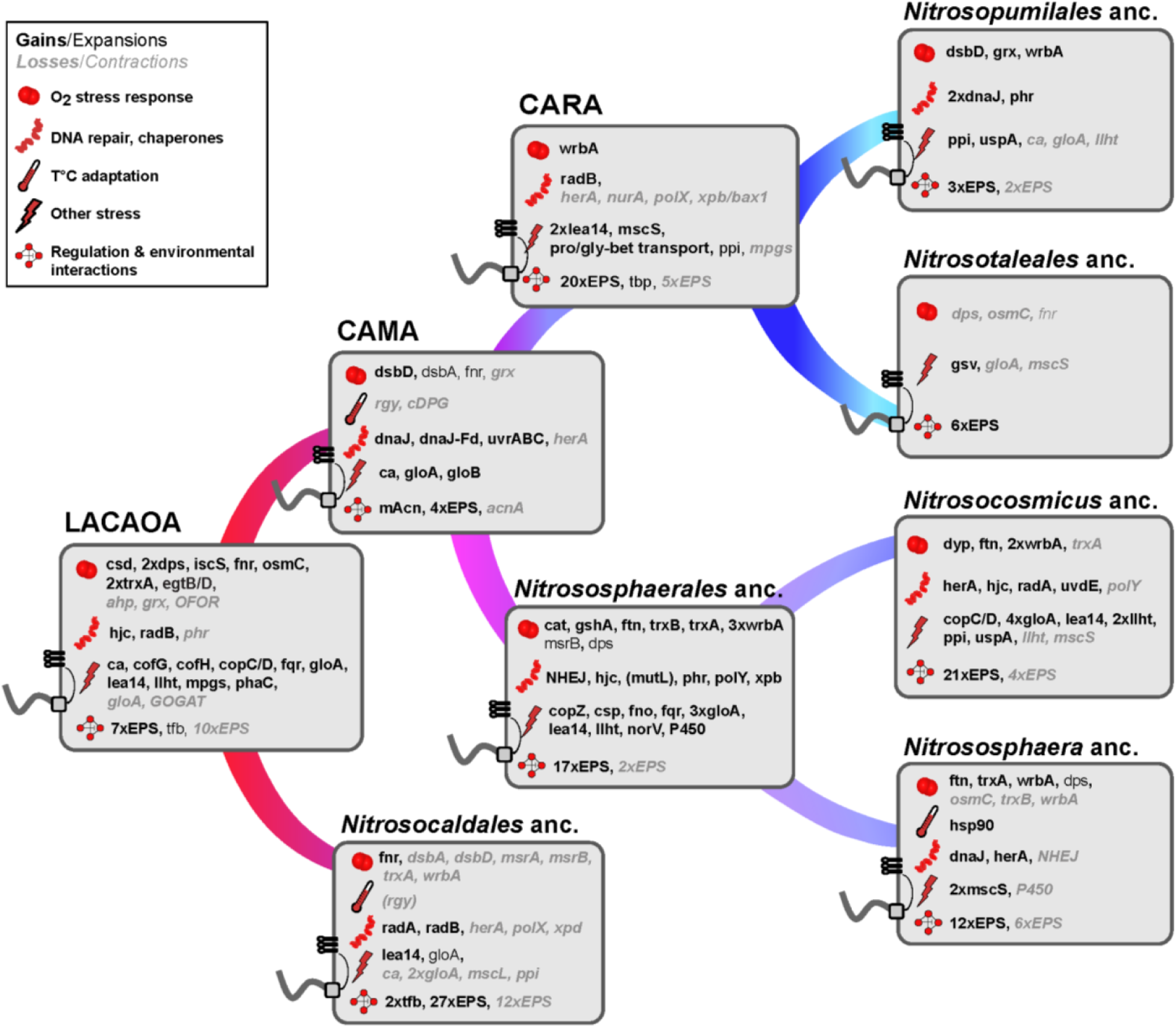
Summary of crucial gene gains and losses in ancestors of AOA and subsequently evolved lineages. The evolutionary events (gains/expansions in black, losses/contractions in grey) related to each stress category are displayed next to the corresponding symbol. Ancestral cells inferred to have been flagellated harbor a schematic flagellum and chemotaxis receptors. For discussion of functions see main text and Table S4 & Supplementary Information). Abbreviations: acnA, AcnA/IRP type aconitase; mAcn, mitochondrial-like aconitase; ahp, alkyl hydroperoxide reductase; ca, carbonic anhydrase; cat, catalase; cDPG, cyclic 2, 3-diphosphoglycerate synthetase; cofG/H, FO synthase subunits 1/2; copZ, copper binding protein; copC/D, copper resistance family proteins; csd/iscS, cysteine desulfurase families; csp, four-helix bundle copper storage protein; dps, DNA protection during starvation family protein; dsbA/D, disulfide bond oxidoreductase A/D; dyp, dyp-type peroxidase; EPS, extracellular polymeric substances synthesis families; fqr, F420H(2)-dependent quinone reductase; fno, F420H2:NADP oxidoreductase; fnr, flavodoxin reductase (ferredoxin-NADPH reductase); fprA/norV, flavorubredoxin; ftn, ferritin-like protein; gloA/B, glyoxalase I/II; GOGAT, glutamate synthase; grx, glutaredoxin; gshA, glutamate-cysteine ligase; gsv, gas-vesicle formation genes; herA, HR helicase; hjc, holliday-junction resolvase; lea14-like, LEA14-like desiccation related protein; llht, luciferase-like hydride transferase; mpgs, mannosyl-3-phosphoglycerate synthase; mscL/S, large-and small-conductance mechanosensitive channel respectively; msrA/B, methionine sulfoxide reductase A/B; mut-L, putative DNA mismatch repair enzyme MutL-like; NHEJ, non-homologous end joining; nurA, HR nuclease; OFOR, 2-oxoacid:ferredoxin oxidoreductase; OsmC, peroxiredoxin; P450, cytochrome p450-domain protein; phaC/E, poly(R)-hydroxyalkanoic acid synthase subunits C/E; phr, photolyase; ppi, peptidylprolyl isomerase; polY/X, DNA repair polymerases Y & X; ProP (proline/glycine-betaine):(H+/Na+) symporter; radA/B, DNA repair and recombination proteins; rgy, reverse gyrase; tbp, TATA-box binding protein; tfb, transcription initiation factor TFIIB homolog; trxA, thioredoxin; trxB, thioredoxin reductase; wrbA, multimeric flavodoxin WrbA; dnaJ/dnaK/grpE/dnaJ-ferredoxin, chaperones; uspA, universal stress protein family A; uvrABC/uvdE, ultraviolet radiation repair excinuclease ABC system/UV damage endonuclease uvdE; xpb/bax1, NER helicase/nuclease pair; xpd, NER helicase.

When surveying the functions of families acquired and lost along the evolution of mesophilic AOA, we observed that the majority of acquisitions in all examined ancestors, ranging from 18-34% of all functionally annotated families (and a few key losses), could be classified into three major categories, discussed further below: i) adaptations to various kinds of abiotic stress, in particular oxygen-related stress ii) specific metabolic regulations and general increase of the regulatory potential, and iii) extension of capacities to engage in complex interactions with the environment (Fig. 4, Fig. S3 Table S4).

### 1. Stress adaptations were crucial to colonize ‘moderate environments’

#### Strategies against oxidative and nitrosative stress

Although LACAOA, being an aerobe, was already equipped with a set of proteins dedicated to reactive oxygen species (ROS) detoxification, redox homeostasis and ROS damage repair (Fig. 3 and Tables S3, S4), a number of additional genes involved in these processes were frequently and continuously acquired along all lines of AOA evolution. Some of these specific acquisitions also shaped in distinct ways the genomic repertoire of entire clades, determining their ecological boundaries.

Thiol oxidoreductases of the thioredoxin (Trx) and disulfide-bond reductase proteins (Dsb) family, involved in reducing oxidized cysteine pairs, ^54^ were continuously acquired throughout all AOA lineages (Fig. 4, Fig. S3, Supplementary Information), while the methionine sulfoxide reductase B (MsrB) family, responsible for the reduction of oxidized methionines, was expanded only in *Nitrososphaerales*. Thioredoxin reductase and MsrA families were present in LACAOA and inferred to be vertically inherited. Dsb and Msr family proteins may also be involved in detoxification of reactive nitrogen species (RNS) such as NO ^55,56^, produced by AOA (as in other ammonia oxidizing bacteria) as an intermediate of ammonia oxidation ^57–59^. Interestingly, only the ancestor of *Nitrososphaerales* acquired a Mn-catalase, apparently from Terrabacteria (as suggested by phylogenetic analyses, see Data S1), which was lost in *N. viennensis*), whereas *Ca*. Nitrosopumilales, *Ca*. Nitrosocaldales and *Ca*. Nitrosotaleales all lack this enzyme, considered to be a hallmark enzyme for aerobic metabolisms (Fig. 5, Fig. S3). Although this absence remains puzzling, in Bacteria catalases are the primary scavengers only at high H_2_O_2_ concentrations, while the activity of alkyl hydroperoxide reductase (Ahp, an enzyme present in LACAOA and vertically inherited in all extant AOA) is sufficient during logarithmic growth ^60^. Nevertheless, most AOA lacking catalase have been shown to be dependent on, or stimulated by, the presence of an external H_2_O_2_ scavenger, such as catalase, dimethylthiourea or α-ketoacids ^49,61^, or were/are dependent on co-cultures with bacteria ^62,63^. Stress reduction is a known factor shaping microbial communities^64^, and these interdependencies often are the reason behind the low success rates in isolating and characterizing novel species, as is the case with AOA.

**Figure 5.**
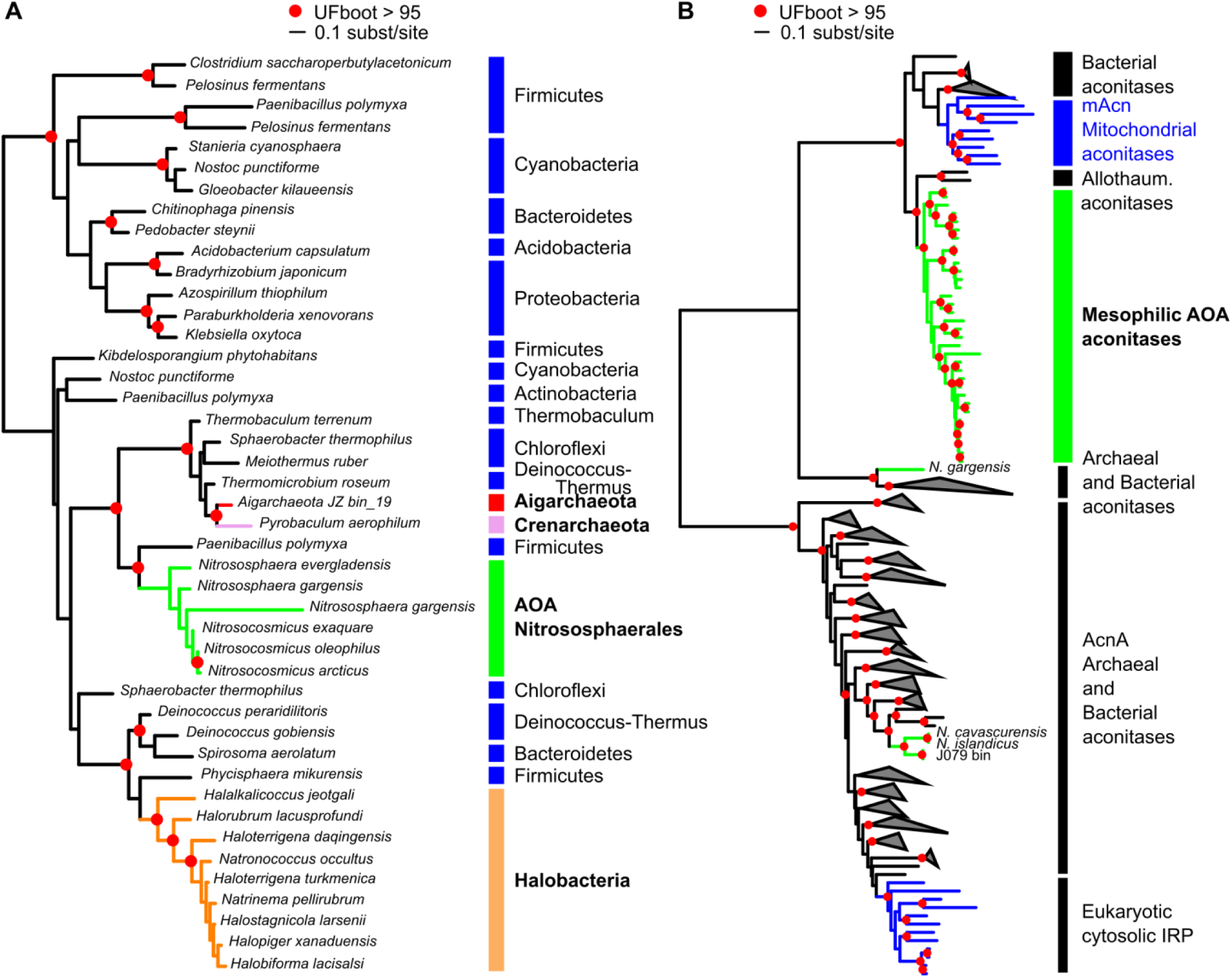
**A**. Phylogenetic tree of the catalase family. A maximum-likelihood phylogenetic tree was obtained for FAM002068. AOA sequences are shown in green, bacterial ones are in black. Archaeal clades are indicated in bold font. **B**. Phylogenetic tree of the Aconitase family. A maximum-likelihood phylogenetic tree was obtained for FAM001301. AOA sequences are shown in green, and eukaryotic ones in blue. Grey triangles correspond to collapsed groups of archaeal and bacterial aconitases. In both panels, branches with UF-Boot support above 95% are indicated by a red circle.

The role of low molecular weight thiols (LMWT) in intracellular redox control and oxidative stress defense is undisputed ^65^. While the most well-known LMWT, glutathione, has not been detected in Archaea, they have nevertheless been shown to accumulate a variety of analogs such as thiosulfate and Coenzyme A ^66^. The key genes (egtB and egtD) for the synthesis of the LMTW ergothioneine (EGT), a histidine betaine derivative with a thiol group synthesized by fungi and bacteria, were gained in LACAOA. In other archaeal phyla they are only found in a few representatives of Methanosarcinales and Thermoplasmatales. A donor could not be identified, but the origin of the genes possibly lies within Actinobacteria ^67^.

Another notable acquisition by the ancestor of *Nitrososphaerales* is the protein family encoding for a gamma-glutamylcysteine ligase (gshA), the evolutionary origin of which could not be unambiguously traced. This enzyme catalyzes the formation of gamma-glutamylcysteine from glutamate and cysteine, typically a precursor of glutathione biosynthesis in Bacteria and Eukaryotes ^68^. Accumulation of this compound was shown in Haloarchaea ^69,70^, making *Nitrososphaerales* the second archaeal group to follow this strategy. The second enzyme responsible for glutathione biosynthesis, gshB is missing in both Haloarchaea and Thaumarchaeota, while candidates for a putative bis-gamma-glutamylcystine reductase can be found among the pyridine nucleotide-disulfide reductase family homologs present in all AOA.

#### Iron, copper and redox homeostasis

Systems that control the levels of unincorporated iron and copper in the cell are upregulated upon oxidative stress in Archaea and Bacteria ^71^ (Supplementary Information). Both Fe^2+^ and Cu(I) can react with H_2_O_2_ and generate hydroxyl radicals (HO•) via the Fenton and Haber-Weiss reactions respectively ^72^, which can subsequently cause severe cellular damage, including DNA lesions. Dps family proteins (DNA-binding proteins from starved cells), known archaeal antioxidants consuming both substrates of the Fenton reaction and physically shielding the DNA ^71,73^, were acquired by LACAOA and expanded in *Nitrososphaerales*. Dps and ferritin-like superfamily proteins were also gained as part of the parallel expansion of the soil lineages in the *Ca*. Nitrosocosmicus and *Nitrososphaera* ancestors (Fig. 4, Fig. S3). Cu(I) can also cause the displacement of iron in enzymatic Fe-S clusters ^74^. Interestingly, a Cys-rich four-helix bundle Copper storage protein (Csp), recently identified in methanotrophs ^75^, was acquired in the ancestor of *Nitrososphaerales*, probably from Bacteria.

Enzymes involved in maintaining redox homeostasis, an essential function to prevent or alleviate oxidative stress, were gained throughout the evolution of AOA. In particular, a ferredoxin:NADP(H) oxidoreductase (FNR) family was gained by LACAOA and expanded in CAMA, while WrbA flavodoxins were respectively acquired by CARA, the ancestor of *Ca*. Nitrosopumilales, the ancestor of *Nitrososphaerales* and the subsequent terrestrial lineages (Table S4, Fig. 4, Fig. S3, Supplementary Information).

#### Frequent exchanges of pathways involved in desiccation, protein stability and osmotic stress relief occurred during the colonization of osmotically variable environments

LACAOA acquired the key enzyme mannosyl-3-phospholgycerate synthase (MPGS) (Data S1), responsible for the first of the two-step synthesis pathway of the compatible solute mannosylglycerate (MG), whose role beyond osmotic stress involves maintenance of protein structure and activity during freezing, desiccation, ROS detox, and thermal denaturation ^76,77^. Interestingly, the pathway for MG synthesis was lost in CARA. The emerging, mostly marine lineages instead acquired the ability to synthesize or take up ectoine/hydroxyectoine (gained by the ancestor of the *Nitrosopumilus* genus ^78^) and proline/glycine-betaine (gained in CARA) during the adaptation to osmotically variable (e.g. estuarine) and high salinity (e.g. open ocean) environments (see Supplementary Information, Fig. 4, Fig. S3). Additional small conductance mechanosensitive channel families (MscS) enabling the rapid outflux of solutes in response to excessive turgor caused by hypo-osmotic shock ^79^ were acquired in CARA and subsequent lineages, while the large conductance MscL was lost in marine lineages adapted to high salinity, as is frequently the case in marine-dwelling Bacteria ^79^ (Table S4).

LEA14-like protein families containing a conserved WHy (water stress and hypersensitive response) domain ^80^ were acquired in LACAOA, in CARA, and repeatedly along AOA’s diversification, ending up with multiple copies in extant genomes (Fig. 4, Fig. S3, Tables S3, S4, Data S1), thus equipping AOA with proteins that act as “molecular shields”, preventing denaturation and inactivation of cellular components under conditions of osmotic imbalance.

#### Gain of common and rare DNA repair systems in mesophilic AOA

Oxidative DNA lesions are hypothesized to be the primary source of DNA damage in oxic habitats and LACAOA was already equipped with the basic components of most DNA repair systems. Interestingly, the bacterial-type UvrABC system common in mesophilic and thermophilic Bacteria ^81^, but not found in the hyperthermophilic AOA ancestors or in non-AOA Thaumarchaeota, was acquired by CAMA, possibly from bacteria or by multiple transfers from Bacteria first to other archaeal clades (methanogens, halophiles, TACK) and then to AOA (Data S1). Genetic exchanges continued to shape the DNA repair arsenal of the different thaumarchaeal lineages (see Fig. 4, Fig. S3 and Supplementary Information). In particular, an impressive inflow of DNA repair families took place into the ancestor of *Nitrososphaerales* with the acquisition of a complete non-homologous end joining (NHEJ) complex from Bacteria (Data S1) while a UvdE endonuclease family involved in the alternate UV damage repair pathway (UVDR) was acquired in *Ca*. Nitrosocosmicus. The former is extremely rare in Archaea ^82^, with a full complex described so far only from *Methanocella paludicola* ^83^.

### 2. Expansion of transcriptional and redox-based metabolic regulation

An intriguing enzyme family exchange took place in CAMA, the last common ancestor of all mesophilic AOA, in which the AcnA/IRP type aconitase, present in *Ca*. Nitrosocaldus genomes, other Archaea, the eukaryotic cytosol and most Bacteria, was exchanged for a mitochondrial-like aconitase (mAcn) present in eukaryotic mitochondria, Bacteroides (as previously observed in ^84^), and only a few other lineages of Bacteria (Spirochaetes, Fibrobacteres, Deltaproteobacteria and Nitrospirae; Fig. 5 and Data S1). The key to this exchange may lie in the differential sensitivity and recovery rate of these Acn types to oxidative inactivation of their Fe-S clusters. While the AcnA/IRP group aconitases are more resistant to oxidative damage and are implicated in oxidative stress responses, their repair is slower as it might require complete cluster disassembly ^85^. On the other hand, the eukaryotic mAcn and bacterial AcnB aconitases, although distantly related, are easily inactivated by ROS, but can be readily reactivated when iron homeostasis is restored ^85,86^. This makes them the enzymes of choice for aerobic respiration in Eukaryotes and some Bacteria as it provides a sophisticated redox regulation mechanism of the central carbon metabolism ^87–89^. Interestingly, two non-AOA Thaumarchaeota genome bins from oceans, likely to be from mesophilic organisms as well, also harbour this mitochondrial-like version of the aconitase, while a replacement has taken place in the moderate thermophile *N. gargensis* (Fig. 5). To our knowledge, this is the first observation of a mitochondrial-type aconitase in Archaea and can be interpreted as a clearly beneficial, perhaps crucial adaptation to an increasingly oxic environment, although this remains to be experimentally validated.

The two basic transcription factors of Archaea (TFB and TBP) which are homologs of the eukaryotic TBP and TFIID are found in extended numbers in AOA (as noted earlier for smaller datasets ^90,91^)(Fig. S3). Our ancestral reconstructions, supported by phylogenetic trees, indicate that LACAOA already encoded multiple TFB copies (probably around three), and extant mesophilic AOA genomes encode 4-11 copies. Interestingly, TBP expansions occurred in CARA with extant genomes encoding 2-5 copies (Table S3, Fig. 4, S3). These expansions enabled multiple potential combinations of TFBs-TBPs, resulting in the emergence of global regulatory networks and rapid physiological adaptation in changing environments, as noted earlier for halophilic Archaea and AOA ^92–94^.

### 3. Interactions with the environment: extracellular structures, cell wall modifications

A variety of enzymes involved in acetamidosugar biosynthesis, EPS production, cell envelope biogenesis and adhesion were gained by every ancestor we reconstructed, with an apparent enhancement of the process in *Nitrososphaerales* (Fig. 4, Tables S3, S4), reflecting their demonstrated ability to form biofilms ^21,95^. Formation of single or multi-species biofilms is an understudied but very successful ecological adaptation in Archaea (reviewed in ^96^), as these structures not only offer protection against environmental stress and nutrient limitation ^97,98^, but can also provide favorable conditions for direct nutrient or electron exchange that facilitate biogeochemical cycling.

Cellular structures play crucial roles in biofilm formation, motility, and in mediating various forms of environmental and cell-cell interactions. Our analysis indicates that at least four different types of archaeal Type IV pili (T4P) have been gained along AOA diversification (Fig. 6 and Supplementary Information), and were sometimes subsequently lost, leading to a complex distribution pattern (Fig. S4). The AOA ancestor already possessed at least one T4P, as indicated by the families inferred to be present and acquired (T4P biosynthesis and chemotaxis genes). AOA possess the archaeal flagellum (“archaellum”) ^99^, a pilus of unknown function related to Ups and Bas pili (involved in cell-cell contact and DNA repair and sugar metabolism respectively in Sulfolobales) ^100,101^, as well as two other adhesion-like pili that could play a role in biofilm formation or biotic interactions (Fig. 6) ^12,102^. Interestingly, *Ca*. Nitrosocosmicus spp. did not harbor any T4P, unlike the soil-associated sister-group *Nitrososphaera*, suggesting a different adaptive strategy to terrestrial environments, as pointed out by a number of differential gains/losses in the two lineages (see above). The precise functions of these four pili remain to be elucidated but their abundance and diversity indicate frequent and varied environmental interactions and exchanges across AOA. Such level of diversity in the pili repertoire of AOA even exceeds the diversity observed in Sulfolobales, and is only paralleled by the diversity we find here in Bathyarchaeota (Fig. 6).

**Figure 6.**
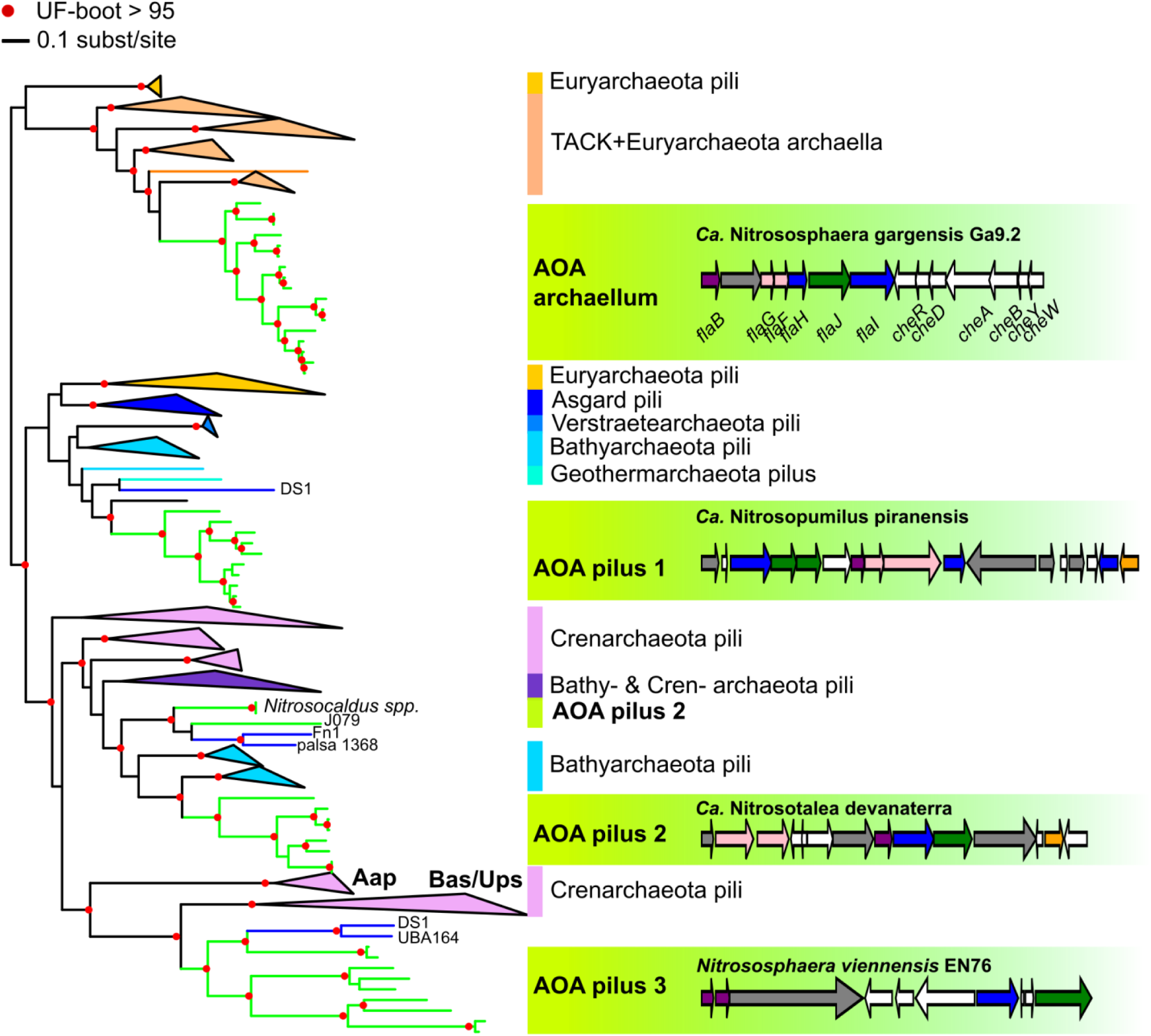
Phylogenetic tree of the Type IV pili ATPase family and genetic structure of AOA pili. A maximum-likelihood phylogenetic tree was obtained for the T4P ATPase in AOA (FAM001536) and representative genomes of other Archaea. Four distinct types of T4P are found in AOA (green branches). Clades other than AOA and non-AOA Thaumarchaeota (blue branches) were collapsed, and colored by taxonomy. Annotations are based on the position of experimentally validated pili (archaeal flagella/archaella, Aap, Ups and Bas pili), otherwise, the generic term “pilus” was used. The genetic architecture of the different types of T4P found in AOA is displayed in green boxes in front of their corresponding position in the ATPase tree. Branches with UF-Boot support above 95% are indicated by a red circle. See also Fig. S4 for the precise genomic distribution within AOA of these different pili.

## General Discussion and Conclusion

Our analysis highlights the many obstacles that AOA might have endured in the course of their diversification and adaptation from hot to moderate temperature environments. Overall, a continuous acquisition of crucial components, many of which are implicated in coping with stress derived from oxygen exposure, seem to have been the key to the ecological success in the different lineages of AOA. We assume that the higher solubility of oxygen at lower temperatures (which is about 4-fold when shifting from 80° to 20°C), together with a metabolic optimization resulting in higher production of ROS and possibly reactive nitrogen species through ammonia oxidation, might have necessitated these extensive adaptations. In summary, we observe a higher proportion of families implicated in oxygen tolerance compared to thermophilic sister lineages in our dataset (Aigarchaeota, Crenarchaeota and some non-AOA Thaumarchaeota) (Fig. S3, Table S4). In addition, moderate habitats are discontinuous and unstable in terms of nutrient availability, hydration content and radiation exposure, in drastically different ways than a hot spring. This is reflected in regulatory adaptations in non-thermophilic AOA, including the acquisition of the regulatable mitochondrial-type aconitase in the common mesophilic ancestor (CAMA), that is found in every mesophilic AOA to date, as well as a considerable expansion of basic transcription factors in all AOA lineages. Thirdly, AOA had to cope increasingly with competition at lower temperatures resulting from higher overall microbial diversity and abundance, which might have triggered their diversification of the outer cell wall and the acquisition of different kinds of type IV pili in most lineages.

The specific differences in adaptations along the evolutionary lineages of marine AOA (*Ca*. Nitrosopumilales) and the two divergently evolved terrestrial lineages (*Ca*. Nitrosocosmicus) and *Nitrososphaera*) reflect their current major ecological distribution. While the ancestors of *Ca*. Nitrosopumilales acquired strategies to cope with higher osmotic pressures and lower temperatures in the ocean, the ancestor of the terrestrial group (*Nitrososphaerales*) acquired a range of families to cope with oxygen, fluctuating nutrient availability and other stressors as well as DNA damage, while it diversified its cell wall modifications and EPS forming capacities. The path into moderate environments of AOA, looks similar to that of halophilic Archaea, where continuous acquisitions of bacterial genes (to Haloarchaea) seem to have contributed to a metabolic switch from anaerobic methanogenic autotrophs to heterotrophy and aerobic respiration, as well as to osmotic adaptations ^93,103–105^. These gradual adaptations contradict the earlier idea of massive lateral gene transfers at the onset of major archaeal clades ^93,103–105^(see also below). Further, we see clear parallels between halophilic Archaea and AOA with respect to their stress adaptations, especially regarding oxidative stress (this study, and ^93,103,106^) and both lineages also have expanded sets of basic transcription factors indicating a possibility to build complex regulatory networks ^94^.

Besides Thaumarchaeota and Halobacteria, Ca. Posidoniales (formerly Marine Group II Archaea) and Class II Methanogens (Methanomicrobia) thrive in oxic habitats ^42,107,108^. Although strictly anaerobic, the latter show particular adaptations in the methanogenesis pathway that result in decreased ROS production together with an enrichment in protein families involved in ROS detoxification and DNA/protein repair ^108^, i.e. similar to adaptations outlined here for AOA, but to a smaller extent.

While we agree that oxygen’s exposure was a driving evolutionary pressure, we here depict a more complex scenario for adaptation of AOA to moderate environments than described in a recent study ^29^. We show that adaptation to higher levels of oxygen occurred in addition to adaptations to various types of stress, and that these gradually occurred involving both innovations, but also pre-existing ancestral sets of genes. In contrast to Ren *et al*. ^29^ we infer an already aerobic ancestor for all AOA which is in agreement with sister lineages to AOA being at least facultative aerobes (Bathy-, Aig-and non-AOA Thaumarchaeota). This is also in line with a more recent diversification of AOA than suggested by the use of molecular clock in Ren et al., 2019 ^29^. The diversification of mesophilic AOA (CAMA) has indeed been estimated to be not older than 950 My based on the acquisition of the fused version of DnaJ-Ferredoxin genes from the ancestor of Viridiplantae ^109^, an acquisition which we could confirm with our comprehensive genomic dataset (DnaJ-Fd on Fig. 4). LACAOA, and at least major mesophilic lineages of AOA might thus have diversified well after the Great Oxidation Event 2.3 Gy ago.

Although the phylogenetic trees we obtained could support Count scenarios and clarify at what point a given family was acquired, in many cases they could not help to identify the donor lineage. This is consistent with another recent study ^29^, and this inability to precisely identify donors persists in spite of the huge increase in taxon sampling since the first studies of lateral gene transfers in Thaumarchaeota ^25,110,111^. It may not only be caused by a limited taxon sampling, but could also result from shifts in evolutionary rates of the acquired families, and the resulting difficulties in solving the gene phylogenies. It is, however, clear that a mix of pre-adaptations (vertical inheritance), and transfers of both bacterial and archaeal genes altogether fell into place to allow the successful radiation of ammonia oxidizing archaea into so many different environments. The complexity of this “genetic cocktail” could explain why the ecological success as seen for AOA appears unique in the domain Archaea.

## Materials & methods

### Genome dataset

We gathered complete genomes of AOA, and enriched the dataset with complete or metagenome-/single-cell-assembled genomes of high quality. We used CheckM (version 1.0.7) ^112^ to assess the completeness and contamination of the genomes gathered (“lineage_wf” parameter), and used a >80% completeness, <5% contamination threshold to include MAGs or SAGs. Yet these criteria were sometimes relaxed in order to broaden the phylogenetic diversity included (see Table S1). In total, we selected 76 genomes for the analysis, including 39 AOA, 13 non-AOA Thaumarchaeota, 8 Aigarchaeota, 5 Bathyarchaeota and 11 Crenarchaeota (Table S1). When no annotations were available for the selected genomes, the Prokka suite (v1.12)^113^ was used to annotate genes and proteins.

### Protein families construction

Protein families were built for the set of genomes selected. A Blast “all sequences against all” was performed, and the sequences representing hits with an e-value below 10^−4^ were used as input for the Silix (v1.2.11) and Hifix (v1.0.6) programs to cluster the sets of similar sequences into families ^114,115^. Two sequences were clustered in a same Silix family when they shared at least 35% of identity and their blast alignment covered at least 70% of the two sequences lengths. The Hifix program was then used on the Silix families for refinement. We obtained 37517 families, of which 12367 had more than one sequence.

### Phylogenomic tree inference

Universal archaeal families described in ^116^ were annotated in our genome dataset using HMMER (v3.1b2) ^117^ as described in ^12^. The best hit was selected in each genome, and a subset of 33 informational protein families that were present in 74 out of 76 genomes of our dataset was selected. The sequences from each family were extracted and aligned using MAFFT (linsi, v7.313) ^118^, and the alignments filtered using BMGE (BLOSUM30, v1.12) ^119^. The filtered alignments were then concatenated, and a maximum-likelihood tree was inferred with IQ-Tree (v1.6.11) using the LG+C60+F model of sequence evolution, and 1000 Ultrafast bootstraps were performed ^120^.

The 33 families were the following: PF00177 Ribosomal protein S7p/S5e, PF00189 Ribosomal protein S3, PF00203 Ribosomal protein S19, PF00237 Ribosomal protein L22p/L17e, PF00238 Ribosomal protein L14p/L23e, PF00252 Ribosomal protein L16p/L10e, PF00276 Ribosomal protein L23, PF00347 Ribosomal protein L6, PF00366 Ribosomal protein S17, PF00380 Ribosomal protein S9/S16, PF00410 Ribosomal protein S8, PF00411 Ribosomal protein S11, PF00416 Ribosomal protein S13/S18, PF00466 Ribosomal protein L10, PF00572 Ribosomal protein L13, PF00573 Ribosomal protein L4/L1 family, PF00673 ribosomal L5P family, PF00687 Ribosomal protein L1p/L10e family, PF00831 Ribosomal L29 protein, PF01090 Ribosomal protein S19e, PF01157 Ribosomal protein L21e PF01200 Ribosomal protein S28e, PF01201 Ribosomal protein S8e, PF01280 Ribosomal protein L19e, PF01667 Ribosomal protein S27, PF01000 RNA polymerase Rpb3/RpoA insert domain, PF01191 RNA polymerase Rpb5, C-terminal domain, PF01192 RNA polymerase Rpb6, PF01912 eIF-6 family, PF09173 Initiation factor eIF2 gamma, PF03439 Early transcription elongation factor of RNA pol II, PF04406 Type IIB DNA topoisomerase, PF11987 Translation-initiation factor 2.

### Inference of ancestral Optimal Growth Temperature

A dataset of 16S rRNA sequences and corresponding OGT was extracted from ^121^ and from the literature (see Table S2). The stem position predictions were obtained from the RNAStrand database (version 2.0) for all the sequences in our dataset that were available (20 rRNA molecules). The 16S rRNA sequences that were available (56 out of 76 genomes, see Table S1) were all aligned together with RNAStrand sequences with SSU-ALIGN (v0.1.1) ^122^, and the consensus of the positions found in stems were conservatively selected to be part of stem regions (878 positions). The GC% of the predicted stems was computed for each sequence. A linear regression between the stem GC% and the OGT was computed. A non-homogeneous model was used in bppml (BppSuite, v2.4.1 ^123^) to estimate the evolutionary model of the 16S rRNA sequences along the reference tree, as in ^31^. The estimated parameters were then used in the program BppAncestor to reconstruct 100 replicates of ancestral sequences along the reference tree. The OGT of the ancestor of extant AOA was inferred using the linear regression coefficients and the bootstrap replicates to estimate a confidence interval, as described in ^31^.

### Count analysis and ancestral genomes reconstruction

A matrix of occurrences of each protein family in extant genomes was created for all the genomes under analysis. This matrix was used as an input for the Count program (v10.04) ^33^ along with the reference phylogeny to compute 1) the rates of gains and losses along the tree independently for each family given a phylogenetic birth-and-death model (gain-loss-duplication model, default parameters), and 2) the posterior probabilities for each family to have been gained and lost given the inferred rates, in each branch of the tree. R (v3.5.1) and Python (v2.7) scripts were used to analyse and extract the resulting data. Evolutionary events inferred on each branch of the reference tree (family gains, losses, expansions: number of family members increase, contractions: number of family members decrease) were selected when their probability was above a fixed threshold of 0.5. This threshold was also used to define the presence and multi-copy state of each family within each node. The sets of families inferred to be present at a given node constituted the ancestral genome of the branches emerging from this node. We therefore could extract the sets of evolutionary events (gains, losses, expansions and contractions) before a lineage diversification, together with the families inferred to have been present in the ancestor of the lineage (Table S3).

### Annotation of protein families and quantification of functional categories

Aside from the annotations provided with the genomes, we obtained annotations using PFAM (version 28.0 PFAM-A, ‘cut-tc’ option, best i-evalue match selected with HMMER) ^124^, TIGRFAM (version 15, ‘cut-tc’ option, best i-evalue match) ^125^ using the corresponding HMM protein profiles and the HMMER program version 3.1b2 ^117^. arCOGs (2014 version) ^126^ and COGs (2014 version) ^127^ were assigned using the COGnitor scripts, with an e-value threshold set to 10^−10^. The MacSyFinder program (v1.0.5) ^128^ was also used to annotate type IV pili in genomes, using models and HMM profiles described earlier ^12^. Manual inspection of the protein families resulted in grouping into custom functional categories, which were then quantified in extant genomes as in Table S4.

### Phylogenetic analyses of gained genes

We first gathered a representative dataset of 281 genomes from Bacteria, Archaea and Eukaryotes (see Table S1). This dataset, composed of complete genomes (Refseq database, NCBI), was selected to cover a vast range of organismal diversity, and was enriched to cover the diversity of organisms from the TACK super-phylum by adding 68 highly complete metagenomic bins (Table S1).

For protein families of interest, we retrieved all the sequences classed as part of the family in our 76 genomes dataset, and used them to query the representative dataset of genomes using Blast (v2.6.0+) ^129^. The first 250 sequences corresponding to the best-score hits with an e-value lower than 10^−10^ were selected. The corresponding sequences were extracted, dereplicated with Uclust (’-cluster_fast’, 100% identity level, Usearch v10.0.240)^130^, and aligned to the sequences from the original protein family using the MAFFT program (’linsi’ algorithm, version 7.397)^118^. The alignments were then filtered using the BMGE program (BLOSUM30)^119^. The filtered alignments were used to build phylogenetic trees by maximum-likelihood (IQ-tree, “-m TESTNEW”, best evolutionary model selected by Bayesian Information Criterion, 1000 UF-Bootstrap and aLRT replicates)^120^. Upon manual inspection of the trees, a few sequences were excluded during protein family phylogenies’ reconstructions, when the corresponding branches had a length in the tree > 1 substitutions/site. In those few cases, alignments and trees were re-constructed without the corresponding sequences. All resulting trees are provided in Data S1, and a list of the generated trees can be found in Table S5.

### Generation of figures

Drawings of trees and genes were generated with iTOl (v4) ^131^ and GeneSpy (v1.1) ^132^ respectively.

## Data availability

The genomes analyzed in this study are publicly available on NCBI or IMG/JGI, and all accession numbers are given in Table S1. All data generated in this study are provided in the Supplementary Tables 2-5 and generated phylogenetic trees in Supplementary Data S1.

## Acknowledgements

This project was supported by ERC Advanced Grant TACKLE (695192) and project P27017 of the Austrian Science Fund (FWF). SSA was funded by a “Marie-Curie Action” fellowship, grant number THAUMECOPHYL 701981. We are thankful to Thomas Rattei and Florian Goldenberg for providing access to the CUBE servers (U. Vienna), to Céline Brochier-Armanet for early discussions on the project, and to Simon Rittmann and Lisa-Maria Mauerhofer for providing optimal growth temperatures of methanogens.

## Author contributions

SSA and CS conceived the study. SSA performed the bioinformatic and phylogenetic/phylogenomic analyses. MK analysed the reconstructed genomes. All authors analysed and discussed results and wrote the manuscript.

