## Supplementary Information for "Ancestral reconstructions decipher major adaptations of ammonia oxidizing archaea upon radiation into moderate terrestrial and marine environments"

**Sophie S. Abby[1,2,+], Melina Kerou[1,+], Christa Schleper[1,*]**
[1] University of Vienna, Archaea Biology and Ecogenomics Division, Dep. of Ecogenomics and Systems Biology, Althanstrasse 14, 1090 Vienna, Austria.
[2] University Grenoble Alpes, CNRS, Grenoble INP, TIMC-IMAG, 38000 Grenoble, France.

[+] these authors contributed equally

*Calibration of a molecular thermometer*

In order to explore if the last common ancestor of AOA was a mesophilic, thermophilic or hyperthermophilic organism, we compiled a dataset containing the stem G+C content of 16S rRNA and OGT for 68 archaeal organisms (Table S2 see Methods) and obtained a highly significant correlation between the stem G+C content and the OGT (Fig. S1, Spearman's P: 0.94, linear regression: adjusted $R^2$=0.88, p-value < 2.2e-16). We checked if the correlation was still significant when taking into account potential phylogenetic inertia using the phylogenetic independent contrasts approach [1]. We thus built a ML tree (IQ-Tree, best model) for the 16S rRNA sequences used for the molecular thermometer calibration (68 sequences in Table S2), and used it to correct for phylogenetic dependencies for both GC and OGT variables among lineages (use of the Cran R package “ape” [2]. We obtained a significant correlation between stem G+C content and OGT, showing our molecular thermometer was robust to phylogenetic inertia and could be robustly used for predictions (Spearman's P: 0.58, linear regression: adjusted $R^2$=0.33, p-value = 1.73e-06). We then reconstructed 100 replicates of ancestral sequences for each ancestor (represented by internal nodes) in our reference phylogeny using a non-homogeneous model of sequence evolution as in [3] and inferred ancestral OGT with confidence intervals (see Materials and Methods).

*Gene gains’ rates plead for gradual acquisitions of new genes along AOA diversification*

We computed the family gain rate as the number of gains inferred divided by the length of the corresponding branch (excluding very short branches <0.005 substitutions per site and incomplete genomes – which gains and losses are biased) for the subtree of Thaumarchaeota. This resulted in a number of gains per substitutions per site. We could observe that the rate was found to be much higher for leaves/contemporary genomes than for deeper branches/ancestral genomes (Fig. S2, Wilcoxon rank sum test, p-value << 0.001). This was expected, as contemporary genomes are known to harbor specific, recently acquired genes (variable part of the genome) that are not destined to be fixed over long evolutionary times, and that by definition, we cannot reconstruct this variable part of the genome for ancestors.

We then focused on the tempo of gains for ancestral genomes. We could find a slightly significant positive correlation between the number of gains and the length of the corresponding branch (Pearson's correlation coefficients: 0.45, p-val ~ 0.004), arguing for an overall constant gain rate along AOA diversification. This trend was more visible (Fig. S2, Pearson's correlation coefficients: 0.64, p-val << 0.001. Linear regressions: adjusted $R_2$ = 0.39, p-val << 0.001), when the number of gains was normalized by the genome size (number of families inferred to be present), and thus computed as a genome portion increase. Such a trend was not visible for losses. We cannot however rule out that in some cases, there might have been e.g. a simultaneous increase for both substitutions and gains' rates, which would bias the ratio and give for instance the false impression of relatively low (high) gain rates in the context of an increased (decreased) substitution rate. Yet, some ancestral branches harbored higher gain rates, such as the one corresponding to the last common ancestor of AOA (Wilcoxon signed rank test, p-value ~ 0.001).

*Exploration of gene gains indicates a flux of functional bricks*

Despite a rather continuous influx of genes, the evolutionary scenarios we obtained were very often suggestive of "en bloc" acquisitions of gene families being part of a same function: the corresponding families were inferred to be acquired by a same ancestor. We can cite as examples the acquisition of the AMO gene cluster, urea-utilization gene set, Uvr genes (*uvrABC*), genes involved in chemotaxis and vitamins (B7, B12) synthesis... Moreover, in several cases, these families were found next to each other in contemporary genomes, reinforcing the idea of a single acquisition event. These observations demonstrated the relevance of the evolutionary scenarios inferred by the Count program. However, phylogenetic reconstructions could prove more powerful in order to study evolutionary scenarios in details, as the patterns of presence/absence and gene family sizes used by Count for its inferences can result from different types of evolutionary events. In order to make sense of the inferred gene gains and losses and make assumptions on their physiological relevance, evolutionary scenarios proposed by Count were therefore refined by comprehensive phylogenetic reconstructions for approximately 80 crucial families discussed in the rest of this manuscript (see Table S5). This conservative approach enabled to correct if needed the evolutionary scenarios inferred by the program Count, although in most cases, both coincided well, emphasizing their relevance. Most of the scenarios we manually rectified were linked to a stringent delineation of gene families: sometimes, homologous sequences were placed in distinct families with our sequence clustering criteria, and therefore not considered as a whole family for the evolutionary reconstruction, making the Count program invoke unnecessary evolutionary events.

Beyond confirming the precise ancestor which acquired the gene families of interest, these phylogenies enabled the investigation of their origins (see Methods, Data S1 and Table S5). Even though in most cases it was difficult from our phylogenies to point out a putative donor clade for the genes gained, we could sometimes make assumption on the donating lineage. Some cases are discussed along main text.

*On the origins of acquired genes*

Because Count infers evolutionary scenarios solely based on presence-absence patterns with no direct link to a gene evolutionary history, we queried a large database of archaeal, bacterial and eukaryotic genomes, and computed phylogenetic trees for ~80 families of interest (Data S1). This allowed to obtain more resolution on these families' and AOA's evolutionary history. Trees were inspected manually, and allowed 1) to confirm or refine the position in the species tree were these particular families were acquired – e.g., in the case of split families in AOA due to high sequence divergence and 2) suggest an origin for the family, when evidence of lateral gene transfers could be found.

*Additional notes on the gene losses occurring on the way to LACAOA*

As we can only account for families that are found in extant genomes for ancestral genome reconstruction, the fraction of specific genes to a genome cannot be reconstructed. It is thus very likely that the LACAOA genome and ancestral AOA genomes had more different families than inferred.

As mentioned, a loss of protein families associated with organic carbon assimilation occurred in LACAOA (Fig. 3 and Tables S3, S4), some of which, such as the OFOR, can also be viewed as adaptations to an increasingly oxygenated environment. Generally OFORs are extremely oxygen-sensitive [4], owing to the inactivation of three [4Fe-4S] clusters, two of which are located on subunit delta (corresponding to an intra-molecular ferredoxin domain, or domain V). Interestingly, the $(ab)_2$-type OFOR retained in LACAOA and all extant AOA and only found in aerobic organisms, contains a sole [4Fe-4S] cluster due to the absence of domain V and thereby is a more oxygen-tolerant enzyme [5–7]. The thaumarchaeal homologs (for example, NVIE_029480 and NVIE_029490) bear a 46.5% and 59% identity to the a- and b-subunits respectively of the broad substrate specificity OFOR from *Sulfolobus tokodaii,* which can utilize pyruvate, 2-oxoglutarate and 2-oxobutyrate [5](Yan et al. 2016). This suggests that this OFOR can play multiple roles in glyconeogenesis, TCA and aminoacid metabolism. Additionally, families corresponding to two subunits of a 2-oxoacid dehydrogenase complex (OADHC) were lost, including genes responsible for the synthesis of the lipoic acid cofactor. The substrates of OADHC complexes in archaea range from acetoin [8] to branched-chain 2-oxoacids resulting from aminoacid catabolism [9].

Losses and contractions were observed in other families involved in aminoacid metabolism, such as the glycine cleavage system, alanine dehydrogenase, carbamate kinase, glutamate synthase and others (see Tables S3, S4). The loss of the crenarchaeal version of the GTP-dependent phosphoenolpyruvate carboxykinase is accompanied by the gain of an ATP-dependent version with high similarity to Firmicutes, however it is unclear whether there is an energetic or other advantage to this exchange.

The genomic inventory for the biosynthesis of all amino acids seems to be present in LACAOA, which gained additionally a putative 5, 10-methylenetetrahydrofolate reductase, enabling methionine salvage from homocysteine.

*Oxidative stress: distribution of detoxification pathways*

The system of periplasmic and membrane-bound thioredoxin superfamily oxidoreductases (DsbA, 76G-FAM004903_0, and DsbD, 76G-FAM002351, respectively), responsible for the formation and repair of disulfide bonds in periplasmic and cell membrane proteins [10–13] was present in LACAOA and the family of soluble periplasmic DsbA-like thiol-disulfide oxidoreductases further expanded in CAMA resulting in up to 11 paralogs in extant AOA genomes (Table S4). While the functional equivalency with either DsbA, DsbC or DsbG in bacteria cannot be established, the essential redox-active CXXC motif is conserved in all thaumarchaeal homologs, which are furthermore predicted to be extracellular. Families of DsbD homologs, the key transmembrane electron hubs of the Dsb system that transfer electrons from cytoplasmic thioredoxins to the various oxidoreductases in the periplasm [10], were also acquired in CAMA and the ancestor of Nitrosopumilales.

Two families of cysteine desulfurases (Csd/Group II and IscS/Group I), acquired by ancestral non-AOA Thaumarchaeota and LACAOA, can further alleviate ROS damage by supplying sulphur from cysteine for the synthesis and repair of [Fe-S] clusters, in addition to the already present SUF system for Fe-S cluster assembly. This ability increases chances of survival upon exposure to environments with lower bioavailability of inorganic sulphur, and therefore facilitates their radiation away from restricted environments, such as hot-springs [14].

*Redox homeostasis*

WrbA flavodoxins (type IV NQOs) catalyze an NADH/NADPH dependent 2-electron reduction of endogenous quinones and thereby prevent the formation of semiquinones, which react with molecular oxygen and produce superoxide [15]. Multiple families were gained throughout the evolution of AOA: in CARA (76G-FAM002176), the ancestor of Nitrososphaerales (76G-FAM010188, 76G-FAM008623, 76G-FAM009189, 76G-FAM005030), the ancestor of Nitrosopumilales (76G-FAM008145), and then mostly in the terrestrial lineages (Fig. 4 and Table S4). CARA gained a WrbA family bearing a high similarity (45-54%) to characterized quinone reductases from *Pseudomonas aeruginosa* and *P. putida*.

Even though the thaumarchaeal ferredoxin:NADP(H) oxidoreductase homologs exhibit low identity to characterized bacterial FNRs (~27% to *E. coli* FNR), the necessary FAD and NAD(P) binding domains are conserved. These flavoenzymes catalyze the reversible electron transfer between the two-electron carrier NADPH and one-electron carriers such as ferredoxins or flavodoxins, a consequence of which is the depletion of cellular NADPH, which in conditions of oxidative stress can rapidly lead to HO• production through Fenton-type chemistry [16,17].

*Additional notes on the evolution of a copper-based metabolism*

A family of two-domain multicopper oxidases (MCO1 in [18], 76G-FAM003386) together with an adjacent ZIP family permease (76G-FAM007716) were gained by LACAOA and are present in almost all extant genomes with the exception of *N. devanaterra*. They are proposed to have an accessory role in nitrite reduction [19], or copper metabolism [18]. Contemporary AOA genomes contain 3-4 distinct membrane-associated or secreted MCOs, indicating an important role for this enzyme family. Additionally, a number of small, blue-copper domain protein families, similar to plastocyanins, were gained in LACAOA to serve as electron carriers. Interestingly, a nitrite reductase (NirK) family protein is not inferred to be present in LACAOA, raising the question of the source of nitric oxide shown to be essential for the archaeal (and bacterial) ammonia oxidation pathway [20,21]. This protein family was inferred as acquired by CAMA (76G-FAM008287), however this could be an example of the limitations of our dataset and of the method used to reconstruct ancestral genomes.

The analysis of other, more diverse lineages of AOA, could eventually enable to make more robust inferences on the genetic repertoire of LACAOA.

Copper detoxification systems in LACAOA consist of the already present copper-exporting P-type ATPase CopA (76G-FAM001010) [22] and the archaeal-specific CopT-like metal-responsive regulator (76G-FAM000666) [23]. Additionally, families of disulfide bond oxidoreductase family proteins (DsbA type, present in LACAOA), can alleviate the consequences of copper-induced protein misfolding in the pseudoperiplasm, as observed in *E.coli* [24]. CopA was lost in the ancestors of the predominantly marine genera Nitrosopelagicus, Cenarchaeum, Nitrosopumilus and Nitrosoarchaeum, in line with the fact that copper has a rather low bioavailability in many marine environments [25].

*T4P in AOA*

We annotated in AOA genomes the different T4P genes from the arCOGs listed by Makarova and colleagues [26] using the HMM protein profiles derived from the arCOG sequences, and a model for T4P provided to the MacSyFinder program [27]. We thus could compare the genetic architecture of the different T4P loci (Fig. S4). The tree of T4P ATPases we built shows four distinct monophyletic groups of pili in AOA. This suggests that these pili might have been acquired by LACAOA and a non-AOA ancestor for some pili, and/or that these pili have been exchanged by lateral gene transfer between AOA. This tree also shows clearly distinct types of T4P, with potentially different functions.

Based on the set of T4P-related families inferred to have been present in LACAOA, we can infer that LACAOA possessed a flagellum. Among the seven T4P families, five are indeed flagellum-specific in extant AOA, while two other families are shared by several pili (including the ATPase, that we used to produce the T4P overview tree). These seven families corresponded to the following arCOGs: arCOG01817, arCOG04148, arCOG01829, arCOG01809, arCOG01824, arCOG01822 and arCOG02298. LACAOA's T4P being a flagellum is also consistent with a parsimony scenario of the acquisition of AOA's flagella in LACAOA followed by vertical transmission and independent losses, because the flagellum phylogeny for AOA is consistent with the species tree.

*The case of $F_{420}$*

The key enzymes for $F_{420}$ biosynthesis CofD (2-phospho-L-lactate transferase, 76G-FAM000786) and CofE ($F_{420}$:γ-L-glutamyl ligase, 76G-FAM000546_0, a multicopy family), were acquired by the ancestor of Aigarchaeota and Thaumarchaeota, and the ancestor of Bathyarchaeota, Aigarchaeota and Thaumarchaeota respectively suggesting a conserved, but as yet elusive, role for this cofactor in these two lineages. In its reduced form, $F_{420}H_2$, a 5-deazaflavin, can reduce via hydride transfer a broad range of organic molecules such as alcohols, alkenes, alkynes, imines and carbon double bonds in the context of catabolic, biosynthetic and detoxification pathways. While an essential electron carrier in the central energy metabolism of methanogenic, methanotrophic and sulphate-reducing archaea, there is a growing appreciation of the widespread occurrence and diverse ecosystem-shaping roles of this cofactor in aerobic bacterial lineages abundant in soil and aquatic ecosystems, such as Actinobacteria, Proteobacteria, Chloroflexi and Firmicutes, and the significant over-representation of $F_{420}$ biosynthetic clusters in aerobic versus anaerobic environments [28,29].

Most studies have focused on the soil-dwelling and pathogenic members of Actinobacteria (mainly mycobacteria and streptomycetes), where loss of the ability to synthesize $F_{420}$ results in hypersensitivity to oxidative, nitrosative and antimicrobial stress. The reduced cofactor is produced via $F_{420}$-dependent glucose 6-phosphate dehydrogenase. Through the action of $F_{420}H_2$-dependent reductases they are able to maintain redox homeostasis, prevent the production of ROS/RNS, maintain the impermeable structure of their cell wall, reductively detoxify ROS/RNS and a variety of toxic compounds of fungal and plant metabolism and xenobiotics such as coumarins, nitroaromatic compounds and polycyclic aromatic hydrocarbons and participate in the biosynthesis of antibiotics [30,31].

These processes are mediated through an extended repertoire of $F_{420}$-dependent enzymes belonging to two superfamilies: the flavin/deazaflavin oxidoreductases (FDOR) and the Luciferase-like hydride transferases (LLHT), both of which were continuously acquired during the evolution of AOA and also present in LACAOA (LLHT). FDORs comprise a highly diverse superfamily with extremely interesting representatives such as the anti-oxidant $F_{420}H_2$-quinone reductases (Fqrs) of mycobacteria, which, by catalyzing a 2-electron reduction of endogenous quinones prevent the formation of semiquinones, which react with molecular oxygen and produce superoxide [32,33]. Additional protection is conferred against nitrosative stress, occurring due to the formation of peroxynitrite by the reaction of superoxide with NO [32,34]. Moreover, reduced $F_{420}$ has itself a protective role against reactive nitrogen species, as it can reduce $NO_2$, a potent oxidant arising from the spontaeous reaction of NO with $O_2$, back to NO [35]. This would confer a double advantage to Thaumarchaeota, as it would simultaneously supplement the NO pool necessary in the second step of ammonia oxidation. Families encoding putative Fqr homologs were gained by LACAOA (76G-FAM007606), and the ancestor of Nitrososphaerales (76G-FAM001715, 76G-FAM009811; Tables S3, S4). Other Thaumarchaeal families comprise putative FDOR enzymes that use other flavin cofactors, and their functions remain unclear.

A continuous expansion in the repertoire of the $F_{420}$-dependent members of the LLHT superfamily occurred in the course of Thaumarchaeota evolution with two families present and one gained in LACAOA (76G-FAM001132, 76G-FAM001111 and 76G-FAM000949 respectively), two additional families gained in the Nitrososphaerales ancestor (76G-FAM003798, 76G-FAM010258) and two more families in the ancestor of Nitrosocosmicus (76G-FAM010783, 76G-FAM011018) (Tables S3, S4). After certain family losses, distribution in extant genomes involves 2 homologs in the order Nitrosopumilales, 3 in Nitrosotaleales, 4-9 in Nitrososphaerales and 3 in Nitrosocaldales, implying an expansion of this enzymatic family in the terrestrial lineages. This enzyme family includes well characterized members with central metabolic roles such as methylenetetrahydromethanopterin reductase (Mer) and $F_{420}$-reducing secondary alcohol dehydrogenases (Adf) from archaea, and $F_{420}$-reducing glucose-6-phosphate dehydrogenase (Fgd) from mycobacteria [31](Greening et al. 2016. It has also been implicated in cell wall modifications essential for mycobacterial survival in the host environment, antibiotics biosynthesis in streptomycetes and degradation of picrate and nitroaromatic compounds by rhodococci, where they are often found in operons [31]. However, this family is also very diverse and the function of the Thaumarchaeal homologs remains elusive, although all homologs contain the conserved residues conferring selectivity for this cofactor.

In the absence of other candidates, generation of reduced $F_{420}$ is likely mediated by a putative $F_{420}H_2$:NADP oxidoreductase (Fno, 76G-FAM003693), a conserved reversible enzyme that either reduces NADP (in methanogens), or transfers electrons from NADP to $F_{420}$ (in bacteria). The ancestor of Aigarchaeota and Thaumarchaeota acquired this family together with the $F_{420}$ biosynthesis genes as indicated by recent phylogenetic studies [29], supporting a conserved mechanism. Interestingly, genes from the LLHT family gained by the ancestor of Nitrososphaerales (76G-FAM000682) share 35-38% identity with $F_{420}$-dependent glucose-6-phosphate dehydrogenases (Fgd) from mycobacteria[36], which serve as the main $F_{420}$ reductants in these organisms. This was already noted by [37], and introduces the possibility of a putative link between central carbon metabolism of Thaumarchaea and production of reduced $F_{420}$.

Persistence is particularly important for soil lineages where they likely encounter severe resource limitation and occasional hypoxia, an observation supported by the distribution of these enzymes in AOA genomes. $F_{420}$ has also been implicated in recovery from hypoxia-induced dormancy, a state in which soil populations are often found[33].

*Xenobiotics degradation*

A number of protein families belonging to the glyoxalase/bleomycin resistance/dioxygenase superfamily were present and additionally gained in LACAOA, CAMA, the ancestor of Nitrososphaerales and in subsequent “soil” lineages [18]. Members of this superfamily are involved in antibiotic resistance and aromatic compounds degradation, traits offering competitive advantages in niche expansion, especially in terrestrial environments. Glyoxalase I (Glo1) is an especially interesting member of this superfamily in AOA, as together with glyoxalase II (Glo2), a family present in LACAOA and additionally gained in CAMA, they may convert methylglyoxal to D-lactate, a precursor of $F_{420}$ biosynthesis [38,39].

*Notes on the mobility of DNA repair systems*

LACAOA was equipped with the basic components of nucleotide excision repair (NER), base excision repair (BER), homologous recombination (HR) machineries, translesion polymerase (PolY family), and a bacterial-type PolX family polymerase, the latter only being acquired by the ancestor of Thaumarchaeaota (Table S4 and Fig. 3).

The NER-associated helicase-nuclease pair XPB-Bax1 [40], and the PolX family polymerase were lost in CARA. Photolyase families [41] responsible for photoreactivation were independently acquired by the ancestors of Nitrosopumilales (76G-FAM001869) and Nitrososphaerales (76G-FAM010262), as well as a putative MutL-like mismatch repair family protein by the latter group (FAM010111). HR recombinase (RadA, RadB), helicase (HerA) and nuclease (NurA) families were gained multiple times, especially as part of the parallel genomic expansion in the ancestors of Nitrososphaera and Nitrosocosmicus (Fig. 4, S3, Tables S3, S4).

### Supplementary Tables

**Table S1.** The 76 genomes used for the evolutionary analysis and the set of 349 representative genomes from the three domains of life used for phylogenetic analyses of acquired families.

**Table S2.** The optimal growth temperature and stem GC content of the 16S rRNA from the 68 archaeal organisms used to calibrate the molecular thermometer.

**Table S3.** Ancestral AOA genomes' content and inferred evolutionary events.

**Table S4.** Annotation tables, and quantification and distribution of the stress-related families acquired along AOA evolution.

**Table S5.** Listing of the trees from families discussed in main text.

### Supplementary Data

**Data S1.** Maximum likelihood trees of stress-related 76 families found in AOA evolutionary analysis. The taxonomy of each sequence is reported on the right. Family names drawn on trees are as found in Tables S3, S4 and are specifically listed in Table S5.

### Supplementary Figures:

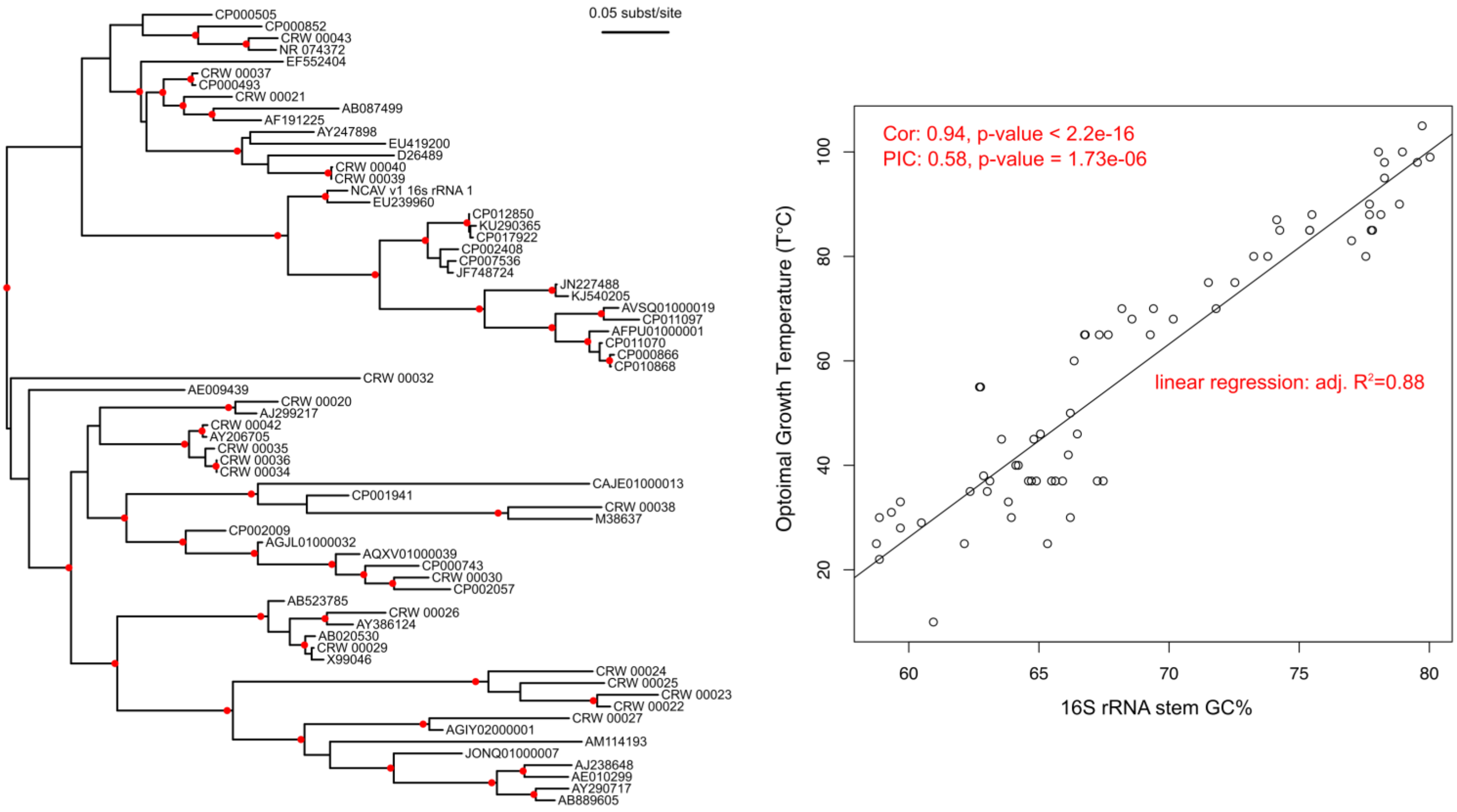


**Figure S1.** Calibration of the molecular thermometer. The maximum likelihood tree of the 16S rRNA tree of the 68 organisms (code in Table S2) is displayed on the left, and the correlation/linear regression between the 16S rRNA stem GC content and the optimal growth temperature of these 68 organisms are displayed on the right. “Cor” stands for “Pearson's correlation coefficient”, ”PIC” for phylogenetic independent contrasts (Pearson's correlation coefficient corrected using the method of contrasts), “adj” stands for “adjusted”. Branches with UF-Boot support above 95% are indicated by a red circle.

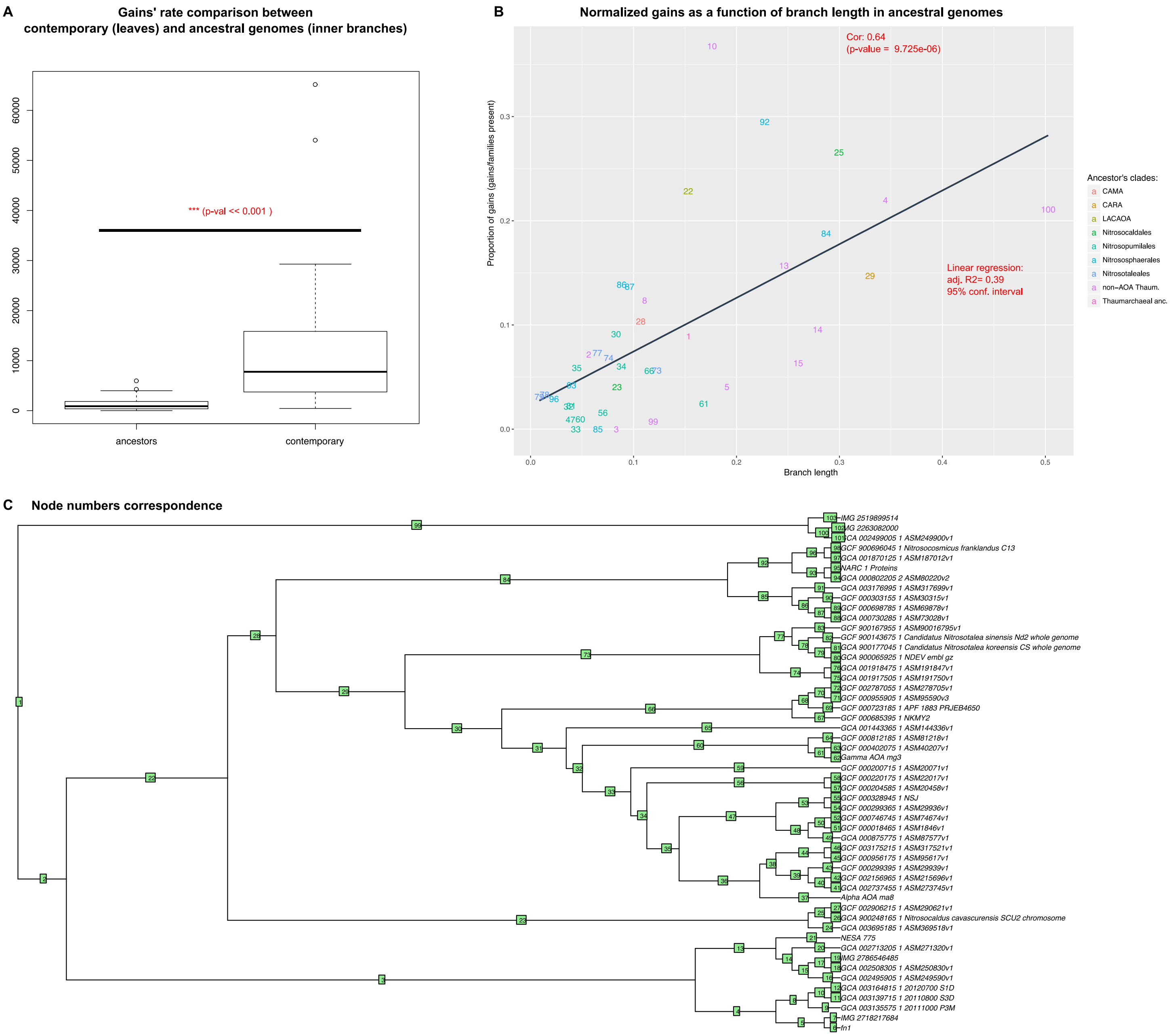


**Figure S2.** **A**. Comparison of gains' rates between contemporary and ancestral genomes (Wilcoxon rank sum test). **B.** Correlation between proportion of gains and number of substitutions per site for Thaumarchaeota's ancestors. "Cor" stands for "Pearson's correlation coefficient", "adj" stands for "adjusted". Nodes are numbered as displayed in **C.** Subtree of Thaumarchaeota with nodes numbered. See Table S1 for genomes' codes.

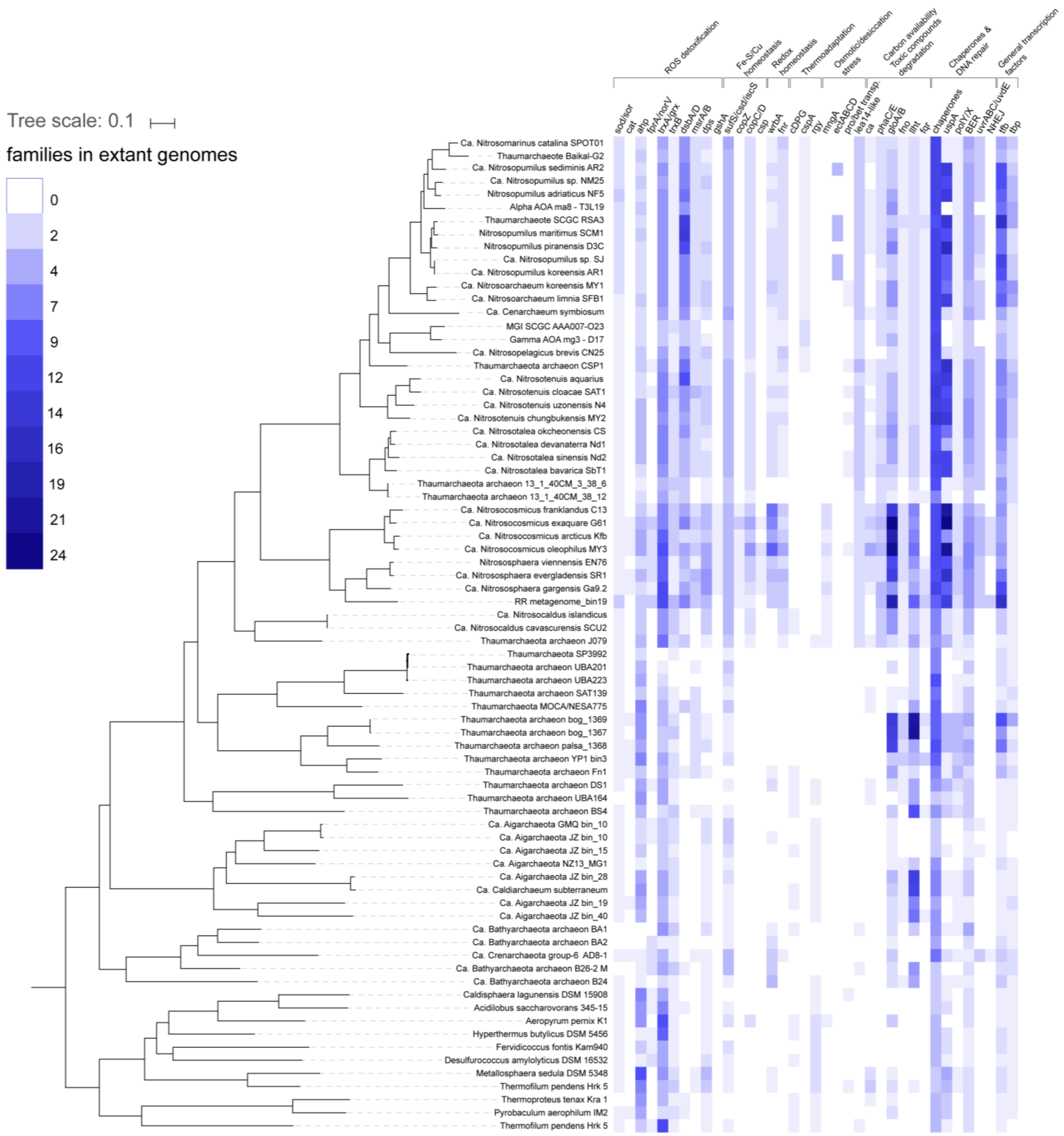


**Figure S3.**

Heatmap of the distribution of genes involved in the main functional categories of ROS detoxification, Fe/Cu/redox homeostasis, thermoadaptation, osmoadaptation, desiccation stress, regulation of carbon availability, toxic compounds degradation, chaperones, DNA repair and general transcpription factors in extant AOA and representatives of Crenarcaeota, Aigarchaeota and Bathyarchaeota used in this study. The phylogenomic tree is the same as Fig. 1, with bootstrap values omitted for clarity. Families in the same functional category were summed up for readability (e.g. sod, superoxide dismutase & sor, superoxide reductase). The number of families per category in each genome are colour-coded as indicated in the legend. The data used to generate this heatmap can be found in Table S4. Abbreviations: cat, catalase; ahp, alkyl hydroperoxide reductase/ Osm-C peroxiredoxin; fprA/norV, flavorubredoxin; trxA/grx, thioredoxin/glutaredoxin; trxB, thioredoxin reductase; dsbA/D, disulfide bond oxidoreductase A/D; msrA/B, methionine sulfoxide reductase A/B; dps, DNA protection during starvation family protein; gshA, glutamate-cysteine ligase; sufS/csd/iscS, cysteine desulfurase families; copZ, copper binding protein; copC/D, copper resistance family proteins; csp, four-helix bundle copper storage protein; wrbA, multimeric flavodoxin WrbA; fnr, flavodoxin reductase (ferredoxin-NADPH reductase); cDPG, cyclic 2, 3-diphosphoglycerate synthetase; cspA, cold-shock protein A; rgy, reverse gyrase; mngA, mannosyl-3-phosphoglycerate synthase; ectABCD, ectoine/hydroxyectoine biosynthesis genes; pro/bet transport, ProP (proline/glycine-betaine):(H+/Na+) symporter; lea14-like, LEA14-like desiccation related protein; ca, carbonic anhydrase; phaC/E, poly(R)-hydroxyalkanoic acid synthase subunits C/E; gloA/B, glyoxalase I/II; llht, luciferase-like hydride transferase; fqr, F420H(2)-dependent quinone reductase; fno, F420H2:NADP oxidoreductase; chaperones, dnaJ/dnaK/grpE/dnaJ-ferredoxin/hsp90/prefoldin/peptidylprolyl isomerase; uspA, universal stress protein family A; polY/X, DNA repair polymerases Y & X; BER, base-excision repair families; uvrABC/uvdE, ultraviolet radiation repair excinuclease ABC system/UV damage endonuclease uvdE; NHEJ, non-homologous end joining.

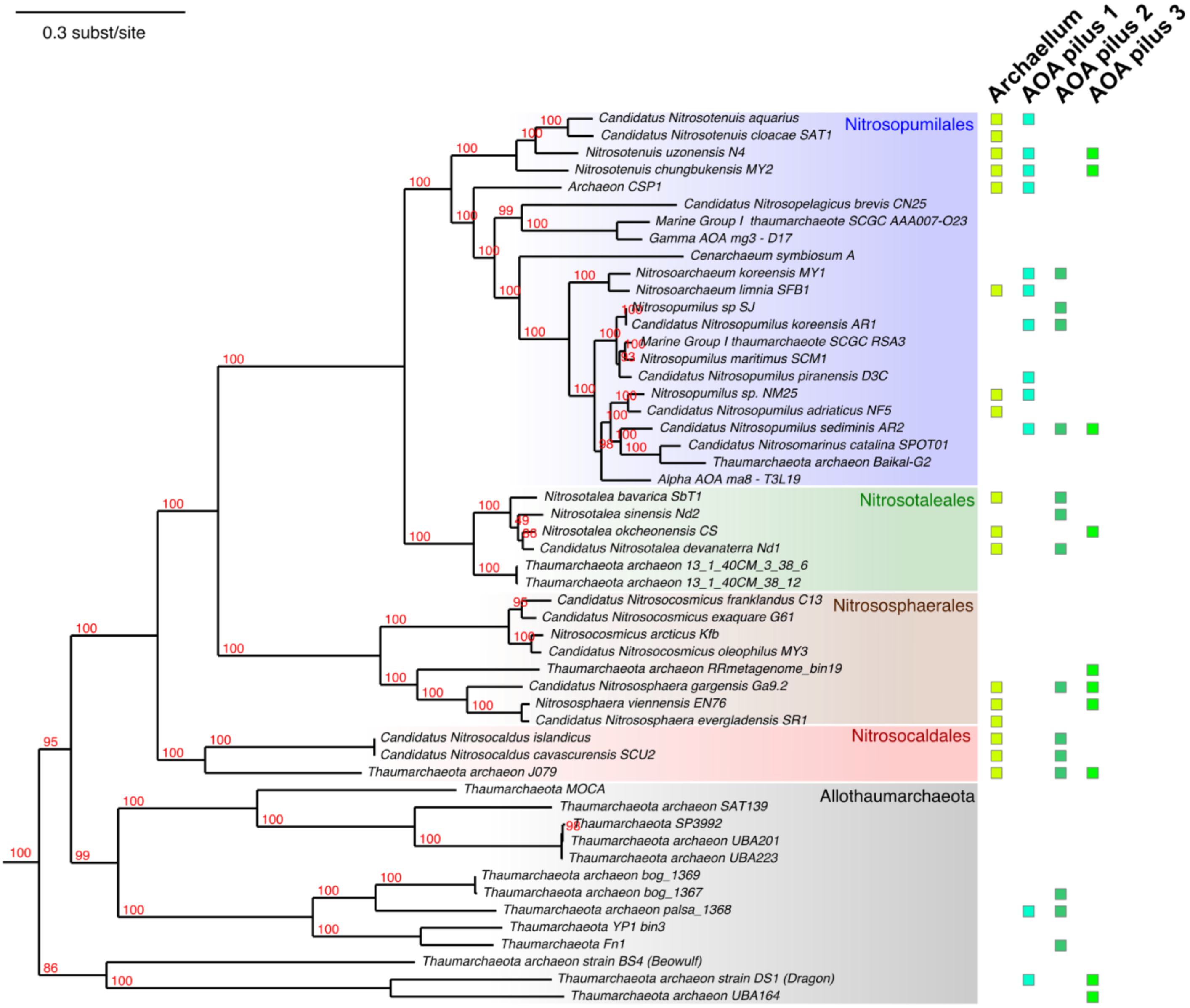


**Figure S4.** Type IV pili distribution in AOA.

Phylogenetic distribution of the four distinct pili found across Thaumarchaeota. The set of different pili found in each Thaumarchaeota genome analysed is shown with boxes of different colors in front of the corresponding species in the tree of Thaumarchaeota. The distinct clades of pili were defined based on a phylogenetic tree of the ATPase (FAM001536, see Fig. 6). Branches with UF-Boot support above 95% are indicated by a red circle.
