## Supplementary material for "Ancestral reconstructions decipher major adaptations of ammonia oxidizing archaea upon radiation into moderate terrestrial and marine environments": Data S1

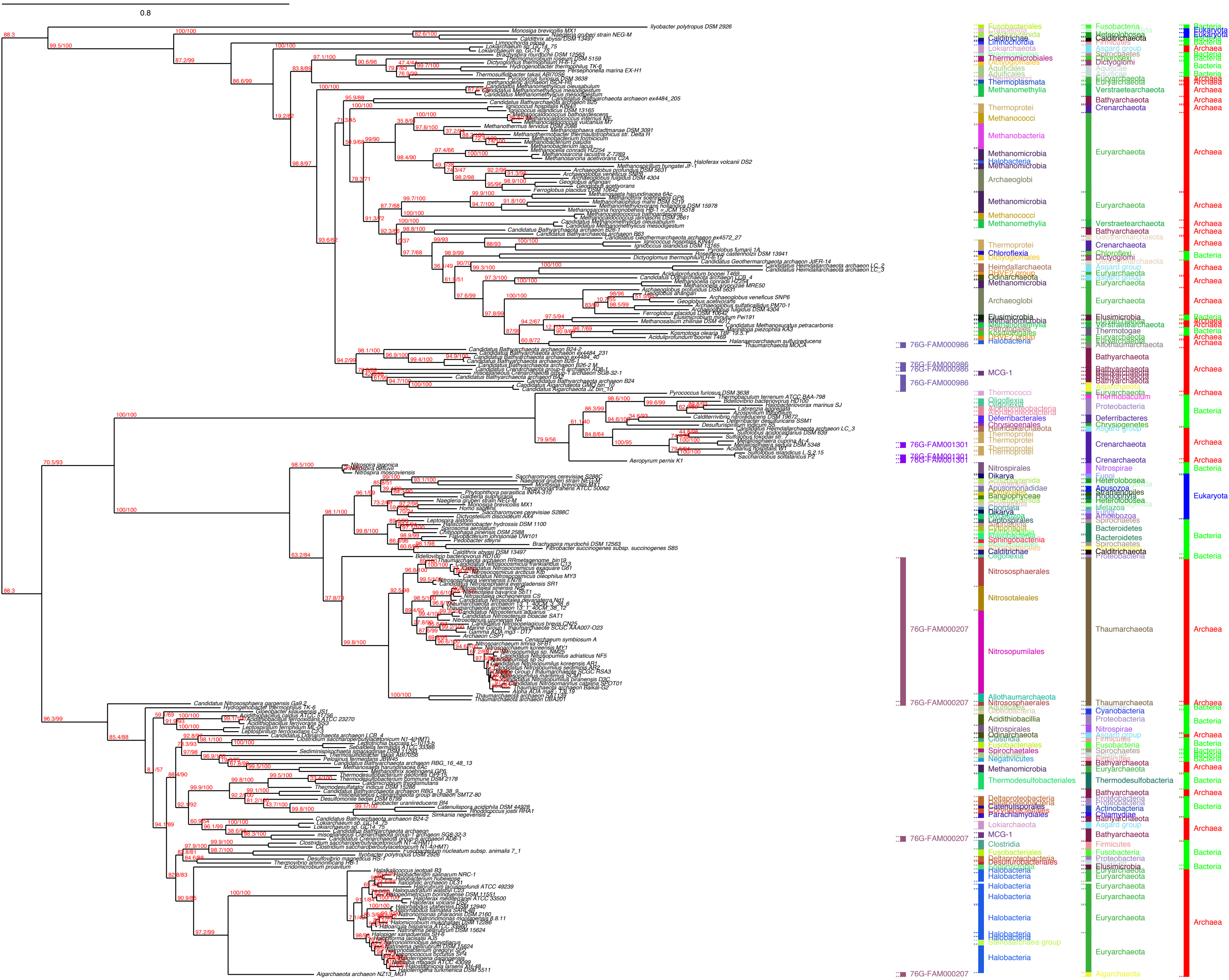

0.8
76G-FAM000986
76G-FAM001301
76G-FAM000207
Archaea
Bacteria
Eukaryota
Euryarchaeota
Thaumarchaeota
Crenarchaeota
Bathyarchaeota
Nitrososphaerales
Nitrosotaleales
Nitrosopumilales
Halobacteria
MCG-1

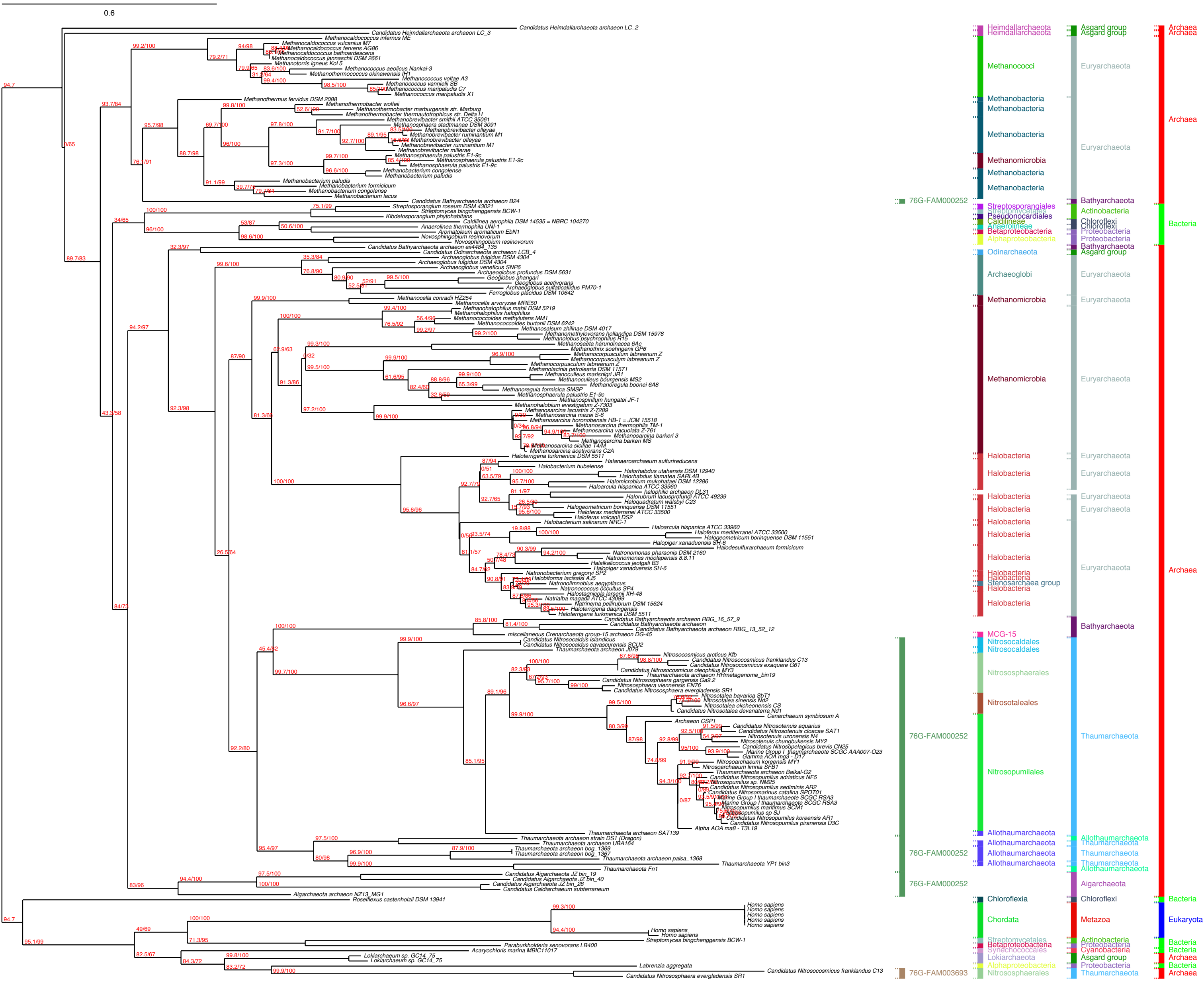

0.6
Heimdallarchaeota
Asgard group
Archaea
Methanococci
Euryarchaeota
Methanobacteria
Methanomicrobia
76G-FAM000252
Bathyarchaeota
Streptosporangiales
Actinobacteria
Streptomycetales
Pseudonocardiales
Chloroflexi
Caldilineae
Anaerolineae
Betaproteobacteria
Proteobacteria
Alphaproteobacteria
Bacteria
Odinarchaeota
Archaeoglobi
Halobacteria
Stenosarchaea group
MCG-15
Nitrosocaldales
Nitrososphaerales
Nitrosotaleales
Thaumarchaeota
Nitrosopumilales
Allothaumarchaeota
Aigarchaeota
Chloroflexia
Chordata
Metazoa
Eukaryota
Synechococcales
Cyanobacteria
Lokiarchaeota
76G-FAM003693

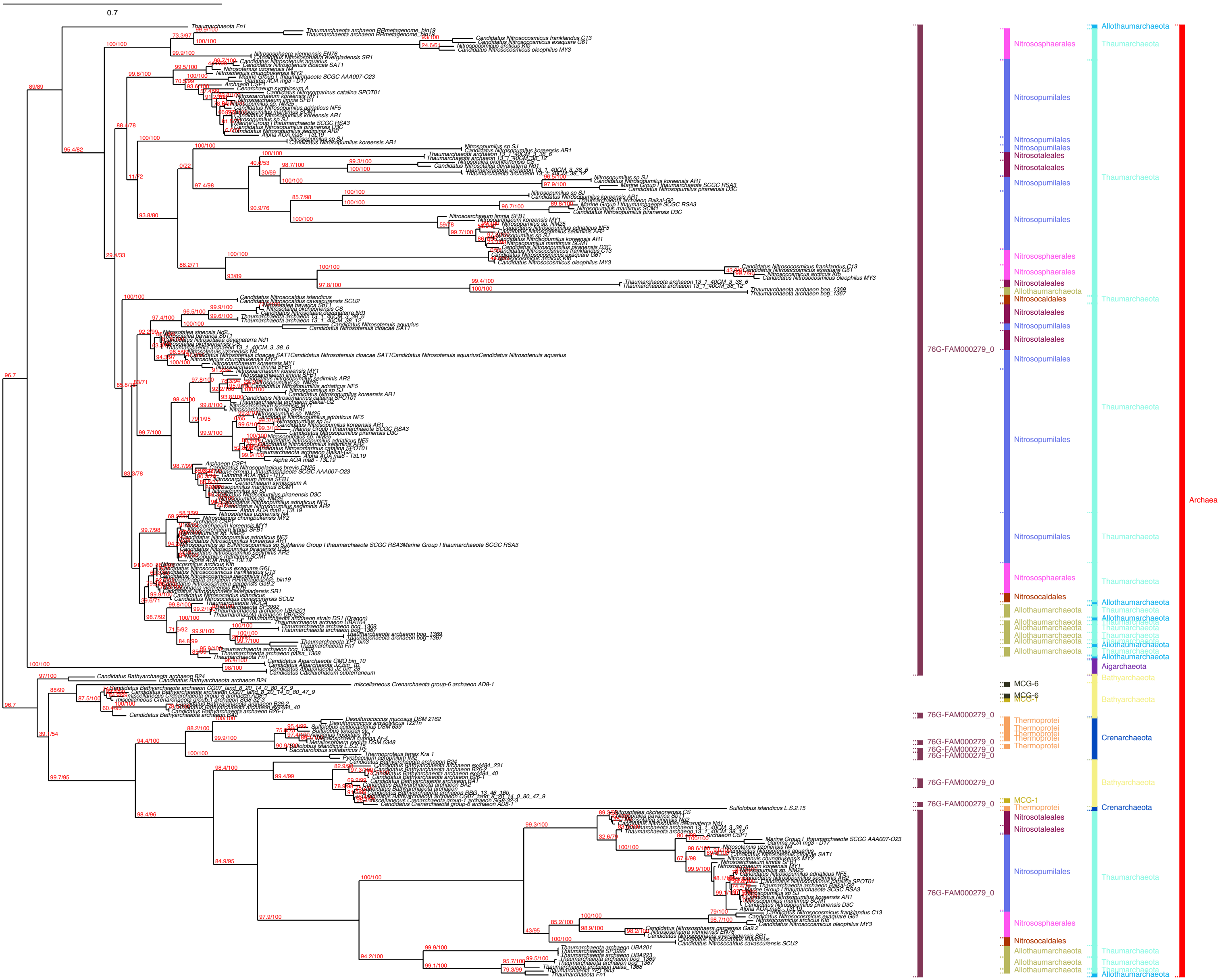
0.7
76G-FAM000279_0
Nitrososphaerales
Nitrosopumilales
Nitrosotaleales
Nitrosocaldales
Allothaumarchaeota
MCG-6
MCG-1
Thermoprotei
Thaumarchaeota
Aigarchaeota
Bathyarchaeota
Crenarchaeota
Archaea

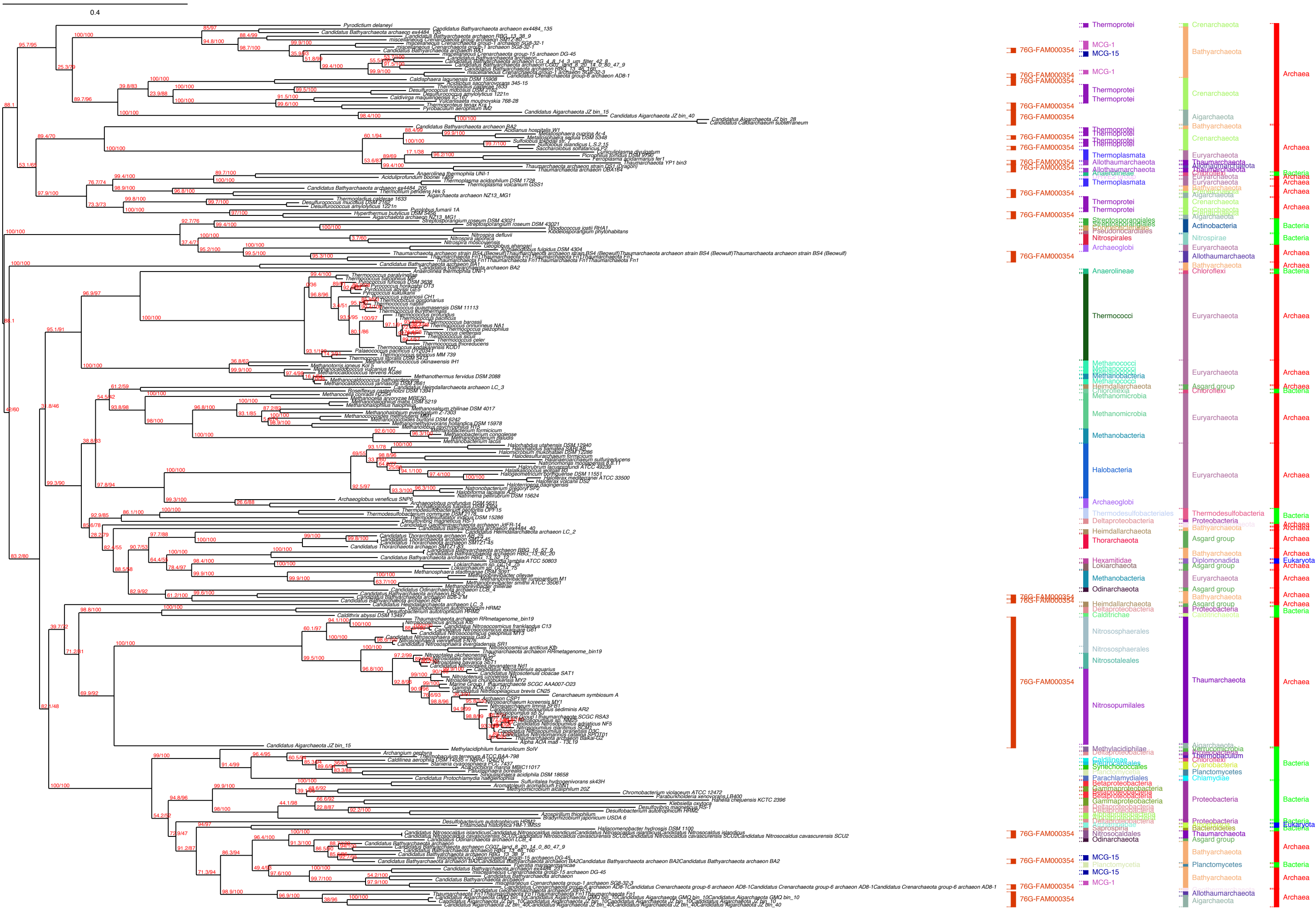

0.4
76G-FAM000354
Thermoprotei
MCG-1
MCG-15
Crenarchaeota
Bathyarchaeota
Aigarchaeota
Archaea
Thermoplasmata
Allothaumarchaeota
Euryarchaeota
Anaerolineae
Chloroflexi
Bacteria
Streptosporangiales
Pseudonocardiales
Actinobacteria
Nitrospirales
Nitrospirae
Archaeoglobi
Thermococci
Methanococci
Methanobacteria
Heimdallarchaeota
Asgard group
Methanomicrobia
Halobacteria
Thermodesulfobacteriales
Thermodesulfobacteria
Deltaproteobacteria
Proteobacteria
Thorarchaeota
Hexamitidae
Diplomonadida
Eukaryota
Lokiarchaeota
Odinarchaeota
Caldithrix
Caldithrichaeota
Nitrososphaerales
Nitrosotaleales
Nitrosopumilales
Thaumarchaeota
Methylacidiphilae
Verrucomicrobia
Thermobaculum
Synechococcales
Cyanobacteria
Planctomycetes
Parachlamydiales
Chlamydiae
Betaproteobacteria
Gammaproteobacteria
Saprospiria
Bacteroidetes
Nitrosocaldales
Planctomycetia

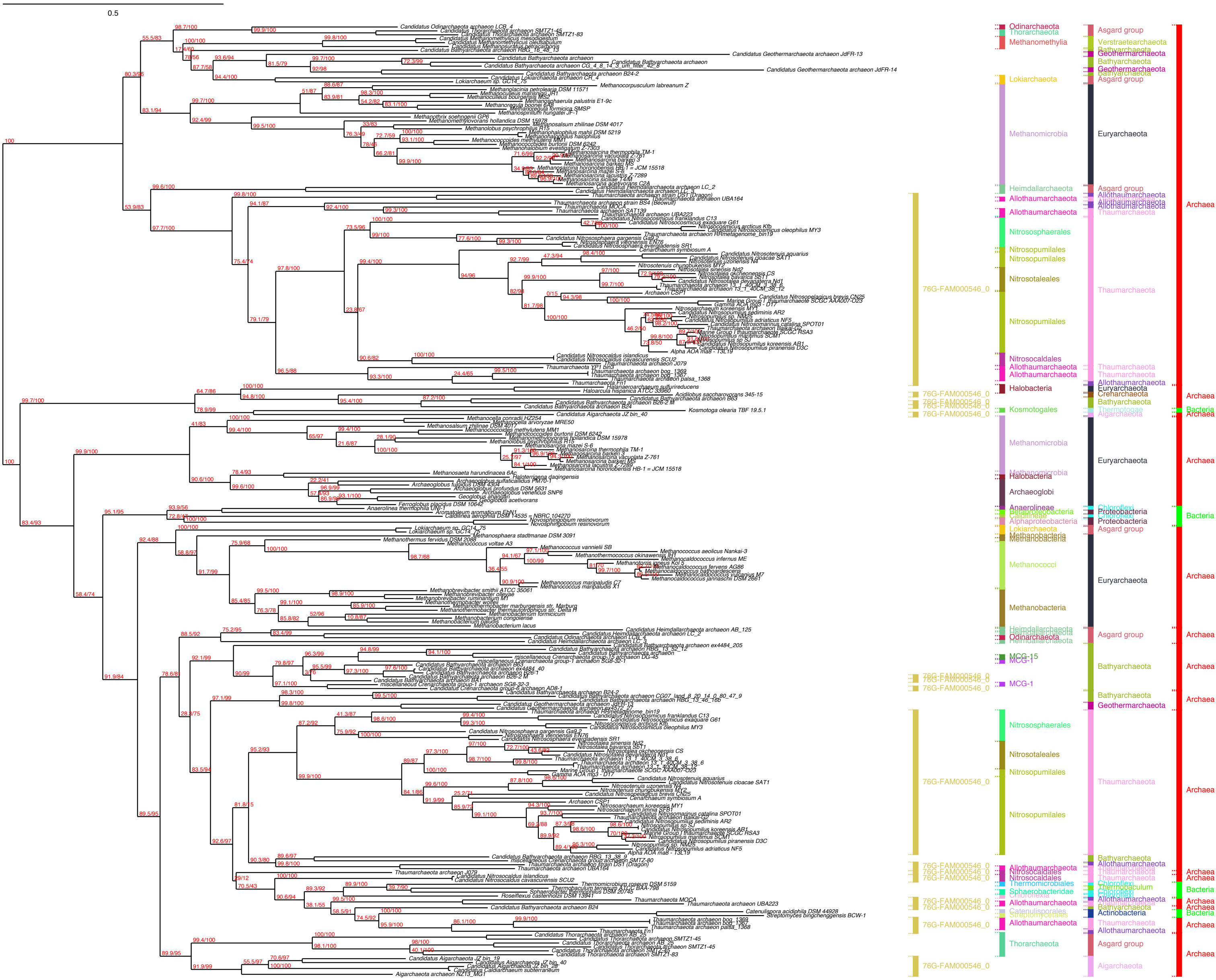
0.5
Odinarchaeota
Thorarchaeota
Methanomethylia
Lokiarchaeota
Methanomicrobia
Heimdallarchaeota
Allothaumarchaeota
Nitrososphaerales
Nitrosopumilales
Nitrosotaleales
Nitrosocaldales
Halobacteria
Kosmotogales
Archaeoglobi
Anaerolineae
Caldilineae
Alphaproteobacteria
Methanobacteria
Methanococci
MCG-15
MCG-1
Thermomicrobiales
Sphaerobacteridae
Catenulisporales
76G-FAM000546_0
Asgard group
Verstraetearchaeota
Geothermarchaeota
Bathyarchaeota
Euryarchaeota
Thaumarchaeota
Crenarchaeota
Thermotogae
Aigarchaeota
Chloroflexi
Proteobacteria
Actinobacteria
Archaea
Bacteria

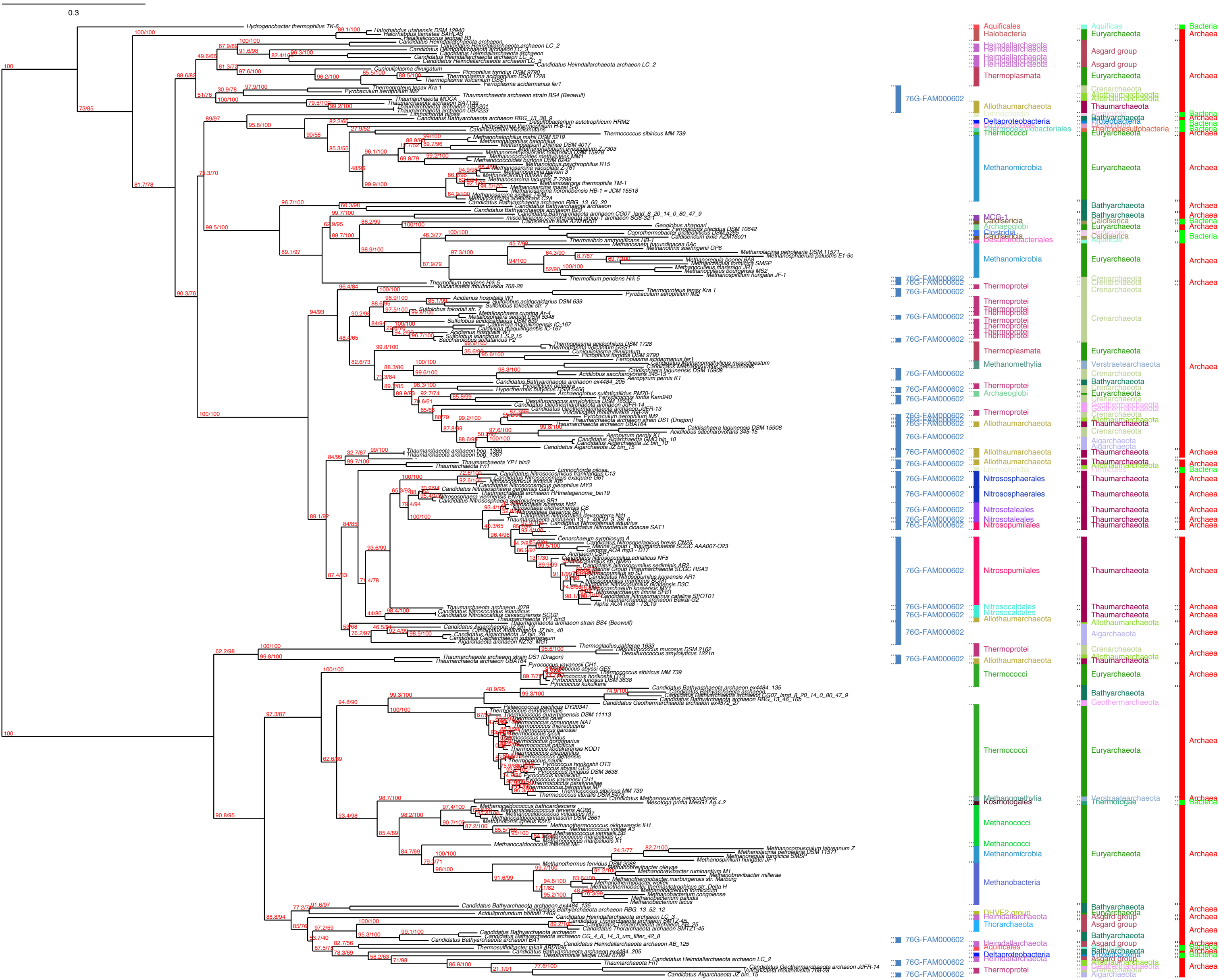

0.3
76G-FAM000602
Aquificales
Halobacteria
Heimdallarchaeota
Thermoplasmata
Allothaumarchaeota
Deltaproteobacteria
Thermococci
Methanomicrobia
Thermoprotei
Methanomethylia
Archaeoglobi
Nitrososphaerales
Nitrosotaleales
Nitrosopumilales
Nitrosocaldales
Methanococci
Kosmotogales
Methanobacteria
Thorarchaeota
Aquificae
Euryarchaeota
Asgard group
Crenarchaeota
Thaumarchaeota
Bathyarchaeota
Proteobacteria
Thermodesulfobacteria
Verstraetearchaeota
Aigarchaeota
Geothermarchaeota
Thermotogae
Bacteria
Archaea

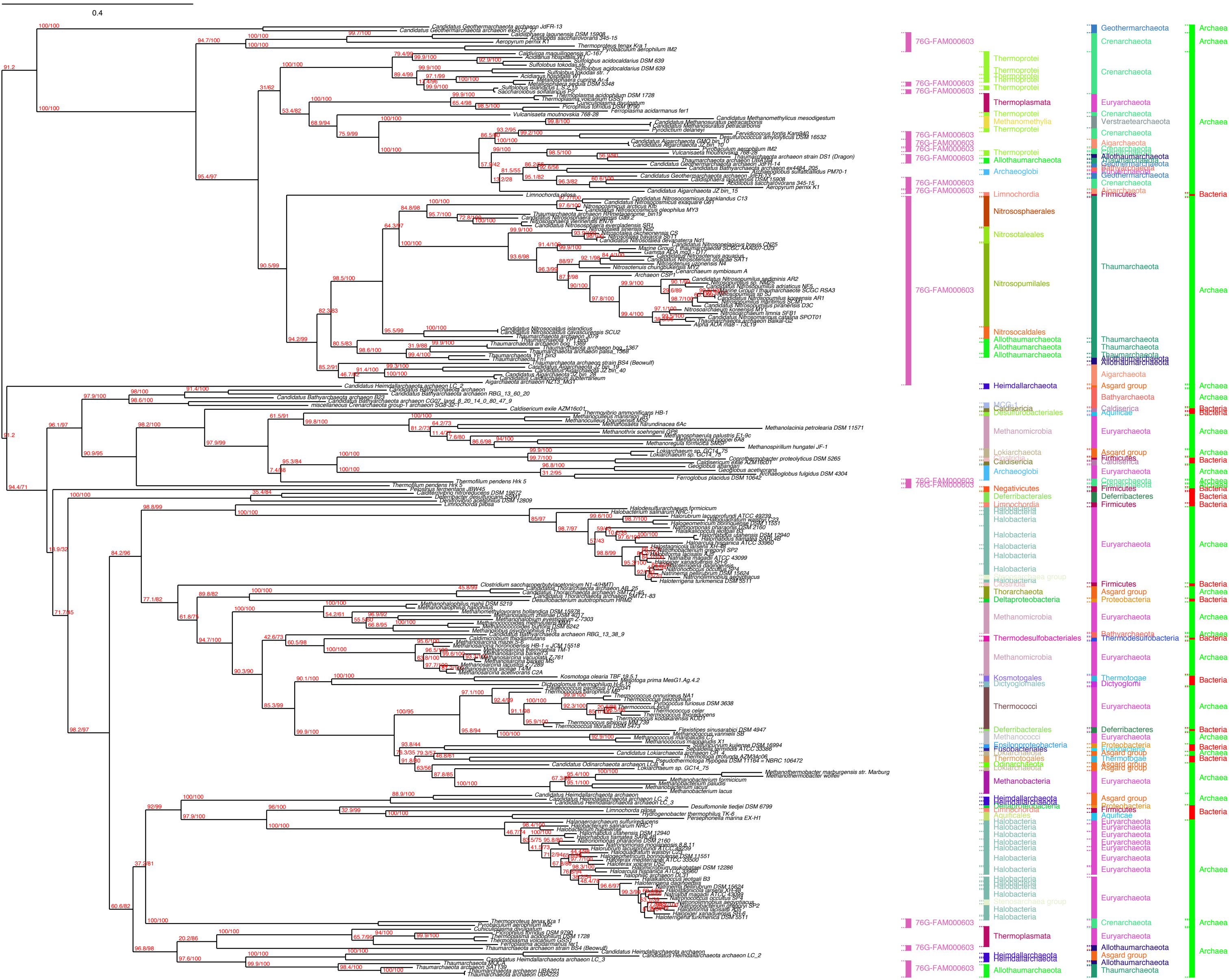
0.4
76G-FAM000603
Thermoprotei
Thermoplasmata
Methanomethylia
Allothaumarchaeota
Archaeoglobi
Limnochordia
Nitrososphaerales
Nitrosotaleales
Nitrosopumilales
Nitrosocaldales
Heimdallarchaeota
Caldisericia
Desulfurobacteriales
Methanomicrobia
Lokiarchaeota
Negativicutes
Deferribacterales
Halobacteria
Thorarchaeota
Deltaproteobacteria
Thermodesulfobacteriales
Kosmotogales
Dictyoglomales
Thermococci
Methanococci
Epsilonproteobacteria
Fusobacteriales
Thermotogales
Odinarchaeota
Methanobacteria
Aquificales
Geothermarchaeota
Crenarchaeota
Euryarchaeota
Verstraetearchaeota
Aigarchaeota
Bathyarchaeota
Firmicutes
Thaumarchaeota
Asgard group
Aquificae
Deferribacteres
Proteobacteria
Thermodesulfobacteria
Thermotogae
Dictyoglomi
Archaea
Bacteria

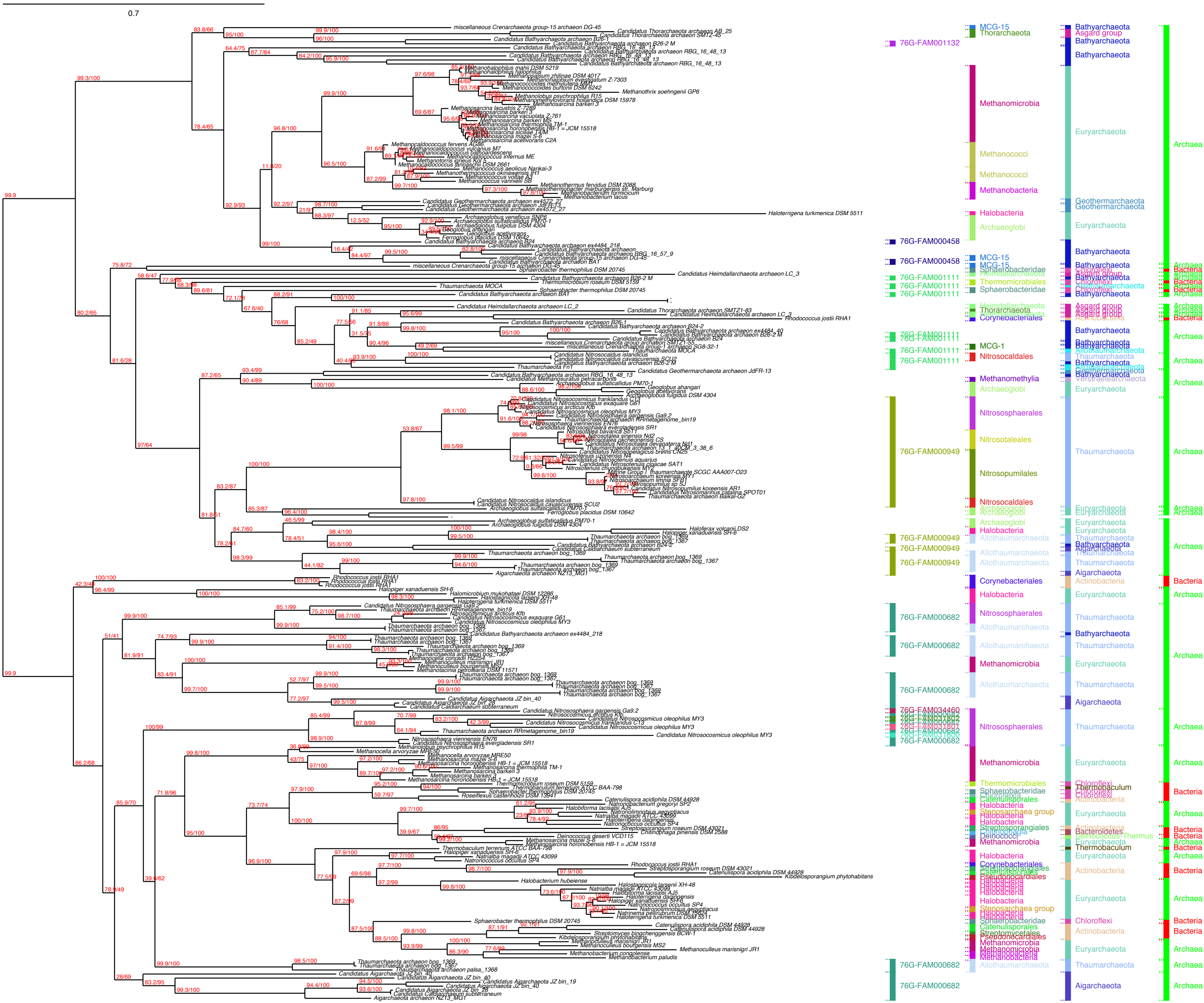
0.7
76G-FAM001132
76G-FAM000458
76G-FAM001111
76G-FAM000949
76G-FAM000682
76G-FAM034460
MCG-15
Thorarchaeota
Methanomicrobia
Methanococci
Methanobacteria
Halobacteria
Archaeoglobi
Sphaerobacteridae
Thermomicrobiales
Heimdallarchaeota
Corynebacteriales
MCG-1
Nitrosocaldales
Methanomethylia
Nitrososphaerales
Nitrosotaleales
Nitrosopumilales
Allothaumarchaeota
Catenulisporales
Stenosarchaea group
Streptosporangiales
Deinococci
Pseudonocardiales
Streptomycetales
Bathyarchaeota
Asgard group
Euryarchaeota
Geothermarchaeota
Chloroflexi
Thaumarchaeota
Verstraetearchaeota
Aigarchaeota
Actinobacteria
Thermobaculum
Bacteroidetes
Deinococcus-Thermus
Archaea
Bacteria

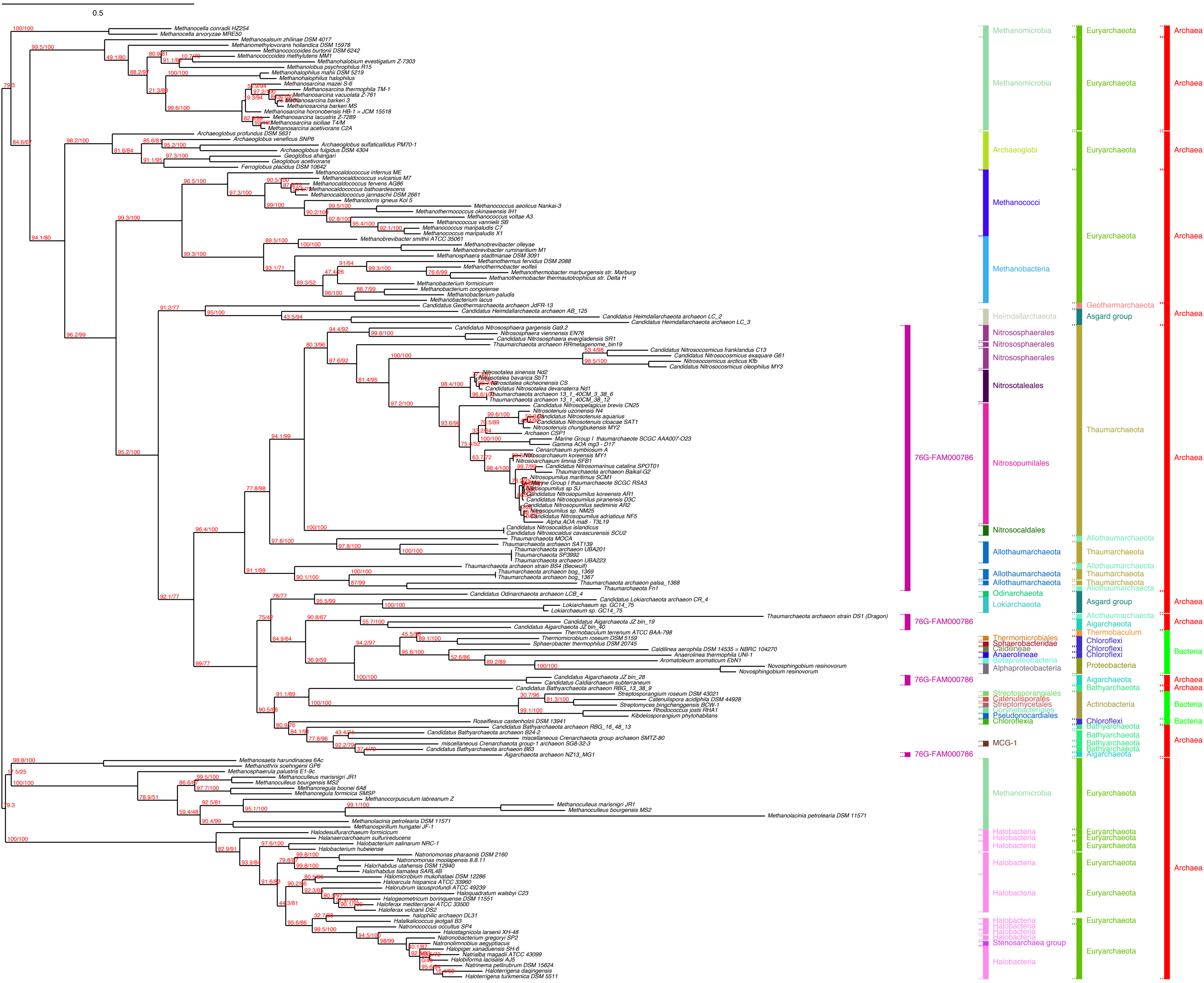

0.5
Methanocella conradii HZ254
Methanocella arvoryzae MRE50
Methanosalsum zhilinae DSM 4017
Methanomethylovorans hollandica DSM 15978
Methanococcoides burtonii DSM 6242
Methanococcoides methylutens MM1
Methanohalobium evestigatum Z-7303
Methanolobus psychrophilus R15
Methanohalophilus mahii DSM 5219
Methanohalophilus halophilus
Methanosarcina mazei S-6
Methanosarcina thermophila TM-1
Methanosarcina vacuolata Z-761
Methanosarcina barkeri 3
Methanosarcina barkeri MS
Methanosarcina horonobensis HB-1 = JCM 15518
Methanosarcina lacustris Z-7289
Methanosarcina siciliae T4/M
Methanosarcina acetivorans C2A
Archaeoglobus profundus DSM 5631
Archaeoglobus veneficus SNP6
Archaeoglobus sulfaticallidus PM70-1
Archaeoglobus fulgidus DSM 4304
Geoglobus ahangari
Geoglobus acetivorans
Ferroglobus placidus DSM 10642
Methanocaldococcus infernus ME
Methanocaldococcus vulcanius M7
Methanocaldococcus fervens AG86
Methanocaldococcus bathoardescens
Methanocaldococcus jannaschii DSM 2661
Methanotorris igneus Kol 5
Methanococcus aeolicus Nankai-3
Methanothermococcus okinawensis IH1
Methanococcus voltae A3
Methanococcus vannielii SB
Methanococcus maripaludis C7
Methanococcus maripaludis X1
Methanobrevibacter smithii ATCC 35061
Methanobrevibacter olleyae
Methanobrevibacter ruminantium M1
Methanosphaera stadtmanae DSM 3091
Methanothermus fervidus DSM 2088
Methanothermobacter wolfeii
Methanothermobacter marburgensis str. Marburg
Methanothermobacter thermautotrophicus str. Delta H
Methanobacterium formicicum
Methanobacterium congolense
Methanobacterium paludis
Methanobacterium lacus
Candidatus Geothermarchaeota archaeon JdFR-13
Candidatus Heimdallarchaeota archaeon AB_125
Candidatus Heimdallarchaeota archaeon LC_2
Candidatus Heimdallarchaeota archaeon LC_3
Candidatus Nitrososphaera gargensis Ga9.2
Nitrososphaera viennensis EN76
Candidatus Nitrososphaera evergladensis SR1
Thaumarchaeota archaeon RRmetagenome_bin19
Candidatus Nitrosocosmicus franklandus C13
Candidatus Nitrosocosmicus exaquare G61
Nitrosocosmicus arcticus Kfb
Candidatus Nitrosocosmicus oleophilus MY3
Nitrosotalea sinensis Nd2
Nitrosotalea bavarica SbT1
Nitrosotalea okcheonensis CS
Candidatus Nitrosotalea devanaterra Nd1
Thaumarchaeota archaeon 13_1_40CM_3_38_6
Thaumarchaeota archaeon 13_1_40CM_38_12
Candidatus Nitrosopelagicus brevis CN25
Nitrosotenuis uzonensis N4
Candidatus Nitrosotenuis aquarius
Candidatus Nitrosotenuis cloacae SAT1
Nitrosotenuis chungbukensis MY2
Archaeon CSP1
Marine Group I thaumarchaeote SCGC AAA007-O23
Gamma AOA mg3 - D17
Cenarchaeum symbiosum A
Nitrosoarchaeum koreensis MY1
Nitrosoarchaeum limnia SFB1
Candidatus Nitrosomarinus catalina SPOT01
Thaumarchaeota archaeon Baikal-G2
Nitrosopumilus maritimus SCM1
Marine Group I thaumarchaeote SCGC RSA3
Nitrosopumilus sp SJ
Candidatus Nitrosopumilus koreensis AR1
Candidatus Nitrosopumilus piranensis D3C
Candidatus Nitrosopumilus sediminis AR2
Nitrosopumilus sp. NM25
Candidatus Nitrosopumilus adriaticus NF5
Alpha AOA ma8 - T3L19
Candidatus Nitrosocaldus islandicus
Candidatus Nitrosocaldus cavascurensis SCU2
Thaumarchaeota MOCA
Thaumarchaeota archaeon SAT139
Thaumarchaeota archaeon UBA201
Thaumarchaeota SP3992
Thaumarchaeota archaeon UBA223
Thaumarchaeota archaeon strain BS4 (Beowulf)
Thaumarchaeota archaeon bog_1369
Thaumarchaeota archaeon bog_1367
Thaumarchaeota archaeon palsa_1368
Thaumarchaeota Fn1
Candidatus Odinarchaeota archaeon LCB_4
Candidatus Lokiarchaeota archaeon CR_4
Lokiarchaeum sp. GC14_75
Lokiarchaeum sp. GC14_75
Thaumarchaeota archaeon strain DS1 (Dragon)
Candidatus Aigarchaeota JZ bin_19
Candidatus Aigarchaeota JZ bin_40
Thermobaculum terrenum ATCC BAA-798
Thermomicrobium roseum DSM 5159
Sphaerobacter thermophilus DSM 20745
Caldilinea aerophila DSM 14535 = NBRC 104270
Anaerolinea thermophila UNI-1
Aromatoleum aromaticum EbN1
Novosphingobium resinovorum
Novosphingobium resinovorum
Candidatus Aigarchaeota JZ bin_28
Candidatus Caldiarchaeum subterraneum
Candidatus Bathyarchaeota archaeon RBG_13_38_9
Streptosporangium roseum DSM 43021
Catenulispora acidiphila DSM 44928
Streptomyces bingchenggensis BCW-1
Rhodococcus jostii RHA1
Kibdelosporangium phytohabitans
Roseiflexus castenholzii DSM 13941
Candidatus Bathyarchaeota archaeon RBG_16_48_13
Candidatus Bathyarchaeota archaeon B24-2
miscellaneous Crenarchaeota group archaeon SMTZ-80
miscellaneous Crenarchaeota group-1 archaeon SG8-32-3
Candidatus Bathyarchaeota archaeon B63
Aigarchaeota archaeon NZ13_MG1
Methanosaeta harundinacea 6Ac
Methanothrix soehngenii GP6
Methanosphaerula palustris E1-9c
Methanoculleus marisnigri JR1
Methanoculleus bourgensis MS2
Methanoregula boonei 6A8
Methanoregula formicica SMSP
Methanocorpusculum labreanum Z
Methanoculleus marisnigri JR1
Methanoculleus bourgensis MS2
Methanolacinia petrolearia DSM 11571
Methanolacinia petrolearia DSM 11571
Methanospirillum hungatei JF-1
Halodesulfurarchaeum formicicum
Halanaeroarchaeum sulfurireducens
Halobacterium salinarum NRC-1
Halobacterium hubeiense
Natronomonas pharaonis DSM 2160
Natronomonas moolapensis 8.8.11
Halorhabdus utahensis DSM 12940
Halorhabdus tiamatea SARL4B
Halomicrobium mukohataei DSM 12286
Haloarcula hispanica ATCC 33960
Halorubrum lacusprofundi ATCC 49239
Haloquadratum walsbyi C23
Halogeometricum borinquense DSM 11551
Haloferax mediterranei ATCC 33500
Haloferax volcanii DS2
halophilic archaeon DL31
Halalkalicoccus jeotgali B3
Natronococcus occultus SP4
Halostagnicola larsenii XH-48
Natronobacterium gregoryi SP2
Natronolimnobius aegyptiacus
Halopiger xanaduensis SH-6
Natrialba magadii ATCC 43099
Halobiforma lacisalsi AJ5
Natrinema pellirubrum DSM 15624
Haloterrigena daqingensis
Haloterrigena turkmenica DSM 5511
76G-FAM000786
76G-FAM000786
76G-FAM000786
76G-FAM000786
Methanomicrobia
Archaeoglobi
Methanococci
Methanobacteria
Heimdallarchaeota
Nitrososphaerales
Nitrosotaleales
Nitrosopumilales
Nitrosocaldales
Allothaumarchaeota
Odinarchaeota
Lokiarchaeota
Thermomicrobiales
Sphaerobacteridae
Caldilineae
Anaerolineae
Betaproteobacteria
Alphaproteobacteria
Streptosporangiales
Catenulisporales
Streptomycetales
Corynebacteriales
Pseudonocardiales
Chloroflexia
MCG-1
Halobacteria
Stenosarchaea group
Euryarchaeota
Geothermarchaeota
Asgard group
Thaumarchaeota
Aigarchaeota
Thermobaculum
Chloroflexi
Proteobacteria
Bathyarchaeota
Actinobacteria
Archaea
Bacteria

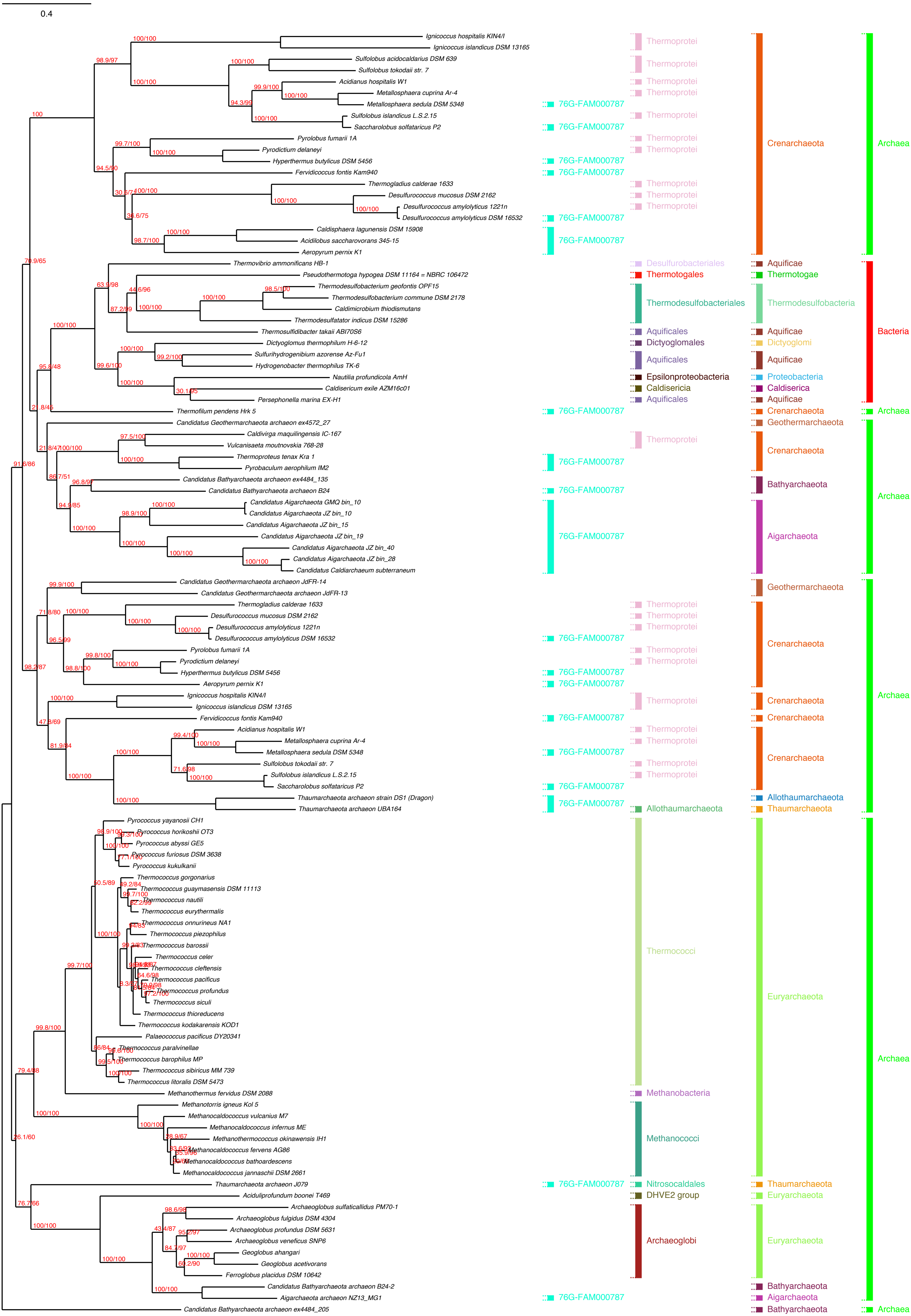
0.4
Ignicoccus hospitalis KIN4/I
Ignicoccus islandicus DSM 13165
Sulfolobus acidocaldarius DSM 639
Sulfolobus tokodaii str. 7
Acidianus hospitalis W1
Metallosphaera cuprina Ar-4
Metallosphaera sedula DSM 5348
Sulfolobus islandicus L.S.2.15
Saccharolobus solfataricus P2
Pyrolobus fumarii 1A
Pyrodictium delaneyi
Hyperthermus butylicus DSM 5456
Fervidicoccus fontis Kam940
Thermogladius calderae 1633
Desulfurococcus mucosus DSM 2162
Desulfurococcus amylolyticus 1221n
Desulfurococcus amylolyticus DSM 16532
Caldisphaera lagunensis DSM 15908
Acidilobus saccharovorans 345-15
Aeropyrum pernix K1
Thermovibrio ammonificans HB-1
Pseudothermotoga hypogea DSM 11164 = NBRC 106472
Thermodesulfobacterium geofontis OPF15
Thermodesulfobacterium commune DSM 2178
Caldimicrobium thiodismutans
Thermodesulfatator indicus DSM 15286
Thermosulfidibacter takaii ABI70S6
Dictyoglomus thermophilum H-6-12
Sulfurihydrogenibium azorense Az-Fu1
Hydrogenobacter thermophilus TK-6
Nautilia profundicola AmH
Caldisericum exile AZM16c01
Persephonella marina EX-H1
Thermofilum pendens Hrk 5
Candidatus Geothermarchaeota archaeon ex4572_27
Caldivirga maquilingensis IC-167
Vulcanisaeta moutnovskia 768-28
Thermoproteus tenax Kra 1
Pyrobaculum aerophilum IM2
Candidatus Bathyarchaeota archaeon ex4484_135
Candidatus Bathyarchaeota archaeon B24
Candidatus Aigarchaeota GMQ bin_10
Candidatus Aigarchaeota JZ bin_10
Candidatus Aigarchaeota JZ bin_15
Candidatus Aigarchaeota JZ bin_19
Candidatus Aigarchaeota JZ bin_40
Candidatus Aigarchaeota JZ bin_28
Candidatus Caldiarchaeum subterraneum
Candidatus Geothermarchaeota archaeon JdFR-14
Candidatus Geothermarchaeota archaeon JdFR-13
Thermogladius calderae 1633
Desulfurococcus mucosus DSM 2162
Desulfurococcus amylolyticus 1221n
Desulfurococcus amylolyticus DSM 16532
Pyrolobus fumarii 1A
Pyrodictium delaneyi
Hyperthermus butylicus DSM 5456
Aeropyrum pernix K1
Ignicoccus hospitalis KIN4/I
Ignicoccus islandicus DSM 13165
Fervidicoccus fontis Kam940
Acidianus hospitalis W1
Metallosphaera cuprina Ar-4
Metallosphaera sedula DSM 5348
Sulfolobus tokodaii str. 7
Sulfolobus islandicus L.S.2.15
Saccharolobus solfataricus P2
Thaumarchaeota archaeon strain DS1 (Dragon)
Thaumarchaeota archaeon UBA164
Pyrococcus yayanosii CH1
Pyrococcus horikoshii OT3
Pyrococcus abyssi GE5
Pyrococcus furiosus DSM 3638
Pyrococcus kukulkanii
Thermococcus gorgonarius
Thermococcus guaymasensis DSM 11113
Thermococcus nautili
Thermococcus eurythermalis
Thermococcus onnurineus NA1
Thermococcus piezophilus
Thermococcus barossii
Thermococcus celer
Thermococcus cleftensis
Thermococcus pacificus
Thermococcus profundus
Thermococcus siculi
Thermococcus thioreducens
Thermococcus kodakarensis KOD1
Palaeococcus pacificus DY20341
Thermococcus paralvinellae
Thermococcus barophilus MP
Thermococcus sibiricus MM 739
Thermococcus litoralis DSM 5473
Methanothermus fervidus DSM 2088
Methanotorris igneus Kol 5
Methanocaldococcus vulcanius M7
Methanocaldococcus infernus ME
Methanothermococcus okinawensis IH1
Methanocaldococcus fervens AG86
Methanocaldococcus bathoardescens
Methanocaldococcus jannaschii DSM 2661
Thaumarchaeota archaeon J079
Aciduliprofundum boonei T469
Archaeoglobus sulfaticallidus PM70-1
Archaeoglobus fulgidus DSM 4304
Archaeoglobus profundus DSM 5631
Archaeoglobus veneficus SNP6
Geoglobus ahangari
Geoglobus acetivorans
Ferroglobus placidus DSM 10642
Candidatus Bathyarchaeota archaeon B24-2
Aigarchaeota archaeon NZ13_MG1
Candidatus Bathyarchaeota archaeon ex4484_205
76G-FAM000787
Thermoprotei
Desulfurobacteriales
Thermotogales
Thermodesulfobacteriales
Aquificales
Dictyoglomales
Epsilonproteobacteria
Caldisericia
Allothaumarchaeota
Thermococci
Methanobacteria
Methanococci
Nitrososcaldales
DHVE2 group
Archaeoglobi
Crenarchaeota
Aquificae
Thermotogae
Thermodesulfobacteria
Dictyoglomi
Proteobacteria
Caldiserica
Geothermarchaeota
Bathyarchaeota
Aigarchaeota
Thaumarchaeota
Euryarchaeota
Archaea
Bacteria

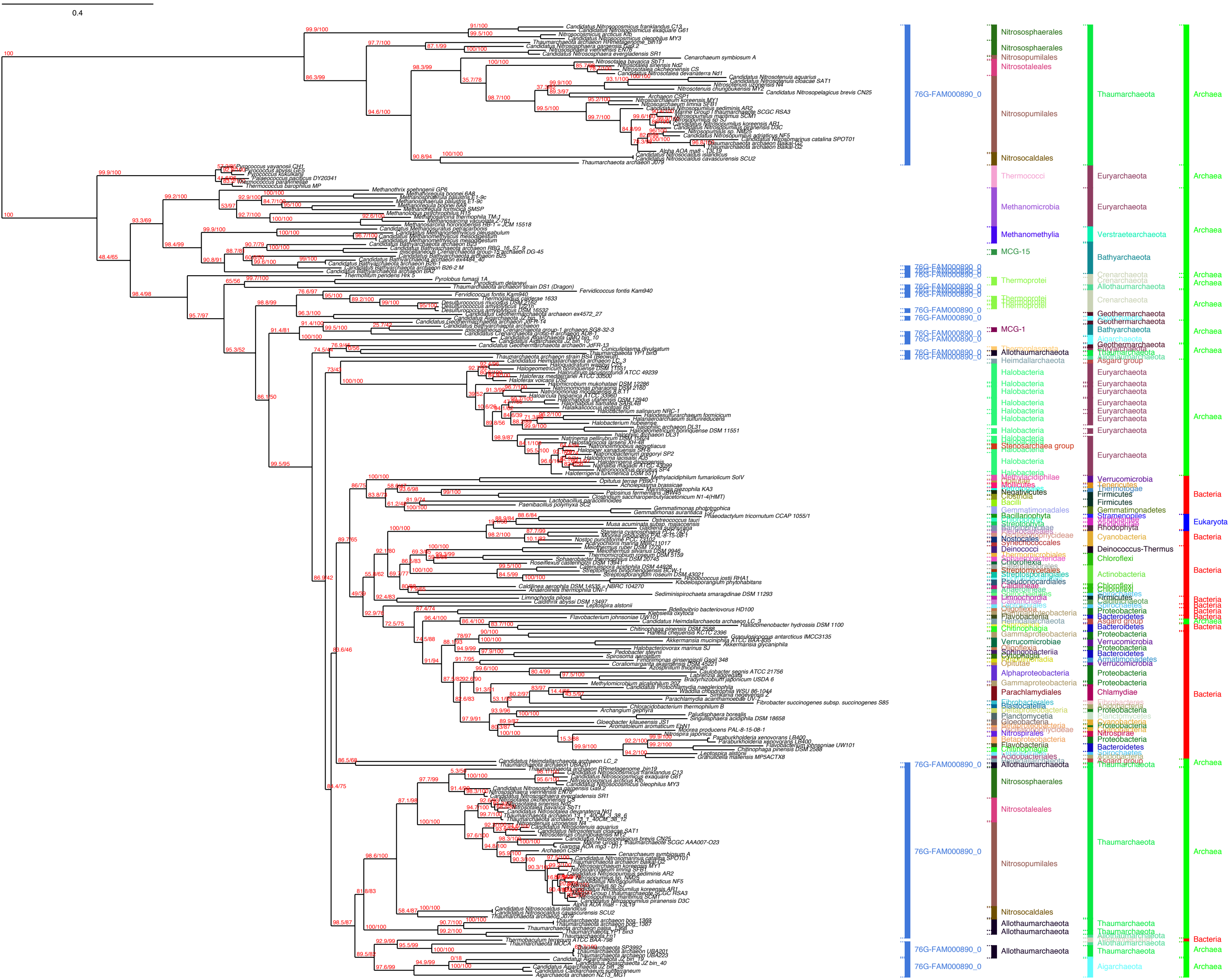

0.4
76G-FAM000890_0
Nitrososphaerales
Nitrosopumilales
Nitrosotaleales
Nitrosocaldales
Thaumarchaeota
Archaea
Thermococci
Methanomicrobia
Methanomethylia
Euryarchaeota
Verstraetearchaeota
MCG-15
Bathyarchaeota
Thermoprotei
Crenarchaeota
Allothaumarchaeota
Geothermarchaeota
MCG-1
Aigarchaeota
Heimdallarchaeota
Asgard group
Halobacteria
Stenosarchaea group
Bacteria
Eukaryota
Verrucomicrobia
Firmicutes
Gemmatimonadetes
Cyanobacteria
Deinococcus-Thermus
Chloroflexi
Actinobacteria
Proteobacteria
Bacteroidetes
Chlamydiae
Nitrospirae

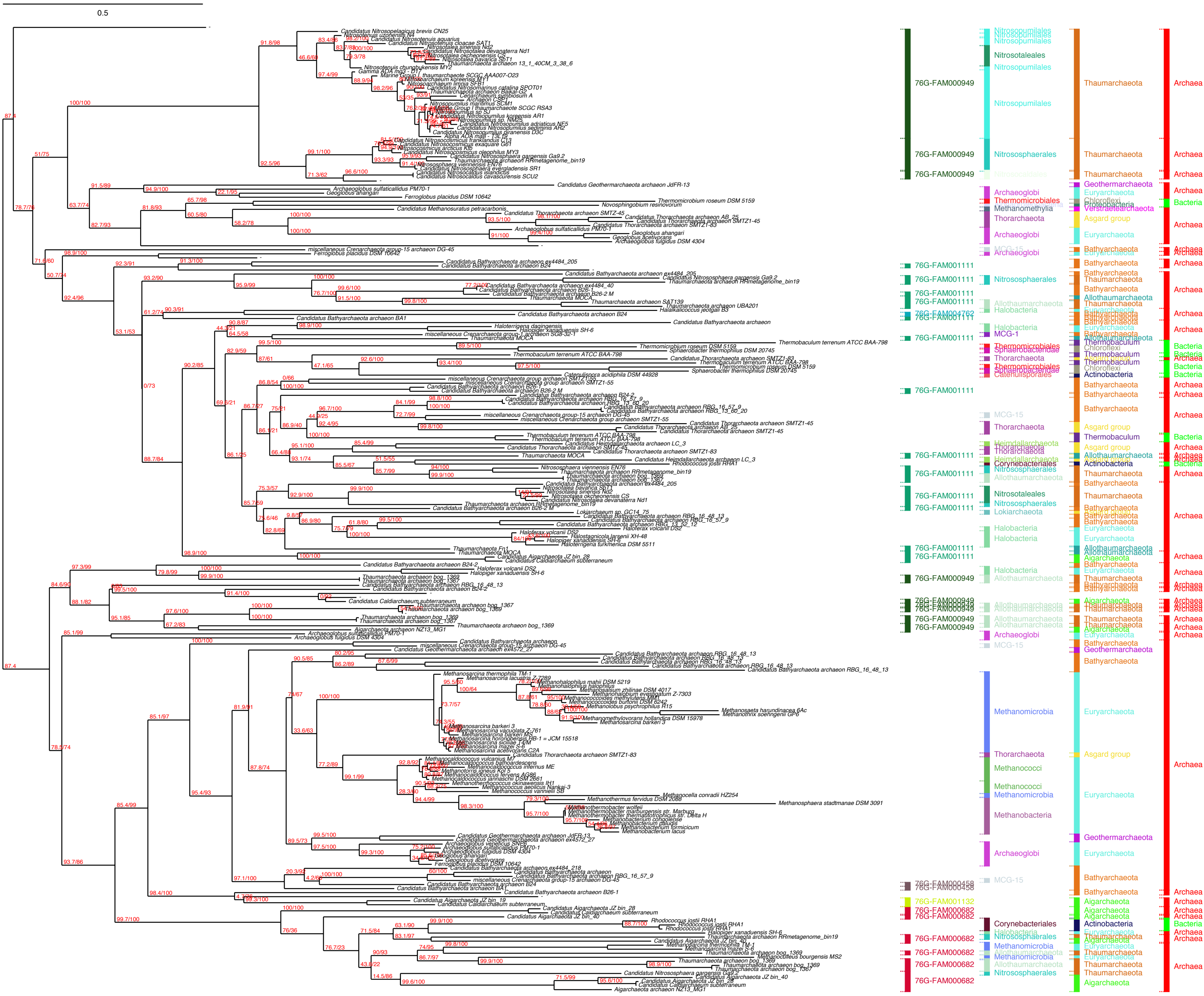

0.5
76G-FAM000949
76G-FAM001111
76G-FAM004762
76G-FAM000458
76G-FAM001132
76G-FAM000682
Nitrosopumilales
Nitrosotaleales
Nitrososphaerales
Archaeoglobi
Thermomicrobiales
Methanomethylia
Thorarchaeota
MCG-15
Allothaumarchaeota
Halobacteria
MCG-1
Sphaerobacteridae
Catenulisporales
Heimdallarchaeota
Corynebacteriales
Lokiarchaeota
Methanomicrobia
Methanococci
Methanobacteria
Thaumarchaeota
Geothermarchaeota
Euryarchaeota
Chloroflexi
Proteobacteria
Verstraetearchaeota
Asgard group
Bathyarchaeota
Thermobaculum
Actinobacteria
Aigarchaeota
Archaea
Bacteria

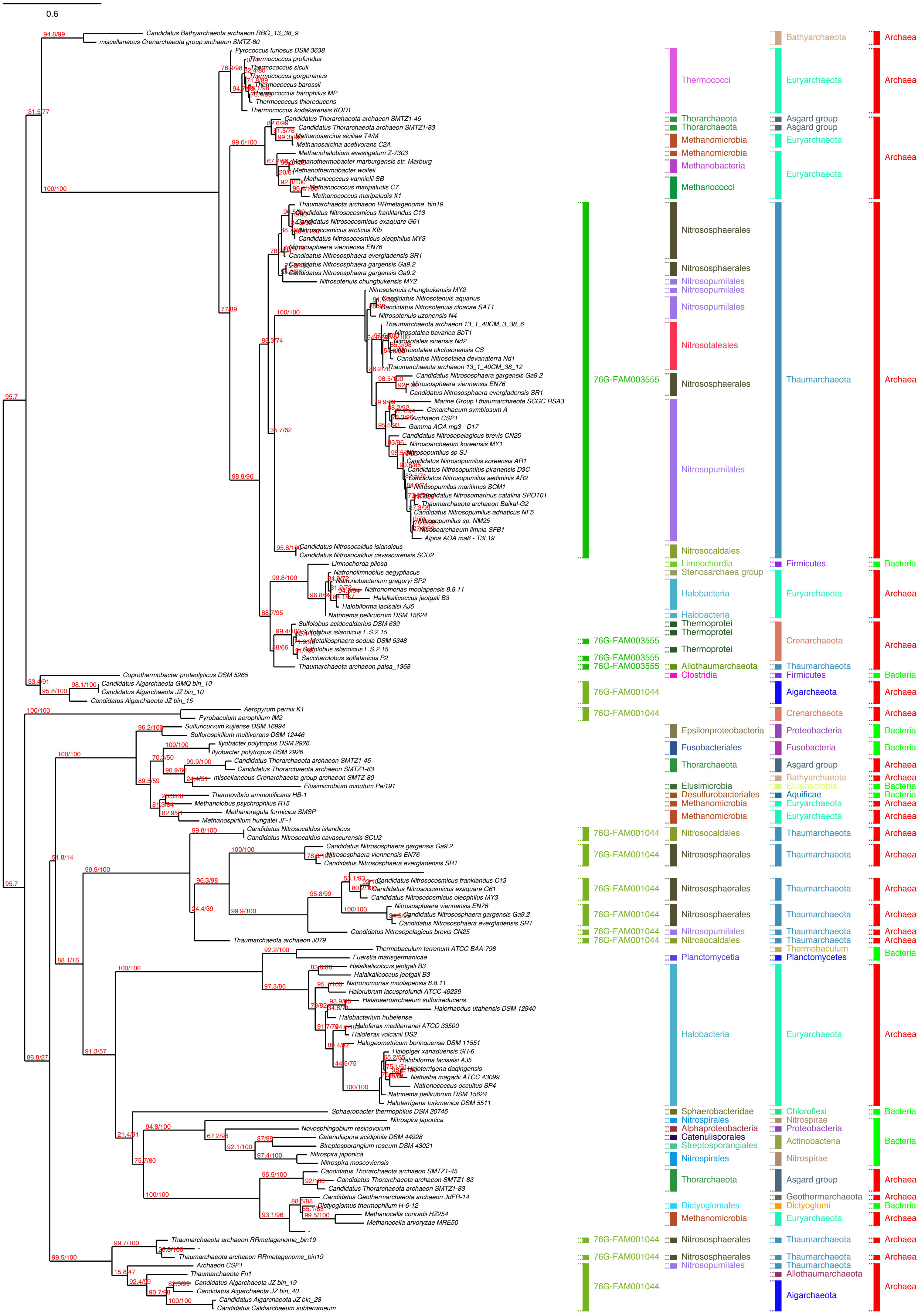

0.6
Candidatus Bathyarchaeota archaeon RBG_13_38_9
miscellaneous Crenarchaeota group archaeon SMTZ-80
Pyrococcus furiosus DSM 3638
Thermococcus profundus
Thermococcus siculi
Thermococcus gorgonarius
Thermococcus barossii
Thermococcus barophilus MP
Thermococcus thioreducens
Thermococcus kodakarensis KOD1
Candidatus Thorarchaeota archaeon SMTZ1-45
Candidatus Thorarchaeota archaeon SMTZ1-83
Methanosarcina siciliae T4/M
Methanosarcina acetivorans C2A
Methanohalobium evestigatum Z-7303
Methanothermobacter marburgensis str. Marburg
Methanothermobacter wolfeii
Methanococcus vannielii SB
Methanococcus maripaludis C7
Methanococcus maripaludis X1
Thaumarchaeota archaeon RRmetagenome_bin19
Candidatus Nitrosocosmicus franklandus C13
Candidatus Nitrosocosmicus exaquare G61
Nitrosocosmicus arcticus Kfb
Candidatus Nitrosocosmicus oleophilus MY3
Nitrososphaera viennensis EN76
Candidatus Nitrososphaera evergladensis SR1
Candidatus Nitrososphaera gargensis Ga9.2
Candidatus Nitrososphaera gargensis Ga9.2
Nitrosotenuis chungbukensis MY2
Nitrosotenuis chungbukensis MY2
Candidatus Nitrosotenuis aquarius
Candidatus Nitrosotenuis cloacae SAT1
Nitrosotenuis uzonensis N4
Thaumarchaeota archaeon 13_1_40CM_3_38_6
Nitrosotalea bavarica SbT1
Nitrosotalea sinensis Nd2
Nitrosotalea okcheonensis CS
Candidatus Nitrosotalea devanaterra Nd1
Thaumarchaeota archaeon 13_1_40CM_38_12
Candidatus Nitrososphaera gargensis Ga9.2
Nitrososphaera viennensis EN76
Candidatus Nitrososphaera evergladensis SR1
Marine Group I thaumarchaeote SCGC RSA3
Cenarchaeum symbiosum A
Archaeon CSP1
Gamma AOA mg3 - D17
Candidatus Nitrosopelagicus brevis CN25
Nitrosoarchaeum koreensis MY1
Nitrosopumilus sp SJ
Candidatus Nitrosopumilus koreensis AR1
Candidatus Nitrosopumilus piranensis D3C
Candidatus Nitrosopumilus sediminis AR2
Nitrosopumilus maritimus SCM1
Candidatus Nitrosomarinus catalina SPOT01
Thaumarchaeota archaeon Baikal-G2
Candidatus Nitrosopumilus adriaticus NF5
Nitrosopumilus sp. NM25
Nitrosoarchaeum limnia SFB1
Alpha AOA ma8 - T3L19
Candidatus Nitrosocaldus islandicus
Candidatus Nitrosocaldus cavascurensis SCU2
Limnochorda pilosa
Natronolimnobius aegyptiacus
Natronobacterium gregoryi SP2
Natronomonas moolapensis 8.8.11
Halalkalicoccus jeotgali B3
Halobiforma lacisalsi AJ5
Natrinema pellirubrum DSM 15624
Sulfolobus acidocaldarius DSM 639
Sulfolobus islandicus L.S.2.15
Metallosphaera sedula DSM 5348
Sulfolobus islandicus L.S.2.15
Saccharolobus solfataricus P2
Thaumarchaeota archaeon palsa_1368
Coprothermobacter proteolyticus DSM 5265
Candidatus Aigarchaeota GMQ bin_10
Candidatus Aigarchaeota JZ bin_10
Candidatus Aigarchaeota JZ bin_15
Aeropyrum pernix K1
Pyrobaculum aerophilum IM2
Sulfuricurvum kujiense DSM 16994
Sulfurospirillum multivorans DSM 12446
Ilyobacter polytropus DSM 2926
Ilyobacter polytropus DSM 2926
Candidatus Thorarchaeota archaeon SMTZ1-45
Candidatus Thorarchaeota archaeon SMTZ1-83
miscellaneous Crenarchaeota group archaeon SMTZ-80
Elusimicrobium minutum Pei191
Thermovibrio ammonificans HB-1
Methanolobus psychrophilus R15
Methanoregula formicica SMSP
Methanospirillum hungatei JF-1
Candidatus Nitrosocaldus islandicus
Candidatus Nitrosocaldus cavascurensis SCU2
Candidatus Nitrososphaera gargensis Ga9.2
Nitrososphaera viennensis EN76
Candidatus Nitrososphaera evergladensis SR1
Candidatus Nitrosocosmicus franklandus C13
Candidatus Nitrosocosmicus exaquare G61
Candidatus Nitrosocosmicus oleophilus MY3
Nitrososphaera viennensis EN76
Candidatus Nitrososphaera gargensis Ga9.2
Candidatus Nitrososphaera evergladensis SR1
Candidatus Nitrosopelagicus brevis CN25
Thaumarchaeota archaeon J079
Thermobaculum terrenum ATCC BAA-798
Fuerstia marisgermanicae
Halalkalicoccus jeotgali B3
Halalkalicoccus jeotgali B3
Natronomonas moolapensis 8.8.11
Halorubrum lacusprofundi ATCC 49239
Halanaeroarchaeum sulfurireducens
Halorhabdus utahensis DSM 12940
Halobacterium hubeiense
Haloferax mediterranei ATCC 33500
Haloferax volcanii DS2
Halogeometricum borinquense DSM 11551
Halopiger xanaduensis SH-6
Halobiforma lacisalsi AJ5
Haloterrigena daqingensis
Natrialba magadii ATCC 43099
Natronococcus occultus SP4
Natrinema pellirubrum DSM 15624
Haloterrigena turkmenica DSM 5511
Sphaerobacter thermophilus DSM 20745
Nitrospira japonica
Novosphingobium resinovorum
Catenulispora acidiphila DSM 44928
Streptosporangium roseum DSM 43021
Nitrospira japonica
Nitrospira moscoviensis
Candidatus Thorarchaeota archaeon SMTZ1-45
Candidatus Thorarchaeota archaeon SMTZ1-83
Candidatus Thorarchaeota archaeon SMTZ1-83
Candidatus Geothermarchaeota archaeon JdFR-14
Dictyoglomus thermophilum H-6-12
Methanocella conradii HZ254
Methanocella arvoryzae MRE50
Thaumarchaeota archaeon RRmetagenome_bin19
Thaumarchaeota archaeon RRmetagenome_bin19
Archaeon CSP1
Thaumarchaeota Fn1
Candidatus Aigarchaeota JZ bin_19
Candidatus Aigarchaeota JZ bin_40
Candidatus Aigarchaeota JZ bin_28
Candidatus Caldiarchaeum subterraneum
76G-FAM003555
76G-FAM001044
Thermococci
Thorarchaeota
Methanomicrobia
Methanobacteria
Methanococci
Nitrososphaerales
Nitrosopumilales
Nitrosotaleales
Nitrosocaldales
Limnochordia
Stenosarchaea group
Halobacteria
Thermoprotei
Allothaumarchaeota
Clostridia
Epsilonproteobacteria
Fusobacteriales
Elusimicrobia
Desulfurobacteriales
Planctomycetia
Sphaerobacteridae
Nitrospirales
Alphaproteobacteria
Catenulisporales
Streptosporangiales
Dictyoglomales
Bathyarchaeota
Euryarchaeota
Asgard group
Thaumarchaeota
Firmicutes
Crenarchaeota
Aigarchaeota
Proteobacteria
Fusobacteria
Aquificae
Thermobaculum
Planctomycetes
Chloroflexi
Nitrospirae
Actinobacteria
Geothermarchaeota
Dictyoglomi
Allothaumarchaeota
Archaea
Bacteria

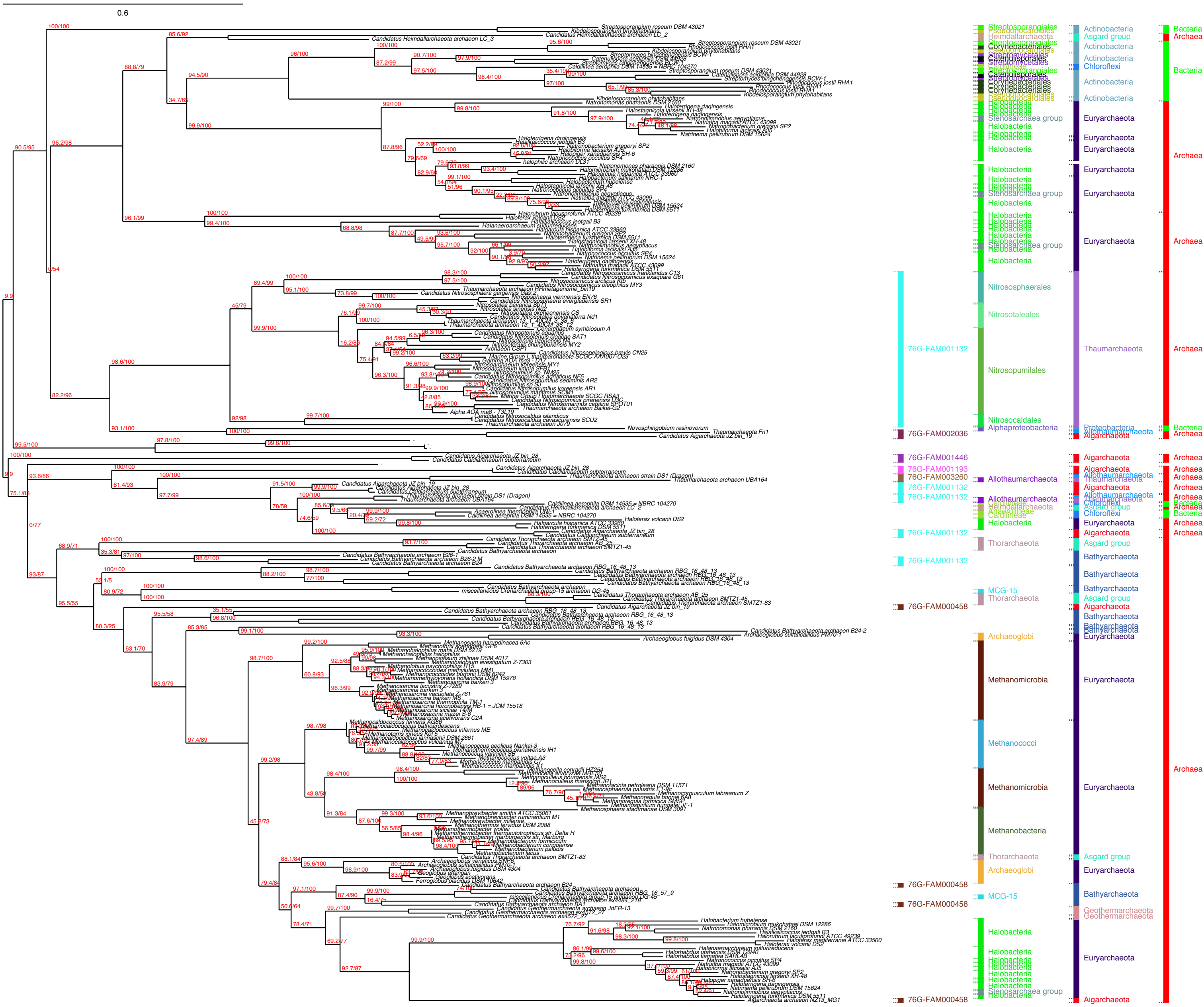

0.6
76G-FAM001132
76G-FAM002036
76G-FAM001446
76G-FAM001193
76G-FAM003260
76G-FAM000458
Actinobacteria
Asgard group
Chloroflexi
Euryarchaeota
Thaumarchaeota
Proteobacteria
Allothaumarchaeota
Aigarchaeota
Bathyarchaeota
Geothermarchaeota
Halobacteria
Stenosarchaea group
Nitrososphaerales
Nitrosotaleales
Nitrosopumilales
Nitrosocaldales
Alphaproteobacteria
Heimdallarchaeota
Thorarchaeota
MCG-15
Archaeoglobi
Methanomicrobia
Methanococci
Methanobacteria
Bacteria
Archaea

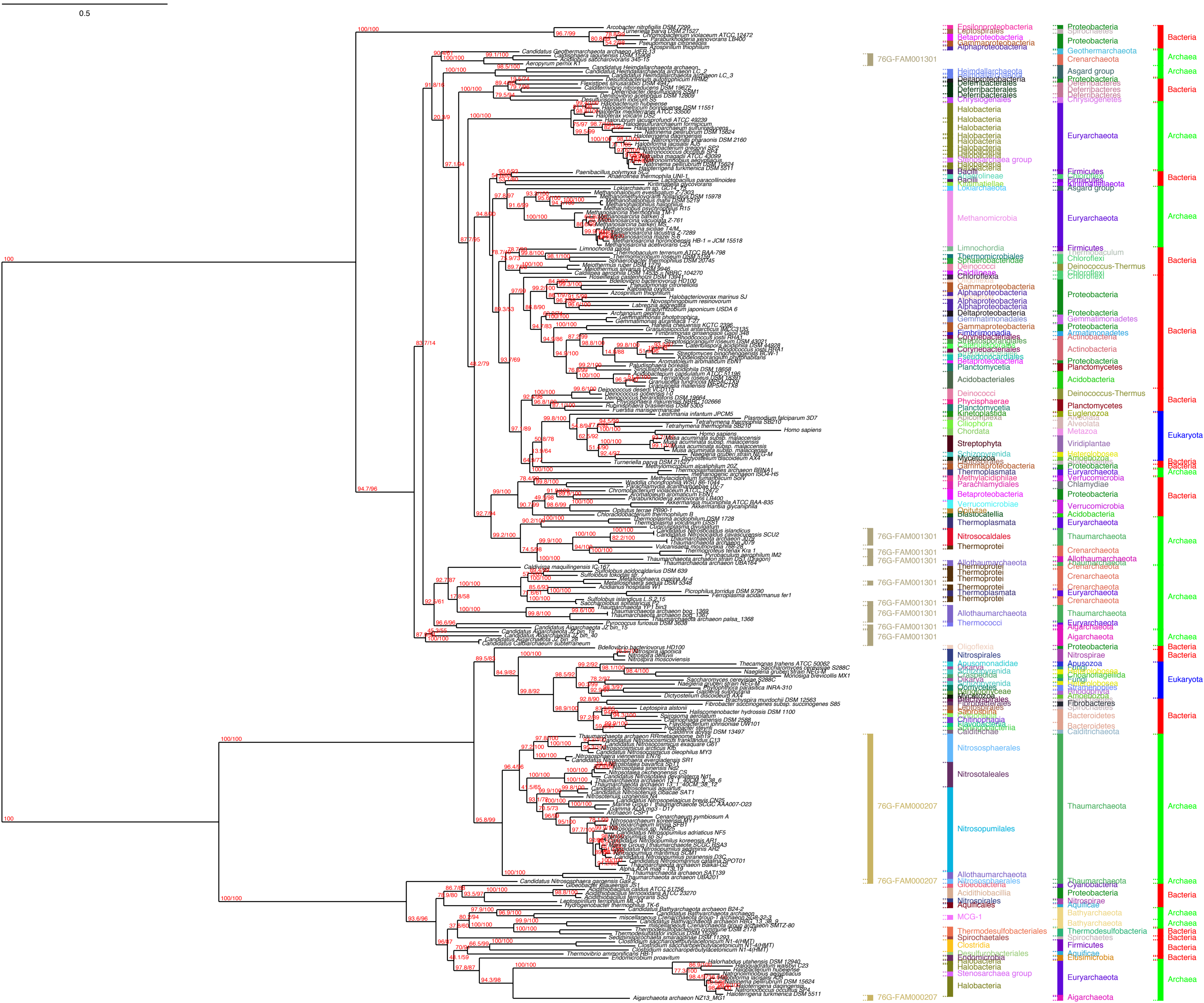

0.5
76G-FAM001301
76G-FAM000207
Epsilonproteobacteria
Betaproteobacteria
Gammaproteobacteria
Alphaproteobacteria
Heimdallarchaeota
Deltaproteobacteria
Deferribacterales
Chrysiogenetes
Halobacteria
Stenosarchaea group
Bacilli
Lokiarchaeota
Methanomicrobia
Limnochordia
Thermomicrobiales
Sphaerobacteridae
Deinococci
Chloroflexia
Gemmatimonadales
Fimbriimonadia
Corynebacteriales
Planctomycetia
Acidobacteriales
Phycisphaerae
Kinetoplastida
Ciliophora
Chordata
Streptophyta
Mycetozoa
Thermoplasmata
Methylacidiphilae
Parachlamydiales
Verrucomicrobiae
Blastocatellia
Nitrosocaldales
Thermoprotei
Allothaumarchaeota
Thermococci
Oligoflexia
Nitrospirales
Apusomonadidae
Dikarya
Saprospiria
Chitinophagia
Flavobacteriia
Caldithrichae
Nitrososphaerales
Nitrosotaleales
Nitrosopumilales
Gloeobacteria
Acidithiobacillia
Aquificae
MCG-1
Thermodesulfobacteriales
Spirochaetales
Clostridia
Desulfurobacteriales
Endomicrobia
Proteobacteria
Spirochaetes
Geothermarchaeota
Crenarchaeota
Asgard group
Euryarchaeota
Firmicutes
Chloroflexi
Kiritimatiellaeota
Thermobaculum
Deinococcus-Thermus
Gemmatimonadetes
Armatimonadetes
Actinobacteria
Planctomycetes
Acidobacteria
Euglenozoa
Alveolata
Metazoa
Viridiplantae
Heterolobosea
Amoebozoa
Verrucomicrobia
Chlamydiae
Thaumarchaeota
Aigarchaeota
Nitrospirae
Apusozoa
Fungi
Choanoflagellida
Stramenopiles
Rhodophyta
Fibrobacteres
Bacteroidetes
Calditrichaeota
Cyanobacteria
Bathyarchaeota
Thermodesulfobacteria
Elusimicrobia
Bacteria
Archaea
Eukaryota

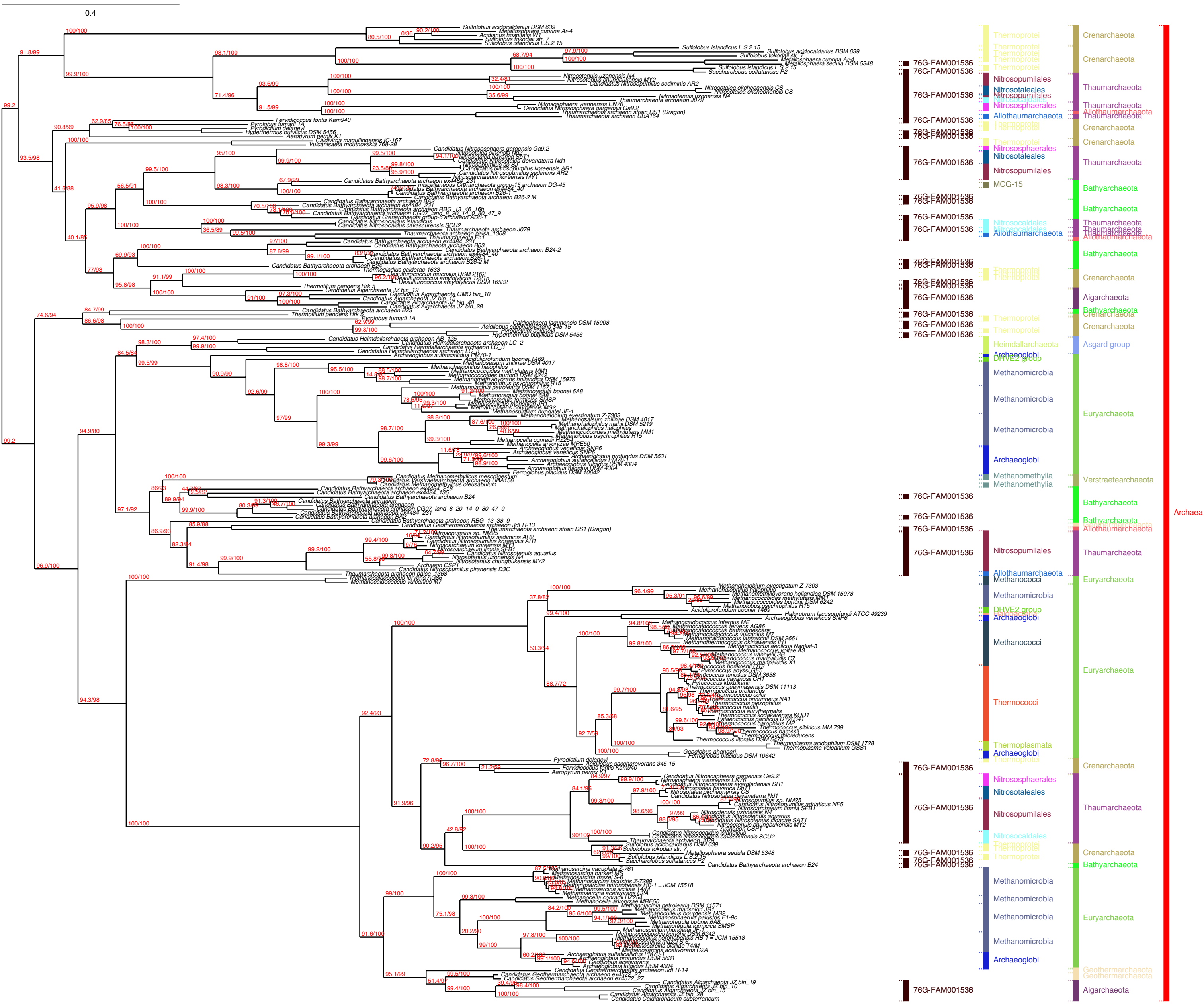
0.4
76G-FAM001536
Thermoprotei
Crenarchaeota
Nitrosopumilales
Nitrosotaleales
Nitrososphaerales
Allothaumarchaeota
Thaumarchaeota
MCG-15
Bathyarchaeota
Nitrosocaldales
Aigarchaeota
Heimdallarchaeota
Archaeoglobi
DHVE2 group
Asgard group
Methanomicrobia
Euryarchaeota
Methanomethylia
Verstraetearchaeota
Methanococci
Thermococci
Thermoplasmata
Geothermarchaeota
Archaea

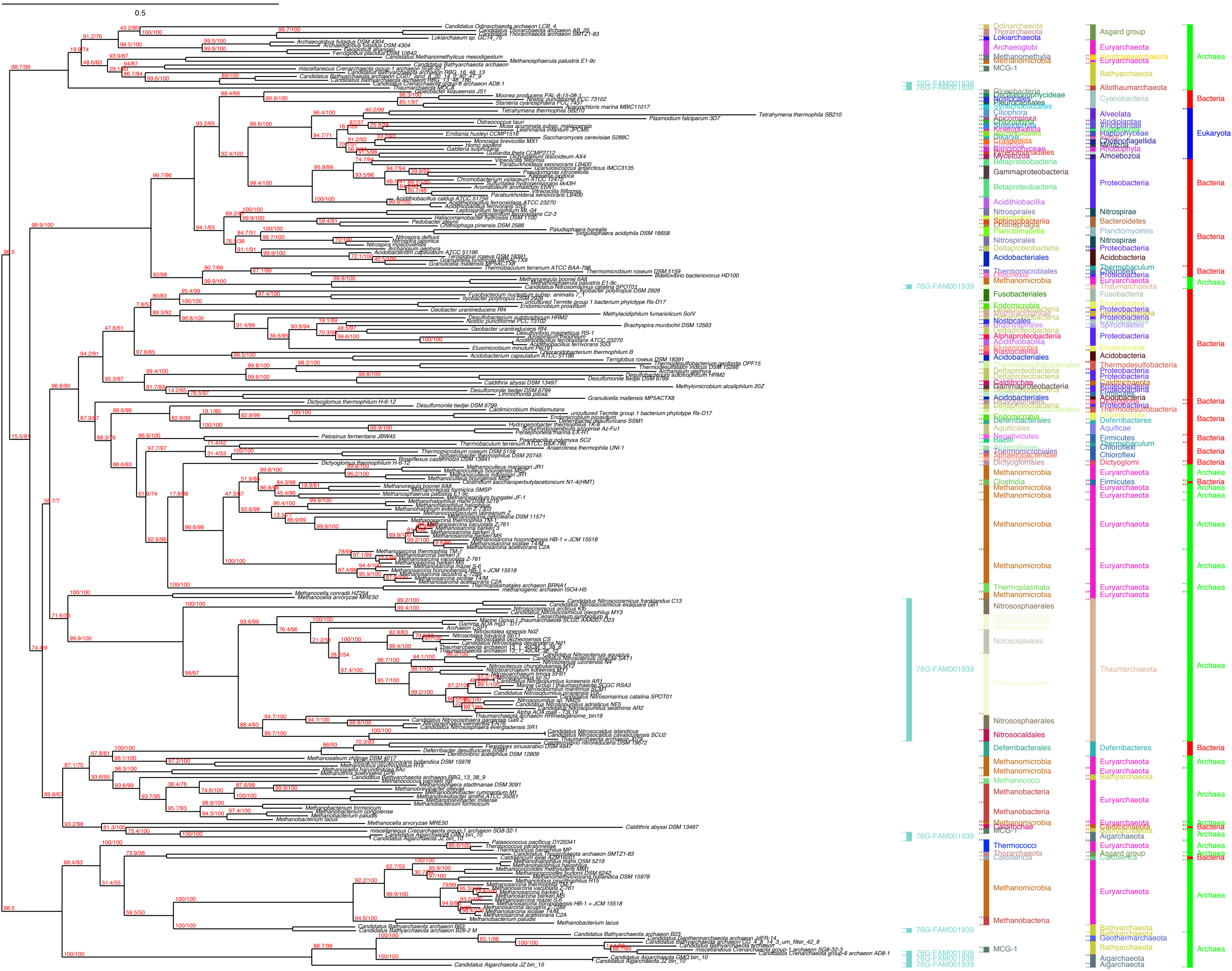
0.5
Archaea
Bacteria
Eukaryota
76G-FAM001939

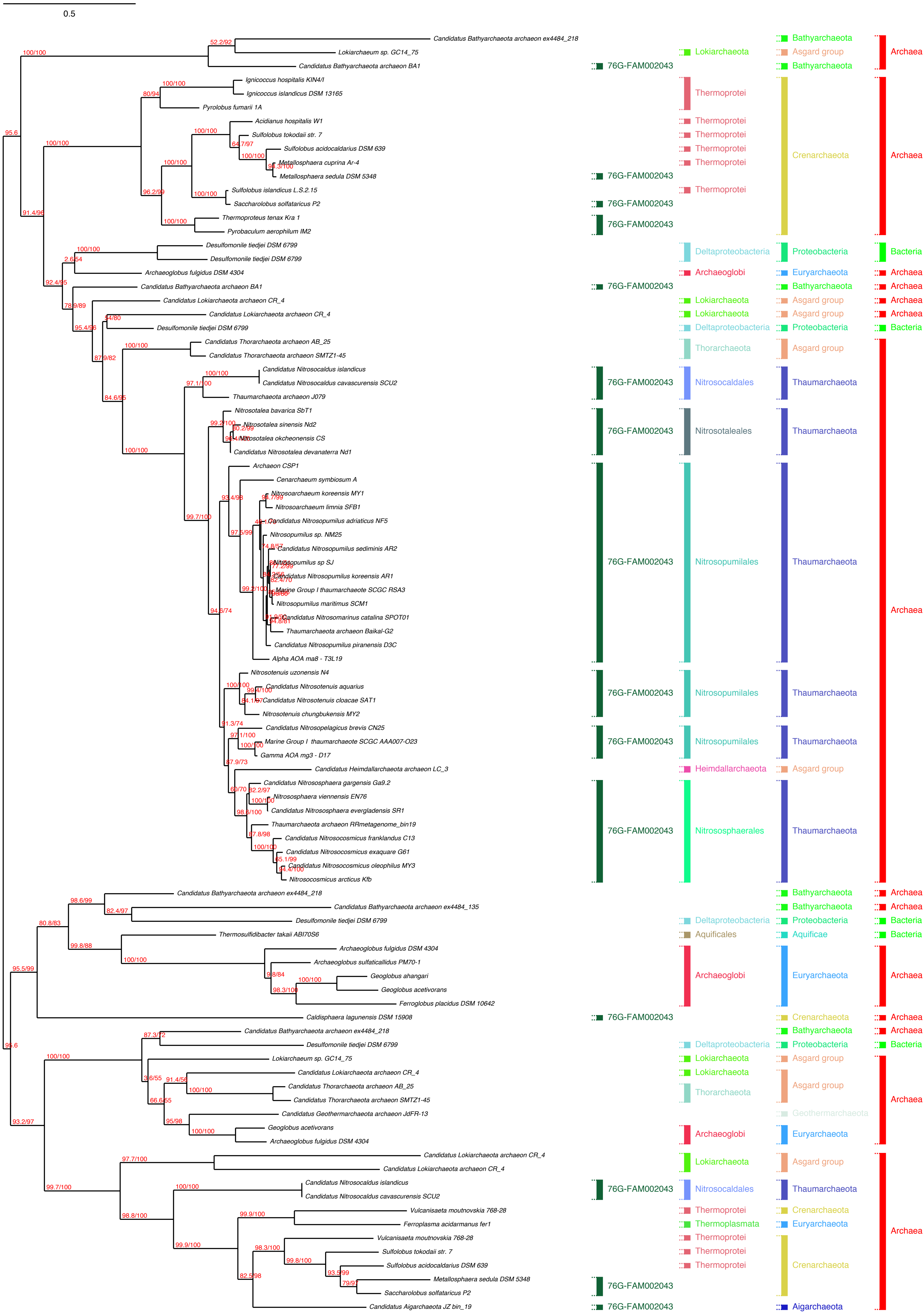
0.5
Candidatus Bathyarchaeota archaeon ex4484_218
Lokiarchaeum sp. GC14_75
Candidatus Bathyarchaeota archaeon BA1
Ignicoccus hospitalis KIN4/I
Ignicoccus islandicus DSM 13165
Pyrolobus fumarii 1A
Acidianus hospitalis W1
Sulfolobus tokodaii str. 7
Sulfolobus acidocaldarius DSM 639
Metallosphaera cuprina Ar-4
Metallosphaera sedula DSM 5348
Sulfolobus islandicus L.S.2.15
Saccharolobus solfataricus P2
Thermoproteus tenax Kra 1
Pyrobaculum aerophilum IM2
Desulfomonile tiedjei DSM 6799
Desulfomonile tiedjei DSM 6799
Archaeoglobus fulgidus DSM 4304
Candidatus Bathyarchaeota archaeon BA1
Candidatus Lokiarchaeota archaeon CR_4
Candidatus Lokiarchaeota archaeon CR_4
Desulfomonile tiedjei DSM 6799
Candidatus Thorarchaeota archaeon AB_25
Candidatus Thorarchaeota archaeon SMTZ1-45
Candidatus Nitrosocaldus islandicus
Candidatus Nitrosocaldus cavascurensis SCU2
Thaumarchaeota archaeon J079
Nitrosotalea bavarica SbT1
Nitrosotalea sinensis Nd2
Nitrosotalea okcheonensis CS
Candidatus Nitrosotalea devanaterra Nd1
Archaeon CSP1
Cenarchaeum symbiosum A
Nitrosoarchaeum koreensis MY1
Nitrosoarchaeum limnia SFB1
Candidatus Nitrosopumilus adriaticus NF5
Nitrosopumilus sp. NM25
Candidatus Nitrosopumilus sediminis AR2
Nitrosopumilus sp SJ
Candidatus Nitrosopumilus koreensis AR1
Marine Group I thaumarchaeote SCGC RSA3
Nitrosopumilus maritimus SCM1
Candidatus Nitrosomarinus catalina SPOT01
Thaumarchaeota archaeon Baikal-G2
Candidatus Nitrosopumilus piranensis D3C
Alpha AOA ma8 - T3L19
Nitrosotenuis uzonensis N4
Candidatus Nitrosotenuis aquarius
Candidatus Nitrosotenuis cloacae SAT1
Nitrosotenuis chungbukensis MY2
Candidatus Nitrosopelagicus brevis CN25
Marine Group I thaumarchaeote SCGC AAA007-O23
Gamma AOA mg3 - D17
Candidatus Heimdallarchaeota archaeon LC_3
Candidatus Nitrososphaera gargensis Ga9.2
Nitrososphaera viennensis EN76
Candidatus Nitrososphaera evergladensis SR1
Thaumarchaeota archaeon RRmetagenome_bin19
Candidatus Nitrosocosmicus franklandus C13
Candidatus Nitrosocosmicus exaquare G61
Candidatus Nitrosocosmicus oleophilus MY3
Nitrosocosmicus arcticus Kfb
Candidatus Bathyarchaeota archaeon ex4484_218
Candidatus Bathyarchaeota archaeon ex4484_135
Desulfomonile tiedjei DSM 6799
Thermosulfidibacter takaii ABI70S6
Archaeoglobus fulgidus DSM 4304
Archaeoglobus sulfaticallidus PM70-1
Geoglobus ahangari
Geoglobus acetivorans
Ferroglobus placidus DSM 10642
Caldisphaera lagunensis DSM 15908
Candidatus Bathyarchaeota archaeon ex4484_218
Desulfomonile tiedjei DSM 6799
Lokiarchaeum sp. GC14_75
Candidatus Lokiarchaeota archaeon CR_4
Candidatus Thorarchaeota archaeon AB_25
Candidatus Thorarchaeota archaeon SMTZ1-45
Candidatus Geothermarchaeota archaeon JdFR-13
Geoglobus acetivorans
Archaeoglobus fulgidus DSM 4304
Candidatus Lokiarchaeota archaeon CR_4
Candidatus Lokiarchaeota archaeon CR_4
Candidatus Nitrosocaldus islandicus
Candidatus Nitrosocaldus cavascurensis SCU2
Vulcanisaeta moutnovskia 768-28
Ferroplasma acidarmanus fer1
Vulcanisaeta moutnovskia 768-28
Sulfolobus tokodaii str. 7
Sulfolobus acidocaldarius DSM 639
Metallosphaera sedula DSM 5348
Saccharolobus solfataricus P2
Candidatus Aigarchaeota archaeon JZ bin_19
76G-FAM002043
Lokiarchaeota
Thermoprotei
Deltaproteobacteria
Archaeoglobi
Thorarchaeota
Nitrosocaldales
Nitrosotaleales
Nitrosopumilales
Heimdallarchaeota
Nitrososphaerales
Aquificales
Thermoplasmata
Bathyarchaeota
Asgard group
Crenarchaeota
Proteobacteria
Euryarchaeota
Thaumarchaeota
Aquificae
Geothermarchaeota
Aigarchaeota
Archaea
Bacteria

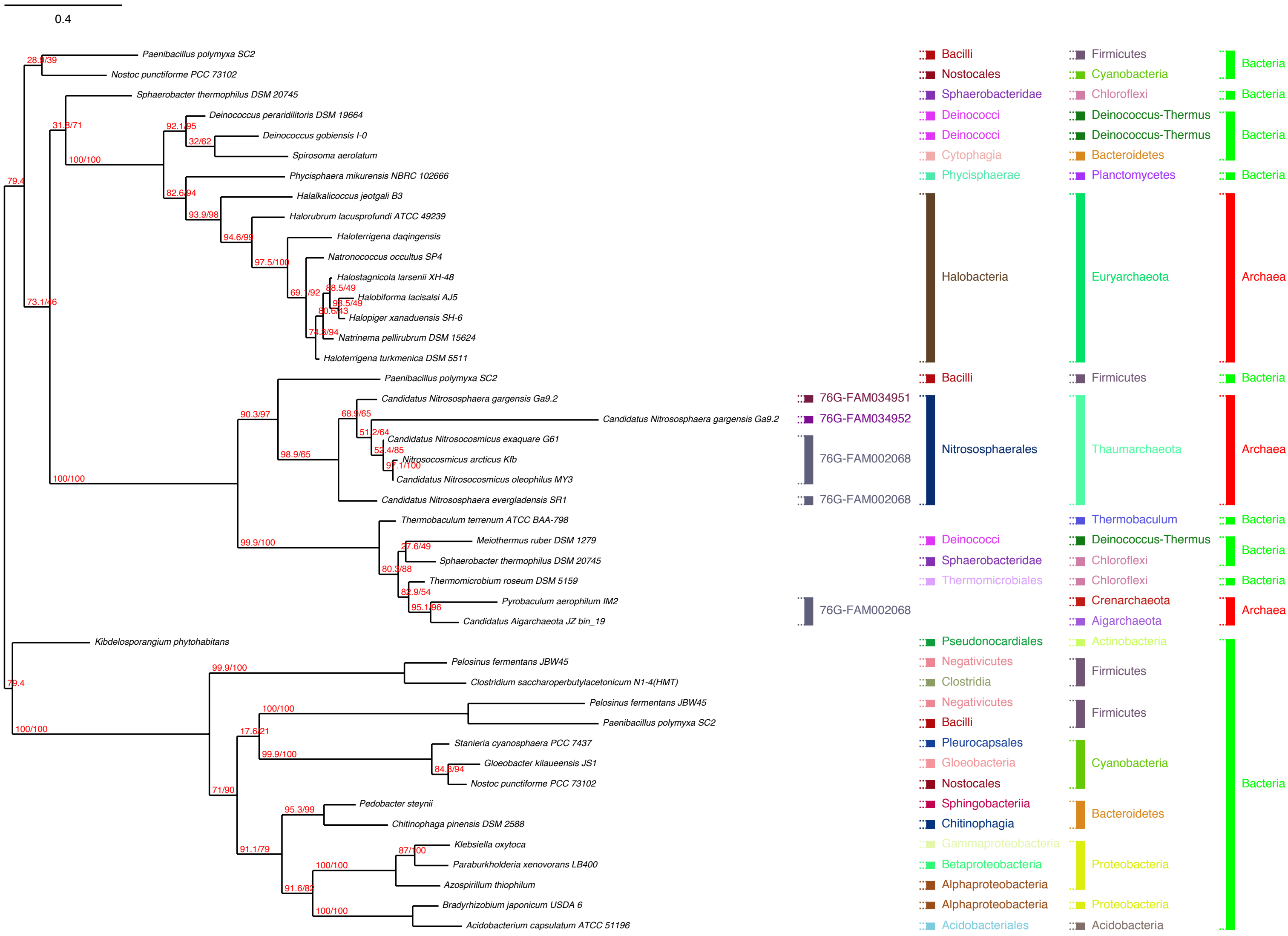
0.4
Paenibacillus polymyxa SC2
Nostoc punctiforme PCC 73102
Sphaerobacter thermophilus DSM 20745
Deinococcus peraridilitoris DSM 19664
Deinococcus gobiensis I-0
Spirosoma aerolatum
Phycisphaera mikurensis NBRC 102666
Halalkalicoccus jeotgali B3
Halorubrum lacusprofundi ATCC 49239
Haloterrigena daqingensis
Natronococcus occultus SP4
Halostagnicola larsenii XH-48
Halobiforma lacisalsi AJ5
Halopiger xanaduensis SH-6
Natrinema pellirubrum DSM 15624
Haloterrigena turkmenica DSM 5511
Paenibacillus polymyxa SC2
Candidatus Nitrososphaera gargensis Ga9.2
Candidatus Nitrososphaera gargensis Ga9.2
Candidatus Nitrosocosmicus exaquare G61
Nitrosocosmicus arcticus Kfb
Candidatus Nitrosocosmicus oleophilus MY3
Candidatus Nitrososphaera evergladensis SR1
Thermobaculum terrenum ATCC BAA-798
Meiothermus ruber DSM 1279
Sphaerobacter thermophilus DSM 20745
Thermomicrobium roseum DSM 5159
Pyrobaculum aerophilum IM2
Candidatus Aigarchaeota JZ bin_19
Kibdelosporangium phytohabitans
Pelosinus fermentans JBW45
Clostridium saccharoperbutylacetonicum N1-4(HMT)
Pelosinus fermentans JBW45
Paenibacillus polymyxa SC2
Stanieria cyanosphaera PCC 7437
Gloeobacter kilaueensis JS1
Nostoc punctiforme PCC 73102
Pedobacter steynii
Chitinophaga pinensis DSM 2588
Klebsiella oxytoca
Paraburkholderia xenovorans LB400
Azospirillum thiophilum
Bradyrhizobium japonicum USDA 6
Acidobacterium capsulatum ATCC 51196
28.9/39
31.8/71
92.1/95
32/62
100/100
79.4
82.6/94
93.9/98
94.6/99
97.5/100
69.1/92
88.5/49
95.5/49
80.6/43
74.3/94
73.1/46
90.3/97
68.9/65
51.2/64
52.4/85
97.1/100
98.9/65
100/100
99.9/100
27.6/49
80.3/88
82.9/54
95.1/96
79.4
99.9/100
100/100
100/100
17.6/21
99.9/100
84.8/94
71/90
95.3/99
87/100
91.1/79
100/100
91.6/82
100/100
76G-FAM034951
76G-FAM034952
76G-FAM002068
76G-FAM002068
76G-FAM002068
Bacilli
Nostocales
Sphaerobacteridae
Deinococci
Deinococci
Cytophagia
Phycisphaerae
Halobacteria
Bacilli
Nitrososphaerales
Deinococci
Sphaerobacteridae
Thermomicrobiales
Pseudonocardiales
Negativicutes
Clostridia
Negativicutes
Bacilli
Pleurocapsales
Gloeobacteria
Nostocales
Sphingobacteriia
Chitinophagia
Gammaproteobacteria
Betaproteobacteria
Alphaproteobacteria
Alphaproteobacteria
Acidobacteriales
Firmicutes
Cyanobacteria
Chloroflexi
Deinococcus-Thermus
Deinococcus-Thermus
Bacteroidetes
Planctomycetes
Euryarchaeota
Firmicutes
Thaumarchaeota
Thermobaculum
Deinococcus-Thermus
Chloroflexi
Chloroflexi
Crenarchaeota
Aigarchaeota
Actinobacteria
Firmicutes
Firmicutes
Cyanobacteria
Bacteroidetes
Proteobacteria
Proteobacteria
Acidobacteria
Bacteria
Bacteria
Bacteria
Bacteria
Archaea
Bacteria
Archaea
Bacteria
Bacteria
Bacteria
Archaea
Bacteria

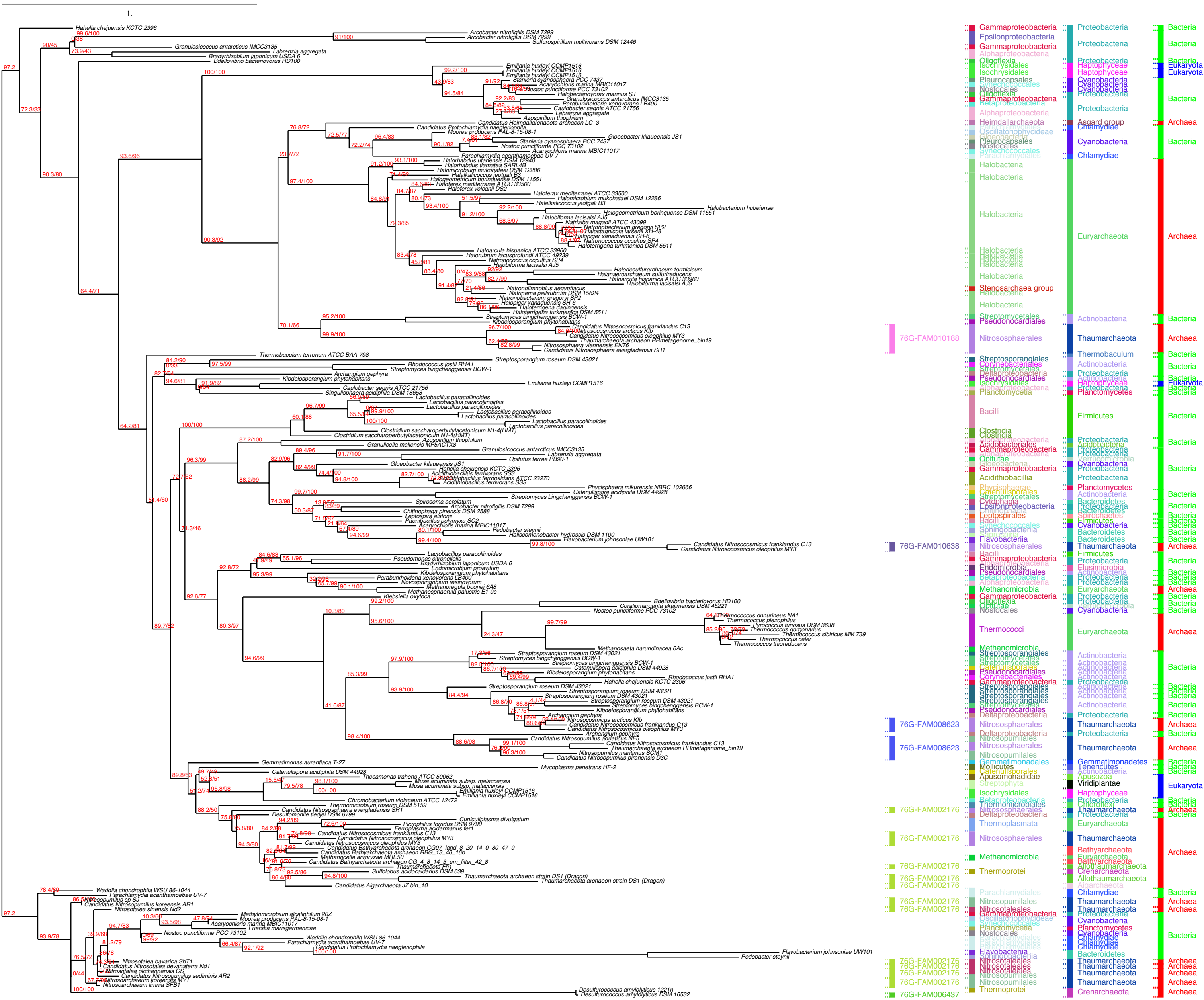

1.
Hahella chejuensis KCTC 2396
Arcobacter nitrofigilis DSM 7299
Granulosicoccus antarcticus IMCC3135
Bradyrhizobium japonicum USDA 6
Bdellovibrio bacteriovorus HD100
Thermobaculum terrenum ATCC BAA-798
Gemmatimonas aurantiaca T-27
Mycoplasma penetrans HF-2
Waddlia chondrophila WSU 86-1044
Flavobacterium johnsoniae UW101
76G-FAM010188
76G-FAM010638
76G-FAM008623
76G-FAM002176
76G-FAM006437
Gammaproteobacteria
Epsilonproteobacteria
Halobacteria
Thermococci
Nitrososphaerales
Nitrosopumilales
Proteobacteria
Cyanobacteria
Euryarchaeota
Thaumarchaeota
Actinobacteria
Firmicutes
Bacteria
Archaea
Eukaryota

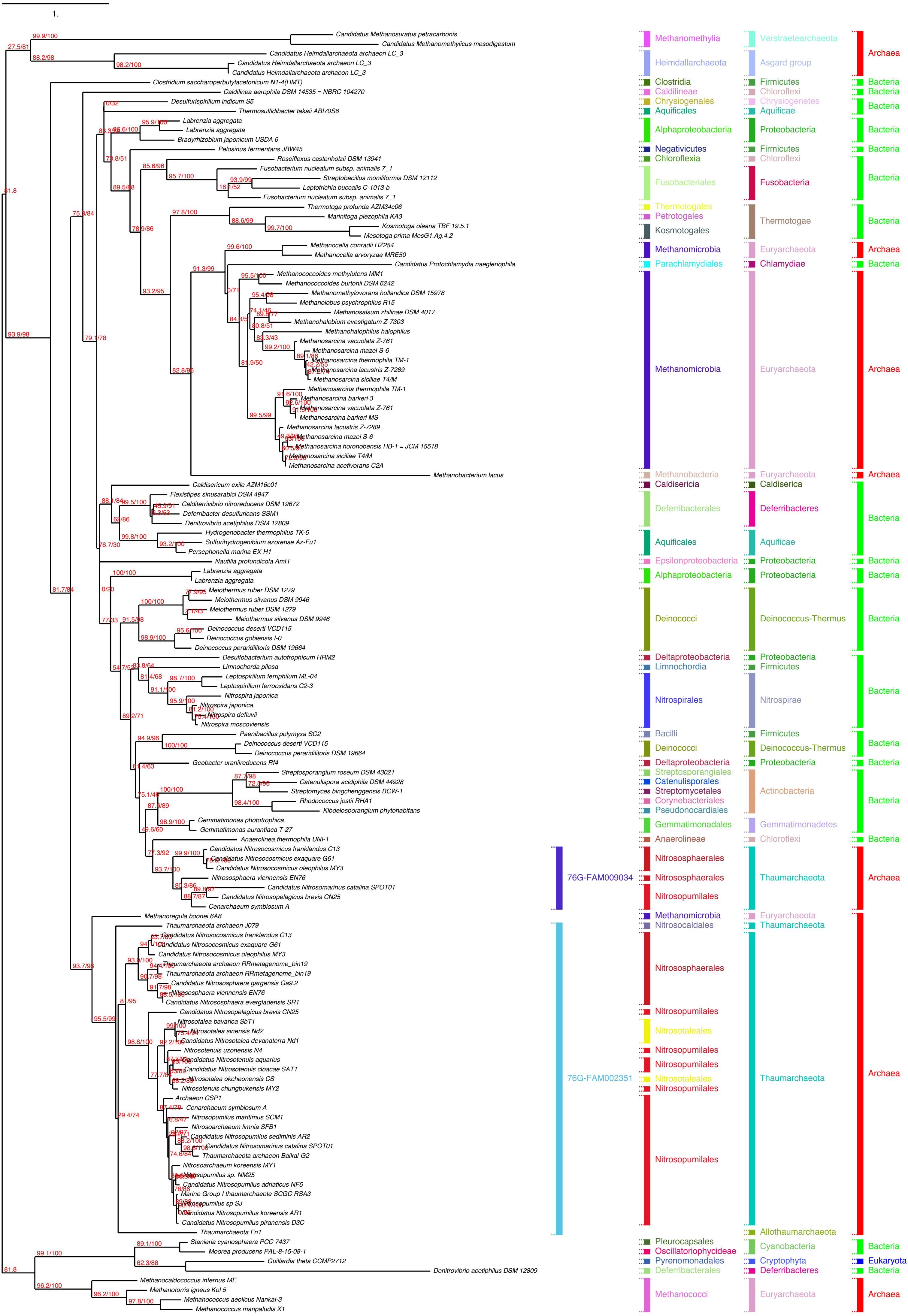

1.
Candidatus Methanosuratus petracarbonis
Candidatus Methanomethylicus mesodigestum
Candidatus Heimdallarchaeota archaeon LC_3
Candidatus Heimdallarchaeota archaeon LC_3
Candidatus Heimdallarchaeota archaeon LC_3
Clostridium saccharoperbutylacetonicum N1-4(HMT)
Caldilinea aerophila DSM 14535 = NBRC 104270
Desulfurispirillum indicum S5
Thermosulfidibacter takaii ABI70S6
Labrenzia aggregata
Labrenzia aggregata
Bradyrhizobium japonicum USDA 6
Pelosinus fermentans JBW45
Roseiflexus castenholzii DSM 13941
Fusobacterium nucleatum subsp. animalis 7_1
Streptobacillus moniliformis DSM 12112
Leptotrichia buccalis C-1013-b
Fusobacterium nucleatum subsp. animalis 7_1
Thermotoga profunda AZM34c06
Marinitoga piezophila KA3
Kosmotoga olearia TBF 19.5.1
Mesotoga prima MesG1.Ag.4.2
Methanocella conradii HZ254
Methanocella arvoryzae MRE50
Candidatus Protochlamydia naegleriophila
Methanococcoides methylutens MM1
Methanococcoides burtonii DSM 6242
Methanomethylovorans hollandica DSM 15978
Methanolobus psychrophilus R15
Methanosalsum zhilinae DSM 4017
Methanohalobium evestigatum Z-7303
Methanohalophilus halophilus
Methanosarcina vacuolata Z-761
Methanosarcina mazei S-6
Methanosarcina thermophila TM-1
Methanosarcina lacustris Z-7289
Methanosarcina siciliae T4/M
Methanosarcina thermophila TM-1
Methanosarcina barkeri 3
Methanosarcina vacuolata Z-761
Methanosarcina barkeri MS
Methanosarcina lacustris Z-7289
Methanosarcina mazei S-6
Methanosarcina horonobensis HB-1 = JCM 15518
Methanosarcina siciliae T4/M
Methanosarcina acetivorans C2A
Methanobacterium lacus
Caldisericum exile AZM16c01
Flexistipes sinusarabici DSM 4947
Calditerrivibrio nitroreducens DSM 19672
Deferribacter desulfuricans SSM1
Denitrovibrio acetiphilus DSM 12809
Hydrogenobacter thermophilus TK-6
Sulfurihydrogenibium azorense Az-Fu1
Persephonella marina EX-H1
Nautilia profundicola AmH
Labrenzia aggregata
Labrenzia aggregata
Meiothermus ruber DSM 1279
Meiothermus silvanus DSM 9946
Meiothermus ruber DSM 1279
Meiothermus silvanus DSM 9946
Deinococcus deserti VCD115
Deinococcus gobiensis I-0
Deinococcus peraridilitoris DSM 19664
Desulfobacterium autotrophicum HRM2
Limnochorda pilosa
Leptospirillum ferriphilum ML-04
Leptospirillum ferrooxidans C2-3
Nitrospira japonica
Nitrospira japonica
Nitrospira defluvii
Nitrospira moscoviensis
Paenibacillus polymyxa SC2
Deinococcus deserti VCD115
Deinococcus peraridilitoris DSM 19664
Geobacter uraniireducens Rf4
Streptosporangium roseum DSM 43021
Catenulispora acidiphila DSM 44928
Streptomyces bingchenggensis BCW-1
Rhodococcus jostii RHA1
Kibdelosporangium phytohabitans
Gemmatimonas phototrophica
Gemmatimonas aurantiaca T-27
Anaerolinea thermophila UNI-1
Candidatus Nitrosocosmicus franklandus C13
Candidatus Nitrosocosmicus exaquare G61
Candidatus Nitrosocosmicus oleophilus MY3
Nitrososphaera viennensis EN76
Candidatus Nitrosomarinus catalina SPOT01
Candidatus Nitrosopelagicus brevis CN25
Cenarchaeum symbiosum A
Methanoregula boonei 6A8
Thaumarchaeota archaeon J079
Candidatus Nitrosocosmicus franklandus C13
Candidatus Nitrosocosmicus exaquare G61
Candidatus Nitrosocosmicus oleophilus MY3
Thaumarchaeota archaeon RRmetagenome_bin19
Thaumarchaeota archaeon RRmetagenome_bin19
Candidatus Nitrososphaera gargensis Ga9.2
Nitrososphaera viennensis EN76
Candidatus Nitrososphaera evergladensis SR1
Candidatus Nitrosopelagicus brevis CN25
Nitrosotalea bavarica SbT1
Nitrosotalea sinensis Nd2
Candidatus Nitrosotalea devanaterra Nd1
Nitrosotenuis uzonensis N4
Candidatus Nitrosotenuis aquarius
Candidatus Nitrosotenuis cloacae SAT1
Nitrosotalea okcheonensis CS
Nitrosotenuis chungbukensis MY2
Archaeon CSP1
Cenarchaeum symbiosum A
Nitrosopumilus maritimus SCM1
Nitrosoarchaeum limnia SFB1
Candidatus Nitrosopumilus sediminis AR2
Candidatus Nitrosomarinus catalina SPOT01
Thaumarchaeota archaeon Baikal-G2
Nitrosoarchaeum koreensis MY1
Nitrosopumilus sp. NM25
Candidatus Nitrosopumilus adriaticus NF5
Marine Group I thaumarchaeote SCGC RSA3
Nitrosopumilus sp SJ
Candidatus Nitrosopumilus koreensis AR1
Candidatus Nitrosopumilus piranensis D3C
Thaumarchaeota Fn1
Stanieria cyanosphaera PCC 7437
Moorea producens PAL-8-15-08-1
Guillardia theta CCMP2712
Denitrovibrio acetiphilus DSM 12809
Methanocaldococcus infernus ME
Methanotorris igneus Kol 5
Methanococcus aeolicus Nankai-3
Methanococcus maripaludis X1
76G-FAM009034
76G-FAM002351
Methanomethylia
Heimdallarchaeota
Clostridia
Caldilineae
Chrysiogenales
Aquificales
Alphaproteobacteria
Negativicutes
Chloroflexia
Fusobacteriales
Thermotogales
Petrotogales
Kosmotogales
Methanomicrobia
Parachlamydiales
Methanomicrobia
Methanobacteria
Caldisericia
Deferribacterales
Aquificales
Epsilonproteobacteria
Alphaproteobacteria
Deinococci
Deltaproteobacteria
Limnochordia
Nitrospirales
Bacilli
Deinococci
Deltaproteobacteria
Streptosporangiales
Catenulisporales
Streptomycetales
Corynebacteriales
Pseudonocardiales
Gemmatimonadales
Anaerolineae
Nitrososphaerales
Nitrososphaerales
Nitrosopumilales
Methanomicrobia
Nitrosocaldales
Nitrososphaerales
Nitrosopumilales
Nitrosotaleales
Nitrosopumilales
Nitrosopumilales
Nitrosotaleales
Nitrosopumilales
Nitrosopumilales
Pleurocapsales
Oscillatoriophycideae
Pyrenomonadales
Deferribacterales
Methanococci
Verstraetearchaeota
Asgard group
Firmicutes
Chloroflexi
Chrysiogenetes
Aquificae
Proteobacteria
Firmicutes
Chloroflexi
Fusobacteria
Thermotogae
Euryarchaeota
Chlamydiae
Euryarchaeota
Euryarchaeota
Caldiserica
Deferribacteres
Aquificae
Proteobacteria
Proteobacteria
Deinococcus-Thermus
Proteobacteria
Firmicutes
Nitrospirae
Firmicutes
Deinococcus-Thermus
Proteobacteria
Actinobacteria
Gemmatimonadetes
Chloroflexi
Thaumarchaeota
Euryarchaeota
Thaumarchaeota
Thaumarchaeota
Allothaumarchaeota
Cyanobacteria
Cryptophyta
Deferribacteres
Euryarchaeota
Archaea
Bacteria
Bacteria
Bacteria
Bacteria
Bacteria
Bacteria
Bacteria
Archaea
Bacteria
Archaea
Archaea
Bacteria
Bacteria
Bacteria
Bacteria
Bacteria
Bacteria
Bacteria
Bacteria
Bacteria
Bacteria
Archaea
Archaea
Bacteria
Eukaryota
Bacteria
Archaea

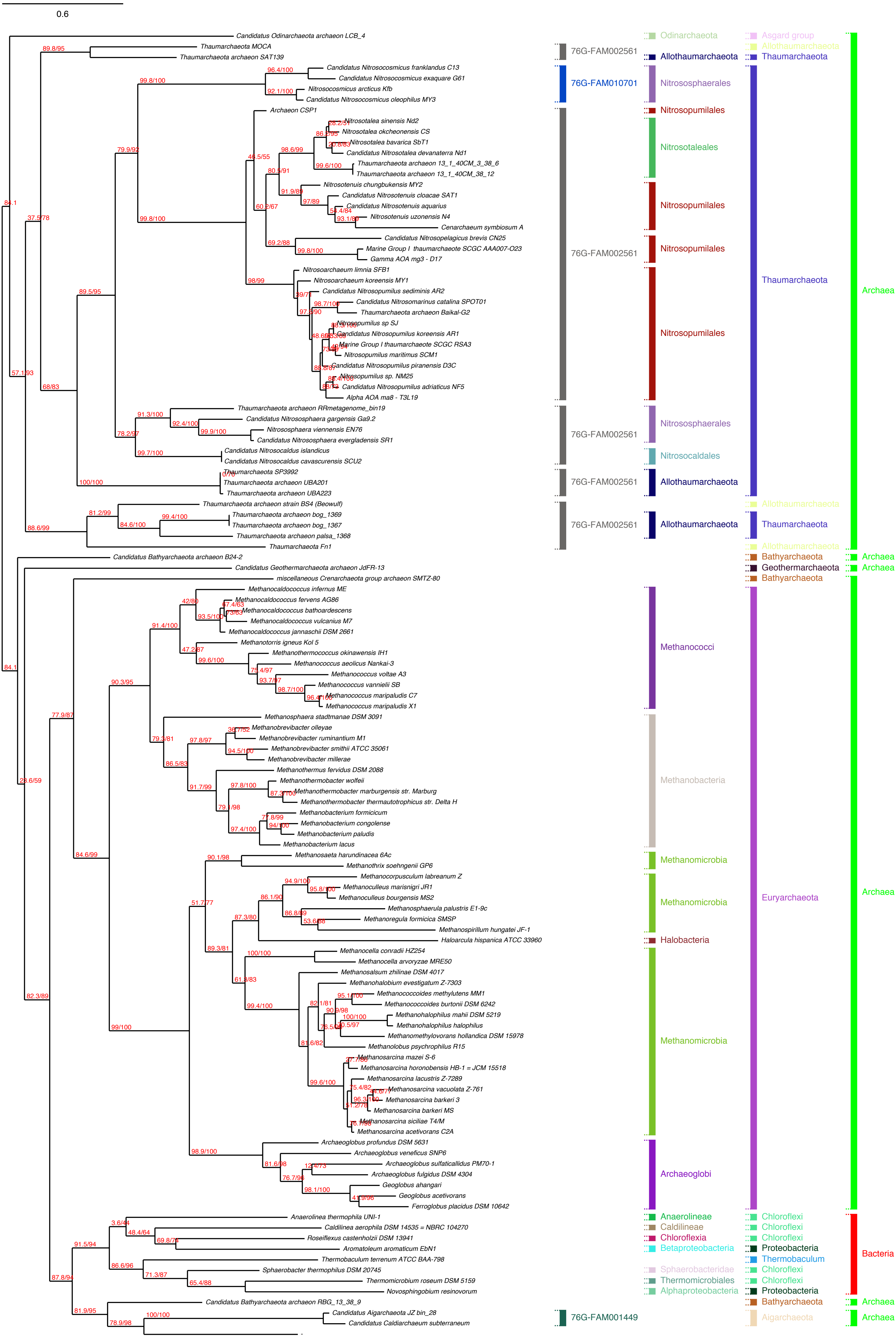

0.6
Candidatus Odinarchaeota archaeon LCB_4
Thaumarchaeota MOCA
Thaumarchaeota archaeon SAT139
Candidatus Nitrosocosmicus franklandus C13
Candidatus Nitrosocosmicus exaquare G61
Nitrosocosmicus arcticus Kfb
Candidatus Nitrosocosmicus oleophilus MY3
Archaeon CSP1
Nitrosotalea sinensis Nd2
Nitrosotalea okcheonensis CS
Nitrosotalea bavarica SbT1
Candidatus Nitrosotalea devanaterra Nd1
Thaumarchaeota archaeon 13_1_40CM_3_38_6
Thaumarchaeota archaeon 13_1_40CM_38_12
Nitrosotenuis chungbukensis MY2
Candidatus Nitrosotenuis cloacae SAT1
Candidatus Nitrosotenuis aquarius
Nitrosotenuis uzonensis N4
Cenarchaeum symbiosum A
Candidatus Nitrosopelagicus brevis CN25
Marine Group I thaumarchaeote SCGC AAA007-O23
Gamma AOA mg3 - D17
Nitrosoarchaeum limnia SFB1
Nitrosoarchaeum koreensis MY1
Candidatus Nitrosopumilus sediminis AR2
Candidatus Nitrosomarinus catalina SPOT01
Thaumarchaeota archaeon Baikal-G2
Nitrosopumilus sp SJ
Candidatus Nitrosopumilus koreensis AR1
Marine Group I thaumarchaeote SCGC RSA3
Nitrosopumilus maritimus SCM1
Candidatus Nitrosopumilus piranensis D3C
Nitrosopumilus sp. NM25
Candidatus Nitrosopumilus adriaticus NF5
Alpha AOA ma8 - T3L19
Thaumarchaeota archaeon RRmetagenome_bin19
Candidatus Nitrososphaera gargensis Ga9.2
Nitrososphaera viennensis EN76
Candidatus Nitrososphaera evergladensis SR1
Candidatus Nitrosocaldus islandicus
Candidatus Nitrosocaldus cavascurensis SCU2
Thaumarchaeota SP3992
Thaumarchaeota archaeon UBA201
Thaumarchaeota archaeon UBA223
Thaumarchaeota archaeon strain BS4 (Beowulf)
Thaumarchaeota archaeon bog_1369
Thaumarchaeota archaeon bog_1367
Thaumarchaeota archaeon palsa_1368
Thaumarchaeota Fn1
Candidatus Bathyarchaeota archaeon B24-2
Candidatus Geothermarchaeota archaeon JdFR-13
miscellaneous Crenarchaeota group archaeon SMTZ-80
Methanocaldococcus infernus ME
Methanocaldococcus fervens AG86
Methanocaldococcus bathoardescens
Methanocaldococcus vulcanius M7
Methanocaldococcus jannaschii DSM 2661
Methanotorris igneus Kol 5
Methanothermococcus okinawensis IH1
Methanococcus aeolicus Nankai-3
Methanococcus voltae A3
Methanococcus vannielii SB
Methanococcus maripaludis C7
Methanococcus maripaludis X1
Methanosphaera stadtmanae DSM 3091
Methanobrevibacter olleyae
Methanobrevibacter ruminantium M1
Methanobrevibacter smithii ATCC 35061
Methanobrevibacter millerae
Methanothermus fervidus DSM 2088
Methanothermobacter wolfeii
Methanothermobacter marburgensis str. Marburg
Methanothermobacter thermautotrophicus str. Delta H
Methanobacterium formicicum
Methanobacterium congolense
Methanobacterium paludis
Methanobacterium lacus
Methanosaeta harundinacea 6Ac
Methanothrix soehngenii GP6
Methanocorpusculum labreanum Z
Methanoculleus marisnigri JR1
Methanoculleus bourgensis MS2
Methanosphaerula palustris E1-9c
Methanoregula formicica SMSP
Methanospirillum hungatei JF-1
Haloarcula hispanica ATCC 33960
Methanocella conradii HZ254
Methanocella arvoryzae MRE50
Methanosalsum zhilinae DSM 4017
Methanohalobium evestigatum Z-7303
Methanococcoides methylutens MM1
Methanococcoides burtonii DSM 6242
Methanohalophilus mahii DSM 5219
Methanohalophilus halophilus
Methanomethylovorans hollandica DSM 15978
Methanolobus psychrophilus R15
Methanosarcina mazei S-6
Methanosarcina horonobensis HB-1 = JCM 15518
Methanosarcina lacustris Z-7289
Methanosarcina vacuolata Z-761
Methanosarcina barkeri 3
Methanosarcina barkeri MS
Methanosarcina siciliae T4/M
Methanosarcina acetivorans C2A
Archaeoglobus profundus DSM 5631
Archaeoglobus veneficus SNP6
Archaeoglobus sulfaticallidus PM70-1
Archaeoglobus fulgidus DSM 4304
Geoglobus ahangari
Geoglobus acetivorans
Ferroglobus placidus DSM 10642
Anaerolinea thermophila UNI-1
Caldilinea aerophila DSM 14535 = NBRC 104270
Roseiflexus castenholzii DSM 13941
Aromatoleum aromaticum EbN1
Thermobaculum terrenum ATCC BAA-798
Sphaerobacter thermophilus DSM 20745
Thermomicrobium roseum DSM 5159
Novosphingobium resinovorum
Candidatus Bathyarchaeota archaeon RBG_13_38_9
Candidatus Aigarchaeota JZ bin_28
Candidatus Caldiarchaeum subterraneum
76G-FAM002561
76G-FAM010701
76G-FAM002561
76G-FAM002561
76G-FAM002561
76G-FAM002561
76G-FAM001449
Odinarchaeota
Allothaumarchaeota
Nitrososphaerales
Nitrosopumilales
Nitrosotaleales
Nitrosopumilales
Nitrosopumilales
Nitrosopumilales
Nitrososphaerales
Nitrosocaldales
Allothaumarchaeota
Allothaumarchaeota
Methanococci
Methanobacteria
Methanomicrobia
Methanomicrobia
Halobacteria
Methanomicrobia
Archaeoglobi
Anaerolineae
Caldilineae
Chloroflexia
Betaproteobacteria
Sphaerobacteridae
Thermomicrobiales
Alphaproteobacteria
Asgard group
Allothaumarchaeota
Thaumarchaeota
Thaumarchaeota
Allothaumarchaeota
Thaumarchaeota
Allothaumarchaeota
Bathyarchaeota
Geothermarchaeota
Bathyarchaeota
Euryarchaeota
Chloroflexi
Chloroflexi
Chloroflexi
Proteobacteria
Thermobaculum
Chloroflexi
Chloroflexi
Proteobacteria
Bathyarchaeota
Aigarchaeota
Archaea
Archaea
Archaea
Archaea
Bacteria
Archaea
Archaea

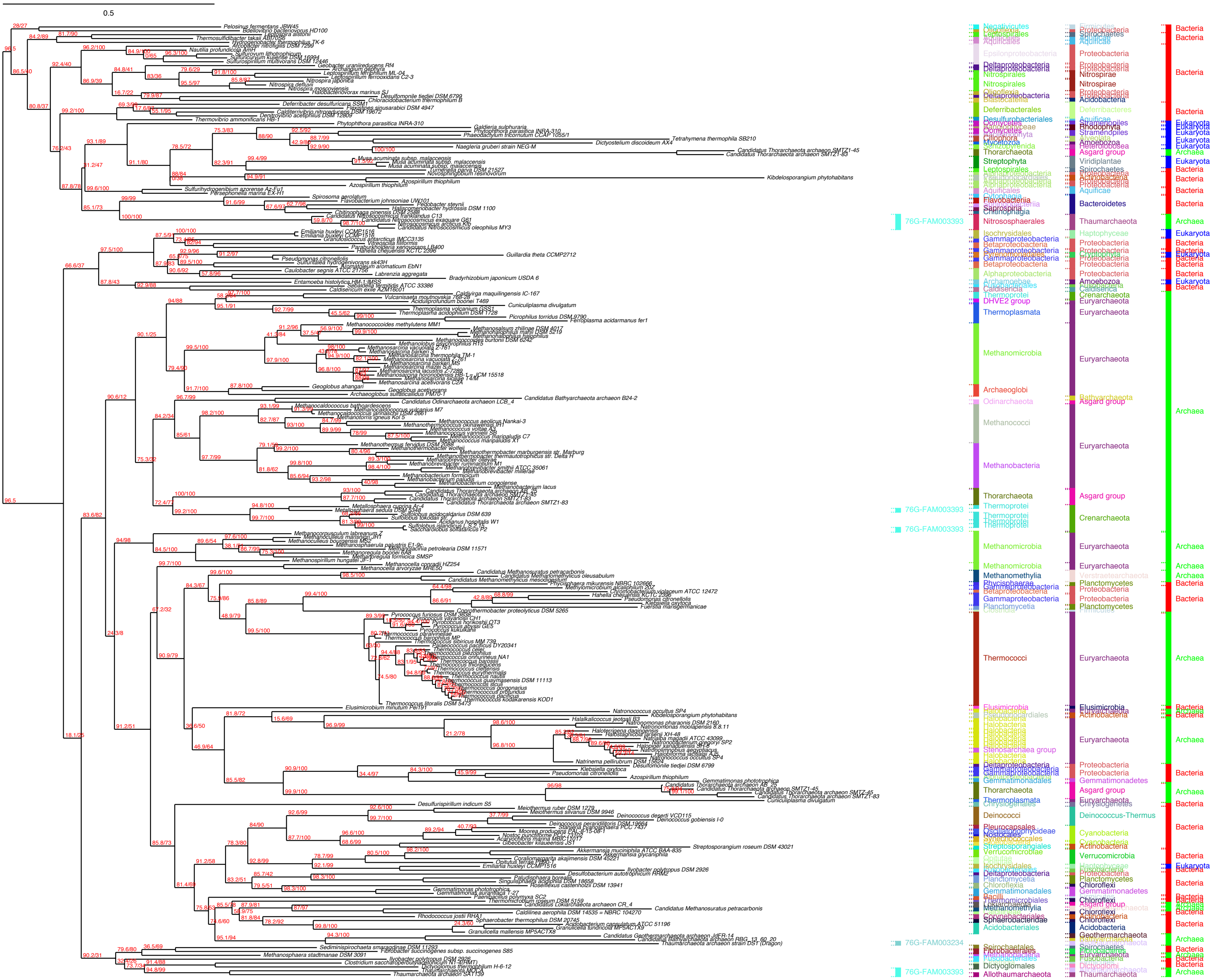

0.5
76G-FAM003393
76G-FAM003393
76G-FAM003393
76G-FAM003234
76G-FAM003393
Negativicutes
Leptospirales
Aquificales
Epsilonproteobacteria
Deltaproteobacteria
Nitrospirales
Deferribacterales
Desulfurobacteriales
Oomycetes
Ciliophora
Mycetozoa
Thorarchaeota
Streptophyta
Leptospirales
Aquificales
Flavobacteriia
Saprospiria
Chitinophagia
Nitrososphaerales
Isochrysidales
Gammaproteobacteria
Betaproteobacteria
Alphaproteobacteria
Archamoebae
Caldisericia
Thermoprotei
DHVE2 group
Thermoplasmata
Methanomicrobia
Archaeoglobi
Odinarchaeota
Methanococci
Methanobacteria
Thorarchaeota
Thermoprotei
Methanomicrobia
Methanomethylia
Planctomycetia
Thermococci
Elusimicrobia
Stenosarchaea group
Halobacteria
Gemmatimonadales
Thermoplasmata
Deinococci
Nostocales
Verrucomicrobiae
Acidobacteriales
Spirochaetales
Dictyoglomales
Allothaumarchaeota
Proteobacteria
Spirochaetes
Aquificae
Nitrospirae
Acidobacteria
Deferribacteres
Stramenopiles
Rhodophyta
Amoebozoa
Heterolobosea
Asgard group
Viridiplantae
Actinobacteria
Bacteroidetes
Thaumarchaeota
Haptophyceae
Cryptophyta
Crenarchaeota
Euryarchaeota
Bathyarchaeota
Planctomycetes
Gemmatimonadetes
Deinococcus-Thermus
Cyanobacteria
Verrucomicrobia
Chloroflexi
Geothermarchaeota
Fusobacteria
Dictyoglomi
Bacteria
Eukaryota
Archaea

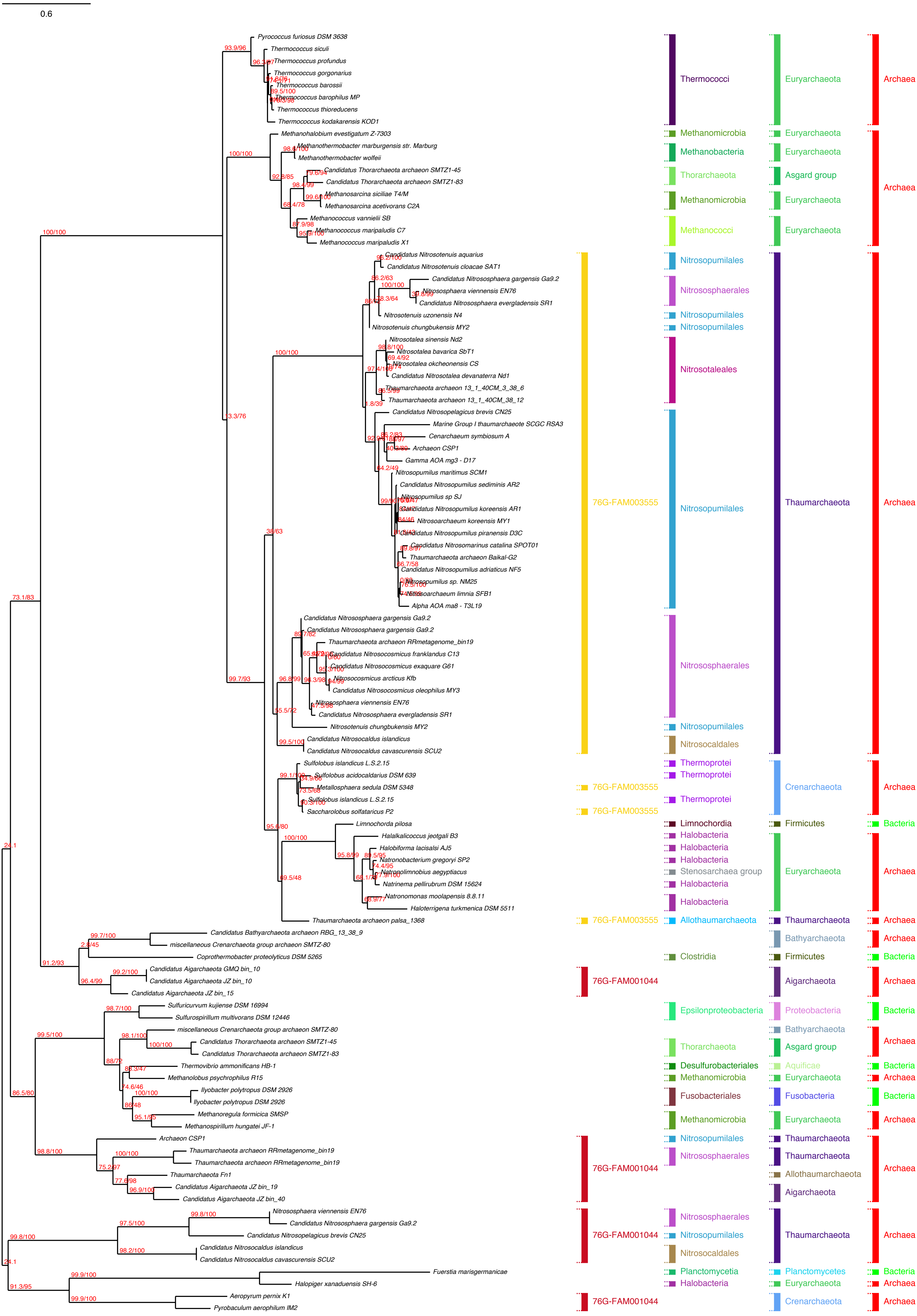

0.6
Pyrococcus furiosus DSM 3638
Thermococcus siculi
Thermococcus profundus
Thermococcus gorgonarius
Thermococcus barossii
Thermococcus barophilus MP
Thermococcus thioreducens
Thermococcus kodakarensis KOD1
Methanohalobium evestigatum Z-7303
Methanothermobacter marburgensis str. Marburg
Methanothermobacter wolfeii
Candidatus Thorarchaeota archaeon SMTZ1-45
Candidatus Thorarchaeota archaeon SMTZ1-83
Methanosarcina siciliae T4/M
Methanosarcina acetivorans C2A
Methanococcus vannielii SB
Methanococcus maripaludis C7
Methanococcus maripaludis X1
Candidatus Nitrosotenuis aquarius
Candidatus Nitrosotenuis cloacae SAT1
Candidatus Nitrososphaera gargensis Ga9.2
Nitrososphaera viennensis EN76
Candidatus Nitrososphaera evergladensis SR1
Nitrosotenuis uzonensis N4
Nitrosotenuis chungbukensis MY2
Nitrosotalea sinensis Nd2
Nitrosotalea bavarica SbT1
Nitrosotalea okcheonensis CS
Candidatus Nitrosotalea devanaterra Nd1
Thaumarchaeota archaeon 13_1_40CM_3_38_6
Thaumarchaeota archaeon 13_1_40CM_38_12
Candidatus Nitrosopelagicus brevis CN25
Marine Group I thaumarchaeote SCGC RSA3
Cenarchaeum symbiosum A
Archaeon CSP1
Gamma AOA mg3 - D17
Nitrosopumilus maritimus SCM1
Candidatus Nitrosopumilus sediminis AR2
Nitrosopumilus sp SJ
Candidatus Nitrosopumilus koreensis AR1
Nitrosoarchaeum koreensis MY1
Candidatus Nitrosopumilus piranensis D3C
Candidatus Nitrosomarinus catalina SPOT01
Thaumarchaeota archaeon Baikal-G2
Candidatus Nitrosopumilus adriaticus NF5
Nitrosopumilus sp. NM25
Nitrosoarchaeum limnia SFB1
Alpha AOA ma8 - T3L19
Candidatus Nitrososphaera gargensis Ga9.2
Candidatus Nitrososphaera gargensis Ga9.2
Thaumarchaeota archaeon RRmetagenome_bin19
Candidatus Nitrosocosmicus franklandus C13
Candidatus Nitrosocosmicus exaquare G61
Nitrosocosmicus arcticus Kfb
Candidatus Nitrosocosmicus oleophilus MY3
Nitrososphaera viennensis EN76
Candidatus Nitrososphaera evergladensis SR1
Nitrosotenuis chungbukensis MY2
Candidatus Nitrosocaldus islandicus
Candidatus Nitrosocaldus cavascurensis SCU2
Sulfolobus islandicus L.S.2.15
Sulfolobus acidocaldarius DSM 639
Metallosphaera sedula DSM 5348
Sulfolobus islandicus L.S.2.15
Saccharolobus solfataricus P2
Limnochorda pilosa
Halalkalicoccus jeotgali B3
Halobiforma lacisalsi AJ5
Natronobacterium gregoryi SP2
Natronolimnobius aegyptiacus
Natrinema pellirubrum DSM 15624
Natronomonas moolapensis 8.8.11
Haloterrigena turkmenica DSM 5511
Thaumarchaeota archaeon palsa_1368
Candidatus Bathyarchaeota archaeon RBG_13_38_9
miscellaneous Crenarchaeota group archaeon SMTZ-80
Coprothermobacter proteolyticus DSM 5265
Candidatus Aigarchaeota GMQ bin_10
Candidatus Aigarchaeota JZ bin_10
Candidatus Aigarchaeota JZ bin_15
Sulfuricurvum kujiense DSM 16994
Sulfurospirillum multivorans DSM 12446
miscellaneous Crenarchaeota group archaeon SMTZ-80
Candidatus Thorarchaeota archaeon SMTZ1-45
Candidatus Thorarchaeota archaeon SMTZ1-83
Thermovibrio ammonificans HB-1
Methanolobus psychrophilus R15
Ilyobacter polytropus DSM 2926
Ilyobacter polytropus DSM 2926
Methanoregula formicica SMSP
Methanospirillum hungatei JF-1
Archaeon CSP1
Thaumarchaeota archaeon RRmetagenome_bin19
Thaumarchaeota archaeon RRmetagenome_bin19
Thaumarchaeota Fn1
Candidatus Aigarchaeota JZ bin_19
Candidatus Aigarchaeota JZ bin_40
Nitrososphaera viennensis EN76
Candidatus Nitrososphaera gargensis Ga9.2
Candidatus Nitrosopelagicus brevis CN25
Candidatus Nitrosocaldus islandicus
Candidatus Nitrosocaldus cavascurensis SCU2
Fuerstia marisgermanicae
Halopiger xanaduensis SH-6
Aeropyrum pernix K1
Pyrobaculum aerophilum IM2
Thermococci
Methanomicrobia
Methanobacteria
Thorarchaeota
Methanomicrobia
Methanococci
Nitrosopumilales
Nitrososphaerales
Nitrosopumilales
Nitrosopumilales
Nitrosotaleales
Nitrosopumilales
Nitrososphaerales
Nitrosopumilales
Nitrosocaldales
Thermoprotei
Thermoprotei
Thermoprotei
Limnochorda
Halobacteria
Halobacteria
Halobacteria
Stenosarchaea group
Halobacteria
Halobacteria
Allothaumarchaeota
Clostridia
Epsilonproteobacteria
Thorarchaeota
Desulfurobacteriales
Methanomicrobia
Fusobacteriales
Methanomicrobia
Nitrosopumilales
Nitrososphaerales
Nitrososphaerales
Nitrosopumilales
Nitrosocaldales
Planctomycetia
Halobacteria
76G-FAM003555
76G-FAM003555
76G-FAM003555
76G-FAM003555
76G-FAM001044
76G-FAM001044
76G-FAM001044
76G-FAM001044
Euryarchaeota
Euryarchaeota
Euryarchaeota
Asgard group
Euryarchaeota
Euryarchaeota
Thaumarchaeota
Crenarchaeota
Firmicutes
Euryarchaeota
Thaumarchaeota
Bathyarchaeota
Firmicutes
Aigarchaeota
Proteobacteria
Bathyarchaeota
Asgard group
Aquificae
Euryarchaeota
Fusobacteria
Euryarchaeota
Thaumarchaeota
Thaumarchaeota
Allothaumarchaeota
Aigarchaeota
Thaumarchaeota
Planctomycetes
Euryarchaeota
Crenarchaeota
Archaea
Archaea
Archaea
Archaea
Bacteria
Archaea
Archaea
Archaea
Bacteria
Archaea
Bacteria
Archaea
Bacteria
Archaea
Bacteria
Archaea
Archaea
Archaea
Bacteria
Archaea
Archaea

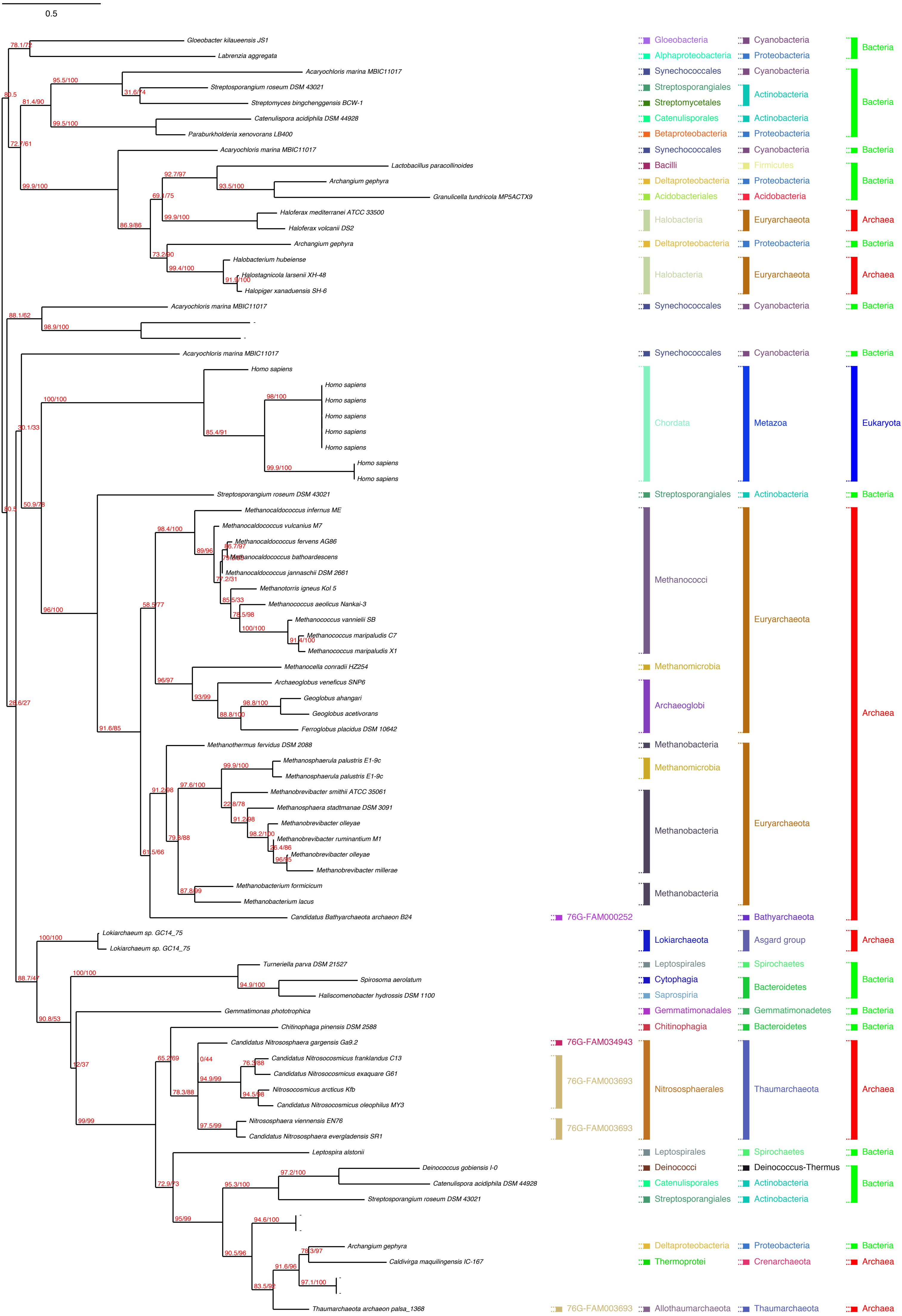

0.5
Gloeobacter kilaueensis JS1
Labrenzia aggregata
Acaryochloris marina MBIC11017
Streptosporangium roseum DSM 43021
Streptomyces bingchenggensis BCW-1
Catenulispora acidiphila DSM 44928
Paraburkholderia xenovorans LB400
Lactobacillus paracollinoides
Archangium gephyra
Granulicella tundricola MP5ACTX9
Haloferax mediterranei ATCC 33500
Haloferax volcanii DS2
Halobacterium hubeiense
Halostagnicola larsenii XH-48
Halopiger xanaduensis SH-6
Homo sapiens
Methanocaldococcus infernus ME
Methanocaldococcus vulcanius M7
Methanocaldococcus fervens AG86
Methanocaldococcus bathoardescens
Methanocaldococcus jannaschii DSM 2661
Methanotorris igneus Kol 5
Methanococcus aeolicus Nankai-3
Methanococcus vannielii SB
Methanococcus maripaludis C7
Methanococcus maripaludis X1
Methanocella conradii HZ254
Archaeoglobus veneficus SNP6
Geoglobus ahangari
Geoglobus acetivorans
Ferroglobus placidus DSM 10642
Methanothermus fervidus DSM 2088
Methanosphaerula palustris E1-9c
Methanobrevibacter smithii ATCC 35061
Methanosphaera stadtmanae DSM 3091
Methanobrevibacter olleyae
Methanobrevibacter ruminantium M1
Methanobrevibacter millerae
Methanobacterium formicicum
Methanobacterium lacus
Candidatus Bathyarchaeota archaeon B24
Lokiarchaeum sp. GC14_75
Turneriella parva DSM 21527
Spirosoma aerolatum
Haliscomenobacter hydrossis DSM 1100
Gemmatimonas phototrophica
Chitinophaga pinensis DSM 2588
Candidatus Nitrososphaera gargensis Ga9.2
Candidatus Nitrosocosmicus franklandus C13
Candidatus Nitrosocosmicus exaquare G61
Nitrosocosmicus arcticus Kfb
Candidatus Nitrosocosmicus oleophilus MY3
Nitrososphaera viennensis EN76
Candidatus Nitrososphaera evergladensis SR1
Leptospira alstonii
Deinococcus gobiensis I-0
Caldivirga maquilingensis IC-167
Thaumarchaeota archaeon palsa_1368
Gloeobacteria
Alphaproteobacteria
Synechococcales
Streptosporangiales
Streptomycetales
Catenulisporales
Betaproteobacteria
Bacilli
Deltaproteobacteria
Acidobacteriales
Halobacteria
Chordata
Methanococci
Methanomicrobia
Archaeoglobi
Methanobacteria
76G-FAM000252
Lokiarchaeota
Leptospirales
Cytophagia
Saprospiria
Gemmatimonadales
Chitinophagia
76G-FAM034943
76G-FAM003693
Nitrososphaerales
Deinococci
Thermoprotei
Allothaumarchaeota
Cyanobacteria
Proteobacteria
Actinobacteria
Firmicutes
Acidobacteria
Euryarchaeota
Metazoa
Bathyarchaeota
Asgard group
Spirochaetes
Bacteroidetes
Gemmatimonadetes
Thaumarchaeota
Deinococcus-Thermus
Crenarchaeota
Bacteria
Archaea
Eukaryota

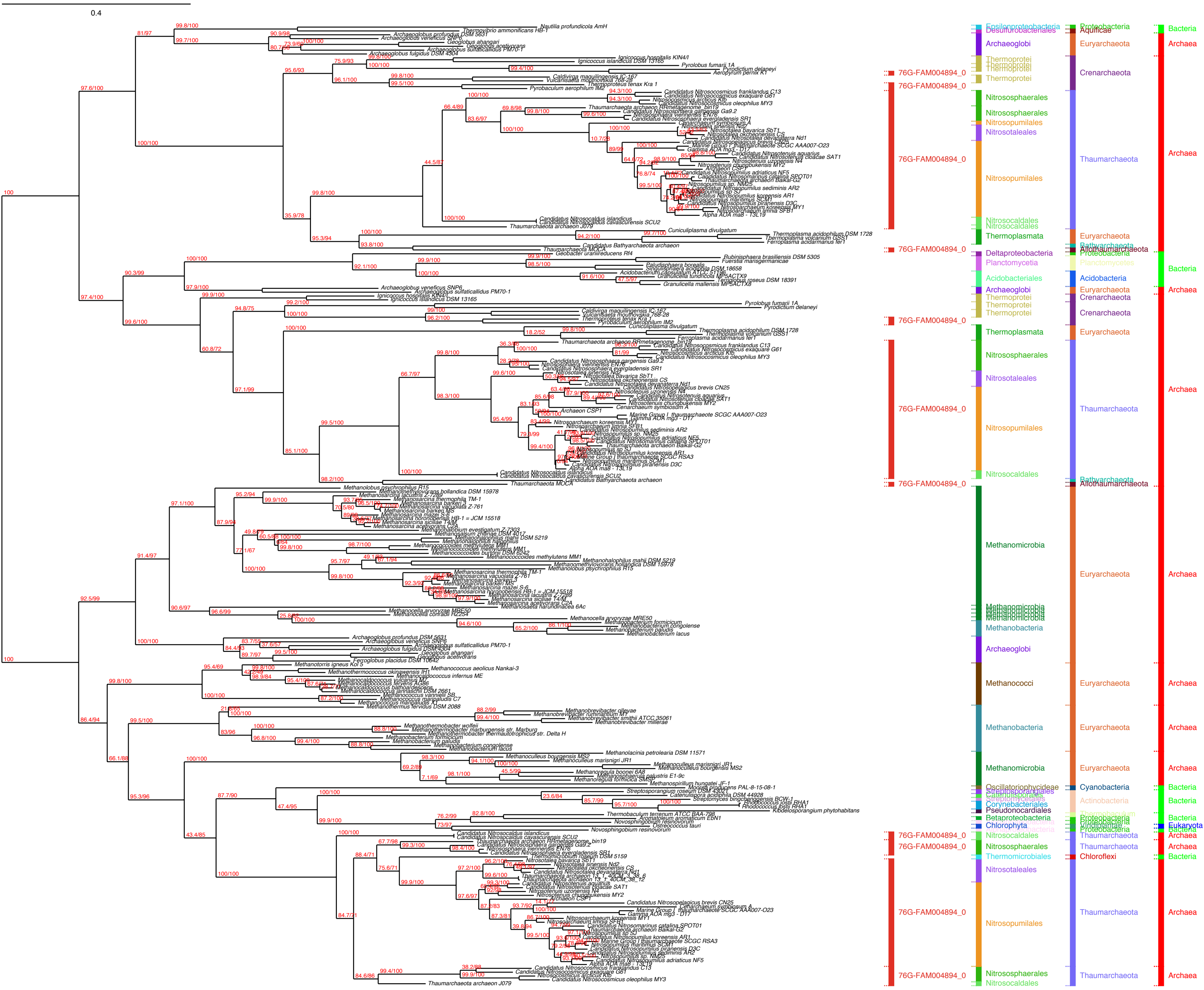
0.4
76G-FAM004894_0
Epsilonproteobacteria
Desulfurobacteriales
Archaeoglobi
Thermoprotei
Nitrososphaerales
Nitrosopumilales
Nitrosotaleales
Nitrosocaldales
Thermoplasmata
Deltaproteobacteria
Planctomycetia
Acidobacteriales
Methanomicrobia
Methanobacteria
Methanococci
Oscillatoriophycideae
Streptosporangiales
Corynebacteriales
Pseudonocardiales
Betaproteobacteria
Chlorophyta
Thermomicrobiales
Proteobacteria
Aquificae
Euryarchaeota
Crenarchaeota
Thaumarchaeota
Bathyarchaeota
Allothaumarchaeota
Planctomycetes
Acidobacteria
Cyanobacteria
Actinobacteria
Viridiplantae
Chloroflexi
Bacteria
Archaea
Eukaryota

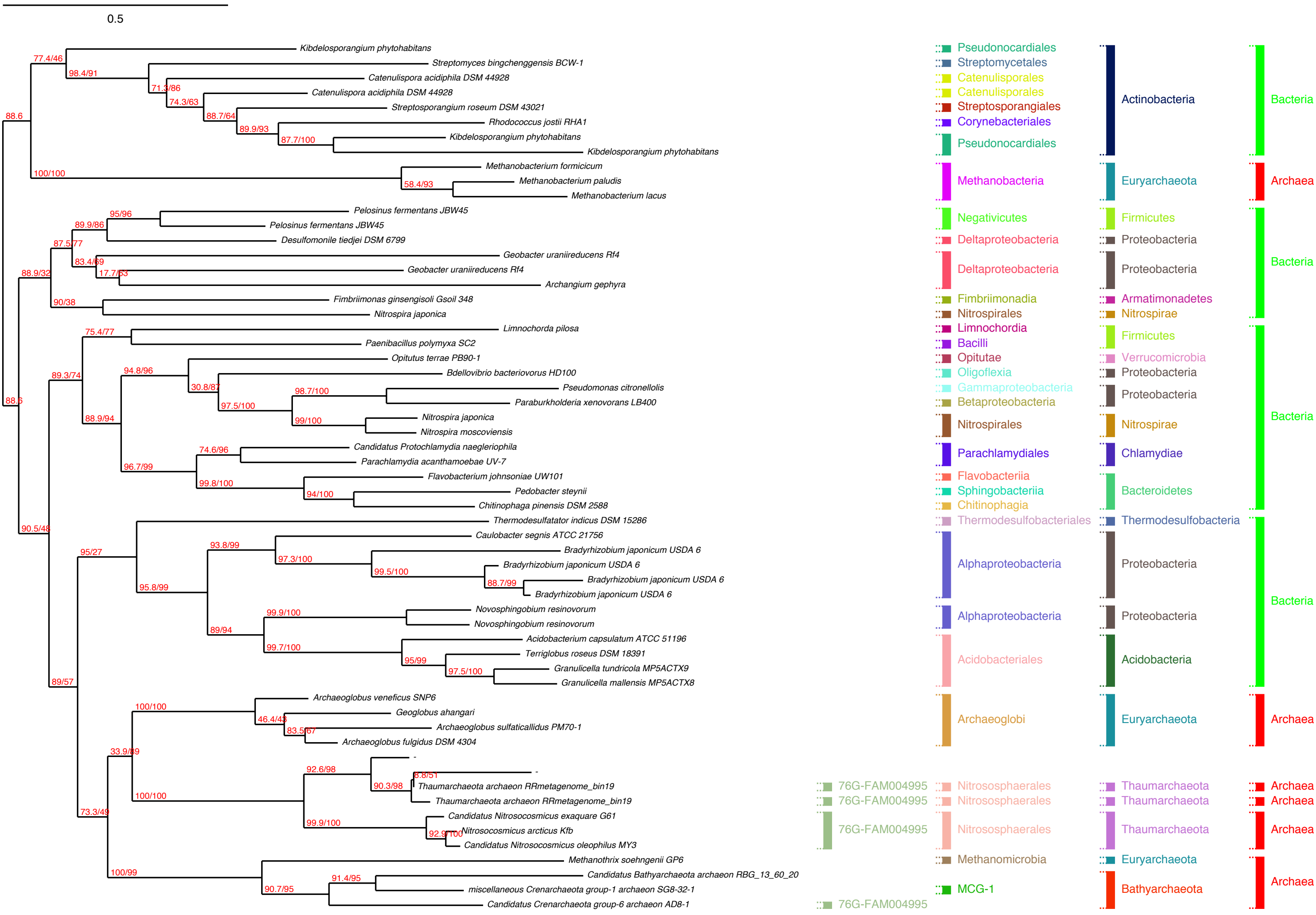

0.5
Kibdelosporangium phytohabitans
Streptomyces bingchenggensis BCW-1
Catenulispora acidiphila DSM 44928
Catenulispora acidiphila DSM 44928
Streptosporangium roseum DSM 43021
Rhodococcus jostii RHA1
Kibdelosporangium phytohabitans
Kibdelosporangium phytohabitans
Methanobacterium formicicum
Methanobacterium paludis
Methanobacterium lacus
Pelosinus fermentans JBW45
Pelosinus fermentans JBW45
Desulfomonile tiedjei DSM 6799
Geobacter uraniireducens Rf4
Geobacter uraniireducens Rf4
Archangium gephyra
Fimbriimonas ginsengisoli Gsoil 348
Nitrospira japonica
Limnochorda pilosa
Paenibacillus polymyxa SC2
Opitutus terrae PB90-1
Bdellovibrio bacteriovorus HD100
Pseudomonas citronellolis
Paraburkholderia xenovorans LB400
Nitrospira japonica
Nitrospira moscoviensis
Candidatus Protochlamydia naegleriophila
Parachlamydia acanthamoebae UV-7
Flavobacterium johnsoniae UW101
Pedobacter steynii
Chitinophaga pinensis DSM 2588
Thermodesulfatator indicus DSM 15286
Caulobacter segnis ATCC 21756
Bradyrhizobium japonicum USDA 6
Bradyrhizobium japonicum USDA 6
Bradyrhizobium japonicum USDA 6
Bradyrhizobium japonicum USDA 6
Novosphingobium resinovorum
Novosphingobium resinovorum
Acidobacterium capsulatum ATCC 51196
Terriglobus roseus DSM 18391
Granulicella tundricola MP5ACTX9
Granulicella mallensis MP5ACTX8
Archaeoglobus veneficus SNP6
Geoglobus ahangari
Archaeoglobus sulfaticallidus PM70-1
Archaeoglobus fulgidus DSM 4304
Thaumarchaeota archaeon RRmetagenome_bin19
Thaumarchaeota archaeon RRmetagenome_bin19
Candidatus Nitrosocosmicus exaquare G61
Nitrosocosmicus arcticus Kfb
Candidatus Nitrosocosmicus oleophilus MY3
Methanothrix soehngenii GP6
Candidatus Bathyarchaeota archaeon RBG_13_60_20
miscellaneous Crenarchaeota group-1 archaeon SG8-32-1
Candidatus Crenarchaeota group-6 archaeon AD8-1
77.4/46
98.4/91
88.6
74.3/63
88.7/64
89.9/93
87.7/100
100/100
58.4/93
95/96
89.9/86
88.9/32
90/38
75.4/77
89.3/74
94.8/96
98.7/100
97.5/100
99/100
88.9/94
74.6/96
96.7/99
99.8/100
94/100
90.5/48
95/27
93.8/99
97.3/100
99.5/100
88.7/99
95.8/99
99.9/100
89/94
99.7/100
95/99
97.5/100
89/57
100/100
46.4/43
83.5/67
92.6/98
90.3/98
100/100
99.9/100
73.3/49
100/99
91.4/95
90.7/95
76G-FAM004995
76G-FAM004995
76G-FAM004995
76G-FAM004995
Pseudonocardiales
Streptomycetales
Catenulisporales
Catenulisporales
Streptosporangiales
Corynebacteriales
Pseudonocardiales
Methanobacteria
Negativicutes
Deltaproteobacteria
Deltaproteobacteria
Fimbriimonadia
Nitrospirales
Limnochordia
Bacilli
Opitutae
Oligoflexia
Gammaproteobacteria
Betaproteobacteria
Nitrospirales
Parachlamydiales
Flavobacteriia
Sphingobacteriia
Chitinophagia
Thermodesulfobacteriales
Alphaproteobacteria
Alphaproteobacteria
Acidobacteriales
Archaeoglobi
Nitrososphaerales
Nitrososphaerales
Nitrososphaerales
Methanomicrobia
MCG-1
Actinobacteria
Euryarchaeota
Firmicutes
Proteobacteria
Proteobacteria
Armatimonadetes
Nitrospirae
Firmicutes
Verrucomicrobia
Proteobacteria
Proteobacteria
Nitrospirae
Chlamydiae
Bacteroidetes
Thermodesulfobacteria
Proteobacteria
Proteobacteria
Acidobacteria
Euryarchaeota
Thaumarchaeota
Thaumarchaeota
Thaumarchaeota
Euryarchaeota
Bathyarchaeota
Bacteria
Archaea
Bacteria
Bacteria
Bacteria
Archaea
Archaea
Archaea
Archaea
Archaea

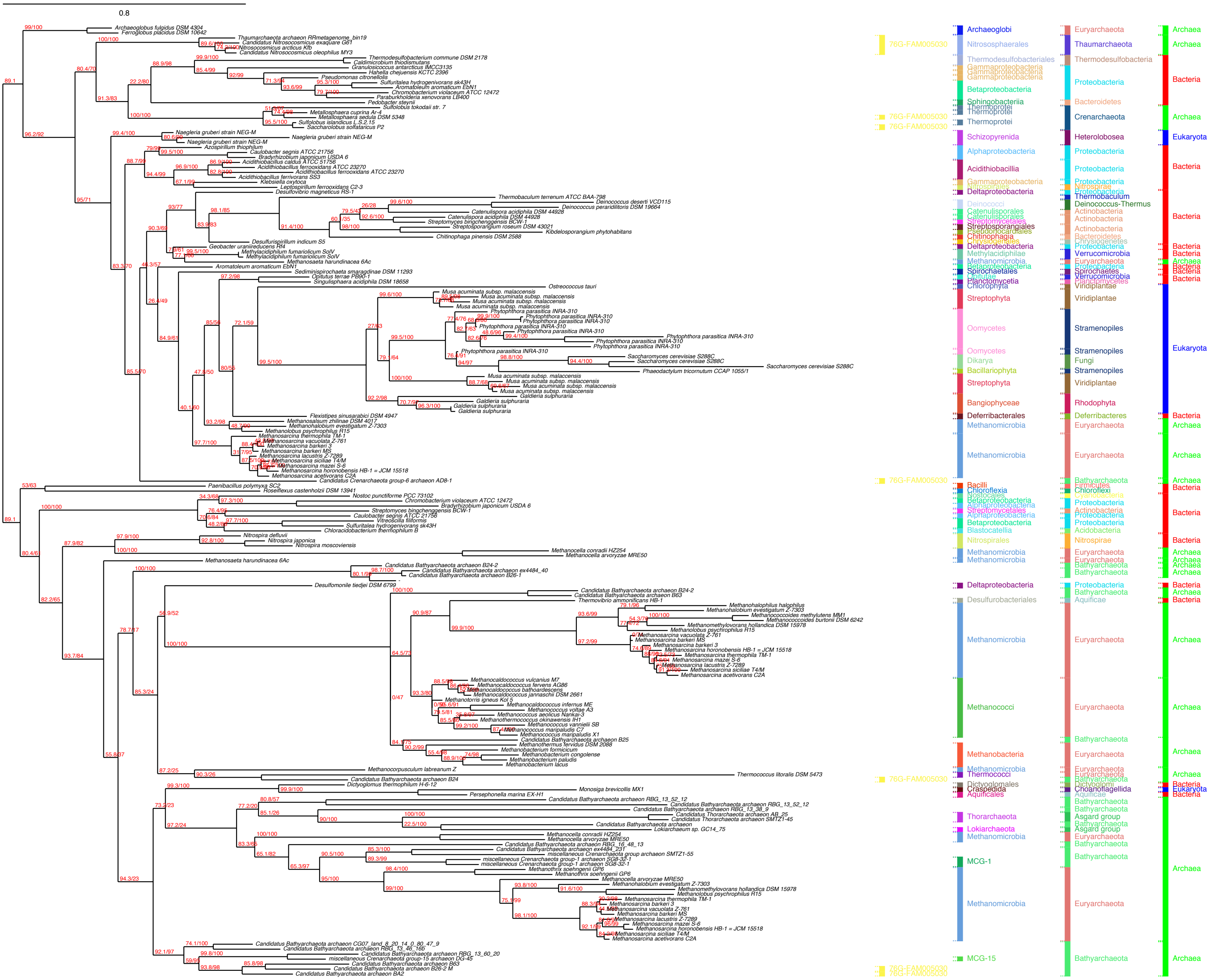

0.8
Archaeoglobus fulgidus DSM 4304
Ferroglobus placidus DSM 10642
Thaumarchaeota archaeon RRmetagenome_bin19
Candidatus Nitrosocosmicus exaquare G61
Nitrosocosmicus arcticus Kfb
Candidatus Nitrosocosmicus oleophilus MY3
Thermodesulfobacterium commune DSM 2178
Caldimicrobium thiodismutans
Granulosicoccus antarcticus IMCC3135
Hahella chejuensis KCTC 2396
Pseudomonas citronellolis
Sulfuritalea hydrogenivorans sk43H
Aromatoleum aromaticum EbN1
Chromobacterium violaceum ATCC 12472
Paraburkholderia xenovorans LB400
Pedobacter steynii
Sulfolobus tokodaii str. 7
Metallosphaera cuprina Ar-4
Metallosphaera sedula DSM 5348
Sulfolobus islandicus L.S.2.15
Saccharolobus solfataricus P2
Naegleria gruberi strain NEG-M
Azospirillum thiophilum
Caulobacter segnis ATCC 21756
Bradyrhizobium japonicum USDA 6
Acidithiobacillus caldus ATCC 51756
Acidithiobacillus ferrooxidans ATCC 23270
Acidithiobacillus ferrivorans SS3
Klebsiella oxytoca
Leptospirillum ferrooxidans C2-3
Desulfovibrio magneticus RS-1
Thermobaculum terrenum ATCC BAA-798
Deinococcus deserti VCD115
Deinococcus peraridilitoris DSM 19664
Catenulispora acidiphila DSM 44928
Streptomyces bingchenggensis BCW-1
Streptosporangium roseum DSM 43021
Kibdelosporangium phytohabitans
Chitinophaga pinensis DSM 2588
Desulfurispirillum indicum S5
Geobacter uraniireducens Rf4
Methylacidiphilum fumariolicum SolV
Methanosaeta harundinacea 6Ac
Aromatoleum aromaticum EbN1
Sediminispirochaeta smaragdinae DSM 11293
Opitutus terrae PB90-1
Singulisphaera acidiphila DSM 18658
Ostreococcus tauri
Musa acuminata subsp. malaccensis
Phytophthora parasitica INRA-310
Saccharomyces cerevisiae S288C
Phaeodactylum tricornutum CCAP 1055/1
Galdieria sulphuraria
Flexistipes sinusarabici DSM 4947
Methanosalsum zhilinae DSM 4017
Methanohalobium evestigatum Z-7303
Methanolobus psychrophilus R15
Methanosarcina thermophila TM-1
Methanosarcina vacuolata Z-761
Methanosarcina barkeri 3
Methanosarcina barkeri MS
Methanosarcina lacustris Z-7289
Methanosarcina siciliae T4/M
Methanosarcina mazei S-6
Methanosarcina horonobensis HB-1 = JCM 15518
Methanosarcina acetivorans C2A
Candidatus Crenarchaeota group-6 archaeon AD8-1
Paenibacillus polymyxa SC2
Roseiflexus castenholzii DSM 13941
Nostoc punctiforme PCC 73102
Chromobacterium violaceum ATCC 12472
Bradyrhizobium japonicum USDA 6
Streptomyces bingchenggensis BCW-1
Caulobacter segnis ATCC 21756
Vitreoscilla filiformis
Sulfuritalea hydrogenivorans sk43H
Chloracidobacterium thermophilum B
Nitrospira defluvii
Nitrospira japonica
Nitrospira moscoviensis
Methanocella conradii HZ254
Methanocella arvoryzae MRE50
Methanosaeta harundinacea 6Ac
Candidatus Bathyarchaeota archaeon B24-2
Candidatus Bathyarchaeota archaeon ex4484_40
Candidatus Bathyarchaeota archaeon B26-1
Desulfomonile tiedjei DSM 6799
Candidatus Bathyarchaeota archaeon B63
Thermovibrio ammonificans HB-1
Methanohalophilus halophilus
Methanohalobium evestigatum Z-7303
Methanococcoides methylutens MM1
Methanococcoides burtonii DSM 6242
Methanomethylovorans hollandica DSM 15978
Methanolobus psychrophilus R15
Methanocaldococcus vulcanius M7
Methanocaldococcus fervens AG86
Methanocaldococcus bathoardescens
Methanocaldococcus jannaschii DSM 2661
Methanotorris igneus Kol 5
Methanocaldococcus infernus ME
Methanococcus voltae A3
Methanococcus aeolicus Nankai-3
Methanothermococcus okinawensis IH1
Methanococcus vannielii SB
Methanococcus maripaludis C7
Methanococcus maripaludis X1
Candidatus Bathyarchaeota archaeon B25
Methanothermus fervidus DSM 2088
Methanobacterium formicicum
Methanobacterium congolense
Methanobacterium paludis
Methanobacterium lacus
Methanocorpusculum labreanum Z
Thermococcus litoralis DSM 5473
Candidatus Bathyarchaeota archaeon B24
Dictyoglomus thermophilum H-6-12
Monosiga brevicollis MX1
Persephonella marina EX-H1
Candidatus Bathyarchaeota archaeon RBG_13_52_12
Candidatus Bathyarchaeota archaeon RBG_13_38_9
Candidatus Thorarchaeota archaeon AB_25
Candidatus Thorarchaeota archaeon SMTZ1-45
Candidatus Bathyarchaeota archaeon
Lokiarchaeum sp. GC14_75
Candidatus Bathyarchaeota archaeon RBG_16_48_13
Candidatus Bathyarchaeota archaeon ex4484_231
miscellaneous Crenarchaeota group archaeon SMTZ1-55
miscellaneous Crenarchaeota group-1 archaeon SG8-32-1
Methanothrix soehngenii GP6
Candidatus Bathyarchaeota archaeon CG07_land_8_20_14_0_80_47_9
Candidatus Bathyarchaeota archaeon RBG_13_46_16b
Candidatus Bathyarchaeota archaeon RBG_13_60_20
miscellaneous Crenarchaeota group-15 archaeon DG-45
Candidatus Bathyarchaeota archaeon B26-2 M
Candidatus Bathyarchaeota archaeon BA2
76G-FAM005030
Archaeoglobi
Nitrososphaerales
Thermodesulfobacteriales
Gammaproteobacteria
Betaproteobacteria
Sphingobacteriia
Thermoprotei
Schizopyrenida
Alphaproteobacteria
Acidithiobacillia
Nitrospirales
Deltaproteobacteria
Deinococci
Catenulisporales
Streptomycetales
Streptosporangiales
Pseudonocardiales
Chitinophagia
Chrysiogenetes
Methylacidiphilae
Methanomicrobia
Spirochaetales
Opitutae
Planctomycetia
Chlorophyta
Streptophyta
Oomycetes
Dikarya
Bacillariophyta
Bangiophyceae
Deferribacterales
Bacilli
Chloroflexia
Nostocales
Blastocatellia
Desulfurobacteriales
Methanococci
Methanobacteria
Thermococci
Dictyoglomales
Craspedida
Aquificales
Thorarchaeota
Lokiarchaeota
MCG-1
MCG-15
Euryarchaeota
Thaumarchaeota
Thermodesulfobacteria
Proteobacteria
Bacteroidetes
Crenarchaeota
Heterolobosea
Thermobaculum
Deinococcus-Thermus
Actinobacteria
Verrucomicrobia
Spirochaetes
Planctomycetes
Viridiplantae
Stramenopiles
Fungi
Rhodophyta
Deferribacteres
Bathyarchaeota
Firmicutes
Chloroflexi
Cyanobacteria
Acidobacteria
Nitrospirae
Aquificae
Dictyoglomi
Choanoflagellida
Asgard group
Archaea
Bacteria
Eukaryota

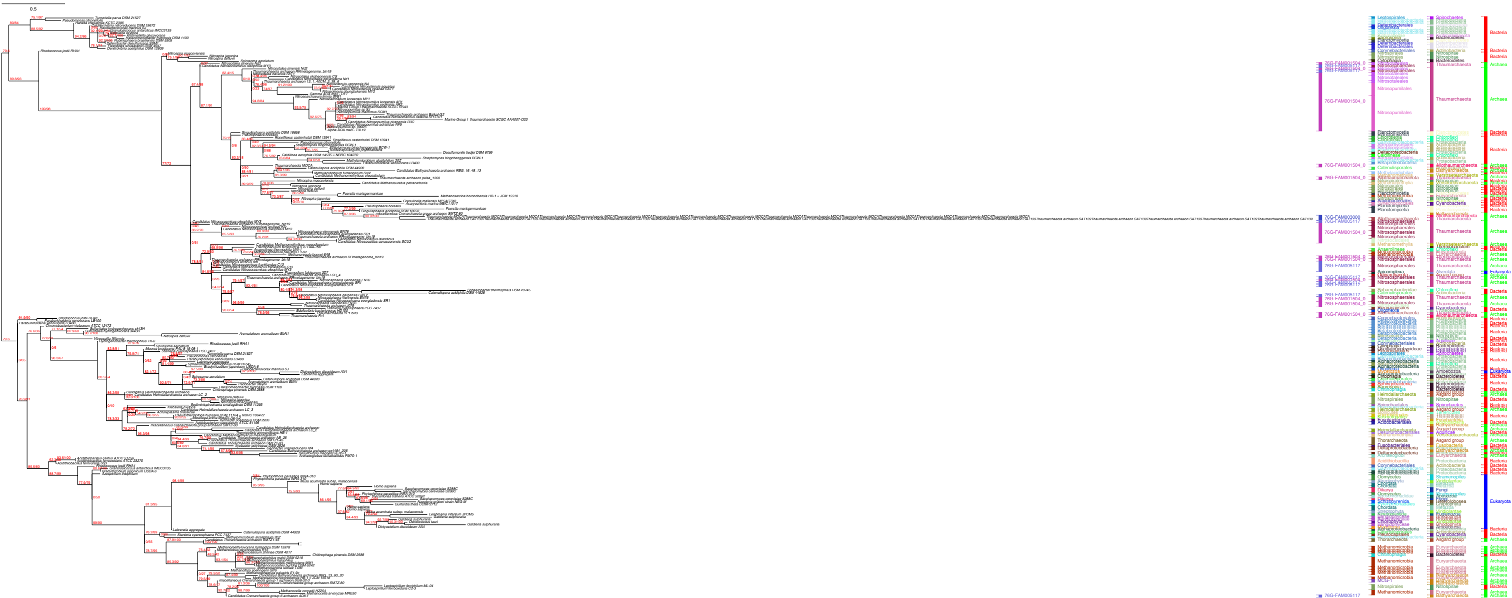

0.5
Bacteria
Archaea
Eukaryota
Thaumarchaeota
Nitrosopumilales
Nitrososphaerales

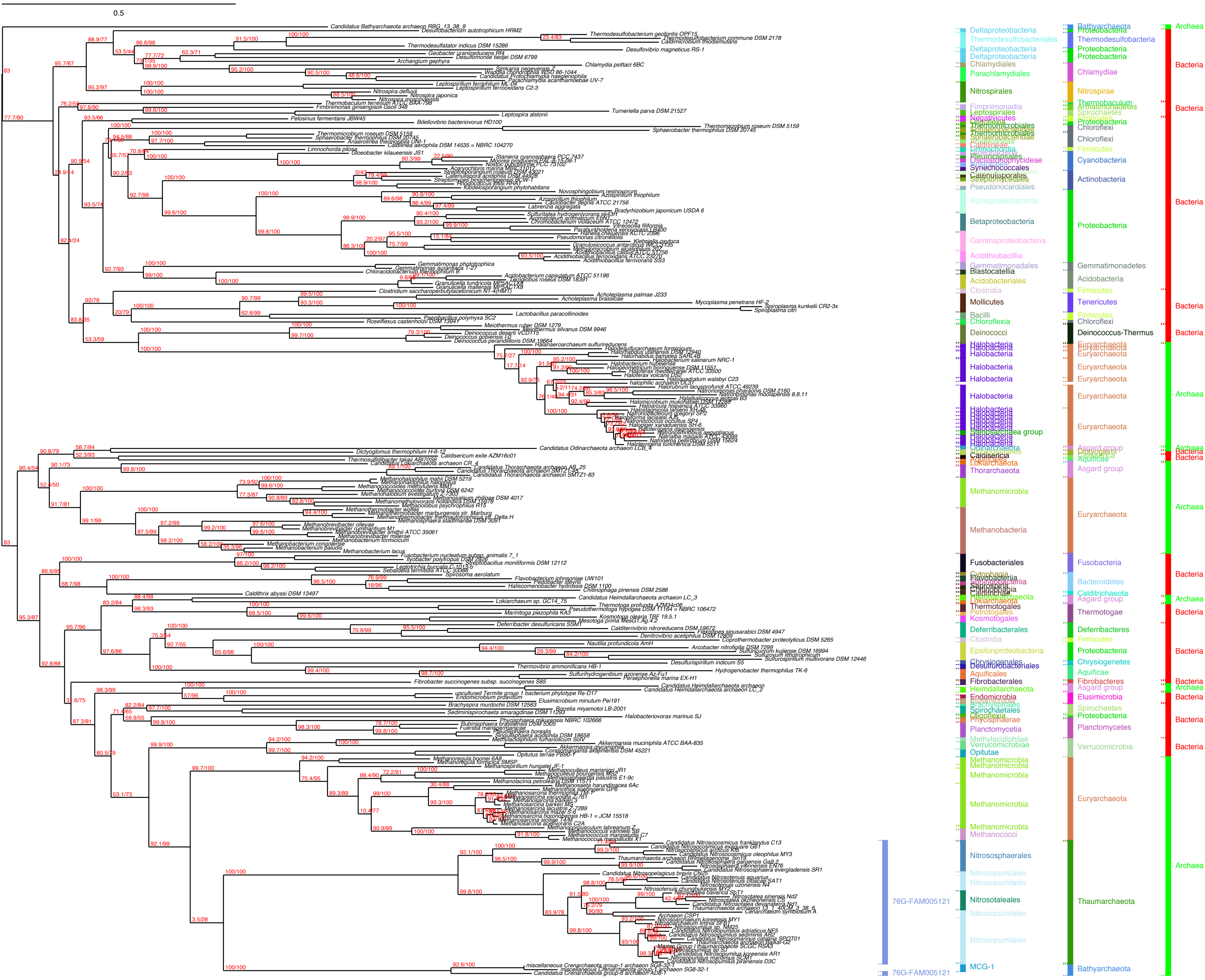
0.5
76G-FAM005121
76G-FAM005121
Deltaproteobacteria
Thermodesulfobacteriales
Chlamydiales
Parachlamydiales
Nitrospirales
Alphaproteobacteria
Betaproteobacteria
Gammaproteobacteria
Acidithiobacillia
Gemmatimonadales
Blastocatellia
Acidobacteriales
Clostridia
Mollicutes
Bacilli
Chloroflexia
Deinococci
Halobacteria
Methanomicrobia
Methanobacteria
Fusobacteriales
Deferribacterales
Epsilonproteobacteria
Fibrobacterales
Heimdallarchaeota
Planctomycetia
Opitutae
Methanococci
Nitrososphaerales
Nitrosopumilales
Nitrosotaleales
MCG-1
Bathyarchaeota
Proteobacteria
Thermodesulfobacteria
Chlamydiae
Nitrospirae
Chloroflexi
Firmicutes
Cyanobacteria
Actinobacteria
Gemmatimonadetes
Acidobacteria
Tenericutes
Deinococcus-Thermus
Euryarchaeota
Asgard group
Fusobacteria
Bacteroidetes
Thermotogae
Deferribacteres
Chrysiogenetes
Aquificae
Fibrobacteres
Elusimicrobia
Spirochaetes
Planctomycetes
Verrucomicrobia
Thaumarchaeota
Archaea
Bacteria

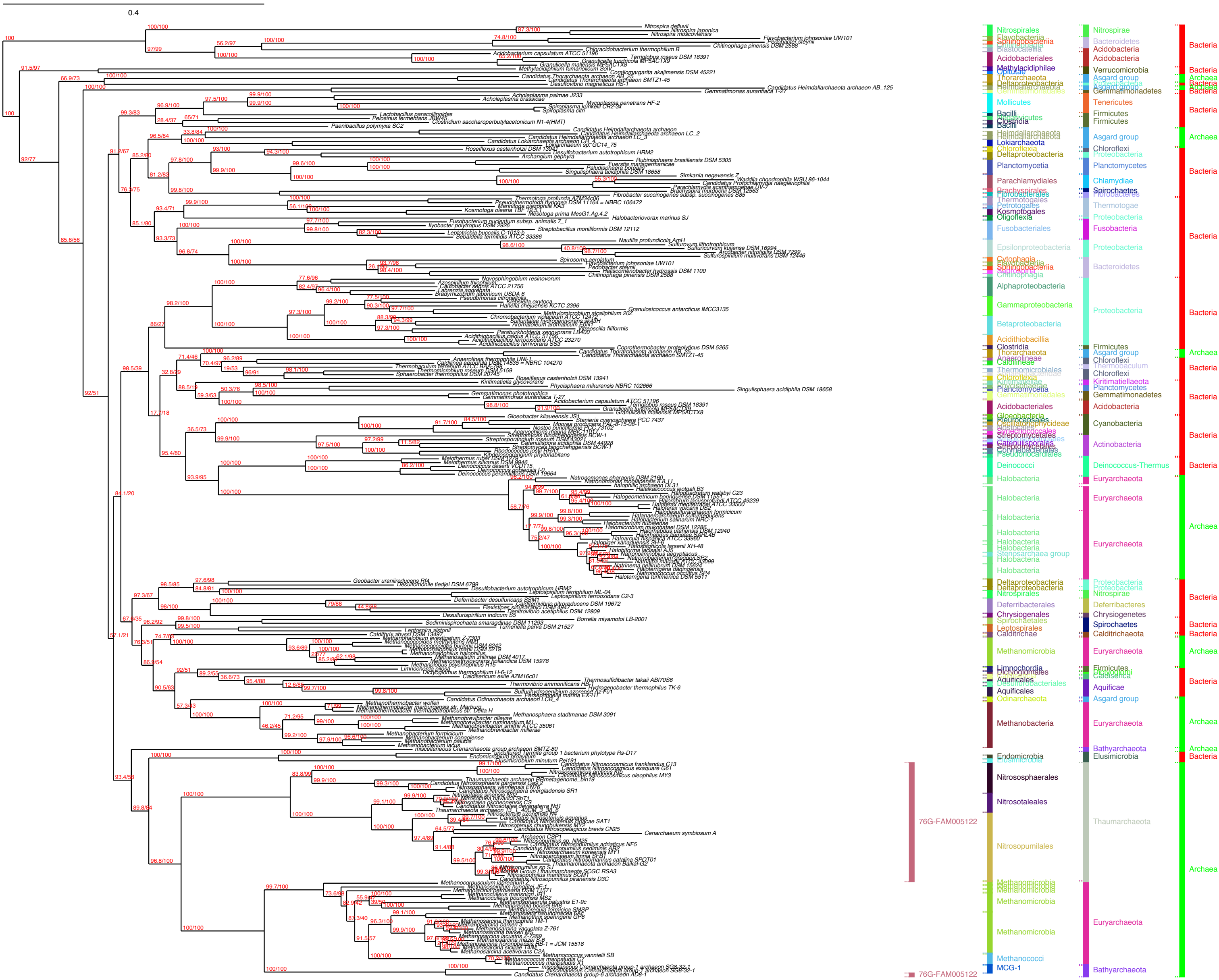

0.4
76G-FAM005122
Nitrospirales
Acidobacteriales
Mollicutes
Planctomycetia
Parachlamydiales
Fusobacteriales
Epsilonproteobacteria
Alphaproteobacteria
Gammaproteobacteria
Betaproteobacteria
Acidithiobacillia
Acidobacteriales
Deinococci
Halobacteria
Methanomicrobia
Methanobacteria
Endomicrobia
Nitrososphaerales
Nitrosotaleales
Nitrosopumilales
Methanococci
MCG-1
Nitrospirae
Bacteroidetes
Acidobacteria
Verrucomicrobia
Asgard group
Tenericutes
Firmicutes
Chloroflexi
Proteobacteria
Planctomycetes
Chlamydiae
Spirochaetes
Thermotogae
Fusobacteria
Gemmatimonadetes
Cyanobacteria
Actinobacteria
Deinococcus-Thermus
Euryarchaeota
Deferribacteres
Chrysiogenetes
Caldithrichaeota
Aquificae
Bathyarchaeota
Elusimicrobia
Thaumarchaeota
Bacteria
Archaea

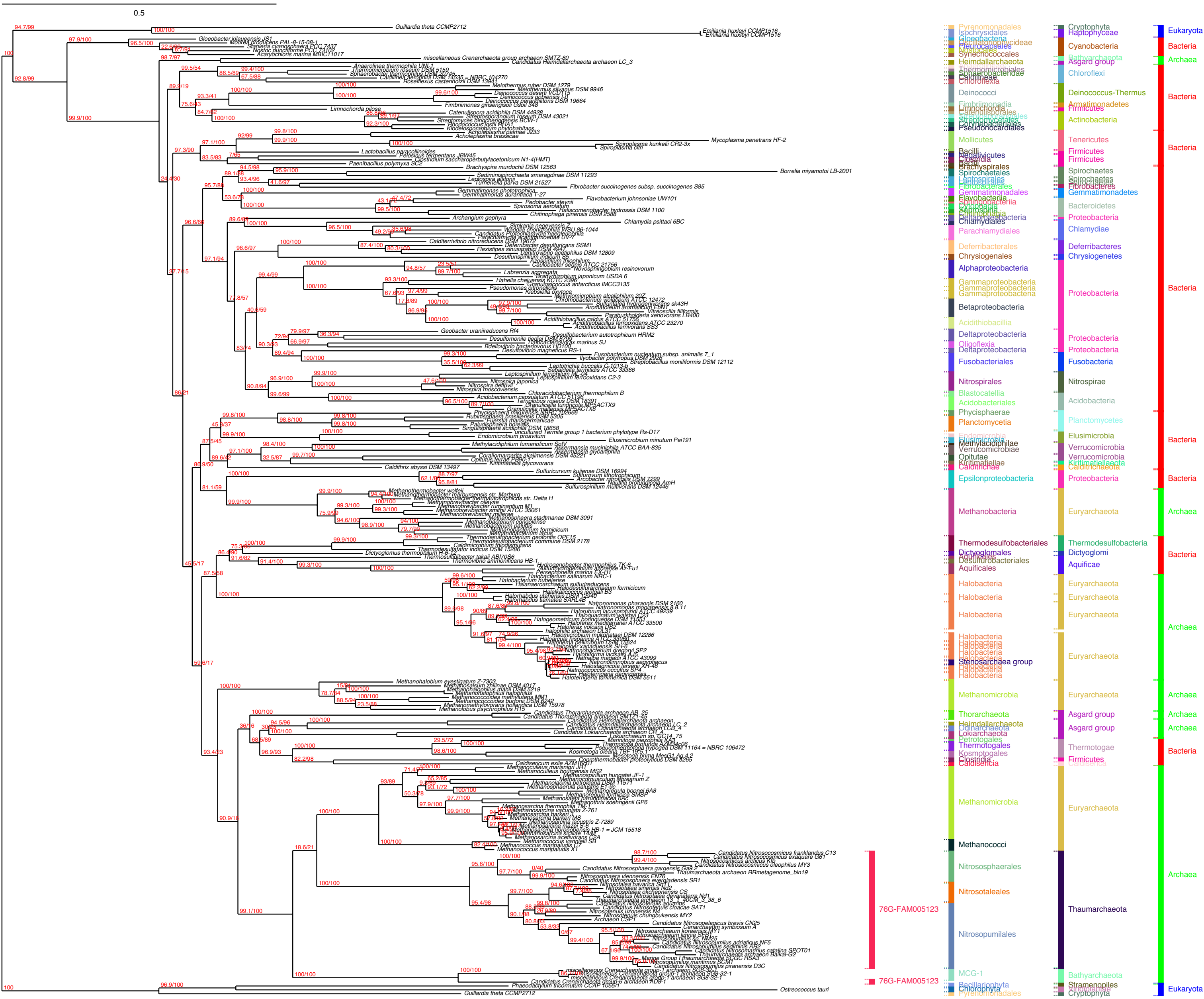

0.5
Pyrenomonadales
Isochrysidales
Gloeobacteria
Oscillatoriophycideae
Pleurocapsales
Nostocales
Synechococcales
Heimdallarchaeota
Thermomicrobiales
Sphaerobacteridae
Caldilineae
Chloroflexia
Deinococci
Fimbriimonadia
Limnochordia
Catenulisporales
Streptosporangiales
Streptomycetales
Corynebacteriales
Pseudonocardiales
Mollicutes
Bacilli
Negativicutes
Clostridia
Brachyspirales
Spirochaetales
Leptospirales
Fibrobacterales
Gemmatimonadetes
Flavobacteriia
Sphingobacteriia
Cytophagia
Saprospiria
Chitinophagia
Deltaproteobacteria
Chlamydiales
Parachlamydiales
Deferribacterales
Chrysiogenales
Alphaproteobacteria
Gammaproteobacteria
Betaproteobacteria
Acidithiobacillia
Deltaproteobacteria
Oligoflexia
Deltaproteobacteria
Fusobacteriales
Nitrospirales
Blastocatellia
Acidobacteriales
Phycisphaerae
Planctomycetia
Endomicrobia
Elusimicrobia
Methylacidiphilae
Verrucomicrobiae
Opitutae
Kiritimatiellae
Caldithrichae
Epsilonproteobacteria
Methanobacteria
Thermodesulfobacteriales
Dictyoglomales
Aquificales
Desulfurobacteriales
Aquificales
Halobacteria
Stenosarchaea group
Methanomicrobia
Thorarchaeota
Heimdallarchaeota
Odinarchaeota
Lokiarchaeota
Petrotogales
Thermotogales
Kosmotogales
Clostridia
Caldisericia
Methanomicrobia
Methanococci
Nitrososphaerales
Nitrosotaleales
Nitrosopumilales
MCG-1
Bacillariophyta
Chlorophyta
Pyrenomonadales
Cryptophyta
Haptophyceae
Cyanobacteria
Bathyarchaeota
Asgard group
Chloroflexi
Deinococcus-Thermus
Armatimonadetes
Firmicutes
Actinobacteria
Tenericutes
Firmicutes
Firmicutes
Spirochaetes
Spirochaetes
Spirochaetes
Fibrobacteres
Gemmatimonadetes
Bacteroidetes
Proteobacteria
Chlamydiae
Deferribacteres
Chrysiogenetes
Proteobacteria
Proteobacteria
Proteobacteria
Fusobacteria
Nitrospirae
Acidobacteria
Planctomycetes
Elusimicrobia
Verrucomicrobia
Verrucomicrobia
Kiritimatiellaeota
Calditrichaeota
Proteobacteria
Euryarchaeota
Thermodesulfobacteria
Dictyoglomi
Aquificae
Euryarchaeota
Euryarchaeota
Euryarchaeota
Euryarchaeota
Euryarchaeota
Asgard group
Asgard group
Thermotogae
Firmicutes
Caldiserica
Euryarchaeota
Thaumarchaeota
Bathyarchaeota
Stramenopiles
Chlorophyta
Cryptophyta
Eukaryota
Bacteria
Archaea
Bacteria
Bacteria
Bacteria
Bacteria
Bacteria
Archaea
Bacteria
Archaea
Archaea
Archaea
Bacteria
Archaea
Eukaryota
76G-FAM005123
76G-FAM005123

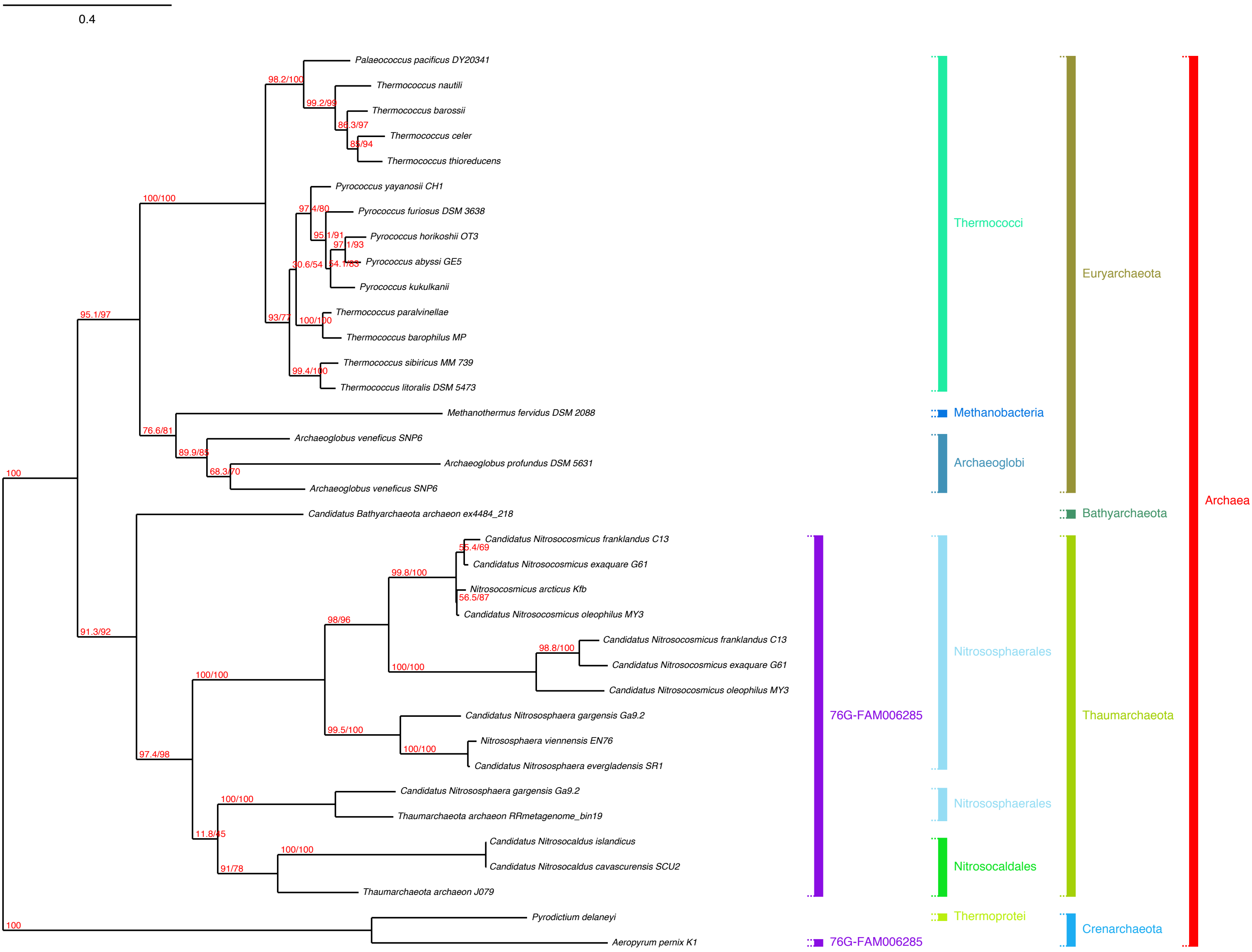

0.4
Palaeococcus pacificus DY20341
Thermococcus nautili
Thermococcus barossii
Thermococcus celer
Thermococcus thioreducens
Pyrococcus yayanosii CH1
Pyrococcus furiosus DSM 3638
Pyrococcus horikoshii OT3
Pyrococcus abyssi GE5
Pyrococcus kukulkanii
Thermococcus paralvinellae
Thermococcus barophilus MP
Thermococcus sibiricus MM 739
Thermococcus litoralis DSM 5473
Methanothermus fervidus DSM 2088
Archaeoglobus veneficus SNP6
Archaeoglobus profundus DSM 5631
Archaeoglobus veneficus SNP6
Candidatus Bathyarchaeota archaeon ex4484_218
Candidatus Nitrosocosmicus franklandus C13
Candidatus Nitrosocosmicus exaquare G61
Nitrosocosmicus arcticus Kfb
Candidatus Nitrosocosmicus oleophilus MY3
Candidatus Nitrosocosmicus franklandus C13
Candidatus Nitrosocosmicus exaquare G61
Candidatus Nitrosocosmicus oleophilus MY3
Candidatus Nitrososphaera gargensis Ga9.2
Nitrososphaera viennensis EN76
Candidatus Nitrososphaera evergladensis SR1
Candidatus Nitrososphaera gargensis Ga9.2
Thaumarchaeota archaeon RRmetagenome_bin19
Candidatus Nitrosocaldus islandicus
Candidatus Nitrosocaldus cavascurensis SCU2
Thaumarchaeota archaeon J079
Pyrodictium delaneyi
Aeropyrum pernix K1
98.2/100
99.2/99
86.3/97
85/94
100/100
97.4/80
95.1/91
97.1/93
30.6/54
54.1/83
93/77
100/100
99.4/100
95.1/97
76.6/81
89.9/85
68.3/70
100
55.4/69
99.8/100
56.5/87
98/96
98.8/100
100/100
91.3/92
100/100
99.5/100
100/100
97.4/98
100/100
11.8/45
100/100
91/78
100
Thermococci
Methanobacteria
Archaeoglobi
Euryarchaeota
Bathyarchaeota
Archaea
76G-FAM006285
Nitrososphaerales
Nitrososphaerales
Nitrosocaldales
Thaumarchaeota
Thermoprotei
76G-FAM006285
Crenarchaeota

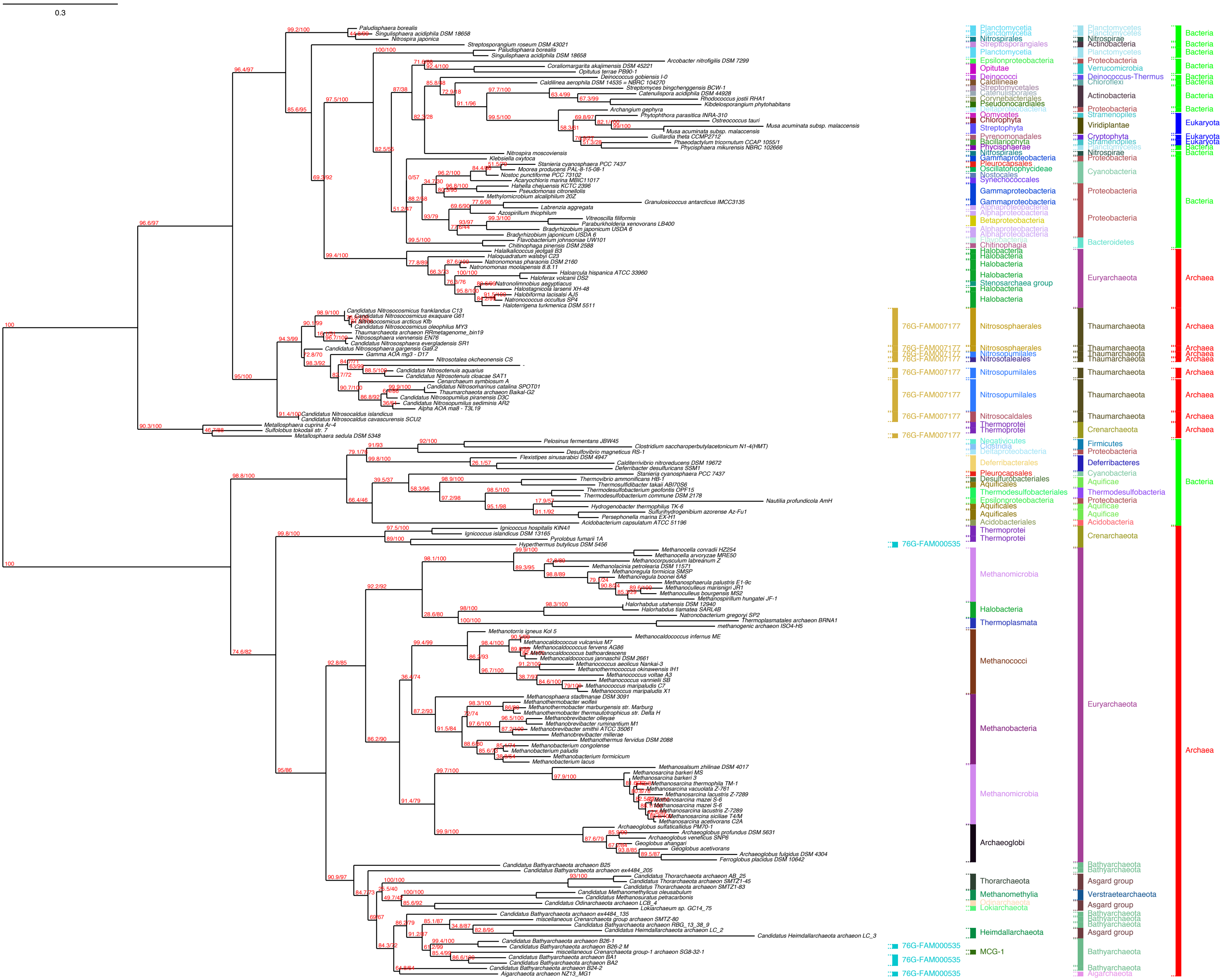
0.3
Paludisphaera borealis
Singulisphaera acidiphila DSM 18658
Nitrospira japonica
Streptosporangium roseum DSM 43021
Paludisphaera borealis
Singulisphaera acidiphila DSM 18658
Arcobacter nitrofigilis DSM 7299
Coraliomargarita akajimensis DSM 45221
Opitutus terrae PB90-1
Deinococcus gobiensis I-0
Caldilinea aerophila DSM 14535 = NBRC 104270
Streptomyces bingchenggensis BCW-1
Catenulispora acidiphila DSM 44928
Rhodococcus jostii RHA1
Kibdelosporangium phytohabitans
Archangium gephyra
Phytophthora parasitica INRA-310
Ostreococcus tauri
Musa acuminata subsp. malaccensis
Musa acuminata subsp. malaccensis
Guillardia theta CCMP2712
Phaeodactylum tricornutum CCAP 1055/1
Phycisphaera mikurensis NBRC 102666
Nitrospira moscoviensis
Klebsiella oxytoca
Stanieria cyanosphaera PCC 7437
Moorea producens PAL-8-15-08-1
Nostoc punctiforme PCC 73102
Acaryochloris marina MBIC11017
Hahella chejuensis KCTC 2396
Pseudomonas citronellolis
Methylomicrobium alcaliphilum 20Z
Granulosicoccus antarcticus IMCC3135
Labrenzia aggregata
Azospirillum thiophilum
Vitreoscilla filiformis
Paraburkholderia xenovorans LB400
Bradyrhizobium japonicum USDA 6
Bradyrhizobium japonicum USDA 6
Flavobacterium johnsoniae UW101
Chitinophaga pinensis DSM 2588
Halalkalicoccus jeotgali B3
Haloquadratum walsbyi C23
Natronomonas pharaonis DSM 2160
Natronomonas moolapensis 8.8.11
Haloarcula hispanica ATCC 33960
Haloferax volcanii DS2
Natronolimnobius aegyptiacus
Halostagnicola larsenii XH-48
Halobiforma lacisalsi AJ5
Natronococcus occultus SP4
Haloterrigena turkmenica DSM 5511
Candidatus Nitrosocosmicus franklandus C13
Candidatus Nitrosocosmicus exaquare G61
Candidatus Nitrosocosmicus arcticus Kfb
Candidatus Nitrosocosmicus oleophilus MY3
Thaumarchaeota archaeon RRmetagenome_bin19
Nitrososphaera viennensis EN76
Candidatus Nitrososphaera evergladensis SR1
Candidatus Nitrososphaera gargensis Ga9.2
Gamma AOA mg3 - D17
Nitrosotalea okcheonensis CS
Candidatus Nitrosotenuis aquarius
Candidatus Nitrosotenuis cloacae SAT1
Cenarchaeum symbiosum A
Candidatus Nitrosomarinus catalina SPOT01
Thaumarchaeota archaeon Baikal-G2
Candidatus Nitrosopumilus piranensis D3C
Candidatus Nitrosopumilus sediminis AR2
Alpha AOA ma8 - T3L19
Candidatus Nitrosocaldus islandicus
Candidatus Nitrosocaldus cavascurensis SCU2
Metallosphaera cuprina Ar-4
Sulfolobus tokodaii str. 7
Metallosphaera sedula DSM 5348
Pelosinus fermentans JBW45
Clostridium saccharoperbutylacetonicum N1-4(HMT)
Desulfovibrio magneticus RS-1
Flexistipes sinusarabici DSM 4947
Calditerrivibrio nitroreducens DSM 19672
Deferribacter desulfuricans SSM1
Stanieria cyanosphaera PCC 7437
Thermovibrio ammonificans HB-1
Thermosulfidibacter takaii ABI70S6
Thermodesulfobacterium geofontis OPF15
Thermodesulfobacterium commune DSM 2178
Nautilia profundicola AmH
Hydrogenobacter thermophilus TK-6
Sulfurihydrogenibium azorense Az-Fu1
Persephonella marina EX-H1
Acidobacterium capsulatum ATCC 51196
Ignicoccus hospitalis KIN4/I
Ignicoccus islandicus DSM 13165
Pyrolobus fumarii 1A
Hyperthermus butylicus DSM 5456
Methanocella conradii HZ254
Methanocella arvoryzae MRE50
Methanocorpusculum labreanum Z
Methanolacinia petrolearia DSM 11571
Methanoregula formicica SMSP
Methanoregula boonei 6A8
Methanosphaerula palustris E1-9c
Methanoculleus marisnigri JR1
Methanoculleus bourgensis MS2
Methanospirillum hungatei JF-1
Halorhabdus utahensis DSM 12940
Halorhabdus tiamatea SARL4B
Natronobacterium gregoryi SP2
Thermoplasmatales archaeon BRNA1
methanogenic archaeon ISO4-H5
Methanotorris igneus Kol 5
Methanocaldococcus infernus ME
Methanocaldococcus vulcanius M7
Methanocaldococcus fervens AG86
Methanocaldococcus bathoardescens
Methanocaldococcus jannaschii DSM 2661
Methanococcus aeolicus Nankai-3
Methanothermococcus okinawensis IH1
Methanococcus voltae A3
Methanococcus vannielii SB
Methanococcus maripaludis C7
Methanococcus maripaludis X1
Methanosphaera stadtmanae DSM 3091
Methanothermobacter wolfeii
Methanothermobacter marburgensis str. Marburg
Methanothermobacter thermautotrophicus str. Delta H
Methanobrevibacter olleyae
Methanobrevibacter ruminantium M1
Methanobrevibacter smithii ATCC 35061
Methanobrevibacter millerae
Methanothermus fervidus DSM 2088
Methanobacterium congolense
Methanobacterium paludis
Methanobacterium formicicum
Methanobacterium lacus
Methanosalsum zhilinae DSM 4017
Methanosarcina barkeri MS
Methanosarcina barkeri 3
Methanosarcina thermophila TM-1
Methanosarcina vacuolata Z-761
Methanosarcina lacustris Z-7289
Methanosarcina mazei S-6
Methanosarcina mazei S-6
Methanosarcina lacustris Z-7289
Methanosarcina siciliae T4/M
Methanosarcina acetivorans C2A
Archaeoglobus sulfaticallidus PM70-1
Archaeoglobus profundus DSM 5631
Archaeoglobus veneficus SNP6
Geoglobus ahangari
Geoglobus acetivorans
Archaeoglobus fulgidus DSM 4304
Ferroglobus placidus DSM 10642
Candidatus Bathyarchaeota archaeon B25
Candidatus Bathyarchaeota archaeon ex4484_205
Candidatus Thorarchaeota archaeon AB_25
Candidatus Thorarchaeota archaeon SMTZ1-45
Candidatus Thorarchaeota archaeon SMTZ1-83
Candidatus Methanomethylicus oleusabulum
Candidatus Methanosuratus petracarbonis
Candidatus Odinarchaeota archaeon LCB_4
Lokiarchaeum sp. GC14_75
Candidatus Bathyarchaeota archaeon ex4484_135
miscellaneous Crenarchaeota group archaeon SMTZ-80
Candidatus Bathyarchaeota archaeon RBG_13_38_9
Candidatus Heimdallarchaeota archaeon LC_2
Candidatus Heimdallarchaeota archaeon LC_3
Candidatus Bathyarchaeota archaeon B26-1
Candidatus Bathyarchaeota archaeon B26-2 M
miscellaneous Crenarchaeota group-1 archaeon SG8-32-1
Candidatus Bathyarchaeota archaeon BA1
Candidatus Bathyarchaeota archaeon BA2
Candidatus Bathyarchaeota archaeon B24-2
Aigarchaeota archaeon NZ13_MG1
76G-FAM007177
76G-FAM000535
Planctomycetia
Nitrospirales
Streptosporangiales
Epsilonproteobacteria
Opitutae
Deinococci
Caldilineae
Streptomycetales
Catenulisporales
Corynebacteriales
Pseudonocardiales
Deltaproteobacteria
Oomycetes
Chlorophyta
Streptophyta
Pyrenomonadales
Bacillariophyta
Phycisphaerae
Gammaproteobacteria
Pleurocapsales
Oscillatoriophycideae
Nostocales
Synechococcales
Alphaproteobacteria
Betaproteobacteria
Flavobacteriia
Chitinophagia
Halobacteria
Stenosarchaea group
Nitrososphaerales
Nitrosopumilales
Nitrosotaleales
Nitrosocaldales
Thermoprotei
Negativicutes
Clostridia
Deferribacterales
Desulfurobacteriales
Aquificales
Thermodesulfobacteriales
Acidobacteriales
Methanomicrobia
Thermoplasmata
Methanococci
Methanobacteria
Archaeoglobi
Thorarchaeota
Methanomethylia
Odinarchaeota
Lokiarchaeota
Heimdallarchaeota
MCG-1
Planctomycetes
Nitrospirae
Actinobacteria
Proteobacteria
Verrucomicrobia
Deinococcus-Thermus
Chloroflexi
Stramenopiles
Viridiplantae
Cryptophyta
Cyanobacteria
Bacteroidetes
Euryarchaeota
Thaumarchaeota
Crenarchaeota
Firmicutes
Deferribacteres
Aquificae
Thermodesulfobacteria
Acidobacteria
Bathyarchaeota
Asgard group
Verstraetearchaeota
Aigarchaeota
Bacteria
Eukaryota
Archaea

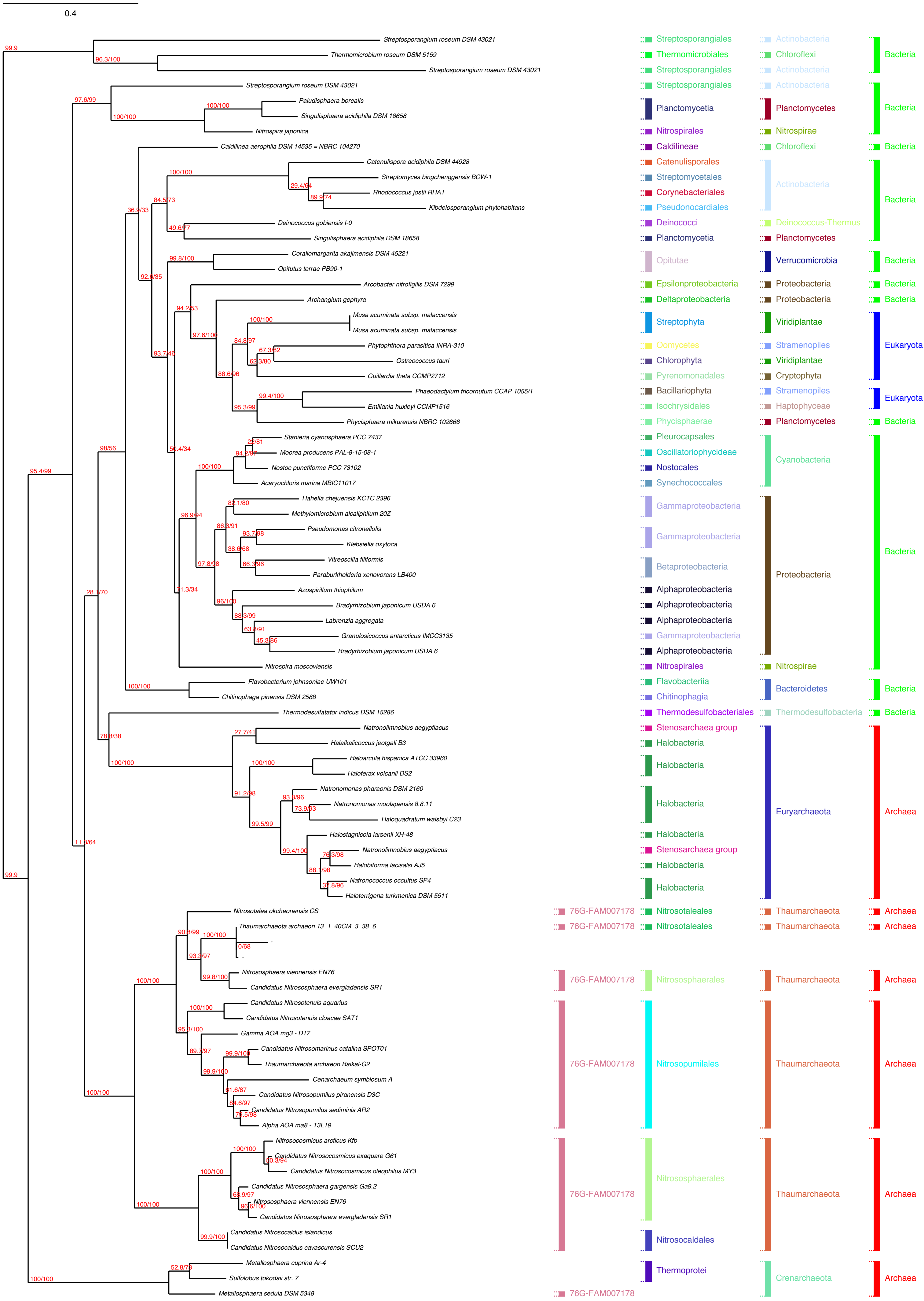
0.4
99.9
96.3/100
97.6/99
100/100
100/100
36.9/33
84.5/73
100/100
29.4/64
89.9/74
49.6/77
92.6/35
99.8/100
94.2/53
97.6/100
100/100
84.8/97
67.3/82
62.3/80
88.6/96
99.4/100
95.3/99
93.7/46
98/56
50.4/34
28/81
94.2/97
100/100
96.9/94
82.1/80
86.3/91
93.7/98
38.6/68
97.8/98
66.3/96
71.3/34
96/100
88.3/99
63.8/91
45.3/86
95.4/99
28.1/70
100/100
78.8/38
27.7/41
100/100
100/100
91.2/98
93.8/96
73.9/93
99.5/99
99.4/100
76.3/98
88.1/98
37.8/96
11.8/64
99.9
90.8/99
100/100
0/68
93.3/97
99.8/100
100/100
100/100
95.8/100
89.7/97
99.9/100
99.9/100
61.6/87
84.6/97
79.5/98
100/100
100/100
100/100
50.3/94
100/100
66.9/97
96.6/100
99.9/100
52.8/78
100/100
Streptosporangium roseum DSM 43021 | Streptosporangiales | Actinobacteria | Bacteria
Thermomicrobium roseum DSM 5159 | Thermomicrobiales | Chloroflexi
Streptosporangium roseum DSM 43021 | Streptosporangiales | Actinobacteria
Streptosporangium roseum DSM 43021 | Streptosporangiales | Actinobacteria | Bacteria
Paludisphaera borealis
Singulisphaera acidiphila DSM 18658 | Planctomycetia | Planctomycetes
Nitrospira japonica | Nitrospirales | Nitrospirae
Caldilinea aerophila DSM 14535 = NBRC 104270 | Caldilineae | Chloroflexi | Bacteria
Catenulispora acidiphila DSM 44928 | Catenulisporales
Streptomyces bingchenggensis BCW-1 | Streptomycetales
Rhodococcus jostii RHA1 | Corynebacteriales | Actinobacteria | Bacteria
Kibdelosporangium phytohabitans | Pseudonocardiales
Deinococcus gobiensis I-0 | Deinococci | Deinococcus-Thermus
Singulisphaera acidiphila DSM 18658 | Planctomycetia | Planctomycetes
Coraliomargarita akajimensis DSM 45221
Opitutus terrae PB90-1 | Opitutae | Verrucomicrobia | Bacteria
Arcobacter nitrofigilis DSM 7299 | Epsilonproteobacteria | Proteobacteria | Bacteria
Archangium gephyra | Deltaproteobacteria | Proteobacteria | Bacteria
Musa acuminata subsp. malaccensis
Musa acuminata subsp. malaccensis | Streptophyta | Viridiplantae
Phytophthora parasitica INRA-310 | Oomycetes | Stramenopiles | Eukaryota
Ostreococcus tauri | Chlorophyta | Viridiplantae
Guillardia theta CCMP2712 | Pyrenomonadales | Cryptophyta
Phaeodactylum tricornutum CCAP 1055/1 | Bacillariophyta | Stramenopiles
Emiliania huxleyi CCMP1516 | Isochrysidales | Haptophyceae | Eukaryota
Phycisphaera mikurensis NBRC 102666 | Phycisphaerae | Planctomycetes | Bacteria
Stanieria cyanosphaera PCC 7437 | Pleurocapsales
Moorea producens PAL-8-15-08-1 | Oscillatoriophycideae | Cyanobacteria
Nostoc punctiforme PCC 73102 | Nostocales
Acaryochloris marina MBIC11017 | Synechococcales
Hahella chejuensis KCTC 2396
Methylomicrobium alcaliphilum 20Z | Gammaproteobacteria
Pseudomonas citronellolis
Klebsiella oxytoca | Gammaproteobacteria
Vitreoscilla filiformis
Paraburkholderia xenovorans LB400 | Betaproteobacteria | Proteobacteria | Bacteria
Azospirillum thiophilum | Alphaproteobacteria
Bradyrhizobium japonicum USDA 6 | Alphaproteobacteria
Labrenzia aggregata | Alphaproteobacteria
Granulosicoccus antarcticus IMCC3135 | Gammaproteobacteria
Bradyrhizobium japonicum USDA 6 | Alphaproteobacteria
Nitrospira moscoviensis | Nitrospirales | Nitrospirae
Flavobacterium johnsoniae UW101 | Flavobacteriia
Chitinophaga pinensis DSM 2588 | Chitinophagia | Bacteroidetes | Bacteria
Thermodesulfatator indicus DSM 15286 | Thermodesulfobacteriales | Thermodesulfobacteria | Bacteria
Natronolimnobius aegyptiacus | Stenosarchaea group
Halalkalicoccus jeotgali B3 | Halobacteria
Haloarcula hispanica ATCC 33960
Haloferax volcanii DS2 | Halobacteria
Natronomonas pharaonis DSM 2160
Natronomonas moolapensis 8.8.11 | Halobacteria | Euryarchaeota | Archaea
Haloquadratum walsbyi C23
Halostagnicola larsenii XH-48 | Halobacteria
Natronolimnobius aegyptiacus | Stenosarchaea group
Halobiforma lacisalsi AJ5 | Halobacteria
Natronococcus occultus SP4
Haloterrigena turkmenica DSM 5511 | Halobacteria
Nitrosotalea okcheonensis CS | 76G-FAM007178 | Nitrosotaleales | Thaumarchaeota | Archaea
Thaumarchaeota archaeon 13_1_40CM_3_38_6 | 76G-FAM007178 | Nitrosotaleales | Thaumarchaeota | Archaea
-
-
Nitrososphaera viennensis EN76
Candidatus Nitrososphaera evergladensis SR1 | 76G-FAM007178 | Nitrososphaerales | Thaumarchaeota | Archaea
Candidatus Nitrosotenuis aquarius
Candidatus Nitrosotenuis cloacae SAT1
Gamma AOA mg3 - D17
Candidatus Nitrosomarinus catalina SPOT01
Thaumarchaeota archaeon Baikal-G2 | 76G-FAM007178 | Nitrosopumilales | Thaumarchaeota | Archaea
Cenarchaeum symbiosum A
Candidatus Nitrosopumilus piranensis D3C
Candidatus Nitrosopumilus sediminis AR2
Alpha AOA ma8 - T3L19
Nitrosocosmicus arcticus Kfb
Candidatus Nitrosocosmicus exaquare G61
Candidatus Nitrosocosmicus oleophilus MY3 | Nitrososphaerales
Candidatus Nitrososphaera gargensis Ga9.2 | 76G-FAM007178 | Thaumarchaeota | Archaea
Nitrososphaera viennensis EN76
Candidatus Nitrososphaera evergladensis SR1
Candidatus Nitrosocaldus islandicus
Candidatus Nitrosocaldus cavascurensis SCU2 | Nitrosocaldales
Metallosphaera cuprina Ar-4
Sulfolobus tokodaii str. 7 | Thermoprotei | Crenarchaeota | Archaea
Metallosphaera sedula DSM 5348 | 76G-FAM007178

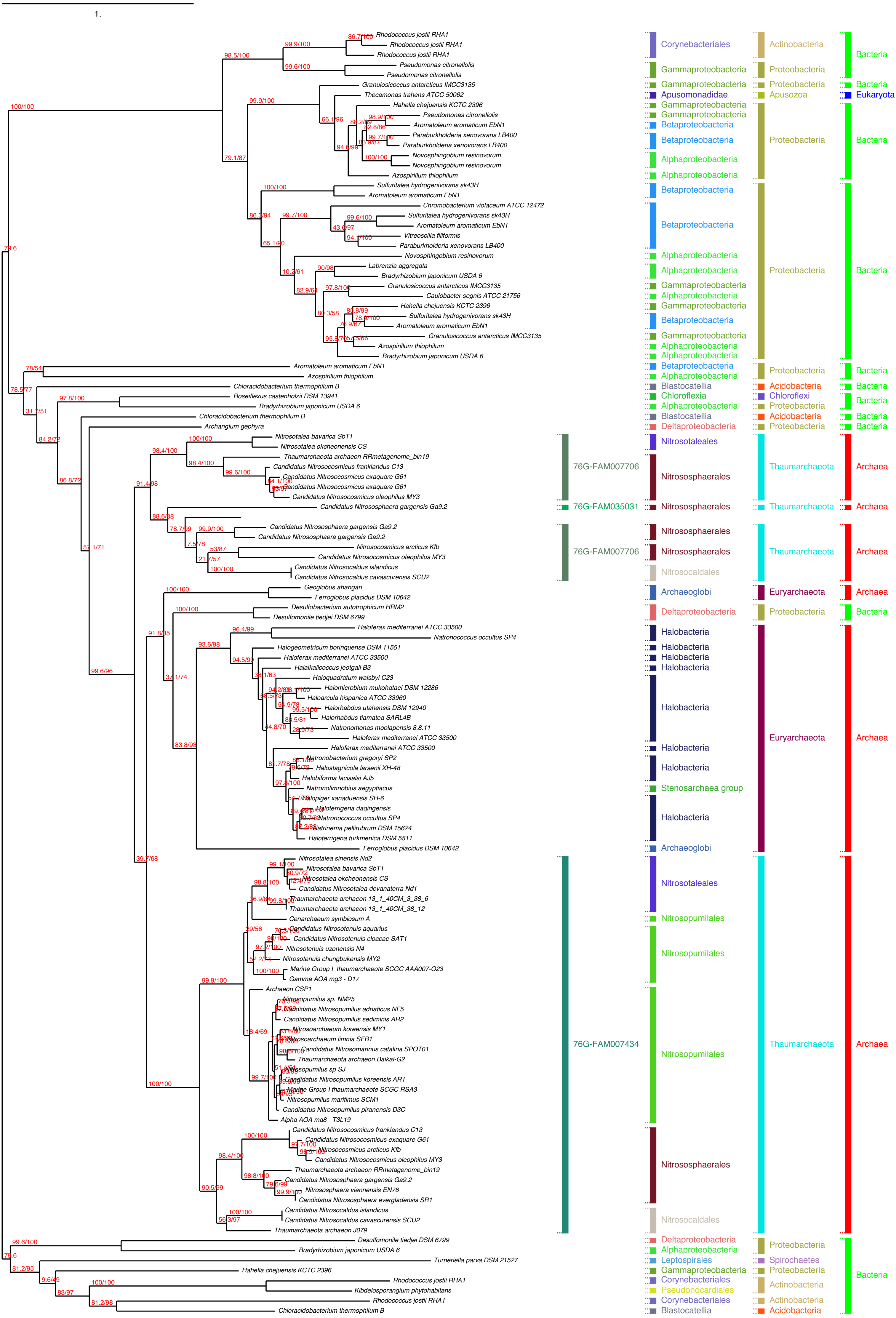

1.
Rhodococcus jostii RHA1
Rhodococcus jostii RHA1
Rhodococcus jostii RHA1
Pseudomonas citronellolis
Pseudomonas citronellolis
Granulosicoccus antarcticus IMCC3135
Thecamonas trahens ATCC 50062
Hahella chejuensis KCTC 2396
Pseudomonas citronellolis
Aromatoleum aromaticum EbN1
Paraburkholderia xenovorans LB400
Paraburkholderia xenovorans LB400
Novosphingobium resinovorum
Novosphingobium resinovorum
Azospirillum thiophilum
Sulfuritalea hydrogenivorans sk43H
Aromatoleum aromaticum EbN1
Chromobacterium violaceum ATCC 12472
Sulfuritalea hydrogenivorans sk43H
Aromatoleum aromaticum EbN1
Vitreoscilla filiformis
Paraburkholderia xenovorans LB400
Novosphingobium resinovorum
Labrenzia aggregata
Bradyrhizobium japonicum USDA 6
Granulosicoccus antarcticus IMCC3135
Caulobacter segnis ATCC 21756
Hahella chejuensis KCTC 2396
Sulfuritalea hydrogenivorans sk43H
Aromatoleum aromaticum EbN1
Granulosicoccus antarcticus IMCC3135
Azospirillum thiophilum
Bradyrhizobium japonicum USDA 6
Aromatoleum aromaticum EbN1
Azospirillum thiophilum
Chloracidobacterium thermophilum B
Roseiflexus castenholzii DSM 13941
Bradyrhizobium japonicum USDA 6
Chloracidobacterium thermophilum B
Archangium gephyra
Nitrosotalea bavarica SbT1
Nitrosotalea okcheonensis CS
Thaumarchaeota archaeon RRmetagenome_bin19
Candidatus Nitrosocosmicus franklandus C13
Candidatus Nitrosocosmicus exaquare G61
Candidatus Nitrosocosmicus exaquare G61
Candidatus Nitrosocosmicus oleophilus MY3
Candidatus Nitrososphaera gargensis Ga9.2
Candidatus Nitrososphaera gargensis Ga9.2
Candidatus Nitrososphaera gargensis Ga9.2
Nitrosocosmicus arcticus Kfb
Candidatus Nitrosocosmicus oleophilus MY3
Candidatus Nitrosocaldus islandicus
Candidatus Nitrosocaldus cavascurensis SCU2
Geoglobus ahangari
Ferroglobus placidus DSM 10642
Desulfobacterium autotrophicum HRM2
Desulfomonile tiedjei DSM 6799
Haloferax mediterranei ATCC 33500
Natronococcus occultus SP4
Halogeometricum borinquense DSM 11551
Haloferax mediterranei ATCC 33500
Halalkalicoccus jeotgali B3
Haloquadratum walsbyi C23
Halomicrobium mukohataei DSM 12286
Haloarcula hispanica ATCC 33960
Halorhabdus utahensis DSM 12940
Halorhabdus tiamatea SARL4B
Natronomonas moolapensis 8.8.11
Haloferax mediterranei ATCC 33500
Haloferax mediterranei ATCC 33500
Natronobacterium gregoryi SP2
Halostagnicola larsenii XH-48
Halobiforma lacisalsi AJ5
Natronolimnobius aegyptiacus
Halopiger xanaduensis SH-6
Haloterrigena daqingensis
Natronococcus occultus SP4
Natrinema pellirubrum DSM 15624
Haloterrigena turkmenica DSM 5511
Ferroglobus placidus DSM 10642
Nitrosotalea sinensis Nd2
Nitrosotalea bavarica SbT1
Nitrosotalea okcheonensis CS
Candidatus Nitrosotalea devanaterra Nd1
Thaumarchaeota archaeon 13_1_40CM_3_38_6
Thaumarchaeota archaeon 13_1_40CM_38_12
Cenarchaeum symbiosum A
Candidatus Nitrosotenuis aquarius
Candidatus Nitrosotenuis cloacae SAT1
Nitrosotenuis uzonensis N4
Nitrosotenuis chungbukensis MY2
Marine Group I thaumarchaeote SCGC AAA007-O23
Gamma AOA mg3 - D17
Archaeon CSP1
Nitrosopumilus sp. NM25
Candidatus Nitrosopumilus adriaticus NF5
Candidatus Nitrosopumilus sediminis AR2
Nitrosoarchaeum koreensis MY1
Nitrosoarchaeum limnia SFB1
Candidatus Nitrosomarinus catalina SPOT01
Thaumarchaeota archaeon Baikal-G2
Nitrosopumilus sp SJ
Candidatus Nitrosopumilus koreensis AR1
Marine Group I thaumarchaeote SCGC RSA3
Nitrosopumilus maritimus SCM1
Candidatus Nitrosopumilus piranensis D3C
Alpha AOA ma8 - T3L19
Candidatus Nitrosocosmicus franklandus C13
Candidatus Nitrosocosmicus exaquare G61
Nitrosocosmicus arcticus Kfb
Candidatus Nitrosocosmicus oleophilus MY3
Thaumarchaeota archaeon RRmetagenome_bin19
Candidatus Nitrososphaera gargensis Ga9.2
Nitrososphaera viennensis EN76
Candidatus Nitrososphaera evergladensis SR1
Candidatus Nitrosocaldus islandicus
Candidatus Nitrosocaldus cavascurensis SCU2
Thaumarchaeota archaeon J079
Desulfomonile tiedjei DSM 6799
Bradyrhizobium japonicum USDA 6
Turneriella parva DSM 21527
Hahella chejuensis KCTC 2396
Rhodococcus jostii RHA1
Kibdelosporangium phytohabitans
Rhodococcus jostii RHA1
Chloracidobacterium thermophilum B
76G-FAM007706
76G-FAM035031
76G-FAM007706
76G-FAM007434
Corynebacteriales
Gammaproteobacteria
Gammaproteobacteria
Apusomonadidae
Gammaproteobacteria
Gammaproteobacteria
Betaproteobacteria
Betaproteobacteria
Alphaproteobacteria
Alphaproteobacteria
Betaproteobacteria
Betaproteobacteria
Alphaproteobacteria
Alphaproteobacteria
Gammaproteobacteria
Alphaproteobacteria
Gammaproteobacteria
Betaproteobacteria
Gammaproteobacteria
Alphaproteobacteria
Alphaproteobacteria
Betaproteobacteria
Alphaproteobacteria
Blastocatellia
Chloroflexia
Alphaproteobacteria
Blastocatellia
Deltaproteobacteria
Nitrosotaleales
Nitrososphaerales
Nitrososphaerales
Nitrososphaerales
Nitrososphaerales
Nitrosocaldales
Archaeoglobi
Deltaproteobacteria
Halobacteria
Halobacteria
Halobacteria
Halobacteria
Halobacteria
Halobacteria
Halobacteria
Stenosarchaea group
Halobacteria
Archaeoglobi
Nitrosotaleales
Nitrosopumilales
Nitrosopumilales
Nitrosopumilales
Nitrososphaerales
Nitrosocaldales
Deltaproteobacteria
Alphaproteobacteria
Leptospirales
Gammaproteobacteria
Corynebacteriales
Pseudonocardiales
Corynebacteriales
Blastocatellia
Actinobacteria
Proteobacteria
Proteobacteria
Apusozoa
Proteobacteria
Proteobacteria
Proteobacteria
Acidobacteria
Chloroflexi
Proteobacteria
Acidobacteria
Proteobacteria
Thaumarchaeota
Thaumarchaeota
Thaumarchaeota
Euryarchaeota
Proteobacteria
Euryarchaeota
Thaumarchaeota
Proteobacteria
Spirochaetes
Proteobacteria
Actinobacteria
Actinobacteria
Acidobacteria
Bacteria
Bacteria
Eukaryota
Bacteria
Bacteria
Bacteria
Bacteria
Bacteria
Bacteria
Bacteria
Archaea
Archaea
Archaea
Archaea
Bacteria
Archaea
Archaea
Bacteria

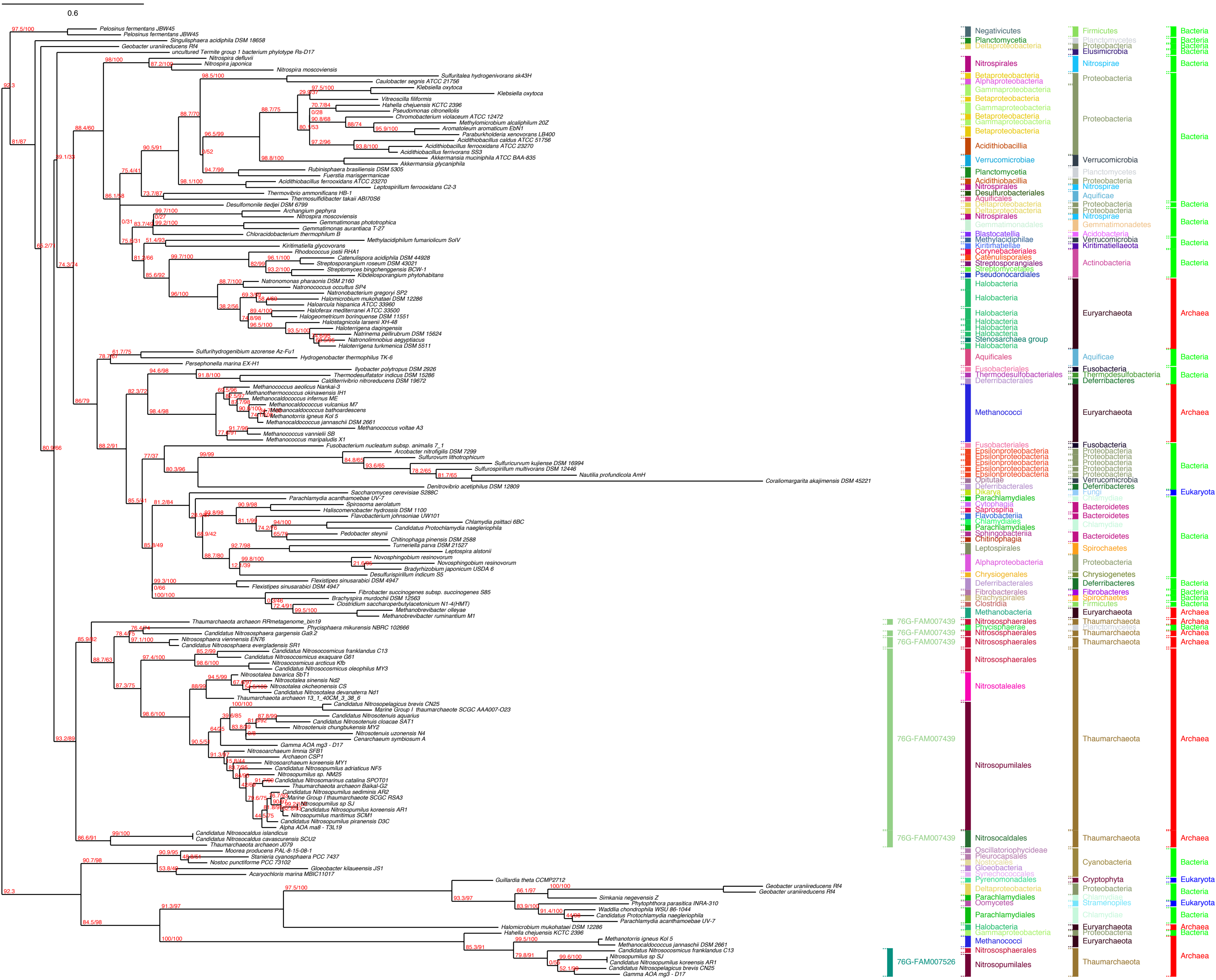
0.6
Pelosinus fermentans JBW45
Pelosinus fermentans JBW45
Singulisphaera acidiphila DSM 18658
Geobacter uraniireducens Rf4
uncultured Termite group 1 bacterium phylotype Rs-D17
Nitrospira defluvii
Nitrospira japonica
Nitrospira moscoviensis
Sulfuritalea hydrogenivorans sk43H
Caulobacter segnis ATCC 21756
Klebsiella oxytoca
Klebsiella oxytoca
Vitreoscilla filiformis
Hahella chejuensis KCTC 2396
Pseudomonas citronellolis
Chromobacterium violaceum ATCC 12472
Methylomicrobium alcaliphilum 20Z
Aromatoleum aromaticum EbN1
Paraburkholderia xenovorans LB400
Acidithiobacillus caldus ATCC 51756
Acidithiobacillus ferrooxidans ATCC 23270
Acidithiobacillus ferrivorans SS3
Akkermansia muciniphila ATCC BAA-835
Akkermansia glycaniphila
Rubinisphaera brasiliensis DSM 5305
Fuerstia marisgermanicae
Acidithiobacillus ferrooxidans ATCC 23270
Leptospirillum ferrooxidans C2-3
Thermovibrio ammonificans HB-1
Thermosulfidibacter takaii ABI70S6
Desulfomonile tiedjei DSM 6799
Archangium gephyra
Nitrospira moscoviensis
Gemmatimonas phototrophica
Gemmatimonas aurantiaca T-27
Chloracidobacterium thermophilum B
Methylacidiphilum fumariolicum SolV
Kiritimatiella glycovorans
Rhodococcus jostii RHA1
Catenulispora acidiphila DSM 44928
Streptosporangium roseum DSM 43021
Streptomyces bingchenggensis BCW-1
Kibdelosporangium phytohabitans
Natronomonas pharaonis DSM 2160
Natronococcus occultus SP4
Natronobacterium gregoryi SP2
Halomicrobium mukohataei DSM 12286
Haloarcula hispanica ATCC 33960
Haloferax mediterranei ATCC 33500
Halogeometricum borinquense DSM 11551
Halostagnicola larsenii XH-48
Haloterrigena daqingensis
Natrinema pellirubrum DSM 15624
Natronolimnobius aegyptiacus
Haloterrigena turkmenica DSM 5511
Sulfurihydrogenibium azorense Az-Fu1
Hydrogenobacter thermophilus TK-6
Persephonella marina EX-H1
Ilyobacter polytropus DSM 2926
Thermodesulfatator indicus DSM 15286
Calditerrivibrio nitroreducens DSM 19672
Methanococcus aeolicus Nankai-3
Methanothermococcus okinawensis IH1
Methanocaldococcus infernus ME
Methanocaldococcus vulcanius M7
Methanocaldococcus bathoardescens
Methanotorris igneus Kol 5
Methanocaldococcus jannaschii DSM 2661
Methanococcus voltae A3
Methanococcus vannielii SB
Methanococcus maripaludis X1
Fusobacterium nucleatum subsp. animalis 7_1
Arcobacter nitrofigilis DSM 7299
Sulfurovum lithotrophicum
Sulfuricurvum kujiense DSM 16994
Sulfurospirillum multivorans DSM 12446
Nautilia profundicola AmH
Coraliomargarita akajimensis DSM 45221
Denitrovibrio acetiphilus DSM 12809
Saccharomyces cerevisiae S288C
Parachlamydia acanthamoebae UV-7
Spirosoma aerolatum
Haliscomenobacter hydrossis DSM 1100
Flavobacterium johnsoniae UW101
Chlamydia psittaci 6BC
Candidatus Protochlamydia naegleriophila
Pedobacter steynii
Chitinophaga pinensis DSM 2588
Turneriella parva DSM 21527
Leptospira alstonii
Novosphingobium resinovorum
Novosphingobium resinovorum
Bradyrhizobium japonicum USDA 6
Desulfurispirillum indicum S5
Flexistipes sinusarabici DSM 4947
Flexistipes sinusarabici DSM 4947
Fibrobacter succinogenes subsp. succinogenes S85
Brachyspira murdochii DSM 12563
Clostridium saccharoperbutylacetonicum N1-4(HMT)
Methanobrevibacter olleyae
Methanobrevibacter ruminantium M1
Thaumarchaeota archaeon RRmetagenome_bin19
Phycisphaera mikurensis NBRC 102666
Candidatus Nitrososphaera gargensis Ga9.2
Nitrososphaera viennensis EN76
Candidatus Nitrososphaera evergladensis SR1
Candidatus Nitrosocosmicus franklandus C13
Candidatus Nitrosocosmicus exaquare G61
Nitrosocosmicus arcticus Kfb
Candidatus Nitrosocosmicus oleophilus MY3
Nitrosotalea bavarica SbT1
Nitrosotalea sinensis Nd2
Nitrosotalea okcheonensis CS
Candidatus Nitrosotalea devanaterra Nd1
Thaumarchaeota archaeon 13_1_40CM_3_38_6
Candidatus Nitrosopelagicus brevis CN25
Marine Group I thaumarchaeote SCGC AAA007-O23
Candidatus Nitrosotenuis aquarius
Candidatus Nitrosotenuis cloacae SAT1
Nitrosotenuis chungbukensis MY2
Nitrosotenuis uzonensis N4
Cenarchaeum symbiosum A
Gamma AOA mg3 - D17
Nitrosoarchaeum limnia SFB1
Archaeon CSP1
Nitrosoarchaeum koreensis MY1
Candidatus Nitrosopumilus adriaticus NF5
Nitrosopumilus sp. NM25
Candidatus Nitrosomarinus catalina SPOT01
Thaumarchaeota archaeon Baikal-G2
Candidatus Nitrosopumilus sediminis AR2
Marine Group I thaumarchaeote SCGC RSA3
Nitrosopumilus sp SJ
Candidatus Nitrosopumilus koreensis AR1
Nitrosopumilus maritimus SCM1
Candidatus Nitrosopumilus piranensis D3C
Alpha AOA ma8 - T3L19
Candidatus Nitrosocaldus islandicus
Candidatus Nitrosocaldus cavascurensis SCU2
Thaumarchaeota archaeon J079
Moorea producens PAL-8-15-08-1
Stanieria cyanosphaera PCC 7437
Nostoc punctiforme PCC 73102
Gloeobacter kilaueensis JS1
Acaryochloris marina MBIC11017
Guillardia theta CCMP2712
Geobacter uraniireducens Rf4
Geobacter uraniireducens Rf4
Simkania negevensis Z
Phytophthora parasitica INRA-310
Waddlia chondrophila WSU 86-1044
Candidatus Protochlamydia naegleriophila
Parachlamydia acanthamoebae UV-7
Halomicrobium mukohataei DSM 12286
Hahella chejuensis KCTC 2396
Methanotorris igneus Kol 5
Methanocaldococcus jannaschii DSM 2661
Candidatus Nitrosocosmicus franklandus C13
Nitrosopumilus sp SJ
Candidatus Nitrosopumilus koreensis AR1
Candidatus Nitrosopelagicus brevis CN25
Gamma AOA mg3 - D17
76G-FAM007439
76G-FAM007439
76G-FAM007439
76G-FAM007439
76G-FAM007439
76G-FAM007526
Negativicutes
Planctomycetia
Deltaproteobacteria
Nitrospirales
Betaproteobacteria
Alphaproteobacteria
Gammaproteobacteria
Acidithiobacillia
Verrucomicrobiae
Desulfurobacteriales
Aquificales
Gemmatimonadales
Blastocatellia
Methylacidiphilae
Kiritimatiellae
Corynebacteriales
Catenulisporales
Streptosporangiales
Streptomycetales
Pseudonocardiales
Halobacteria
Stenosarchaea group
Fusobacteriales
Thermodesulfobacteriales
Deferribacterales
Methanococci
Epsilonproteobacteria
Opitutae
Dikarya
Parachlamydiales
Cytophagia
Saprospiria
Flavobacteriia
Chlamydiales
Sphingobacteriia
Chitinophagia
Leptospirales
Chrysiogenales
Fibrobacterales
Brachyspirales
Clostridia
Methanobacteria
Nitrososphaerales
Phycisphaerae
Nitrosotaleales
Nitrosopumilales
Nitrosocaldales
Oscillatoriophycideae
Pleurocapsales
Nostocales
Gloeobacteria
Synechococcales
Pyrenomonadales
Oomycetes
Firmicutes
Planctomycetes
Proteobacteria
Elusimicrobia
Nitrospirae
Verrucomicrobia
Aquificae
Gemmatimonadetes
Acidobacteria
Kiritimatiellaeota
Actinobacteria
Euryarchaeota
Fusobacteria
Thermodesulfobacteria
Deferribacteres
Fungi
Chlamydiae
Bacteroidetes
Spirochaetes
Chrysiogenetes
Fibrobacteres
Thaumarchaeota
Cyanobacteria
Cryptophyta
Stramenopiles
Bacteria
Archaea
Eukaryota

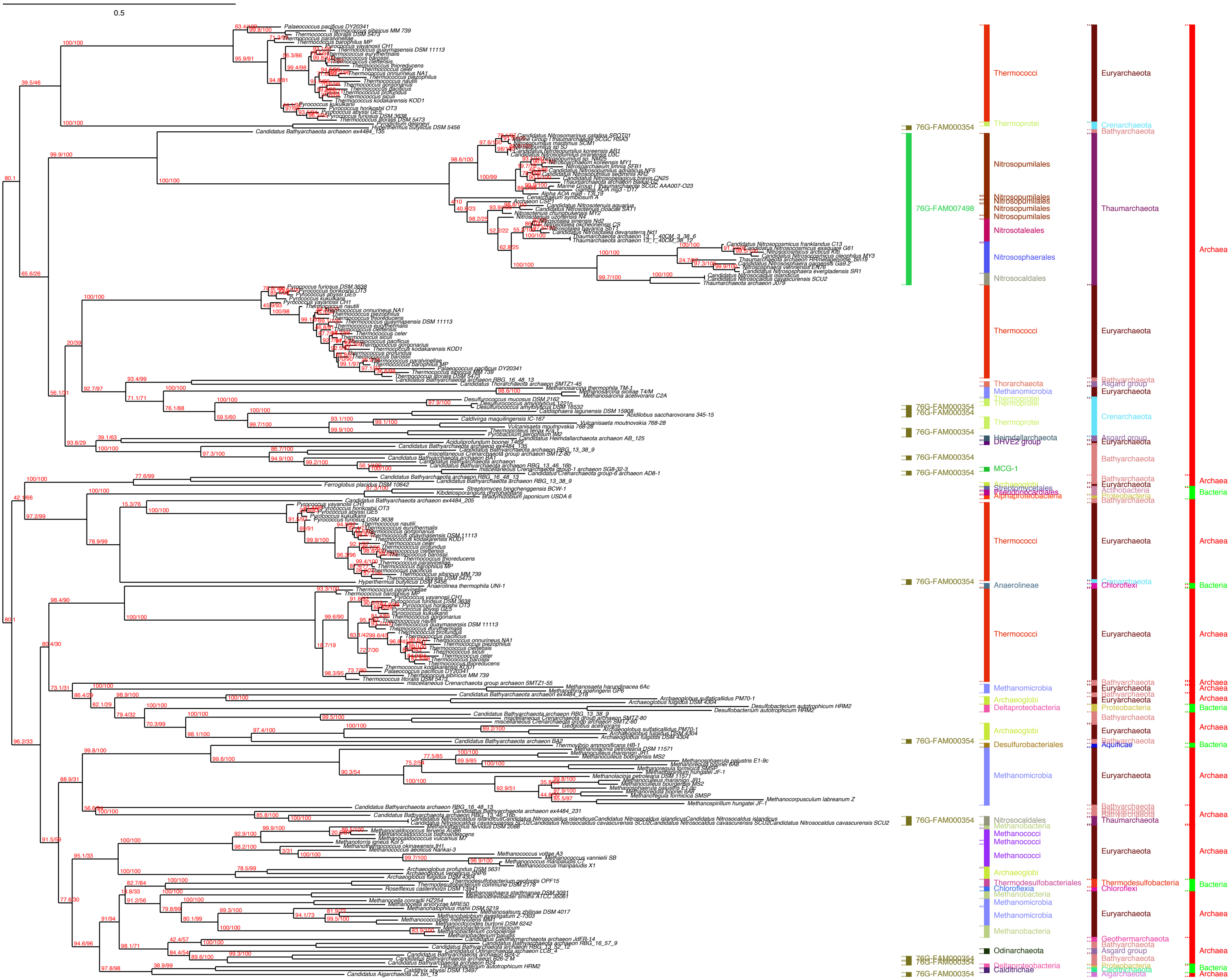
0.5
76G-FAM000354
76G-FAM007498
Thermococci
Thermoprotei
Nitrosopumilales
Nitrosotaleales
Nitrososphaerales
Nitrosocaldales
Thorarchaeota
Methanomicrobia
Heimdallarchaeota
DHVE2 group
MCG-1
Archaeoglobi
Streptomycetales
Pseudonocardiales
Alphaproteobacteria
Anaerolineae
Deltaproteobacteria
Desulfurobacteriales
Methanobacteria
Methanococci
Thermodesulfobacteriales
Chloroflexia
Odinarchaeota
Caldithrichae
Euryarchaeota
Crenarchaeota
Bathyarchaeota
Thaumarchaeota
Asgard group
Actinobacteria
Proteobacteria
Chloroflexi
Aquificae
Thermodesulfobacteria
Geothermarchaeota
Calditrichaeota
Aigarchaeota
Archaea
Bacteria

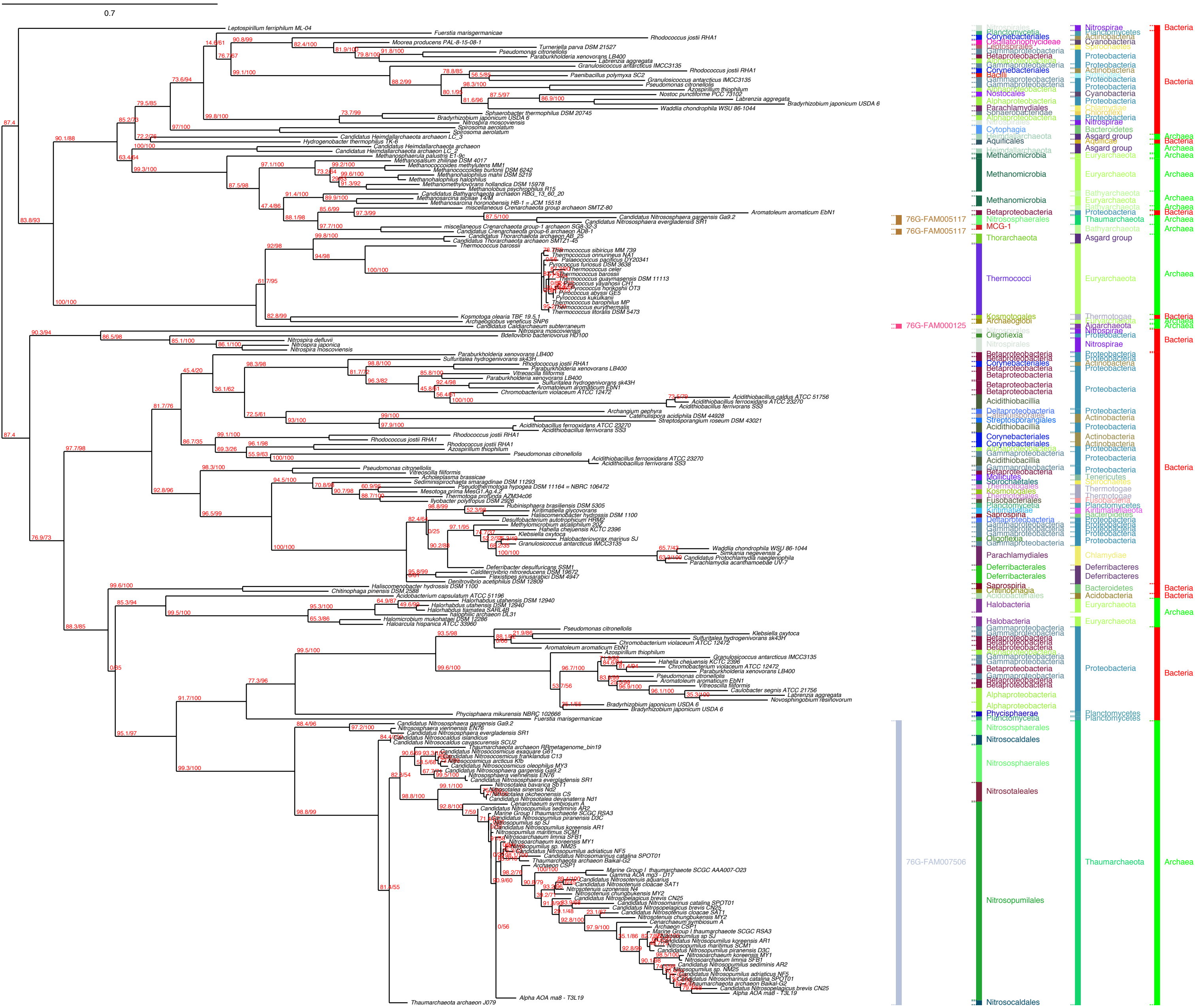
0.7
76G-FAM005117
76G-FAM005117
76G-FAM000125
76G-FAM007506
Methanomicrobia
Thermococci
Thorarchaeota
Nitrososphaerales
Nitrosocaldales
Nitrosotaleales
Nitrosopumilales
Parachlamydiales
Deferribacterales
Halobacteria
Alphaproteobacteria
Betaproteobacteria
Gammaproteobacteria
Proteobacteria
Euryarchaeota
Thaumarchaeota
Asgard group
Nitrospirae
Bacteria
Archaea

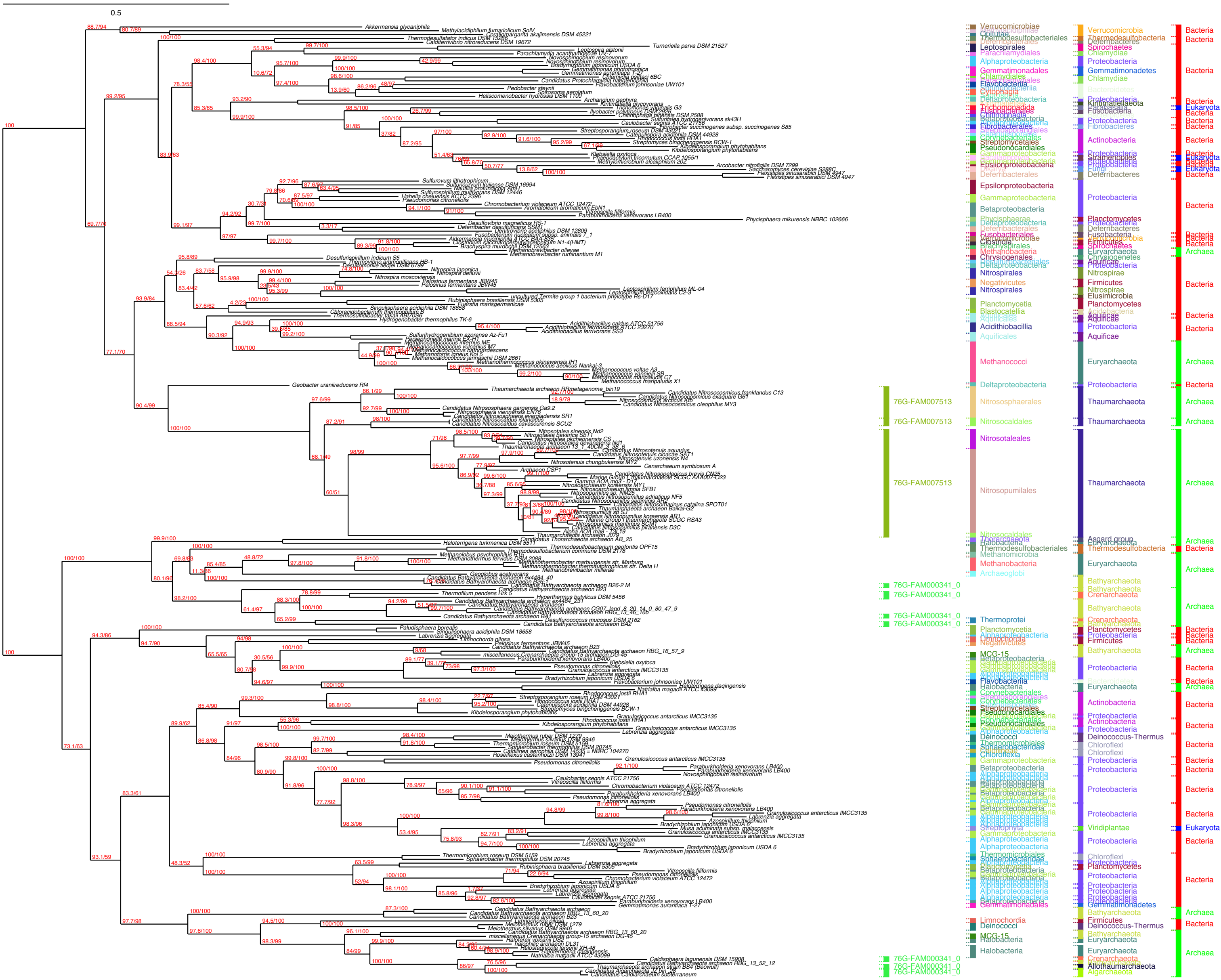

0.5
76G-FAM007513
76G-FAM000341_0
Nitrosotaleales
Nitrosopumilales
Nitrosocaldales
Nitrososphaerales
Methanococci
Thaumarchaeota
Euryarchaeota
Proteobacteria
Bathyarchaeota
Actinobacteria
Verrucomicrobia
Bacteria
Archaea
Eukaryota

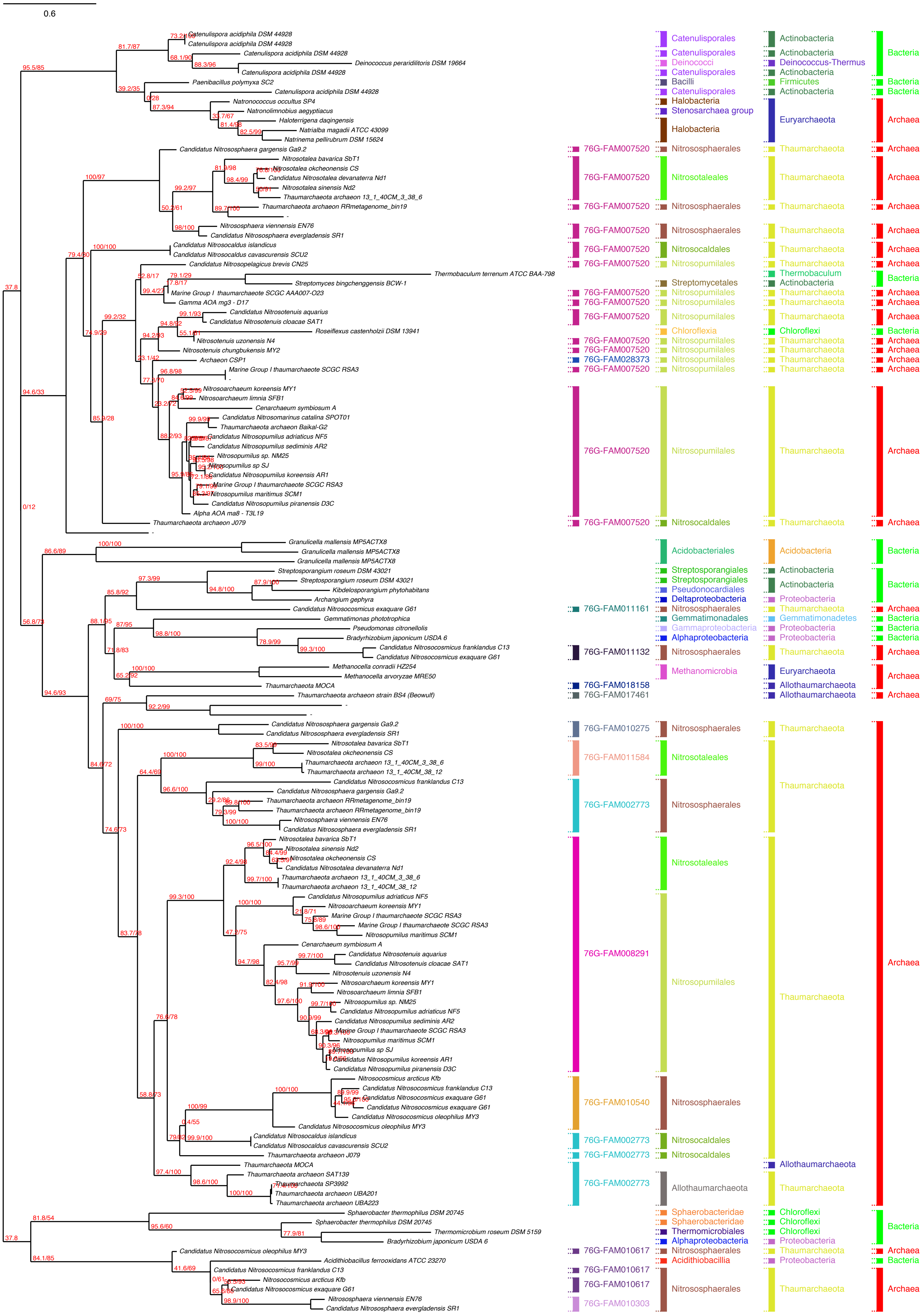

0.6
Catenulispora acidiphila DSM 44928
Catenulispora acidiphila DSM 44928
Catenulispora acidiphila DSM 44928
Deinococcus peraridilitoris DSM 19664
Catenulispora acidiphila DSM 44928
Paenibacillus polymyxa SC2
Catenulispora acidiphila DSM 44928
Natronococcus occultus SP4
Natronolimnobius aegyptiacus
Haloterrigena daqingensis
Natrialba magadii ATCC 43099
Natrinema pellirubrum DSM 15624
Candidatus Nitrososphaera gargensis Ga9.2
Nitrosotalea bavarica SbT1
Nitrosotalea okcheonensis CS
Candidatus Nitrosotalea devanaterra Nd1
Nitrosotalea sinensis Nd2
Thaumarchaeota archaeon 13_1_40CM_3_38_6
Thaumarchaeota archaeon RRmetagenome_bin19
Nitrososphaera viennensis EN76
Candidatus Nitrososphaera evergladensis SR1
Candidatus Nitrosocaldus islandicus
Candidatus Nitrosocaldus cavascurensis SCU2
Candidatus Nitrosopelagicus brevis CN25
Thermobaculum terrenum ATCC BAA-798
Streptomyces bingchenggensis BCW-1
Marine Group I thaumarchaeote SCGC AAA007-O23
Gamma AOA mg3 - D17
Candidatus Nitrosotenuis aquarius
Candidatus Nitrosotenuis cloacae SAT1
Roseiflexus castenholzii DSM 13941
Nitrosotenuis uzonensis N4
Nitrosotenuis chungbukensis MY2
Archaeon CSP1
Marine Group I thaumarchaeote SCGC RSA3
Nitrosoarchaeum koreensis MY1
Nitrosoarchaeum limnia SFB1
Cenarchaeum symbiosum A
Candidatus Nitrosomarinus catalina SPOT01
Thaumarchaeota archaeon Baikal-G2
Candidatus Nitrosopumilus adriaticus NF5
Candidatus Nitrosopumilus sediminis AR2
Nitrosopumilus sp. NM25
Nitrosopumilus sp SJ
Candidatus Nitrosopumilus koreensis AR1
Marine Group I thaumarchaeote SCGC RSA3
Nitrosopumilus maritimus SCM1
Candidatus Nitrosopumilus piranensis D3C
Alpha AOA ma8 - T3L19
Thaumarchaeota archaeon J079
Granulicella mallensis MP5ACTX8
Granulicella mallensis MP5ACTX8
Granulicella mallensis MP5ACTX8
Streptosporangium roseum DSM 43021
Streptosporangium roseum DSM 43021
Kibdelosporangium phytohabitans
Archangium gephyra
Candidatus Nitrosocosmicus exaquare G61
Gemmatimonas phototrophica
Pseudomonas citronellolis
Bradyrhizobium japonicum USDA 6
Candidatus Nitrosocosmicus franklandus C13
Candidatus Nitrosocosmicus exaquare G61
Methanocella conradii HZ254
Methanocella arvoryzae MRE50
Thaumarchaeota MOCA
Thaumarchaeota archaeon strain BS4 (Beowulf)
Candidatus Nitrososphaera gargensis Ga9.2
Candidatus Nitrososphaera evergladensis SR1
Nitrosotalea bavarica SbT1
Nitrosotalea okcheonensis CS
Thaumarchaeota archaeon 13_1_40CM_3_38_6
Thaumarchaeota archaeon 13_1_40CM_38_12
Candidatus Nitrosocosmicus franklandus C13
Candidatus Nitrososphaera gargensis Ga9.2
Thaumarchaeota archaeon RRmetagenome_bin19
Thaumarchaeota archaeon RRmetagenome_bin19
Nitrososphaera viennensis EN76
Candidatus Nitrososphaera evergladensis SR1
Nitrosotalea bavarica SbT1
Nitrosotalea sinensis Nd2
Nitrosotalea okcheonensis CS
Candidatus Nitrosotalea devanaterra Nd1
Thaumarchaeota archaeon 13_1_40CM_3_38_6
Thaumarchaeota archaeon 13_1_40CM_38_12
Candidatus Nitrosopumilus adriaticus NF5
Nitrosoarchaeum koreensis MY1
Marine Group I thaumarchaeote SCGC RSA3
Marine Group I thaumarchaeote SCGC RSA3
Nitrosopumilus maritimus SCM1
Cenarchaeum symbiosum A
Candidatus Nitrosotenuis aquarius
Candidatus Nitrosotenuis cloacae SAT1
Nitrosotenuis uzonensis N4
Nitrosoarchaeum koreensis MY1
Nitrosoarchaeum limnia SFB1
Nitrosopumilus sp. NM25
Candidatus Nitrosopumilus adriaticus NF5
Candidatus Nitrosopumilus sediminis AR2
Marine Group I thaumarchaeote SCGC RSA3
Nitrosopumilus maritimus SCM1
Nitrosopumilus sp SJ
Candidatus Nitrosopumilus koreensis AR1
Candidatus Nitrosopumilus piranensis D3C
Nitrosocosmicus arcticus Kfb
Candidatus Nitrosocosmicus franklandus C13
Candidatus Nitrosocosmicus exaquare G61
Candidatus Nitrosocosmicus exaquare G61
Candidatus Nitrosocosmicus oleophilus MY3
Candidatus Nitrosocosmicus oleophilus MY3
Candidatus Nitrosocaldus islandicus
Candidatus Nitrosocaldus cavascurensis SCU2
Thaumarchaeota archaeon J079
Thaumarchaeota MOCA
Thaumarchaeota archaeon SAT139
Thaumarchaeota SP3992
Thaumarchaeota archaeon UBA201
Thaumarchaeota archaeon UBA223
Sphaerobacter thermophilus DSM 20745
Sphaerobacter thermophilus DSM 20745
Thermomicrobium roseum DSM 5159
Bradyrhizobium japonicum USDA 6
Candidatus Nitrosocosmicus oleophilus MY3
Acidithiobacillus ferrooxidans ATCC 23270
Candidatus Nitrosocosmicus franklandus C13
Nitrosocosmicus arcticus Kfb
Candidatus Nitrosocosmicus exaquare G61
Nitrososphaera viennensis EN76
Candidatus Nitrososphaera evergladensis SR1
76G-FAM007520
76G-FAM011161
76G-FAM011132
76G-FAM018158
76G-FAM017461
76G-FAM028373
76G-FAM010275
76G-FAM011584
76G-FAM002773
76G-FAM008291
76G-FAM010540
76G-FAM010617
76G-FAM010303
Catenulisporales
Deinococci
Bacilli
Halobacteria
Stenosarchaea group
Nitrososphaerales
Nitrosotaleales
Nitrosocaldales
Nitrosopumilales
Streptomycetales
Chloroflexia
Acidobacteriales
Streptosporangiales
Pseudonocardiales
Deltaproteobacteria
Gemmatimonadales
Gammaproteobacteria
Alphaproteobacteria
Methanomicrobia
Allothaumarchaeota
Sphaerobacteridae
Thermomicrobiales
Acidithiobacillia
Actinobacteria
Deinococcus-Thermus
Firmicutes
Euryarchaeota
Thaumarchaeota
Thermobaculum
Chloroflexi
Acidobacteria
Proteobacteria
Gemmatimonadetes
Bacteria
Archaea

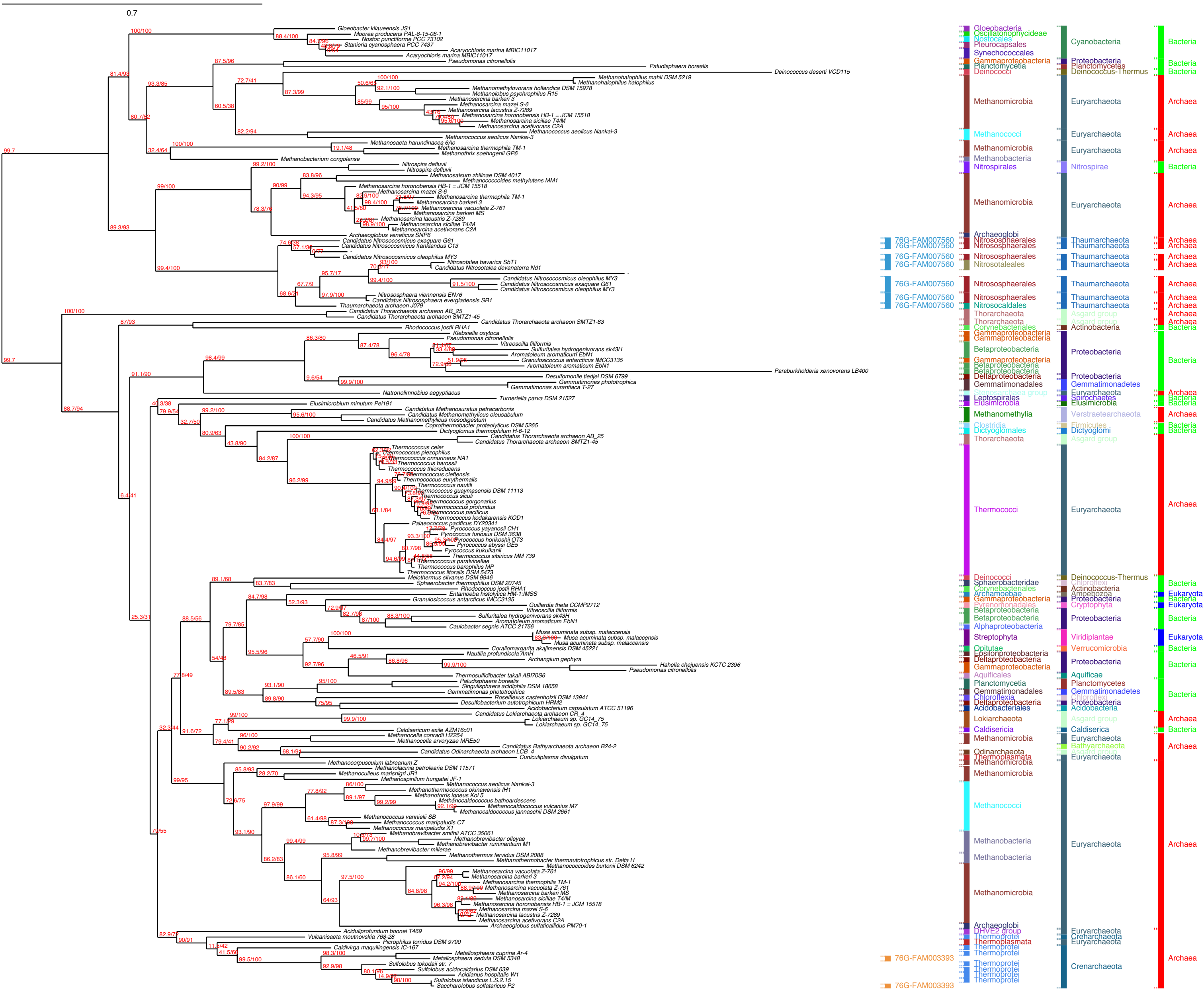

0.7
Gloeobacter kilaueensis JS1
Moorea producens PAL-8-15-08-1
Nostoc punctiforme PCC 73102
Stanieria cyanosphaera PCC 7437
Acaryochloris marina MBIC11017
Acaryochloris marina MBIC11017
Pseudomonas citronellolis
Paludisphaera borealis
Deinococcus deserti VCD115
Methanohalophilus mahii DSM 5219
Methanohalophilus halophilus
Methanomethylovorans hollandica DSM 15978
Methanolobus psychrophilus R15
Methanosarcina barkeri 3
Methanosarcina mazei S-6
Methanosarcina lacustris Z-7289
Methanosarcina horonobensis HB-1 = JCM 15518
Methanosarcina siciliae T4/M
Methanosarcina acetivorans C2A
Methanococcus aeolicus Nankai-3
Methanococcus aeolicus Nankai-3
Methanosaeta harundinacea 6Ac
Methanosarcina thermophila TM-1
Methanothrix soehngenii GP6
Methanobacterium congolense
Nitrospira defluvii
Nitrospira defluvii
Methanosalsum zhilinae DSM 4017
Methanococcoides methylutens MM1
Methanosarcina horonobensis HB-1 = JCM 15518
Methanosarcina mazei S-6
Methanosarcina thermophila TM-1
Methanosarcina barkeri 3
Methanosarcina vacuolata Z-761
Methanosarcina barkeri MS
Methanosarcina lacustris Z-7289
Methanosarcina siciliae T4/M
Methanosarcina acetivorans C2A
Archaeoglobus veneficus SNP6
Candidatus Nitrosocosmicus exaquare G61
Candidatus Nitrosocosmicus franklandus C13
Candidatus Nitrosocosmicus oleophilus MY3
Nitrosotalea bavarica SbT1
Candidatus Nitrosotalea devanaterra Nd1
Candidatus Nitrosocosmicus oleophilus MY3
Candidatus Nitrosocosmicus exaquare G61
Candidatus Nitrosocosmicus oleophilus MY3
Nitrososphaera viennensis EN76
Candidatus Nitrososphaera evergladensis SR1
Thaumarchaeota archaeon J079
Candidatus Thorarchaeota archaeon AB_25
Candidatus Thorarchaeota archaeon SMTZ1-45
Candidatus Thorarchaeota archaeon SMTZ1-83
Rhodococcus jostii RHA1
Klebsiella oxytoca
Pseudomonas citronellolis
Vitreoscilla filiformis
Sulfuritalea hydrogenivorans sk43H
Aromatoleum aromaticum EbN1
Granulosicoccus antarcticus IMCC3135
Aromatoleum aromaticum EbN1
Paraburkholderia xenovorans LB400
Desulfomonile tiedjei DSM 6799
Gemmatimonas phototrophica
Gemmatimonas aurantiaca T-27
Natronolimnobius aegyptiacus
Turneriella parva DSM 21527
Elusimicrobium minutum Pei191
Candidatus Methanosuratus petracarbonis
Candidatus Methanomethylicus oleusabulum
Candidatus Methanomethylicus mesodigestum
Coprothermobacter proteolyticus DSM 5265
Dictyoglomus thermophilum H-6-12
Candidatus Thorarchaeota archaeon AB_25
Candidatus Thorarchaeota archaeon SMTZ1-45
Thermococcus celer
Thermococcus piezophilus
Thermococcus onnurineus NA1
Thermococcus barossii
Thermococcus thioreducens
Thermococcus cleftensis
Thermococcus eurythermalis
Thermococcus nautili
Thermococcus guaymasensis DSM 11113
Thermococcus siculi
Thermococcus gorgonarius
Thermococcus profundus
Thermococcus pacificus
Thermococcus kodakarensis KOD1
Palaeococcus pacificus DY20341
Pyrococcus yayanosii CH1
Pyrococcus furiosus DSM 3638
Pyrococcus horikoshii OT3
Pyrococcus abyssi GE5
Pyrococcus kukulkanii
Thermococcus sibiricus MM 739
Thermococcus paralvinellae
Thermococcus barophilus MP
Thermococcus litoralis DSM 5473
Meiothermus silvanus DSM 9946
Sphaerobacter thermophilus DSM 20745
Rhodococcus jostii RHA1
Entamoeba histolytica HM-1:IMSS
Granulosicoccus antarcticus IMCC3135
Guillardia theta CCMP2712
Vitreoscilla filiformis
Sulfuritalea hydrogenivorans sk43H
Aromatoleum aromaticum EbN1
Caulobacter segnis ATCC 21756
Musa acuminata subsp. malaccensis
Musa acuminata subsp. malaccensis
Musa acuminata subsp. malaccensis
Coraliomargarita akajimensis DSM 45221
Nautilia profundicola AmH
Archangium gephyra
Hahella chejuensis KCTC 2396
Pseudomonas citronellolis
Thermosulfidibacter takaii ABI70S6
Paludisphaera borealis
Singulisphaera acidiphila DSM 18658
Gemmatimonas phototrophica
Roseiflexus castenholzii DSM 13941
Desulfobacterium autotrophicum HRM2
Acidobacterium capsulatum ATCC 51196
Candidatus Lokiarchaeota archaeon CR_4
Lokiarchaeum sp. GC14_75
Lokiarchaeum sp. GC14_75
Caldisericum exile AZM16c01
Methanocella conradii HZ254
Methanocella arvoryzae MRE50
Candidatus Bathyarchaeota archaeon B24-2
Candidatus Odinarchaeota archaeon LCB_4
Cuniculiplasma divulgatum
Methanocorpusculum labreanum Z
Methanolacinia petrolearia DSM 11571
Methanoculleus marisnigri JR1
Methanospirillum hungatei JF-1
Methanococcus aeolicus Nankai-3
Methanothermococcus okinawensis IH1
Methanotorris igneus Kol 5
Methanocaldococcus bathoardescens
Methanocaldococcus vulcanius M7
Methanocaldococcus jannaschii DSM 2661
Methanococcus vannielii SB
Methanococcus maripaludis C7
Methanococcus maripaludis X1
Methanobrevibacter smithii ATCC 35061
Methanobrevibacter olleyae
Methanobrevibacter ruminantium M1
Methanobrevibacter millerae
Methanothermus fervidus DSM 2088
Methanothermobacter thermautotrophicus str. Delta H
Methanococcoides burtonii DSM 6242
Methanosarcina vacuolata Z-761
Methanosarcina barkeri 3
Methanosarcina thermophila TM-1
Methanosarcina vacuolata Z-761
Methanosarcina barkeri MS
Methanosarcina siciliae T4/M
Methanosarcina horonobensis HB-1 = JCM 15518
Methanosarcina mazei S-6
Methanosarcina lacustris Z-7289
Methanosarcina acetivorans C2A
Archaeoglobus sulfaticallidus PM70-1
Aciduliprofundum boonei T469
Vulcanisaeta moutnovskia 768-28
Picrophilus torridus DSM 9790
Caldivirga maquilingensis IC-167
Metallosphaera cuprina Ar-4
Metallosphaera sedula DSM 5348
Sulfolobus tokodaii str. 7
Sulfolobus acidocaldarius DSM 639
Acidianus hospitalis W1
Sulfolobus islandicus L.S.2.15
Saccharolobus solfataricus P2
76G-FAM007560
76G-FAM007560
76G-FAM007560
76G-FAM007560
76G-FAM007560
76G-FAM007560
76G-FAM007560
76G-FAM003393
76G-FAM003393
Gloeobacteria
Oscillatoriophycideae
Nostocales
Pleurocapsales
Synechococcales
Gammaproteobacteria
Planctomycetia
Deinococci
Methanomicrobia
Methanococci
Methanomicrobia
Methanobacteria
Nitrospirales
Methanomicrobia
Archaeoglobi
Nitrososphaerales
Nitrososphaerales
Nitrososphaerales
Nitrosotaleales
Nitrososphaerales
Nitrososphaerales
Nitrosocaldales
Thorarchaeota
Thorarchaeota
Corynebacteriales
Gammaproteobacteria
Gammaproteobacteria
Betaproteobacteria
Gammaproteobacteria
Betaproteobacteria
Betaproteobacteria
Deltaproteobacteria
Gemmatimonadales
Leptospirales
Elusimicrobia
Methanomethylia
Clostridia
Dictyoglomales
Thorarchaeota
Thermococci
Deinococci
Sphaerobacteridae
Corynebacteriales
Archamoebae
Gammaproteobacteria
Pyrenomonadales
Betaproteobacteria
Betaproteobacteria
Alphaproteobacteria
Streptophyta
Opitutae
Epsilonproteobacteria
Deltaproteobacteria
Gammaproteobacteria
Aquificales
Planctomycetia
Gemmatimonadales
Chloroflexia
Deltaproteobacteria
Acidobacteriales
Lokiarchaeota
Caldisericia
Methanomicrobia
Odinarchaeota
Thermoplasmata
Methanomicrobia
Methanomicrobia
Methanococci
Methanobacteria
Methanobacteria
Methanomicrobia
Archaeoglobi
DHVE2 group
Thermoprotei
Thermoplasmata
Thermoprotei
Thermoprotei
Thermoprotei
Thermoprotei
Thermoprotei
Thermoprotei
Cyanobacteria
Proteobacteria
Planctomycetes
Deinococcus-Thermus
Euryarchaeota
Euryarchaeota
Euryarchaeota
Nitrospirae
Euryarchaeota
Thaumarchaeota
Thaumarchaeota
Thaumarchaeota
Thaumarchaeota
Thaumarchaeota
Thaumarchaeota
Thaumarchaeota
Asgard group
Asgard group
Actinobacteria
Proteobacteria
Proteobacteria
Gemmatimonadetes
Euryarchaeota
Spirochaetes
Elusimicrobia
Verstraetearchaeota
Firmicutes
Dictyoglomi
Asgard group
Euryarchaeota
Deinococcus-Thermus
Chloroflexi
Actinobacteria
Amoebozoa
Proteobacteria
Cryptophyta
Proteobacteria
Viridiplantae
Verrucomicrobia
Proteobacteria
Aquificae
Planctomycetes
Gemmatimonadetes
Chloroflexi
Proteobacteria
Acidobacteria
Asgard group
Caldiserica
Euryarchaeota
Bathyarchaeota
Asgard group
Euryarchaeota
Euryarchaeota
Euryarchaeota
Crenarchaeota
Euryarchaeota
Crenarchaeota
Bacteria
Bacteria
Bacteria
Archaea
Archaea
Archaea
Bacteria
Archaea
Archaea
Archaea
Archaea
Archaea
Archaea
Archaea
Archaea
Archaea
Archaea
Bacteria
Bacteria
Archaea
Bacteria
Bacteria
Archaea
Bacteria
Bacteria
Archaea
Bacteria
Eukaryota
Bacteria
Eukaryota
Bacteria
Eukaryota
Bacteria
Bacteria
Bacteria
Archaea
Bacteria
Archaea
Archaea
Archaea

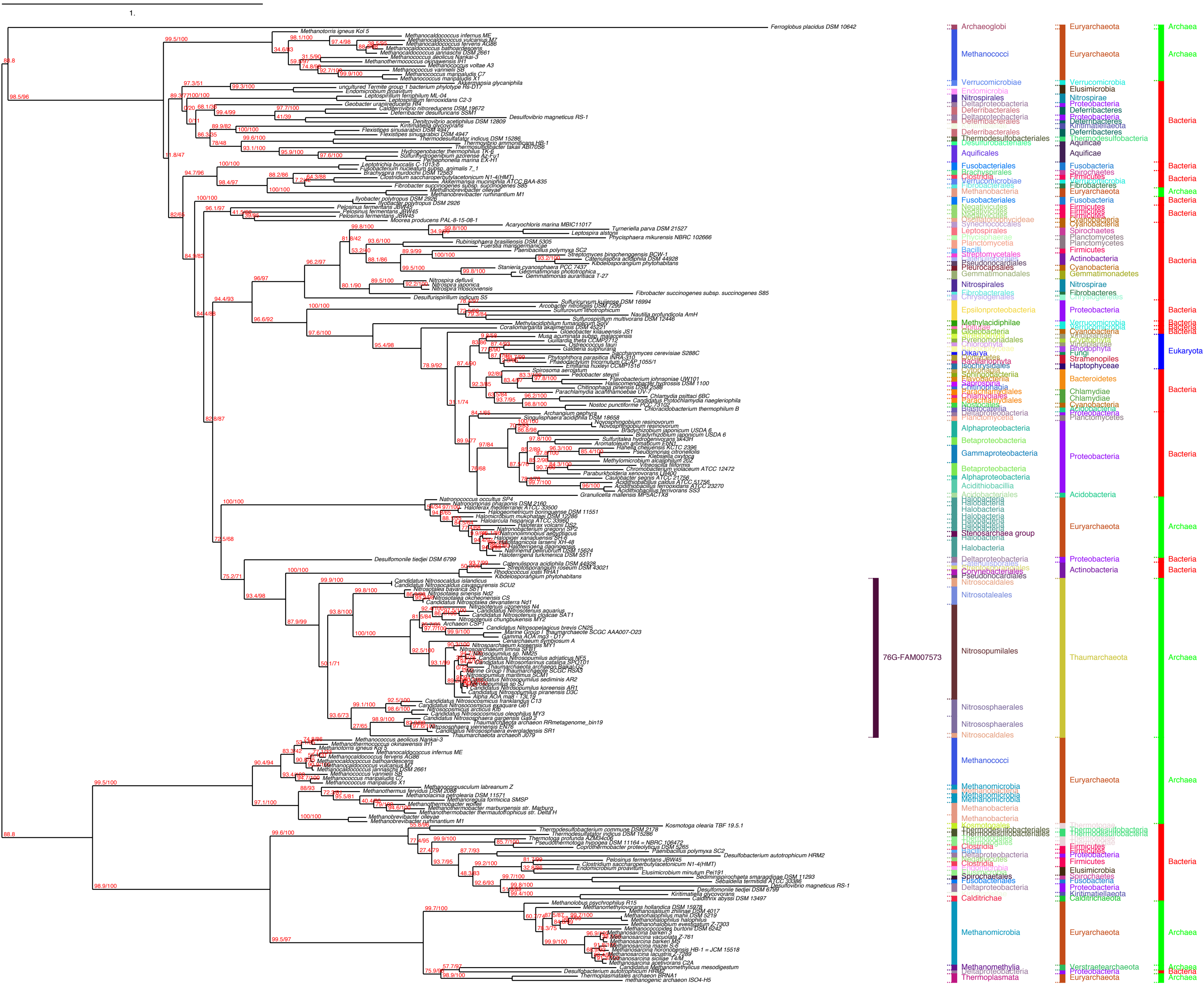
1.
76G-FAM007573
Archaeoglobi
Methanococci
Verrucomicrobiae
Nitrospirales
Deferribacterales
Thermodesulfobacteriales
Aquificales
Fusobacteriales
Methanobacteria
Leptospirales
Planctomycetia
Pseudonocardiales
Gemmatimonadales
Epsilonproteobacteria
Methylacidiphilae
Alphaproteobacteria
Betaproteobacteria
Gammaproteobacteria
Aciditihiobacillia
Halobacteria
Stenosarchaea group
Deltaproteobacteria
Corynebacteriales
Nitrosocaldales
Nitrosotaleales
Nitrosopumilales
Nitrososphaerales
Methanomicrobia
Clostridia
Spirochaetales
Calditrichae
Methanomethylia
Thermoplasmata
Euryarchaeota
Elusimicrobia
Nitrospirae
Proteobacteria
Deferribacteres
Aquificae
Fusobacteria
Fibrobacteres
Firmicutes
Cyanobacteria
Spirochaetes
Planctomycetes
Actinobacteria
Gemmatimonadetes
Fungi
Stramenopiles
Haptophyceae
Bacteroidetes
Chlamydiae
Acidobacteria
Thaumarchaeota
Verstraetearchaeota
Archaea
Bacteria
Eukaryota

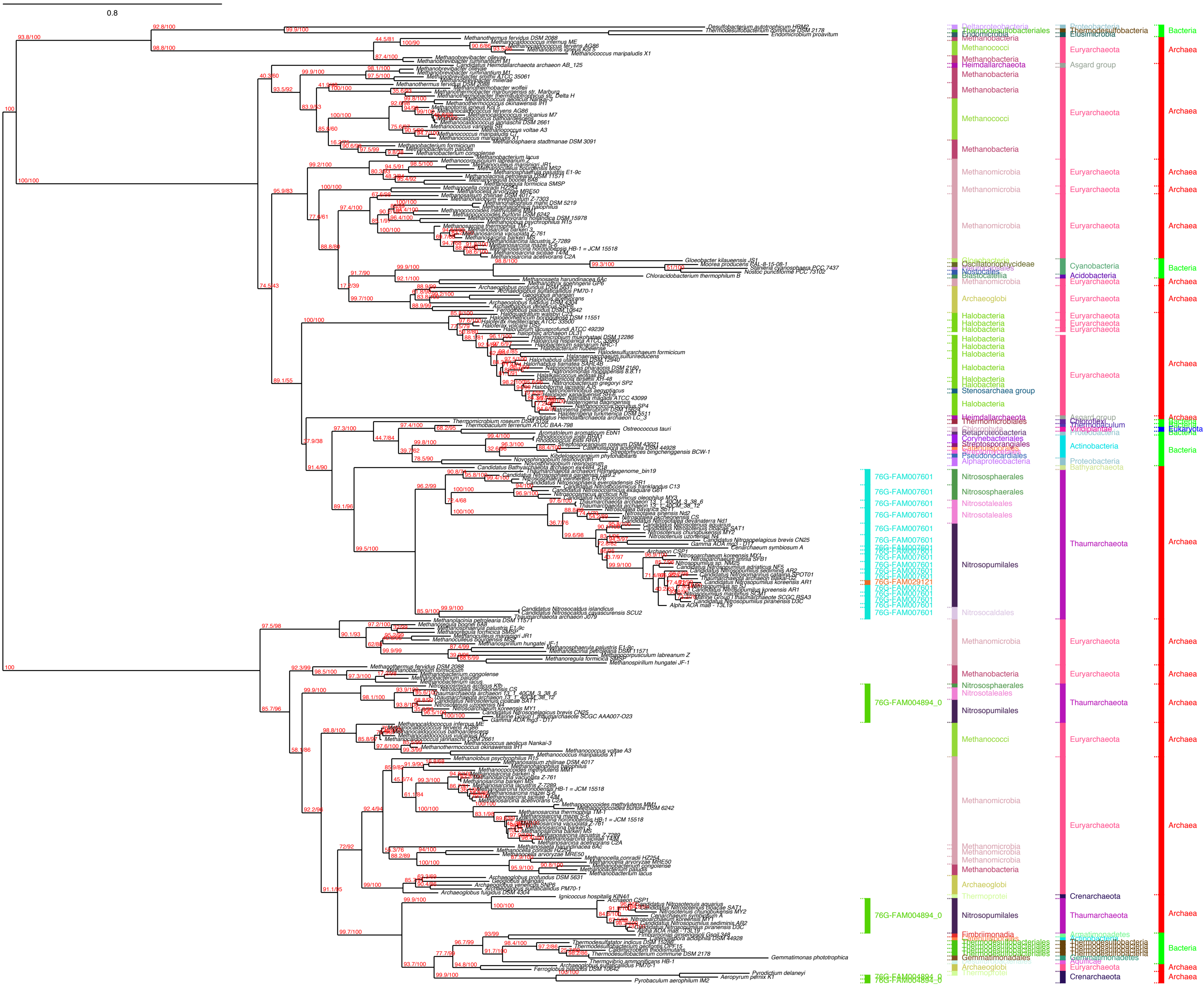
0.8
76G-FAM007601
76G-FAM029121
76G-FAM004894_0
Methanococci
Methanobacteria
Heimdallarchaeota
Methanomicrobia
Archaeoglobi
Halobacteria
Stenosarchaea group
Nitrososphaerales
Nitrosotaleales
Nitrosopumilales
Nitrosocaldales
Thermoprotei
Fimbriimonadia
Gemmatimonadales
Oscillatoriophycideae
Nostocales
Thermomicrobiales
Betaproteobacteria
Corynebacteriales
Streptosporangiales
Pseudonocardiales
Alphaproteobacteria
Thermodesulfobacteriales
Endomicrobia
Euryarchaeota
Asgard group
Cyanobacteria
Acidobacteria
Actinobacteria
Proteobacteria
Bathyarchaeota
Thaumarchaeota
Crenarchaeota
Thermodesulfobacteria
Elusimicrobia
Chloroflexi
Viridiplantae
Bacteria
Archaea
Eukaryota

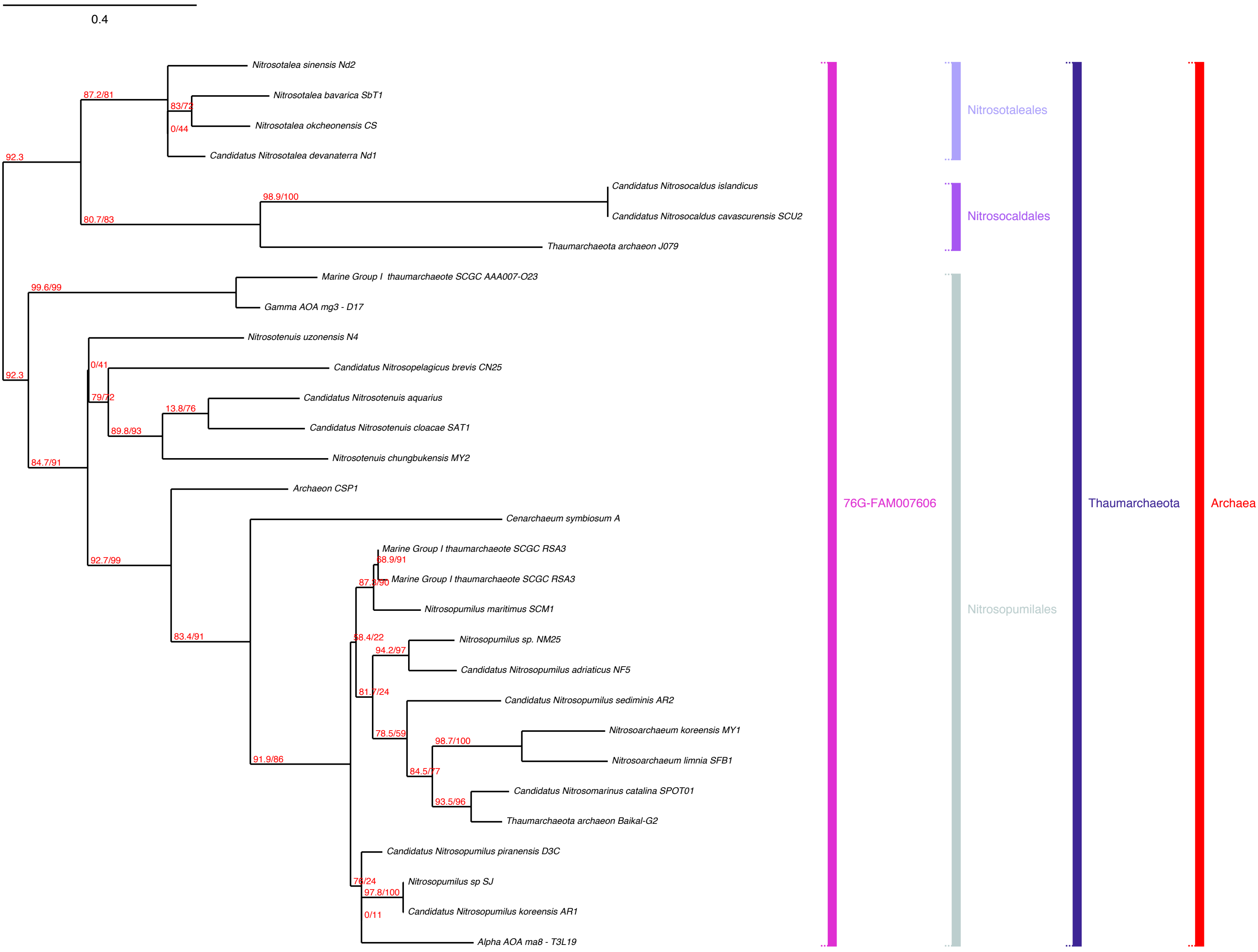

0.4
Nitrosotalea sinensis Nd2
Nitrosotalea bavarica SbT1
Nitrosotalea okcheonensis CS
Candidatus Nitrosotalea devanaterra Nd1
Candidatus Nitrosocaldus islandicus
Candidatus Nitrosocaldus cavascurensis SCU2
Thaumarchaeota archaeon J079
Marine Group I thaumarchaeote SCGC AAA007-O23
Gamma AOA mg3 - D17
Nitrosotenuis uzonensis N4
Candidatus Nitrosopelagicus brevis CN25
Candidatus Nitrosotenuis aquarius
Candidatus Nitrosotenuis cloacae SAT1
Nitrosotenuis chungbukensis MY2
Archaeon CSP1
Cenarchaeum symbiosum A
Marine Group I thaumarchaeote SCGC RSA3
Marine Group I thaumarchaeote SCGC RSA3
Nitrosopumilus maritimus SCM1
Nitrosopumilus sp. NM25
Candidatus Nitrosopumilus adriaticus NF5
Candidatus Nitrosopumilus sediminis AR2
Nitrosoarchaeum koreensis MY1
Nitrosoarchaeum limnia SFB1
Candidatus Nitrosomarinus catalina SPOT01
Thaumarchaeota archaeon Baikal-G2
Candidatus Nitrosopumilus piranensis D3C
Nitrosopumilus sp SJ
Candidatus Nitrosopumilus koreensis AR1
Alpha AOA ma8 - T3L19
87.2/81
83/72
0/44
92.3
98.9/100
80.7/83
99.6/99
0/41
92.3
79/72
13.8/76
89.8/93
84.7/91
92.7/99
68.9/91
87.3/90
83.4/91
58.4/22
94.2/97
81.7/24
78.5/59
98.7/100
91.9/86
84.5/77
93.5/96
76/24
97.8/100
0/11
Nitrosotaleales
Nitrosocaldales
Nitrosopumilales
76G-FAM007606
Thaumarchaeota
Archaea

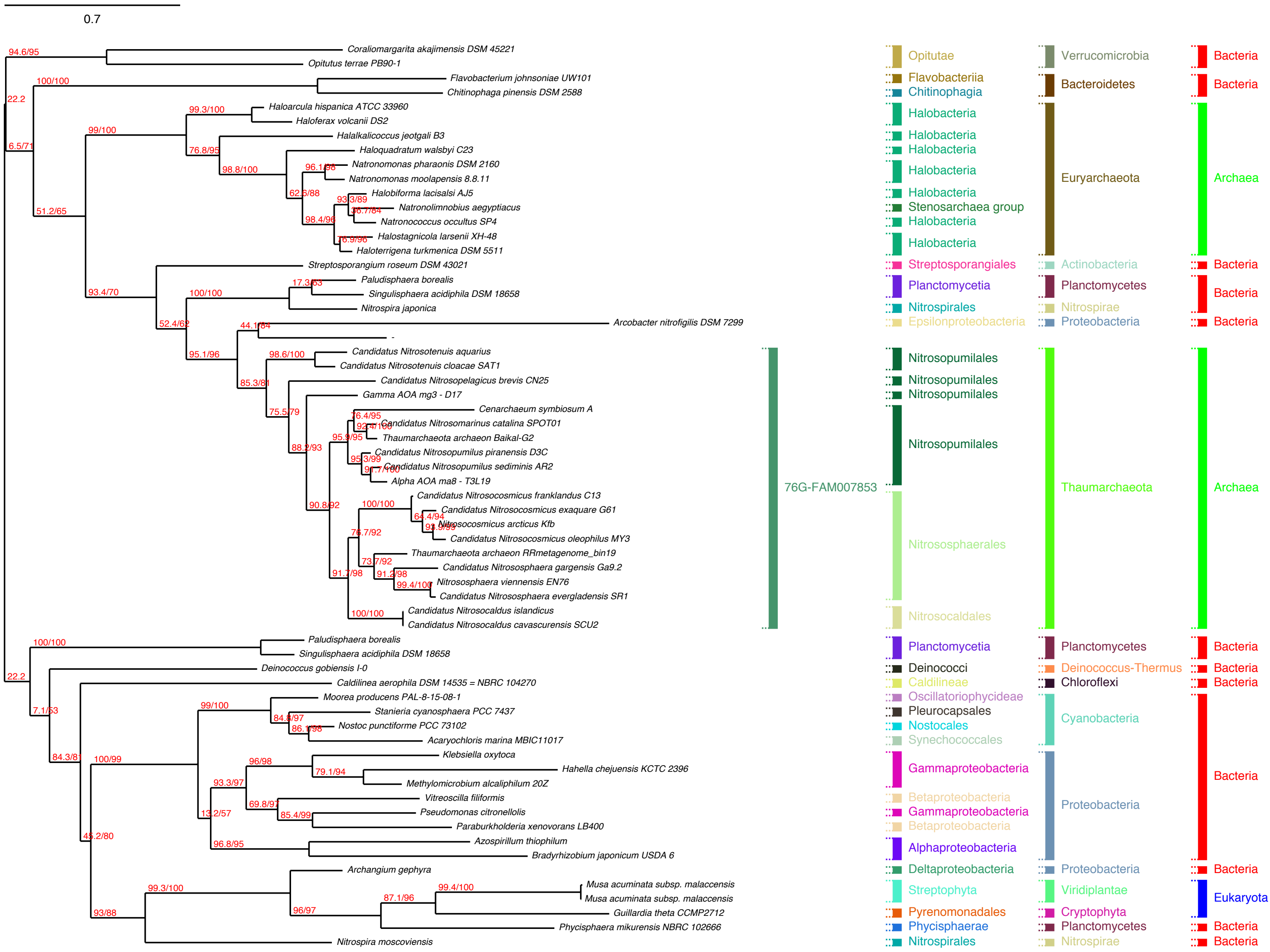
0.7
Coraliomargarita akajimensis DSM 45221
Opitutus terrae PB90-1
Flavobacterium johnsoniae UW101
Chitinophaga pinensis DSM 2588
Haloarcula hispanica ATCC 33960
Haloferax volcanii DS2
Halalkalicoccus jeotgali B3
Haloquadratum walsbyi C23
Natronomonas pharaonis DSM 2160
Natronomonas moolapensis 8.8.11
Halobiforma lacisalsi AJ5
Natronolimnobius aegyptiacus
Natronococcus occultus SP4
Halostagnicola larsenii XH-48
Haloterrigena turkmenica DSM 5511
Streptosporangium roseum DSM 43021
Paludisphaera borealis
Singulisphaera acidiphila DSM 18658
Nitrospira japonica
Arcobacter nitrofigilis DSM 7299
Candidatus Nitrosotenuis aquarius
Candidatus Nitrosotenuis cloacae SAT1
Candidatus Nitrosopelagicus brevis CN25
Gamma AOA mg3 - D17
Cenarchaeum symbiosum A
Candidatus Nitrosomarinus catalina SPOT01
Thaumarchaeota archaeon Baikal-G2
Candidatus Nitrosopumilus piranensis D3C
Candidatus Nitrosopumilus sediminis AR2
Alpha AOA ma8 - T3L19
Candidatus Nitrosocosmicus franklandus C13
Candidatus Nitrosocosmicus exaquare G61
Nitrosocosmicus arcticus Kfb
Candidatus Nitrosocosmicus oleophilus MY3
Thaumarchaeota archaeon RRmetagenome_bin19
Candidatus Nitrososphaera gargensis Ga9.2
Nitrososphaera viennensis EN76
Candidatus Nitrososphaera evergladensis SR1
Candidatus Nitrosocaldus islandicus
Candidatus Nitrosocaldus cavascurensis SCU2
Paludisphaera borealis
Singulisphaera acidiphila DSM 18658
Deinococcus gobiensis I-0
Caldilinea aerophila DSM 14535 = NBRC 104270
Moorea producens PAL-8-15-08-1
Stanieria cyanosphaera PCC 7437
Nostoc punctiforme PCC 73102
Acaryochloris marina MBIC11017
Klebsiella oxytoca
Hahella chejuensis KCTC 2396
Methylomicrobium alcaliphilum 20Z
Vitreoscilla filiformis
Pseudomonas citronellolis
Paraburkholderia xenovorans LB400
Azospirillum thiophilum
Bradyrhizobium japonicum USDA 6
Archangium gephyra
Musa acuminata subsp. malaccensis
Musa acuminata subsp. malaccensis
Guillardia theta CCMP2712
Phycisphaera mikurensis NBRC 102666
Nitrospira moscoviensis
76G-FAM007853
Opitutae
Flavobacteriia
Chitinophagia
Halobacteria
Stenosarchaea group
Streptosporangiales
Planctomycetia
Nitrospirales
Epsilonproteobacteria
Nitrosopumilales
Nitrososphaerales
Nitrosocaldales
Deinococci
Caldilineae
Oscillatoriophycideae
Pleurocapsales
Nostocales
Synechococcales
Gammaproteobacteria
Betaproteobacteria
Alphaproteobacteria
Deltaproteobacteria
Streptophyta
Pyrenomonadales
Phycisphaerae
Verrucomicrobia
Bacteroidetes
Euryarchaeota
Actinobacteria
Planctomycetes
Nitrospirae
Proteobacteria
Thaumarchaeota
Deinococcus-Thermus
Chloroflexi
Cyanobacteria
Viridiplantae
Cryptophyta
Bacteria
Archaea
Eukaryota

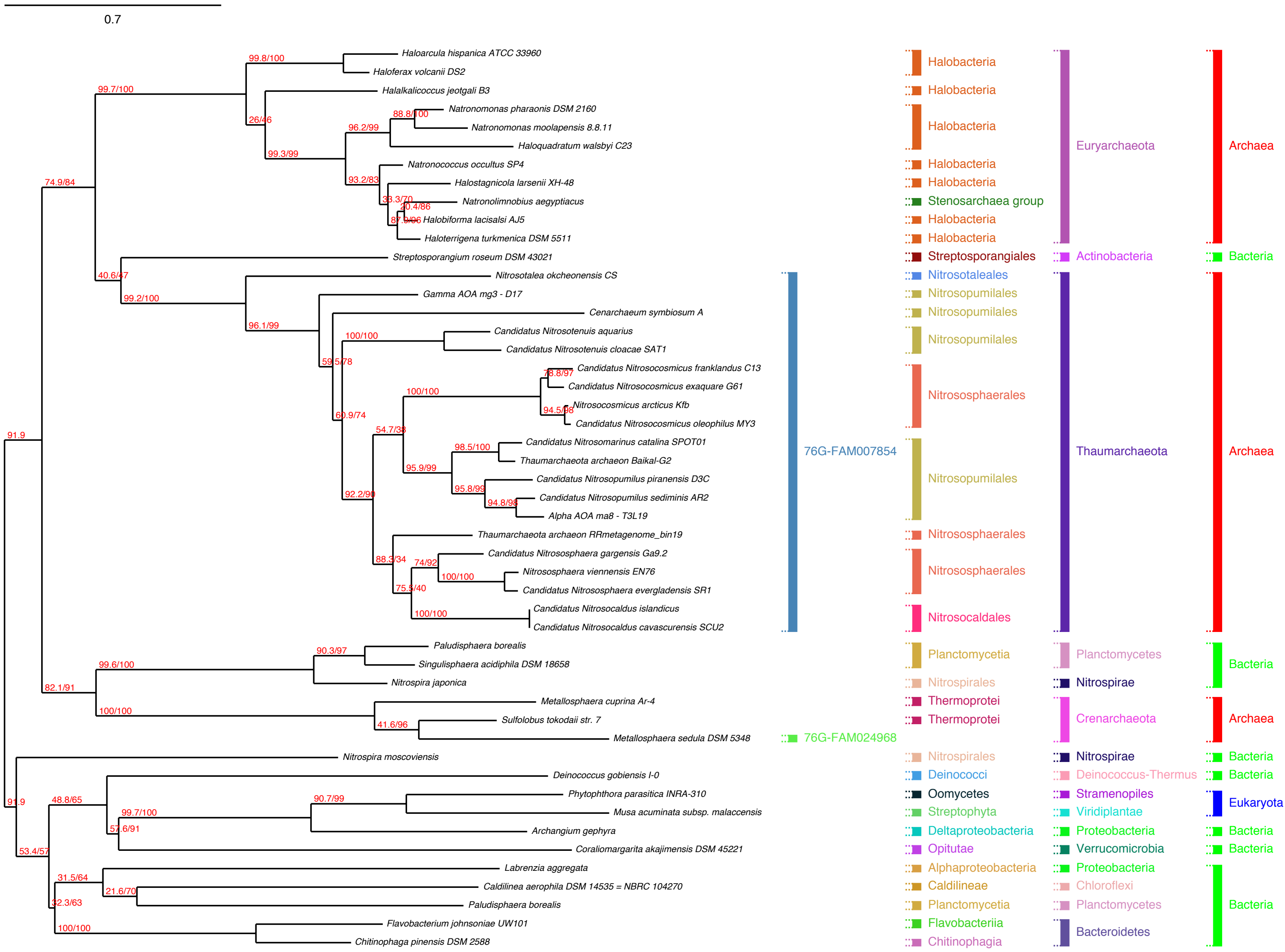
0.7
Haloarcula hispanica ATCC 33960
Haloferax volcanii DS2
Halalkalicoccus jeotgali B3
Natronomonas pharaonis DSM 2160
Natronomonas moolapensis 8.8.11
Haloquadratum walsbyi C23
Natronococcus occultus SP4
Halostagnicola larsenii XH-48
Natronolimnobius aegyptiacus
Halobiforma lacisalsi AJ5
Haloterrigena turkmenica DSM 5511
Streptosporangium roseum DSM 43021
Nitrosotalea okcheonensis CS
Gamma AOA mg3 - D17
Cenarchaeum symbiosum A
Candidatus Nitrosotenuis aquarius
Candidatus Nitrosotenuis cloacae SAT1
Candidatus Nitrosocosmicus franklandus C13
Candidatus Nitrosocosmicus exaquare G61
Nitrosocosmicus arcticus Kfb
Candidatus Nitrosocosmicus oleophilus MY3
Candidatus Nitrosomarinus catalina SPOT01
Thaumarchaeota archaeon Baikal-G2
Candidatus Nitrosopumilus piranensis D3C
Candidatus Nitrosopumilus sediminis AR2
Alpha AOA ma8 - T3L19
Thaumarchaeota archaeon RRmetagenome_bin19
Candidatus Nitrososphaera gargensis Ga9.2
Nitrososphaera viennensis EN76
Candidatus Nitrososphaera evergladensis SR1
Candidatus Nitrosocaldus islandicus
Candidatus Nitrosocaldus cavascurensis SCU2
Paludisphaera borealis
Singulisphaera acidiphila DSM 18658
Nitrospira japonica
Metallosphaera cuprina Ar-4
Sulfolobus tokodaii str. 7
Metallosphaera sedula DSM 5348
Nitrospira moscoviensis
Deinococcus gobiensis I-0
Phytophthora parasitica INRA-310
Musa acuminata subsp. malaccensis
Archangium gephyra
Coraliomargarita akajimensis DSM 45221
Labrenzia aggregata
Caldilinea aerophila DSM 14535 = NBRC 104270
Paludisphaera borealis
Flavobacterium johnsoniae UW101
Chitinophaga pinensis DSM 2588
99.8/100
99.7/100
26/46
88.8/100
96.2/99
99.3/99
93.2/83
33.3/70
20.4/86
87.9/96
74.9/84
40.6/47
99.2/100
96.1/99
100/100
59.5/78
78.8/97
100/100
94.5/98
60.9/74
54.7/38
98.5/100
95.9/99
95.8/99
94.8/98
92.2/90
88.3/34
74/92
100/100
75.5/40
100/100
91.9
90.3/97
99.6/100
82.1/91
100/100
41.6/96
91.9
48.8/65
90.7/99
99.7/100
57.6/91
53.4/57
31.5/64
21.6/70
32.3/63
100/100
76G-FAM007854
76G-FAM024968
Halobacteria
Halobacteria
Halobacteria
Halobacteria
Halobacteria
Stenosarchaea group
Halobacteria
Halobacteria
Streptosporangiales
Nitrosotaleales
Nitrosopumilales
Nitrosopumilales
Nitrosopumilales
Nitrososphaerales
Nitrosopumilales
Nitrososphaerales
Nitrososphaerales
Nitrosocaldales
Planctomycetia
Nitrospirales
Thermoprotei
Thermoprotei
Nitrospirales
Deinococci
Oomycetes
Streptophyta
Deltaproteobacteria
Opitutae
Alphaproteobacteria
Caldilineae
Planctomycetia
Flavobacteriia
Chitinophagia
Euryarchaeota
Actinobacteria
Thaumarchaeota
Planctomycetes
Nitrospirae
Crenarchaeota
Nitrospirae
Deinococcus-Thermus
Stramenopiles
Viridiplantae
Proteobacteria
Verrucomicrobia
Proteobacteria
Chloroflexi
Planctomycetes
Bacteroidetes
Archaea
Bacteria
Archaea
Bacteria
Archaea
Bacteria
Bacteria
Eukaryota
Bacteria
Bacteria
Bacteria

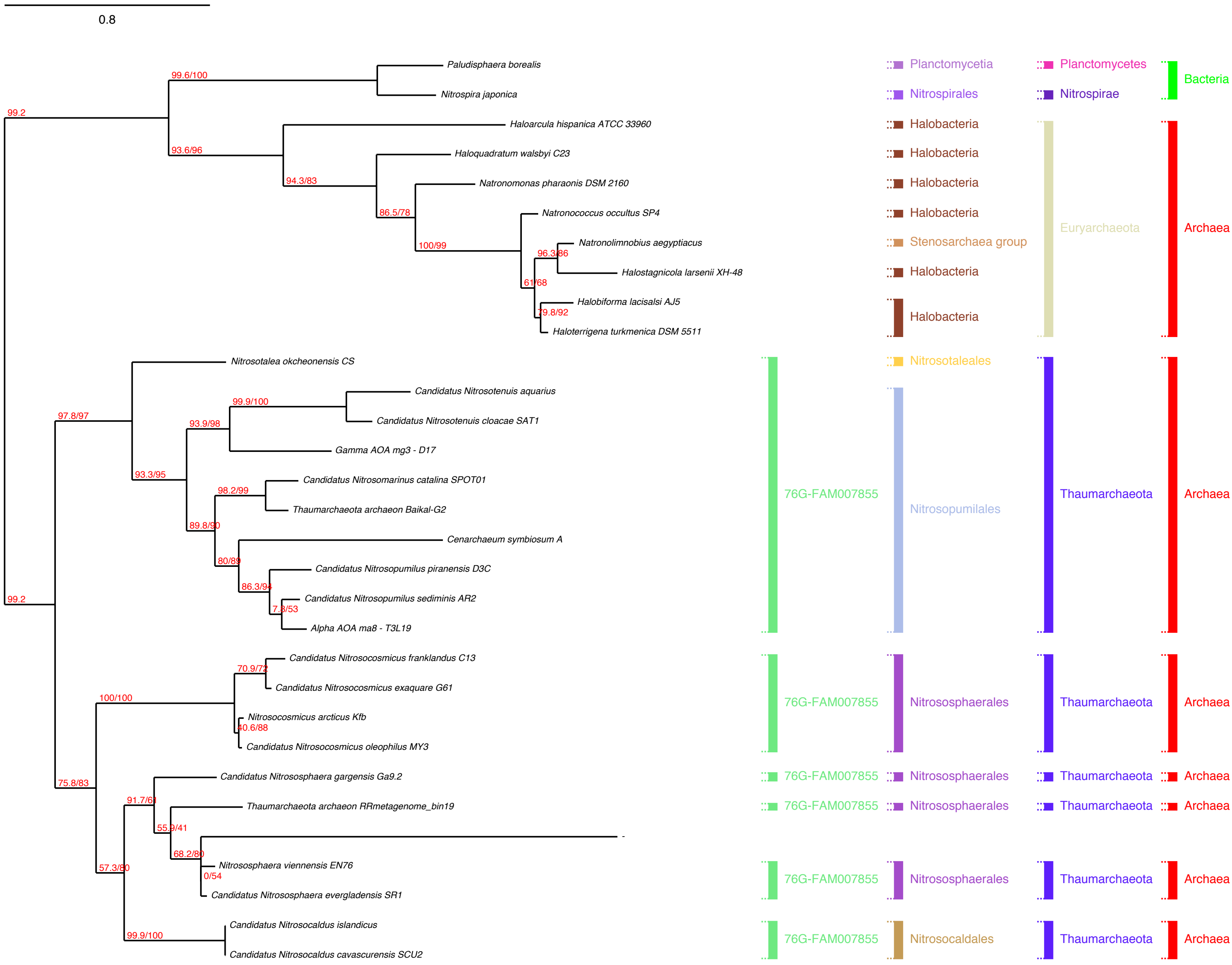

0.8
Paludisphaera borealis
Nitrospira japonica
Haloarcula hispanica ATCC 33960
Haloquadratum walsbyi C23
Natronomonas pharaonis DSM 2160
Natronococcus occultus SP4
Natronolimnobius aegyptiacus
Halostagnicola larsenii XH-48
Halobiforma lacisalsi AJ5
Haloterrigena turkmenica DSM 5511
Nitrosotalea okcheonensis CS
Candidatus Nitrosotenuis aquarius
Candidatus Nitrosotenuis cloacae SAT1
Gamma AOA mg3 - D17
Candidatus Nitrosomarinus catalina SPOT01
Thaumarchaeota archaeon Baikal-G2
Cenarchaeum symbiosum A
Candidatus Nitrosopumilus piranensis D3C
Candidatus Nitrosopumilus sediminis AR2
Alpha AOA ma8 - T3L19
Candidatus Nitrosocosmicus franklandus C13
Candidatus Nitrosocosmicus exaquare G61
Nitrosocosmicus arcticus Kfb
Candidatus Nitrosocosmicus oleophilus MY3
Candidatus Nitrososphaera gargensis Ga9.2
Thaumarchaeota archaeon RRmetagenome_bin19
-
Nitrososphaera viennensis EN76
Candidatus Nitrososphaera evergladensis SR1
Candidatus Nitrosocaldus islandicus
Candidatus Nitrosocaldus cavascurensis SCU2
99.2
99.6/100
93.6/96
94.3/83
86.5/78
100/99
96.3/86
61/68
79.8/92
99.2
97.8/97
99.9/100
93.9/98
93.3/95
98.2/99
89.8/90
80/89
86.3/94
7.8/53
70.9/72
100/100
40.6/88
75.8/83
91.7/61
55.9/41
68.2/80
0/54
57.3/80
99.9/100
Planctomycetia
Nitrospirales
Halobacteria
Halobacteria
Halobacteria
Halobacteria
Stenosarchaea group
Halobacteria
Halobacteria
Nitrosotaleales
Nitrosopumilales
Nitrososphaerales
Nitrososphaerales
Nitrososphaerales
Nitrososphaerales
Nitrosocaldales
Planctomycetes
Nitrospirae
Euryarchaeota
Thaumarchaeota
Thaumarchaeota
Thaumarchaeota
Thaumarchaeota
Thaumarchaeota
Thaumarchaeota
Bacteria
Archaea
Archaea
Archaea
Archaea
Archaea
Archaea
Archaea
76G-FAM007855
76G-FAM007855
76G-FAM007855
76G-FAM007855
76G-FAM007855
76G-FAM007855

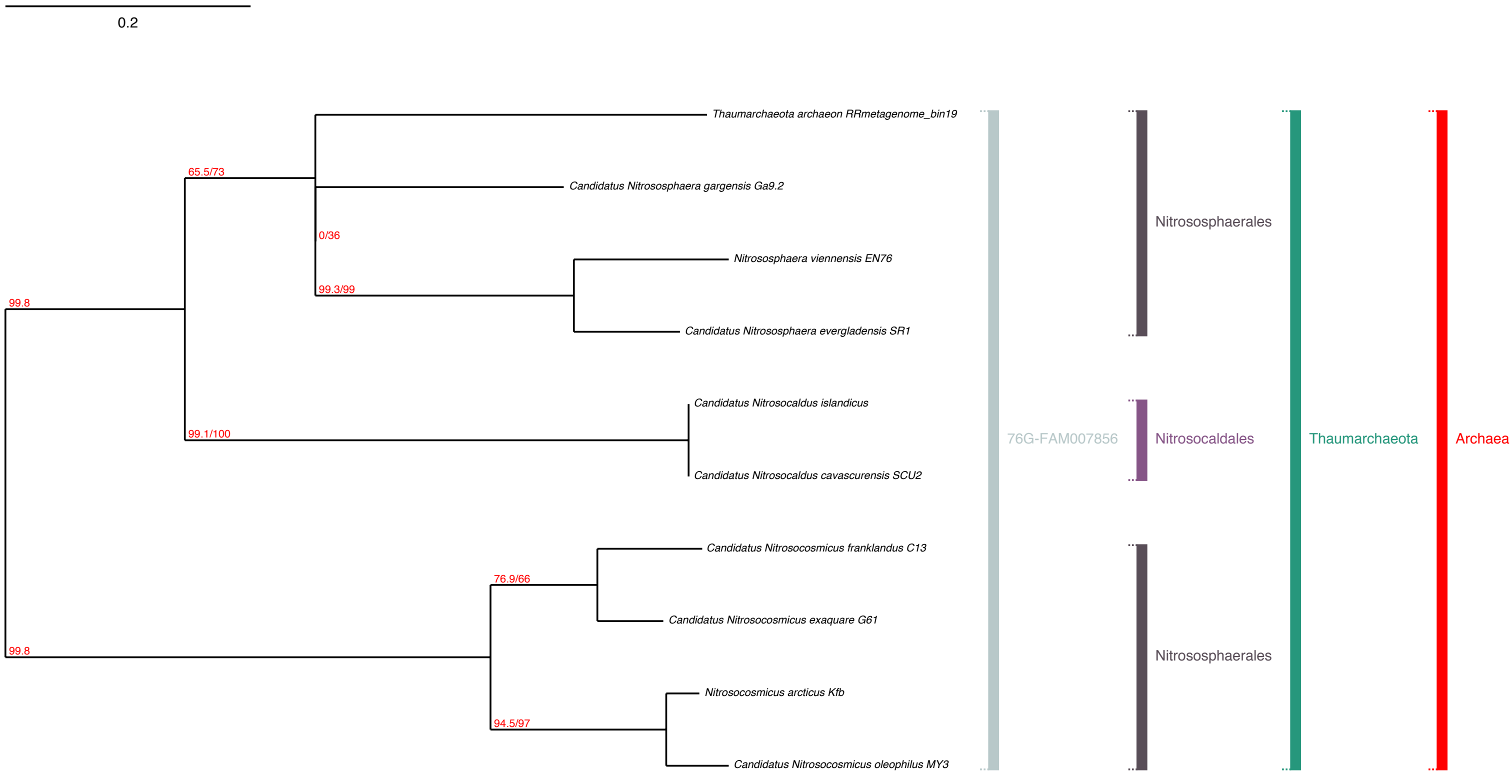

0.2
Thaumarchaeota archaeon RRmetagenome_bin19
Candidatus Nitrososphaera gargensis Ga9.2
Nitrososphaera viennensis EN76
Candidatus Nitrososphaera evergladensis SR1
Candidatus Nitrosocaldus islandicus
Candidatus Nitrosocaldus cavascurensis SCU2
Candidatus Nitrosocosmicus franklandus C13
Candidatus Nitrosocosmicus exaquare G61
Nitrosocosmicus arcticus Kfb
Candidatus Nitrosocosmicus oleophilus MY3
65.5/73
0/36
99.3/99
99.8
99.1/100
76.9/66
99.8
94.5/97
76G-FAM007856
Nitrososphaerales
Nitrosocaldales
Nitrososphaerales
Thaumarchaeota
Archaea

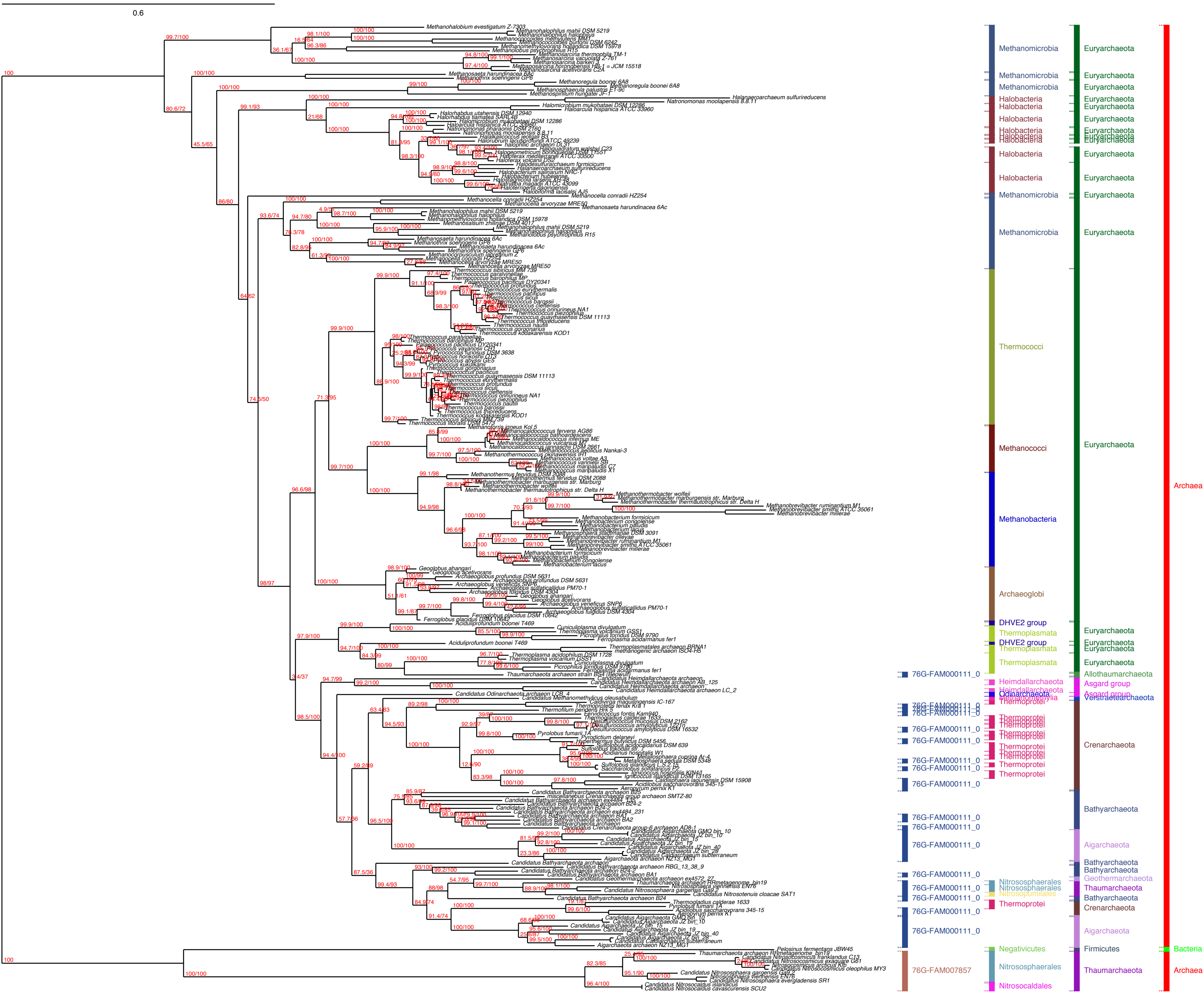
0.6
Methanomicrobia
Halobacteria
Thermococci
Methanococci
Methanobacteria
Archaeoglobi
DHVE2 group
Thermoplasmata
76G-FAM000111_0
Heimdallarchaeota
Odinarchaeota
Methanomethylia
Thermoprotei
Nitrososphaerales
Nitrosopumilales
Negativicutes
76G-FAM007857
Nitrosocaldales
Euryarchaeota
Allothaumarchaeota
Asgard group
Verstraetearchaeota
Crenarchaeota
Bathyarchaeota
Aigarchaeota
Geothermarchaeota
Thaumarchaeota
Firmicutes
Archaea
Bacteria

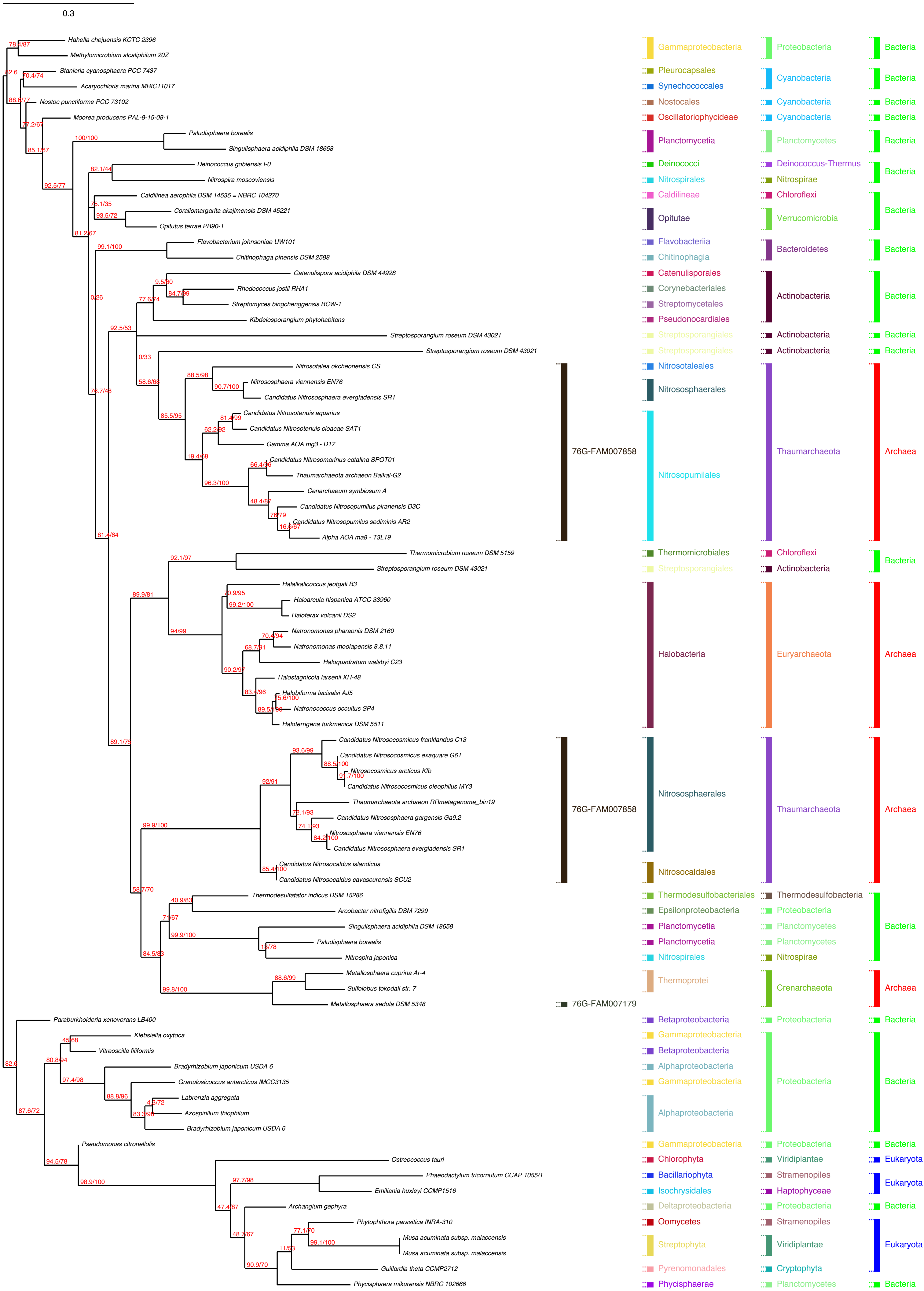

0.3
Hahella chejuensis KCTC 2396
Methylomicrobium alcaliphilum 20Z
Stanieria cyanosphaera PCC 7437
Acaryochloris marina MBIC11017
Nostoc punctiforme PCC 73102
Moorea producens PAL-8-15-08-1
Paludisphaera borealis
Singulisphaera acidiphila DSM 18658
Deinococcus gobiensis I-0
Nitrospira moscoviensis
Caldilinea aerophila DSM 14535 = NBRC 104270
Coraliomargarita akajimensis DSM 45221
Opitutus terrae PB90-1
Flavobacterium johnsoniae UW101
Chitinophaga pinensis DSM 2588
Catenulispora acidiphila DSM 44928
Rhodococcus jostii RHA1
Streptomyces bingchenggensis BCW-1
Kibdelosporangium phytohabitans
Streptosporangium roseum DSM 43021
Streptosporangium roseum DSM 43021
Nitrosotalea okcheonensis CS
Nitrososphaera viennensis EN76
Candidatus Nitrososphaera evergladensis SR1
Candidatus Nitrosotenuis aquarius
Candidatus Nitrosotenuis cloacae SAT1
Gamma AOA mg3 - D17
Candidatus Nitrosomarinus catalina SPOT01
Thaumarchaeota archaeon Baikal-G2
Cenarchaeum symbiosum A
Candidatus Nitrosopumilus piranensis D3C
Candidatus Nitrosopumilus sediminis AR2
Alpha AOA ma8 - T3L19
Thermomicrobium roseum DSM 5159
Streptosporangium roseum DSM 43021
Halalkalicoccus jeotgali B3
Haloarcula hispanica ATCC 33960
Haloferax volcanii DS2
Natronomonas pharaonis DSM 2160
Natronomonas moolapensis 8.8.11
Haloquadratum walsbyi C23
Halostagnicola larsenii XH-48
Halobiforma lacisalsi AJ5
Natronococcus occultus SP4
Haloterrigena turkmenica DSM 5511
Candidatus Nitrosocosmicus franklandus C13
Candidatus Nitrosocosmicus exaquare G61
Nitrosocosmicus arcticus Kfb
Candidatus Nitrosocosmicus oleophilus MY3
Thaumarchaeota archaeon RRmetagenome_bin19
Candidatus Nitrososphaera gargensis Ga9.2
Nitrososphaera viennensis EN76
Candidatus Nitrososphaera evergladensis SR1
Candidatus Nitrosocaldus islandicus
Candidatus Nitrosocaldus cavascurensis SCU2
Thermodesulfatator indicus DSM 15286
Arcobacter nitrofigilis DSM 7299
Singulisphaera acidiphila DSM 18658
Paludisphaera borealis
Nitrospira japonica
Metallosphaera cuprina Ar-4
Sulfolobus tokodaii str. 7
Metallosphaera sedula DSM 5348
Paraburkholderia xenovorans LB400
Klebsiella oxytoca
Vitreoscilla filiformis
Bradyrhizobium japonicum USDA 6
Granulosicoccus antarcticus IMCC3135
Labrenzia aggregata
Azospirillum thiophilum
Bradyrhizobium japonicum USDA 6
Pseudomonas citronellolis
Ostreococcus tauri
Phaeodactylum tricornutum CCAP 1055/1
Emiliania huxleyi CCMP1516
Archangium gephyra
Phytophthora parasitica INRA-310
Musa acuminata subsp. malaccensis
Musa acuminata subsp. malaccensis
Guillardia theta CCMP2712
Phycisphaera mikurensis NBRC 102666
76G-FAM007858
76G-FAM007858
76G-FAM007179
Gammaproteobacteria
Pleurocapsales
Synechococcales
Nostocales
Oscillatoriophycideae
Planctomycetia
Deinococci
Nitrospirales
Caldilineae
Opitutae
Flavobacteriia
Chitinophagia
Catenulisporales
Corynebacteriales
Streptomycetales
Pseudonocardiales
Streptosporangiales
Streptosporangiales
Nitrosotaleales
Nitrososphaerales
Nitrosopumilales
Thermomicrobiales
Streptosporangiales
Halobacteria
Nitrososphaerales
Nitrosocaldales
Thermodesulfobacteriales
Epsilonproteobacteria
Planctomycetia
Planctomycetia
Nitrospirales
Thermoprotei
Betaproteobacteria
Gammaproteobacteria
Betaproteobacteria
Alphaproteobacteria
Gammaproteobacteria
Alphaproteobacteria
Gammaproteobacteria
Chlorophyta
Bacillariophyta
Isochrysidales
Deltaproteobacteria
Oomycetes
Streptophyta
Pyrenomonadales
Phycisphaerae
Proteobacteria
Cyanobacteria
Cyanobacteria
Cyanobacteria
Planctomycetes
Deinococcus-Thermus
Nitrospirae
Chloroflexi
Verrucomicrobia
Bacteroidetes
Actinobacteria
Actinobacteria
Actinobacteria
Thaumarchaeota
Chloroflexi
Actinobacteria
Euryarchaeota
Thaumarchaeota
Thermodesulfobacteria
Proteobacteria
Planctomycetes
Planctomycetes
Nitrospirae
Crenarchaeota
Proteobacteria
Proteobacteria
Proteobacteria
Viridiplantae
Stramenopiles
Haptophyceae
Proteobacteria
Stramenopiles
Viridiplantae
Cryptophyta
Planctomycetes
Bacteria
Archaea
Eukaryota

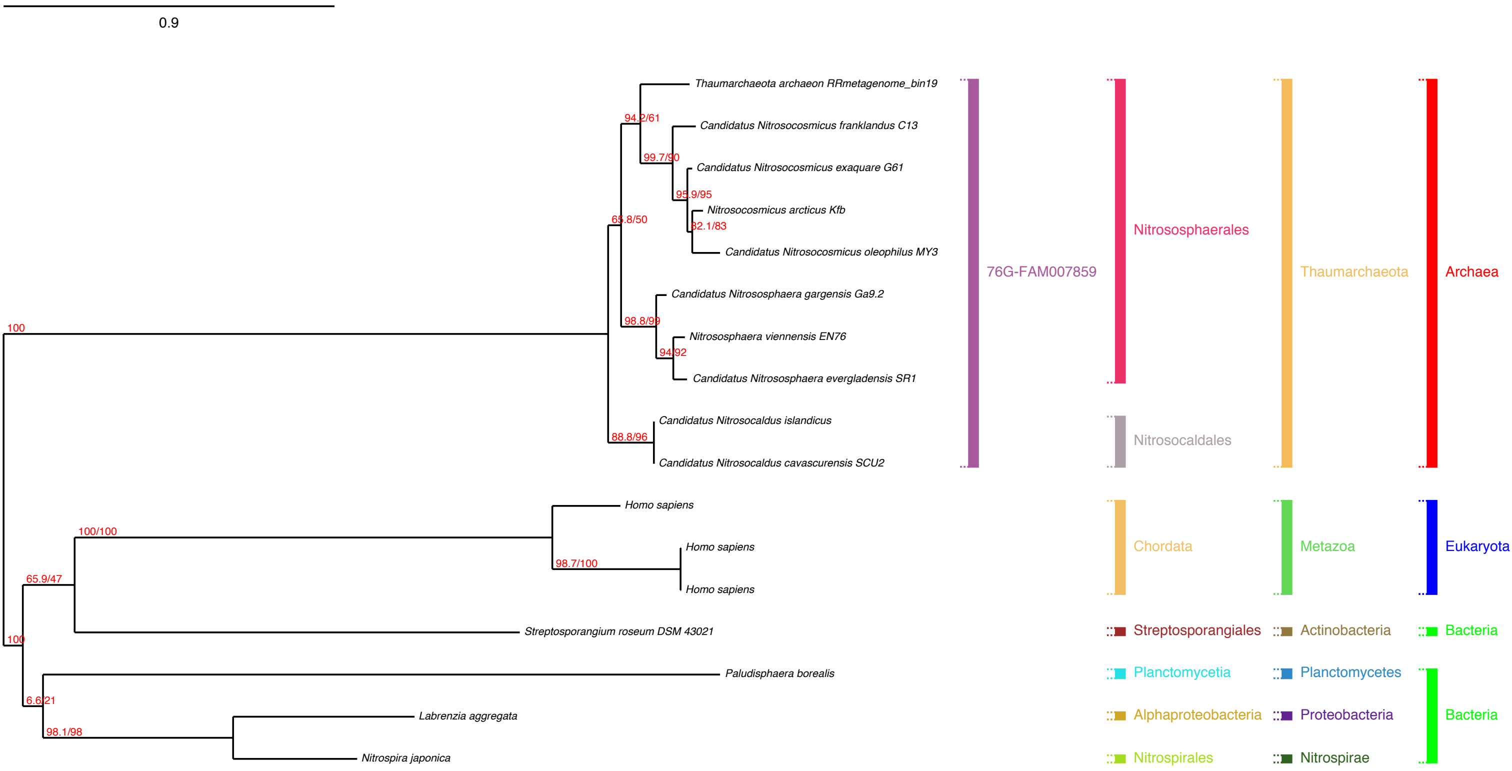
0.9
Thaumarchaeota archaeon RRmetagenome_bin19
Candidatus Nitrosocosmicus franklandus C13
Candidatus Nitrosocosmicus exaquare G61
Nitrosocosmicus arcticus Kfb
Candidatus Nitrosocosmicus oleophilus MY3
Candidatus Nitrososphaera gargensis Ga9.2
Nitrososphaera viennensis EN76
Candidatus Nitrososphaera evergladensis SR1
Candidatus Nitrosocaldus islandicus
Candidatus Nitrosocaldus cavascurensis SCU2
Homo sapiens
Homo sapiens
Homo sapiens
Streptosporangium roseum DSM 43021
Paludisphaera borealis
Labrenzia aggregata
Nitrospira japonica
100
94.2/61
99.7/90
95.9/95
82.1/83
65.8/50
98.8/99
94/92
88.8/96
100/100
98.7/100
65.9/47
100
6.6/21
98.1/98
76G-FAM007859
Nitrososphaerales
Nitrosocaldales
Chordata
Streptosporangiales
Planctomycetia
Alphaproteobacteria
Nitrospirales
Thaumarchaeota
Metazoa
Actinobacteria
Planctomycetes
Proteobacteria
Nitrospirae
Archaea
Eukaryota
Bacteria
Bacteria

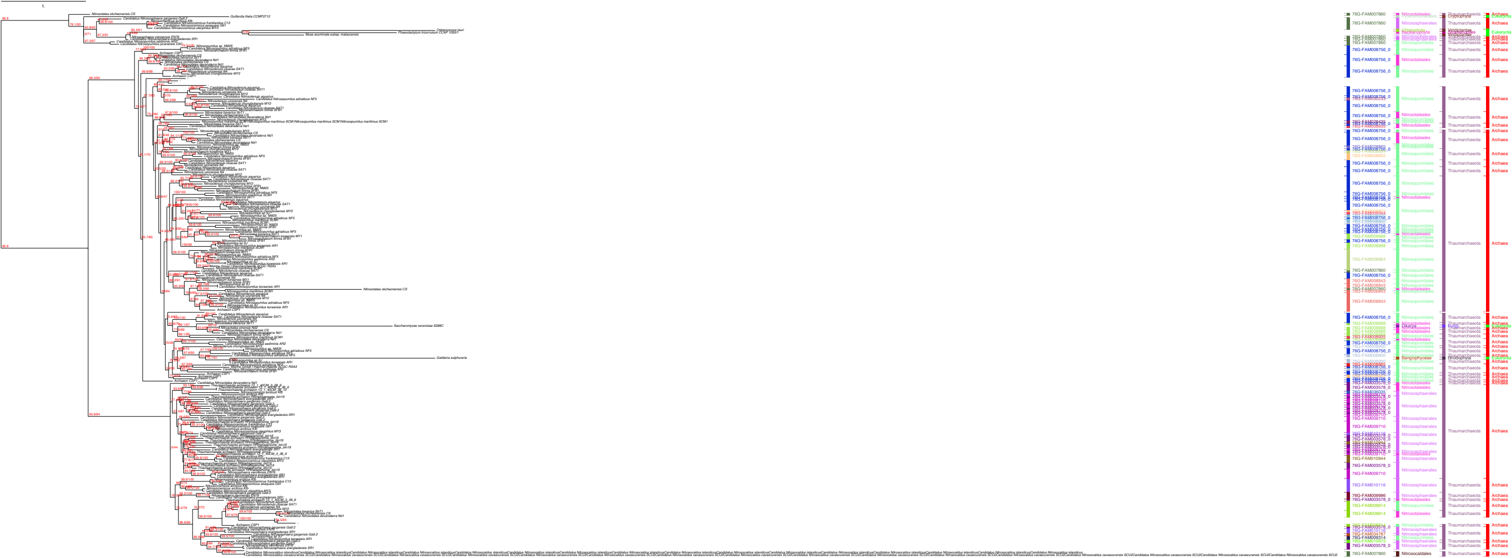

Nitrosopumilales
Nitrososphaerales
Nitrosotaleales
Nitrosocaldales
Thaumarchaeota
Archaea

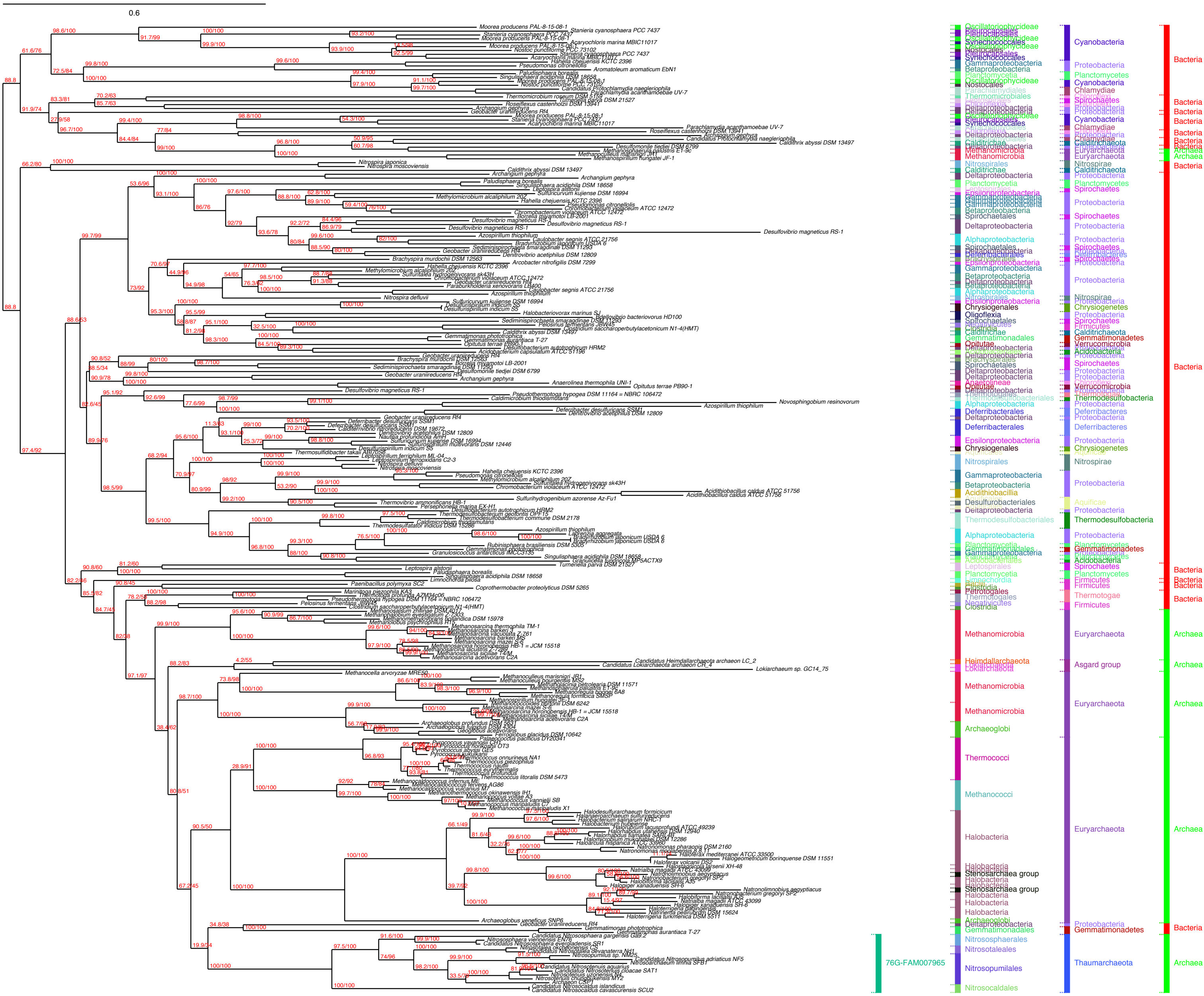
0.6
Bacteria
Archaea
Cyanobacteria
Proteobacteria
Euryarchaeota
Asgard group
Thaumarchaeota
76G-FAM007965

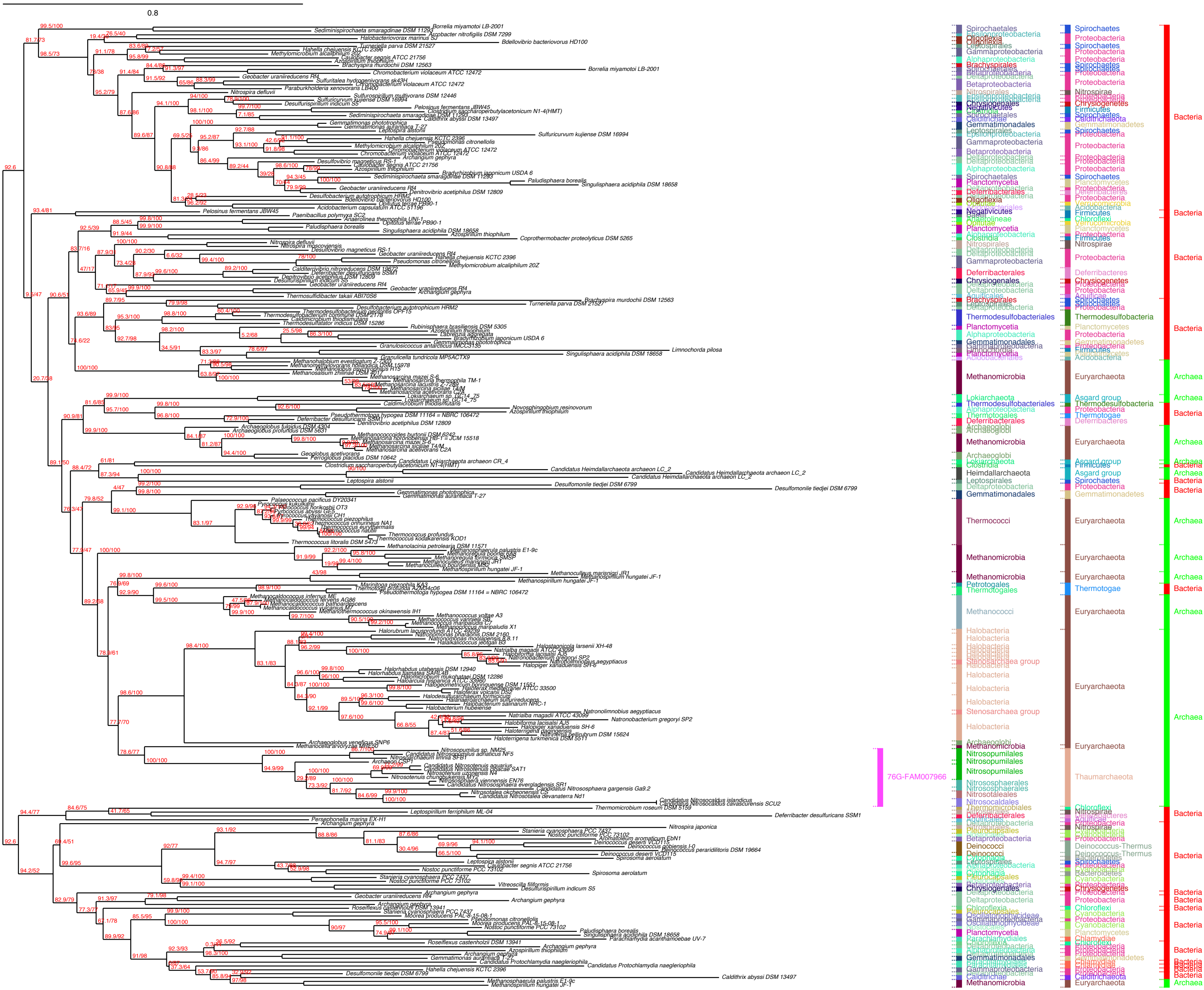

0.8
Bacteria
Archaea
76G-FAM007966

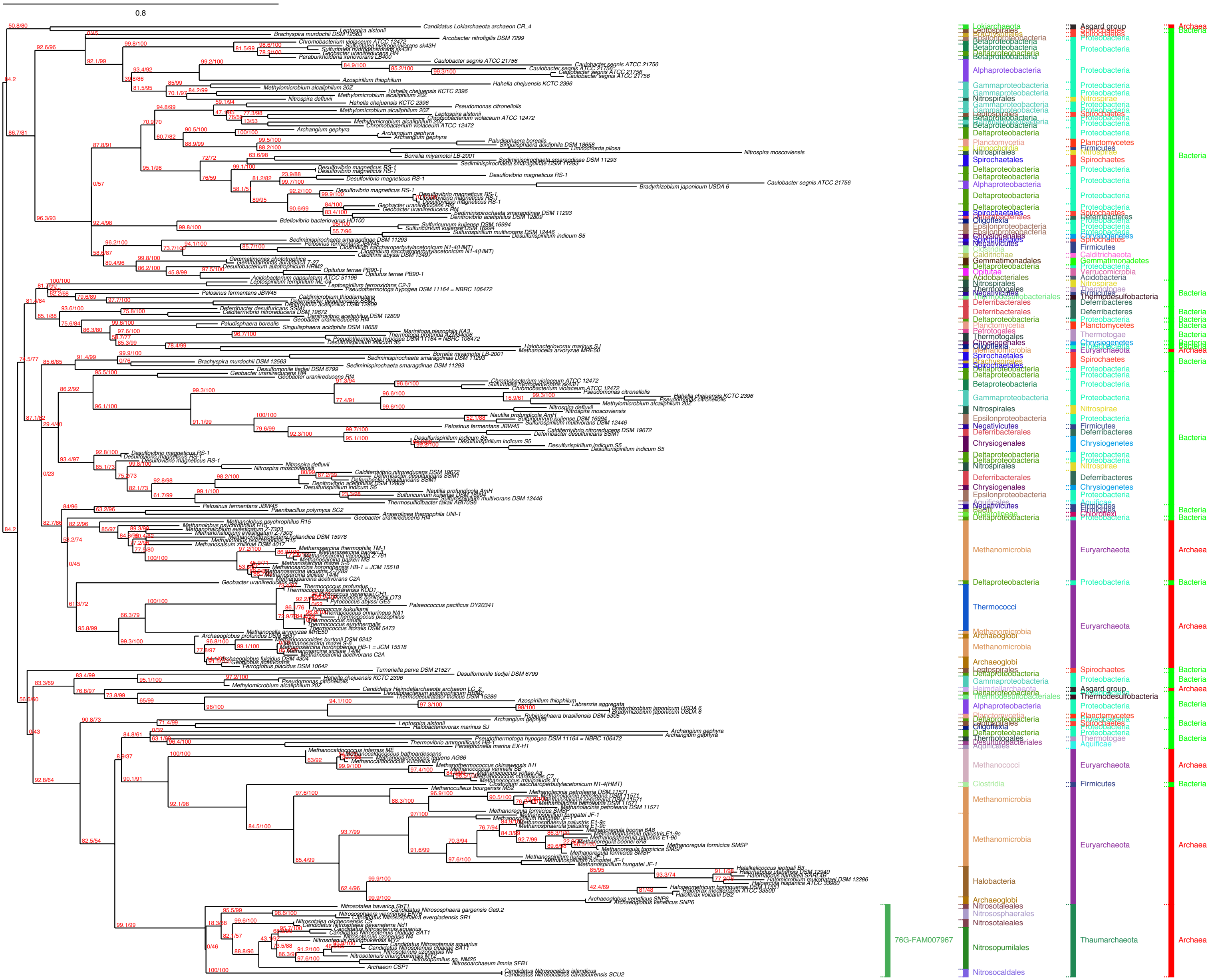
0.8
76G-FAM007967
Lokiarchaeota
Asgard group
Archaea
Bacteria
Betaproteobacteria
Alphaproteobacteria
Gammaproteobacteria
Deltaproteobacteria
Proteobacteria
Nitrospirales
Nitrospirae
Planctomycetia
Planctomycetes
Spirochaetales
Spirochaetes
Firmicutes
Gemmatimonadales
Gemmatimonadetes
Deferribacterales
Deferribacteres
Chrysiogenales
Chrysiogenetes
Thermotogales
Thermotogae
Epsilonproteobacteria
Negativicutes
Methanomicrobia
Euryarchaeota
Thermococci
Archaeoglobi
Methanococci
Clostridia
Halobacteria
Nitrosotaleales
Nitrososphaerales
Nitrosopumilales
Nitrosocaldales
Thaumarchaeota
Thermodesulfobacteria
Heimdallarchaeota

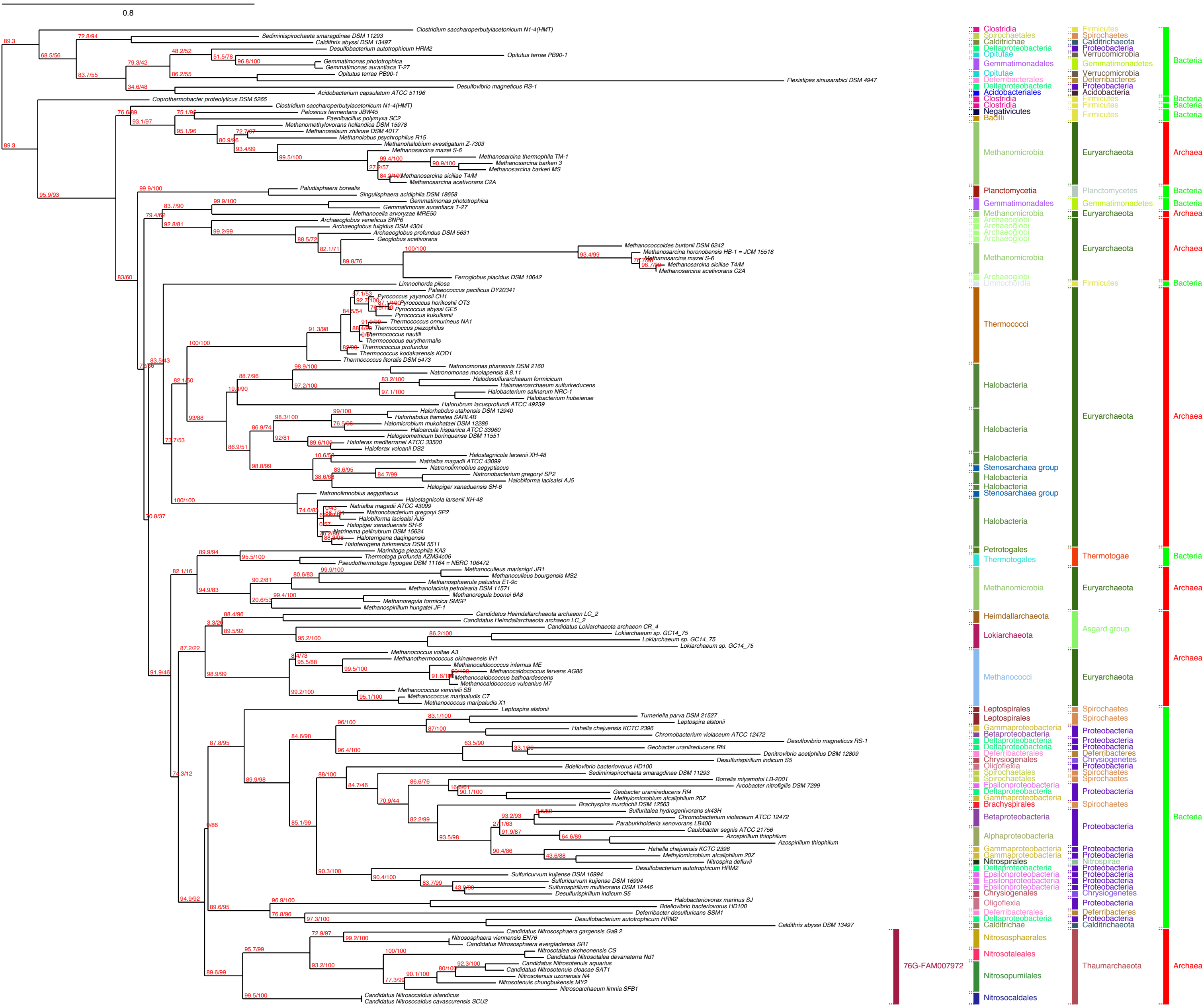

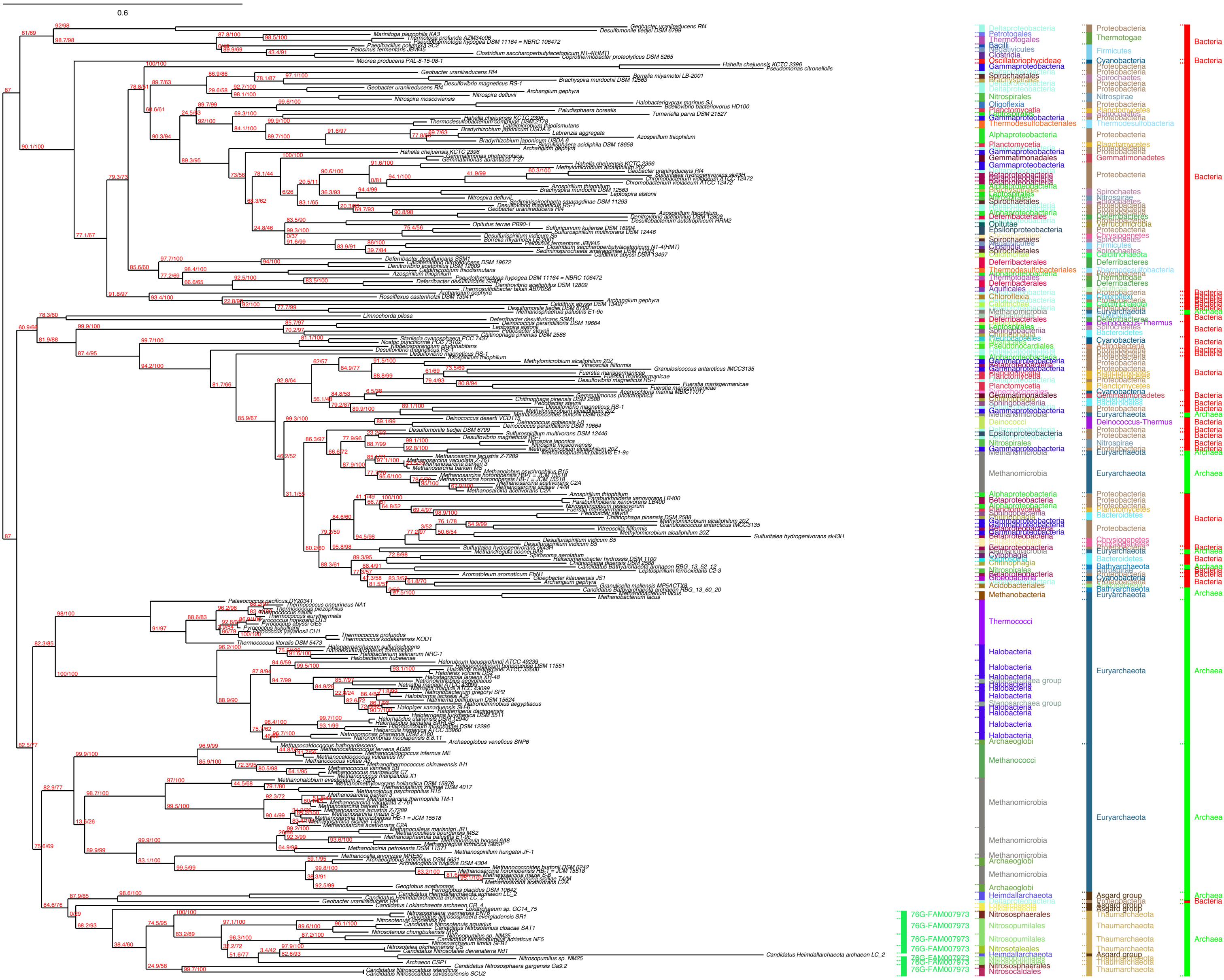

0.6
Bacteria
Archaea

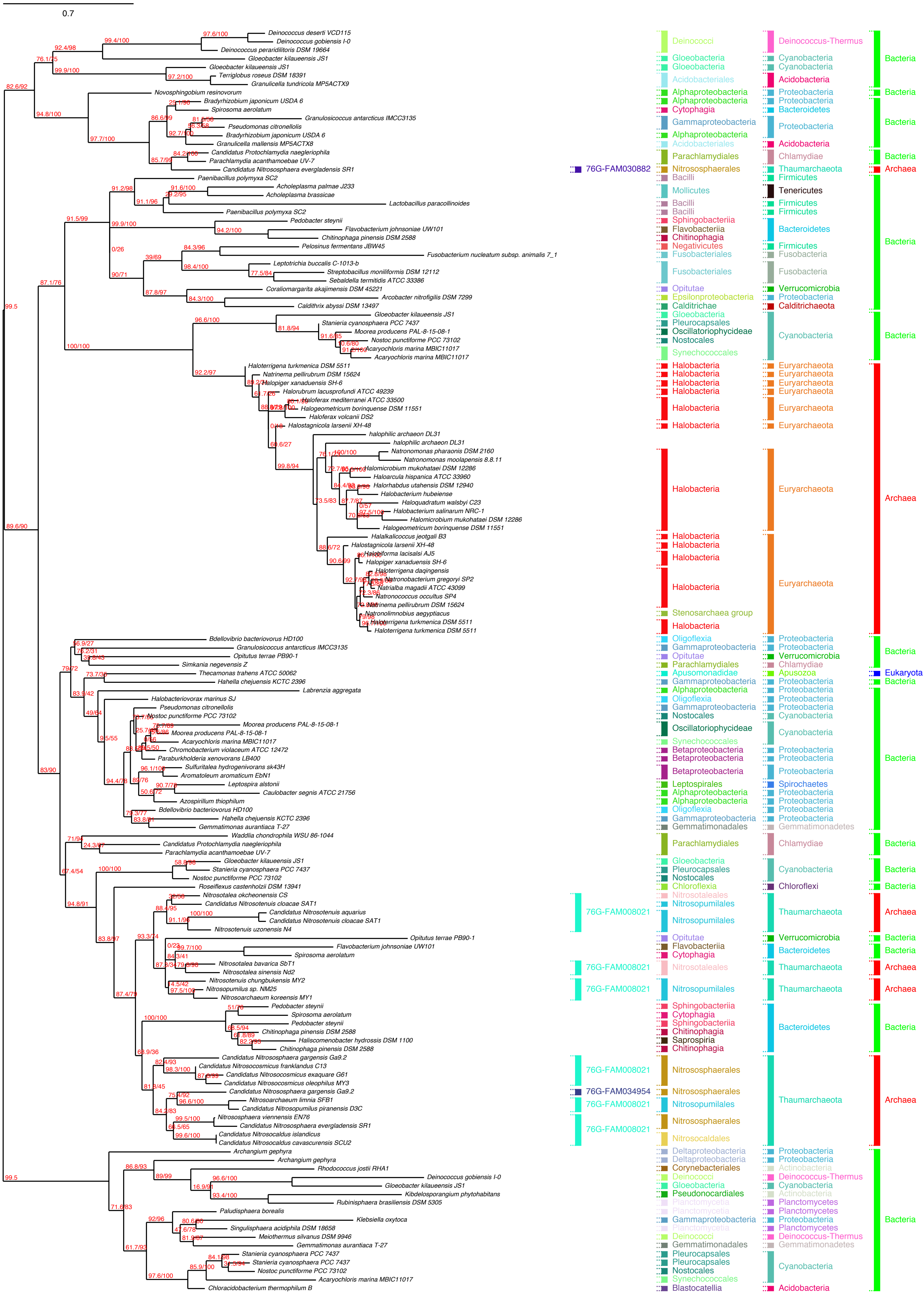

0.7
Deinococcus deserti VCD115
Deinococcus gobiensis I-0
Deinococcus peraridilitoris DSM 19664
Gloeobacter kilaueensis JS1
Gloeobacter kilaueensis JS1
Terriglobus roseus DSM 18391
Granulicella tundricola MP5ACTX9
Novosphingobium resinovorum
Bradyrhizobium japonicum USDA 6
Spirosoma aerolatum
Granulosicoccus antarcticus IMCC3135
Pseudomonas citronellolis
Bradyrhizobium japonicum USDA 6
Granulicella mallensis MP5ACTX8
Candidatus Protochlamydia naegleriophila
Parachlamydia acanthamoebae UV-7
Candidatus Nitrososphaera evergladensis SR1
Paenibacillus polymyxa SC2
Acholeplasma palmae J233
Acholeplasma brassicae
Lactobacillus paracollinoides
Paenibacillus polymyxa SC2
Pedobacter steynii
Flavobacterium johnsoniae UW101
Chitinophaga pinensis DSM 2588
Pelosinus fermentans JBW45
Fusobacterium nucleatum subsp. animalis 7_1
Leptotrichia buccalis C-1013-b
Streptobacillus moniliformis DSM 12112
Sebaldella termitidis ATCC 33386
Coraliomargarita akajimensis DSM 45221
Arcobacter nitrofigilis DSM 7299
Caldithrix abyssi DSM 13497
Gloeobacter kilaueensis JS1
Stanieria cyanosphaera PCC 7437
Moorea producens PAL-8-15-08-1
Nostoc punctiforme PCC 73102
Acaryochloris marina MBIC11017
Acaryochloris marina MBIC11017
Haloterrigena turkmenica DSM 5511
Natrinema pellirubrum DSM 15624
Halopiger xanaduensis SH-6
Halorubrum lacusprofundi ATCC 49239
Haloferax mediterranei ATCC 33500
Halogeometricum borinquense DSM 11551
Haloferax volcanii DS2
Halostagnicola larsenii XH-48
halophilic archaeon DL31
halophilic archaeon DL31
Natronomonas pharaonis DSM 2160
Natronomonas moolapensis 8.8.11
Halomicrobium mukohataei DSM 12286
Haloarcula hispanica ATCC 33960
Halorhabdus utahensis DSM 12940
Halobacterium hubeiense
Haloquadratum walsbyi C23
Halobacterium salinarum NRC-1
Halomicrobium mukohataei DSM 12286
Halogeometricum borinquense DSM 11551
Halalkalicoccus jeotgali B3
Halostagnicola larsenii XH-48
Halobiforma lacisalsi AJ5
Halopiger xanaduensis SH-6
Haloterrigena daqingensis
Natronobacterium gregoryi SP2
Natrialba magadii ATCC 43099
Natronococcus occultus SP4
Natrinema pellirubrum DSM 15624
Natronolimnobius aegyptiacus
Haloterrigena turkmenica DSM 5511
Haloterrigena turkmenica DSM 5511
Bdellovibrio bacteriovorus HD100
Granulosicoccus antarcticus IMCC3135
Opitutus terrae PB90-1
Simkania negevensis Z
Thecamonas trahens ATCC 50062
Hahella chejuensis KCTC 2396
Labrenzia aggregata
Halobacteriovorax marinus SJ
Pseudomonas citronellolis
Nostoc punctiforme PCC 73102
Moorea producens PAL-8-15-08-1
Moorea producens PAL-8-15-08-1
Acaryochloris marina MBIC11017
Chromobacterium violaceum ATCC 12472
Paraburkholderia xenovorans LB400
Sulfuritalea hydrogenivorans sk43H
Aromatoleum aromaticum EbN1
Leptospira alstonii
Caulobacter segnis ATCC 21756
Azospirillum thiophilum
Bdellovibrio bacteriovorus HD100
Hahella chejuensis KCTC 2396
Gemmatimonas aurantiaca T-27
Waddlia chondrophila WSU 86-1044
Candidatus Protochlamydia naegleriophila
Parachlamydia acanthamoebae UV-7
Gloeobacter kilaueensis JS1
Stanieria cyanosphaera PCC 7437
Nostoc punctiforme PCC 73102
Roseiflexus castenholzii DSM 13941
Nitrosotalea okcheonensis CS
Candidatus Nitrosotenuis cloacae SAT1
Candidatus Nitrosotenuis aquarius
Candidatus Nitrosotenuis cloacae SAT1
Nitrosotenuis uzonensis N4
Opitutus terrae PB90-1
Flavobacterium johnsoniae UW101
Spirosoma aerolatum
Nitrosotalea bavarica SbT1
Nitrosotalea sinensis Nd2
Nitrosotenuis chungbukensis MY2
Nitrosopumilus sp. NM25
Nitrosoarchaeum koreensis MY1
Pedobacter steynii
Spirosoma aerolatum
Pedobacter steynii
Chitinophaga pinensis DSM 2588
Haliscomenobacter hydrossis DSM 1100
Chitinophaga pinensis DSM 2588
Candidatus Nitrososphaera gargensis Ga9.2
Candidatus Nitrosocosmicus franklandus C13
Candidatus Nitrosocosmicus exaquare G61
Candidatus Nitrosocosmicus oleophilus MY3
Candidatus Nitrososphaera gargensis Ga9.2
Nitrosoarchaeum limnia SFB1
Candidatus Nitrosopumilus piranensis D3C
Nitrososphaera viennensis EN76
Candidatus Nitrososphaera evergladensis SR1
Candidatus Nitrosocaldus islandicus
Candidatus Nitrosocaldus cavascurensis SCU2
Archangium gephyra
Archangium gephyra
Rhodococcus jostii RHA1
Deinococcus gobiensis I-0
Gloeobacter kilaueensis JS1
Kibdelosporangium phytohabitans
Rubinisphaera brasiliensis DSM 5305
Paludisphaera borealis
Klebsiella oxytoca
Singulisphaera acidiphila DSM 18658
Meiothermus silvanus DSM 9946
Gemmatimonas aurantiaca T-27
Stanieria cyanosphaera PCC 7437
Stanieria cyanosphaera PCC 7437
Nostoc punctiforme PCC 73102
Acaryochloris marina MBIC11017
Chloracidobacterium thermophilum B
76G-FAM030882
76G-FAM008021
76G-FAM008021
76G-FAM008021
76G-FAM008021
76G-FAM034954
76G-FAM008021
76G-FAM008021
Deinococci
Gloeobacteria
Gloeobacteria
Acidobacteriales
Alphaproteobacteria
Alphaproteobacteria
Cytophagia
Gammaproteobacteria
Alphaproteobacteria
Acidobacteriales
Parachlamydiales
Nitrososphaerales
Bacilli
Mollicutes
Bacilli
Bacilli
Sphingobacteriia
Flavobacteriia
Chitinophagia
Negativicutes
Fusobacteriales
Fusobacteriales
Opitutae
Epsilonproteobacteria
Calditrichae
Gloeobacteria
Pleurocapsales
Oscillatoriophycideae
Nostocales
Synechococcales
Halobacteria
Stenosarchaea group
Oligoflexia
Gammaproteobacteria
Opitutae
Parachlamydiales
Apusomonadidae
Gammaproteobacteria
Alphaproteobacteria
Oligoflexia
Gammaproteobacteria
Nostocales
Oscillatoriophycideae
Synechococcales
Betaproteobacteria
Betaproteobacteria
Betaproteobacteria
Leptospirales
Alphaproteobacteria
Alphaproteobacteria
Oligoflexia
Gammaproteobacteria
Gemmatimonadales
Parachlamydiales
Gloeobacteria
Pleurocapsales
Nostocales
Chloroflexia
Nitrosotaleales
Nitrosopumilales
Nitrosopumilales
Opitutae
Flavobacteriia
Cytophagia
Nitrosotaleales
Nitrosopumilales
Sphingobacteriia
Cytophagia
Sphingobacteriia
Chitinophagia
Saprospiria
Chitinophagia
Nitrososphaerales
Nitrososphaerales
Nitrosopumilales
Nitrososphaerales
Nitrosocaldales
Deltaproteobacteria
Deltaproteobacteria
Corynebacteriales
Deinococci
Gloeobacteria
Pseudonocardiales
Planctomycetia
Planctomycetia
Gammaproteobacteria
Planctomycetia
Deinococci
Gemmatimonadales
Pleurocapsales
Pleurocapsales
Nostocales
Synechococcales
Blastocatellia
Deinococcus-Thermus
Cyanobacteria
Acidobacteria
Proteobacteria
Bacteroidetes
Chlamydiae
Thaumarchaeota
Firmicutes
Tenericutes
Fusobacteria
Verrucomicrobia
Calditrichaeota
Euryarchaeota
Apusozoa
Spirochaetes
Gemmatimonadetes
Chloroflexi
Actinobacteria
Planctomycetes
Bacteria
Archaea
Eukaryota

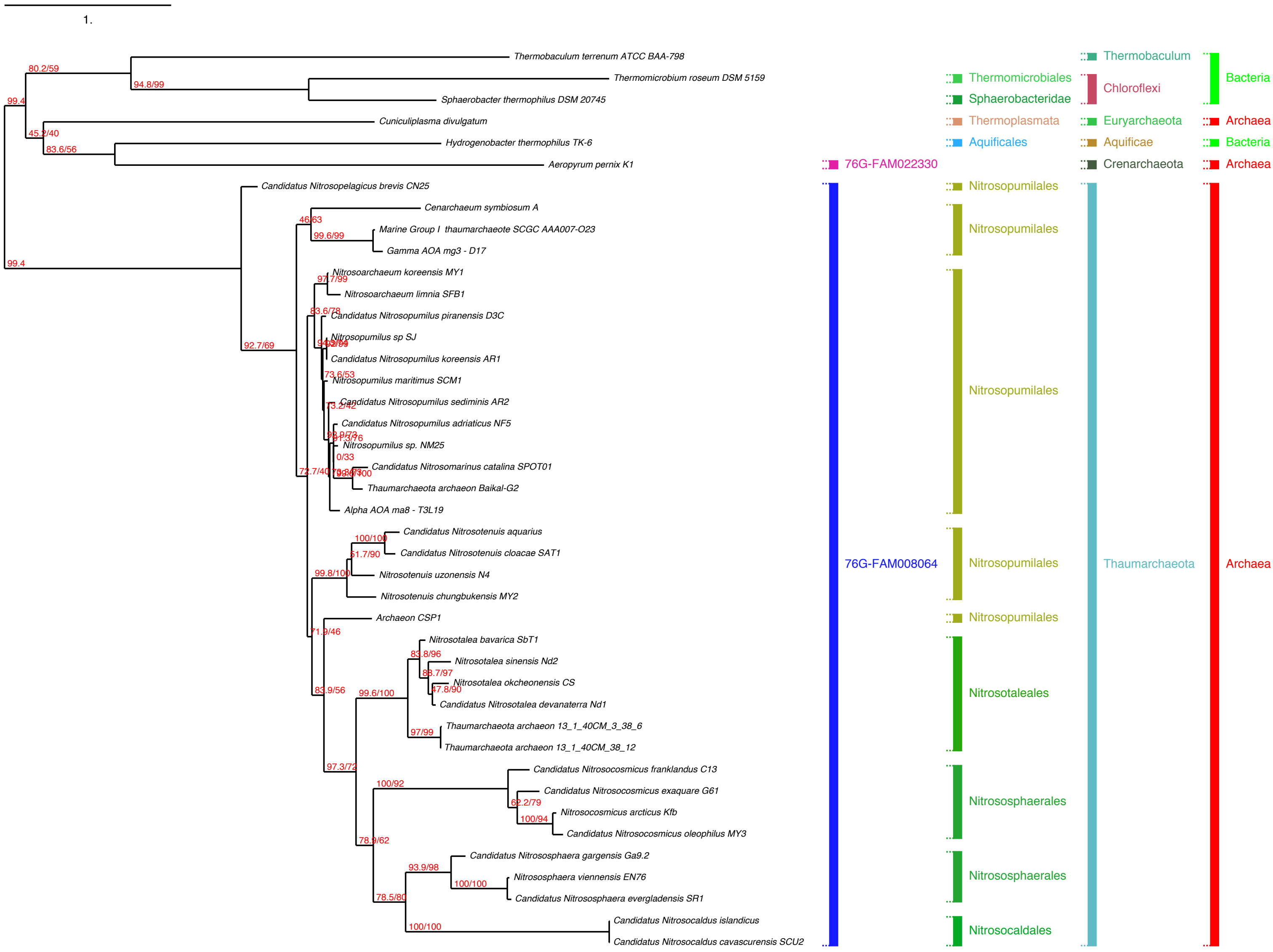
1.
Thermobaculum terrenum ATCC BAA-798
Thermomicrobium roseum DSM 5159
Sphaerobacter thermophilus DSM 20745
Cuniculiplasma divulgatum
Hydrogenobacter thermophilus TK-6
Aeropyrum pernix K1
Candidatus Nitrosopelagicus brevis CN25
Cenarchaeum symbiosum A
Marine Group I thaumarchaeote SCGC AAA007-O23
Gamma AOA mg3 - D17
Nitrosoarchaeum koreensis MY1
Nitrosoarchaeum limnia SFB1
Candidatus Nitrosopumilus piranensis D3C
Nitrosopumilus sp SJ
Candidatus Nitrosopumilus koreensis AR1
Nitrosopumilus maritimus SCM1
Candidatus Nitrosopumilus sediminis AR2
Candidatus Nitrosopumilus adriaticus NF5
Nitrosopumilus sp. NM25
Candidatus Nitrosomarinus catalina SPOT01
Thaumarchaeota archaeon Baikal-G2
Alpha AOA ma8 - T3L19
Candidatus Nitrosotenuis aquarius
Candidatus Nitrosotenuis cloacae SAT1
Nitrosotenuis uzonensis N4
Nitrosotenuis chungbukensis MY2
Archaeon CSP1
Nitrosotalea bavarica SbT1
Nitrosotalea sinensis Nd2
Nitrosotalea okcheonensis CS
Candidatus Nitrosotalea devanaterra Nd1
Thaumarchaeota archaeon 13_1_40CM_3_38_6
Thaumarchaeota archaeon 13_1_40CM_38_12
Candidatus Nitrosocosmicus franklandus C13
Candidatus Nitrosocosmicus exaquare G61
Nitrosocosmicus arcticus Kfb
Candidatus Nitrosocosmicus oleophilus MY3
Candidatus Nitrososphaera gargensis Ga9.2
Nitrososphaera viennensis EN76
Candidatus Nitrososphaera evergladensis SR1
Candidatus Nitrosocaldus islandicus
Candidatus Nitrosocaldus cavascurensis SCU2
80.2/59
94.8/99
99.4
45.2/40
83.6/56
99.4
92.7/69
46/63
99.6/99
97.7/99
83.6/78
73.6/53
73.2/42
0/33
72.7/40
100/100
51.7/90
99.8/100
71.9/46
83.8/96
88.7/97
47.8/90
83.9/56
99.6/100
97/99
97.3/72
100/92
62.2/79
100/94
78.9/62
93.9/98
100/100
78.5/80
100/100
76G-FAM022330
76G-FAM008064
Thermomicrobiales
Sphaerobacteridae
Thermoplasmata
Aquificales
Nitrosopumilales
Nitrosopumilales
Nitrosopumilales
Nitrosopumilales
Nitrosopumilales
Nitrosotaleales
Nitrososphaerales
Nitrososphaerales
Nitrosocaldales
Thermobaculum
Chloroflexi
Euryarchaeota
Aquificae
Crenarchaeota
Thaumarchaeota
Bacteria
Archaea
Bacteria
Archaea
Archaea

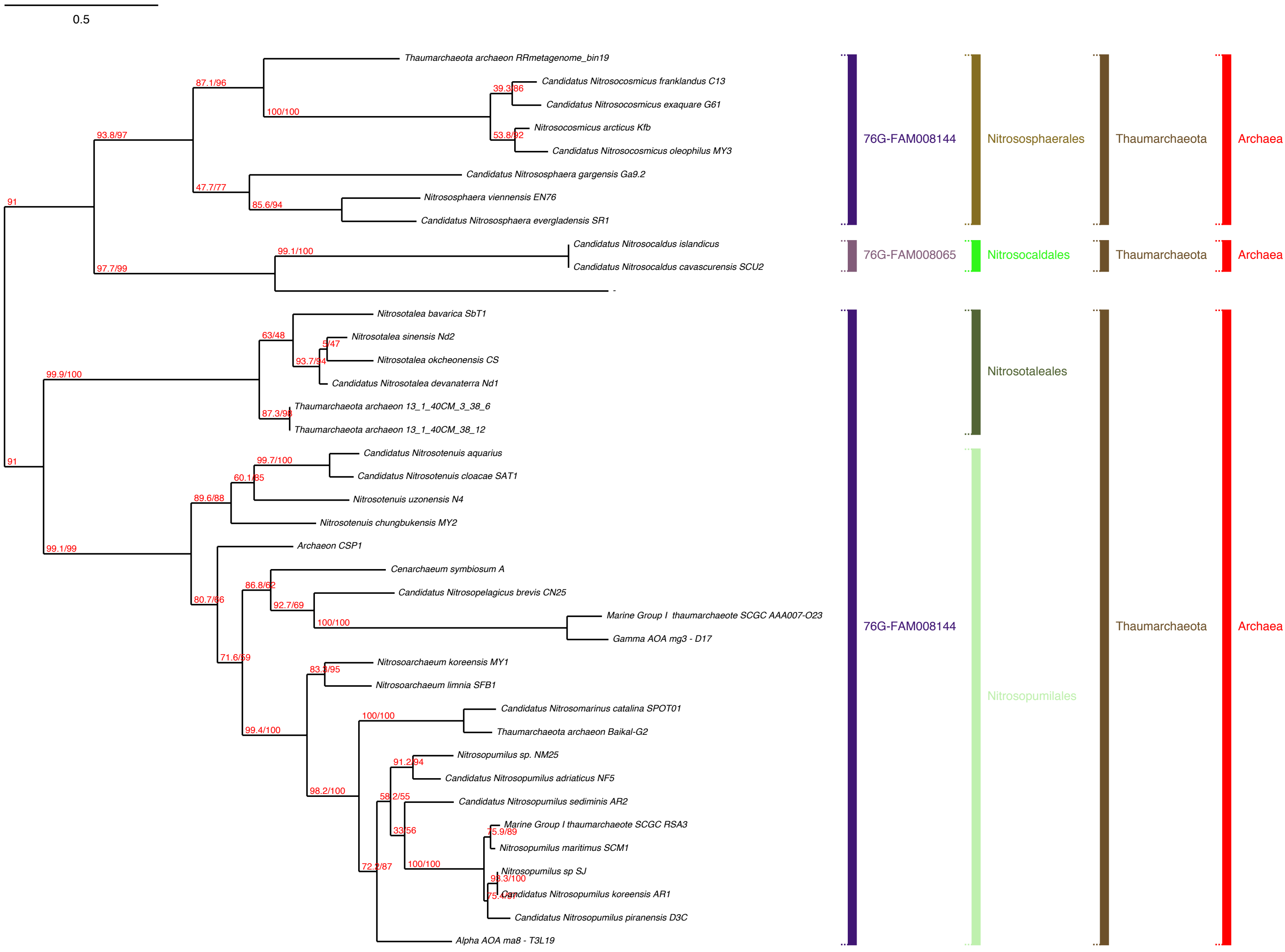
0.5
Thaumarchaeota archaeon RRmetagenome_bin19
Candidatus Nitrosocosmicus franklandus C13
Candidatus Nitrosocosmicus exaquare G61
Nitrosocosmicus arcticus Kfb
Candidatus Nitrosocosmicus oleophilus MY3
Candidatus Nitrososphaera gargensis Ga9.2
Nitrososphaera viennensis EN76
Candidatus Nitrososphaera evergladensis SR1
Candidatus Nitrosocaldus islandicus
Candidatus Nitrosocaldus cavascurensis SCU2
Nitrosotalea bavarica SbT1
Nitrosotalea sinensis Nd2
Nitrosotalea okcheonensis CS
Candidatus Nitrosotalea devanaterra Nd1
Thaumarchaeota archaeon 13_1_40CM_3_38_6
Thaumarchaeota archaeon 13_1_40CM_38_12
Candidatus Nitrosotenuis aquarius
Candidatus Nitrosotenuis cloacae SAT1
Nitrosotenuis uzonensis N4
Nitrosotenuis chungbukensis MY2
Archaeon CSP1
Cenarchaeum symbiosum A
Candidatus Nitrosopelagicus brevis CN25
Marine Group I thaumarchaeote SCGC AAA007-O23
Gamma AOA mg3 - D17
Nitrosoarchaeum koreensis MY1
Nitrosoarchaeum limnia SFB1
Candidatus Nitrosomarinus catalina SPOT01
Thaumarchaeota archaeon Baikal-G2
Nitrosopumilus sp. NM25
Candidatus Nitrosopumilus adriaticus NF5
Candidatus Nitrosopumilus sediminis AR2
Marine Group I thaumarchaeote SCGC RSA3
Nitrosopumilus maritimus SCM1
Nitrosopumilus sp SJ
Candidatus Nitrosopumilus koreensis AR1
Candidatus Nitrosopumilus piranensis D3C
Alpha AOA ma8 - T3L19
76G-FAM008144
Nitrososphaerales
Thaumarchaeota
Archaea
76G-FAM008065
Nitrosocaldales
Thaumarchaeota
Archaea
Nitrosotaleales
76G-FAM008144
Nitrosopumilales
Thaumarchaeota
Archaea

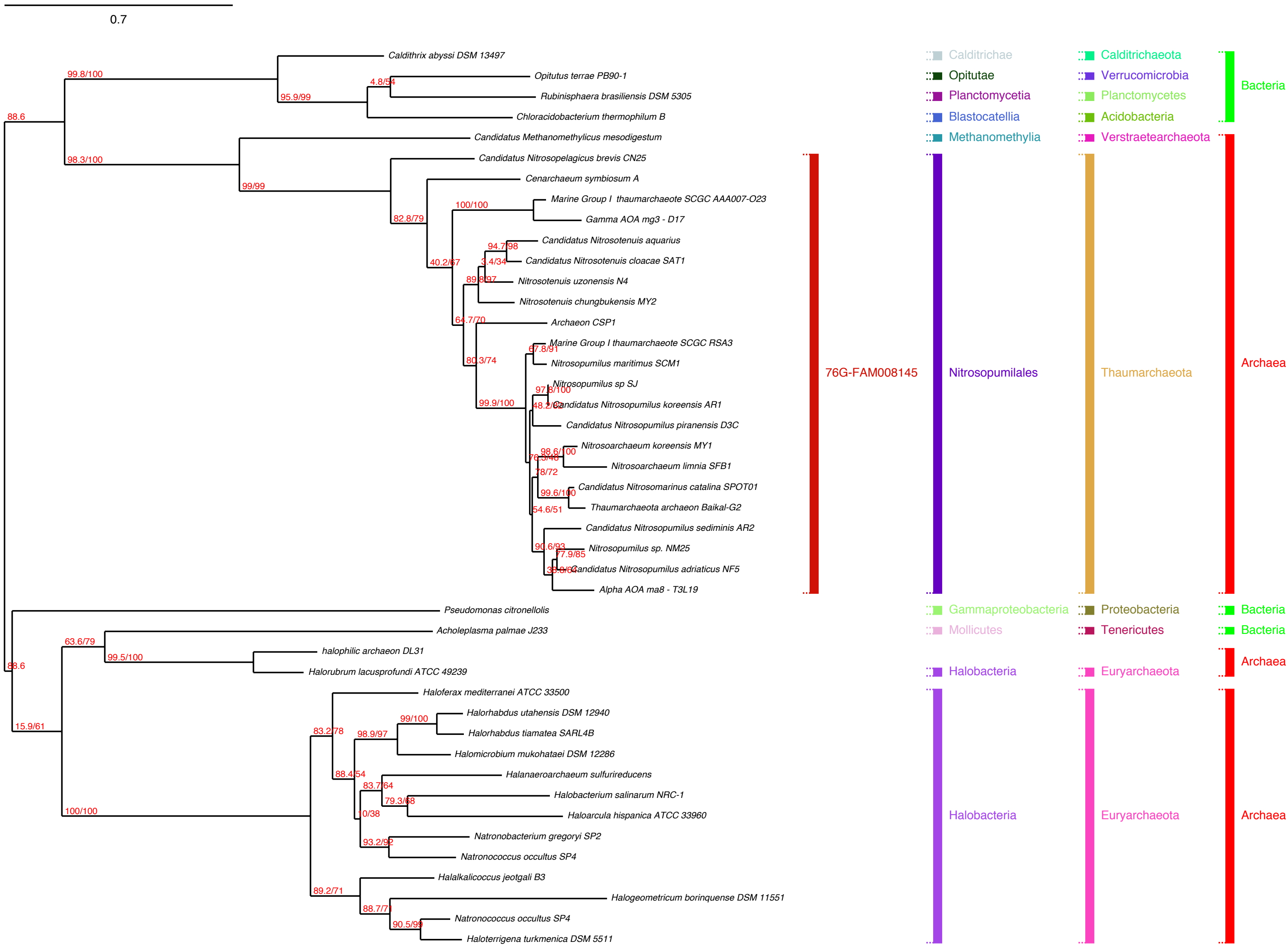

0.7
Caldithrix abyssi DSM 13497
Opitutus terrae PB90-1
Rubinisphaera brasiliensis DSM 5305
Chloracidobacterium thermophilum B
Candidatus Methanomethylicus mesodigestum
Candidatus Nitrosopelagicus brevis CN25
Cenarchaeum symbiosum A
Marine Group I thaumarchaeote SCGC AAA007-O23
Gamma AOA mg3 - D17
Candidatus Nitrosotenuis aquarius
Candidatus Nitrosotenuis cloacae SAT1
Nitrosotenuis uzonensis N4
Nitrosotenuis chungbukensis MY2
Archaeon CSP1
Marine Group I thaumarchaeote SCGC RSA3
Nitrosopumilus maritimus SCM1
Nitrosopumilus sp SJ
Candidatus Nitrosopumilus koreensis AR1
Candidatus Nitrosopumilus piranensis D3C
Nitrosoarchaeum koreensis MY1
Nitrosoarchaeum limnia SFB1
Candidatus Nitrosomarinus catalina SPOT01
Thaumarchaeota archaeon Baikal-G2
Candidatus Nitrosopumilus sediminis AR2
Nitrosopumilus sp. NM25
Candidatus Nitrosopumilus adriaticus NF5
Alpha AOA ma8 - T3L19
Pseudomonas citronellolis
Acholeplasma palmae J233
halophilic archaeon DL31
Halorubrum lacusprofundi ATCC 49239
Haloferax mediterranei ATCC 33500
Halorhabdus utahensis DSM 12940
Halorhabdus tiamatea SARL4B
Halomicrobium mukohataei DSM 12286
Halanaeroarchaeum sulfurireducens
Halobacterium salinarum NRC-1
Haloarcula hispanica ATCC 33960
Natronobacterium gregoryi SP2
Natronococcus occultus SP4
Halalkalicoccus jeotgali B3
Halogeometricum borinquense DSM 11551
Natronococcus occultus SP4
Haloterrigena turkmenica DSM 5511
88.6
99.8/100
95.9/99
4.8/54
98.3/100
99/99
82.8/79
40.2/67
100/100
94.7/98
89.8/97
64.7/70
80.3/74
67.8/91
99.9/100
97.8/100
48.2/62
98.6/100
78/72
99.6/100
54.6/51
90.6/93
77.9/85
88.6
63.6/79
99.5/100
15.9/61
100/100
83.2/78
99/100
98.9/97
88.4/54
83.7/64
79.3/68
10/38
93.2/92
89.2/71
88.7/71
90.5/99
76G-FAM008145
Calditrichae
Opitutae
Planctomycetia
Blastocatellia
Methanomethylia
Nitrosopumilales
Gammaproteobacteria
Mollicutes
Halobacteria
Halobacteria
Calditrichaeota
Verrucomicrobia
Planctomycetes
Acidobacteria
Verstraetearchaeota
Thaumarchaeota
Proteobacteria
Tenericutes
Euryarchaeota
Euryarchaeota
Bacteria
Archaea
Bacteria
Bacteria
Archaea
Archaea

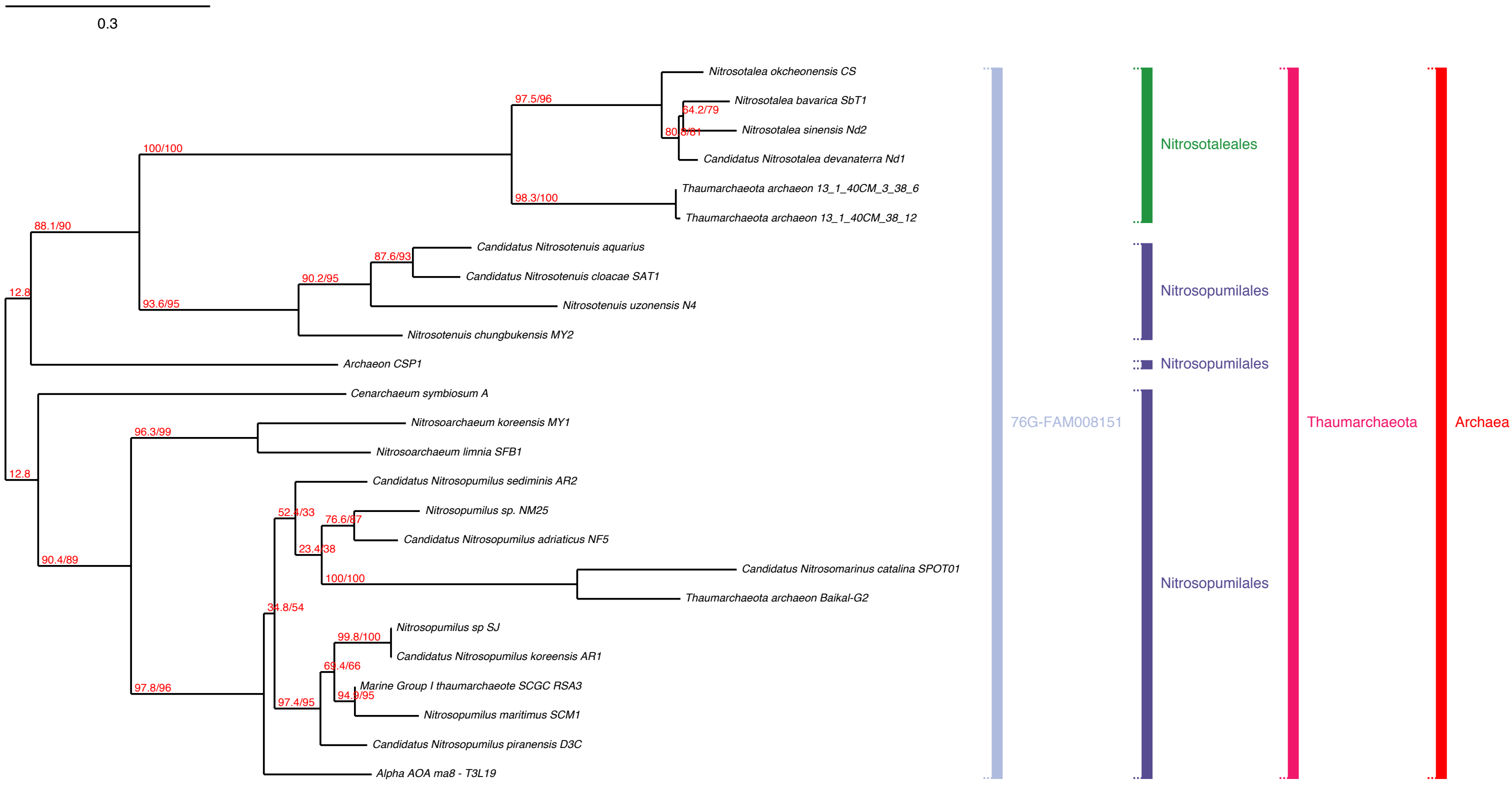

0.3
Nitrosotalea okcheonensis CS
Nitrosotalea bavarica SbT1
Nitrosotalea sinensis Nd2
Candidatus Nitrosotalea devanaterra Nd1
Thaumarchaeota archaeon 13_1_40CM_3_38_6
Thaumarchaeota archaeon 13_1_40CM_38_12
Candidatus Nitrosotenuis aquarius
Candidatus Nitrosotenuis cloacae SAT1
Nitrosotenuis uzonensis N4
Nitrosotenuis chungbukensis MY2
Archaeon CSP1
Cenarchaeum symbiosum A
Nitrosoarchaeum koreensis MY1
Nitrosoarchaeum limnia SFB1
Candidatus Nitrosopumilus sediminis AR2
Nitrosopumilus sp. NM25
Candidatus Nitrosopumilus adriaticus NF5
Candidatus Nitrosomarinus catalina SPOT01
Thaumarchaeota archaeon Baikal-G2
Nitrosopumilus sp SJ
Candidatus Nitrosopumilus koreensis AR1
Marine Group I thaumarchaeote SCGC RSA3
Nitrosopumilus maritimus SCM1
Candidatus Nitrosopumilus piranensis D3C
Alpha AOA ma8 - T3L19
97.5/96
64.2/79
80.8/81
100/100
98.3/100
88.1/90
87.6/93
90.2/95
12.8
93.6/95
96.3/99
12.8
52.4/33
76.6/87
23.4/38
90.4/89
100/100
34.8/54
99.8/100
69.4/66
94.9/95
97.8/96
97.4/95
76G-FAM008151
Nitrosotaleales
Nitrosopumilales
Nitrosopumilales
Nitrosopumilales
Thaumarchaeota
Archaea

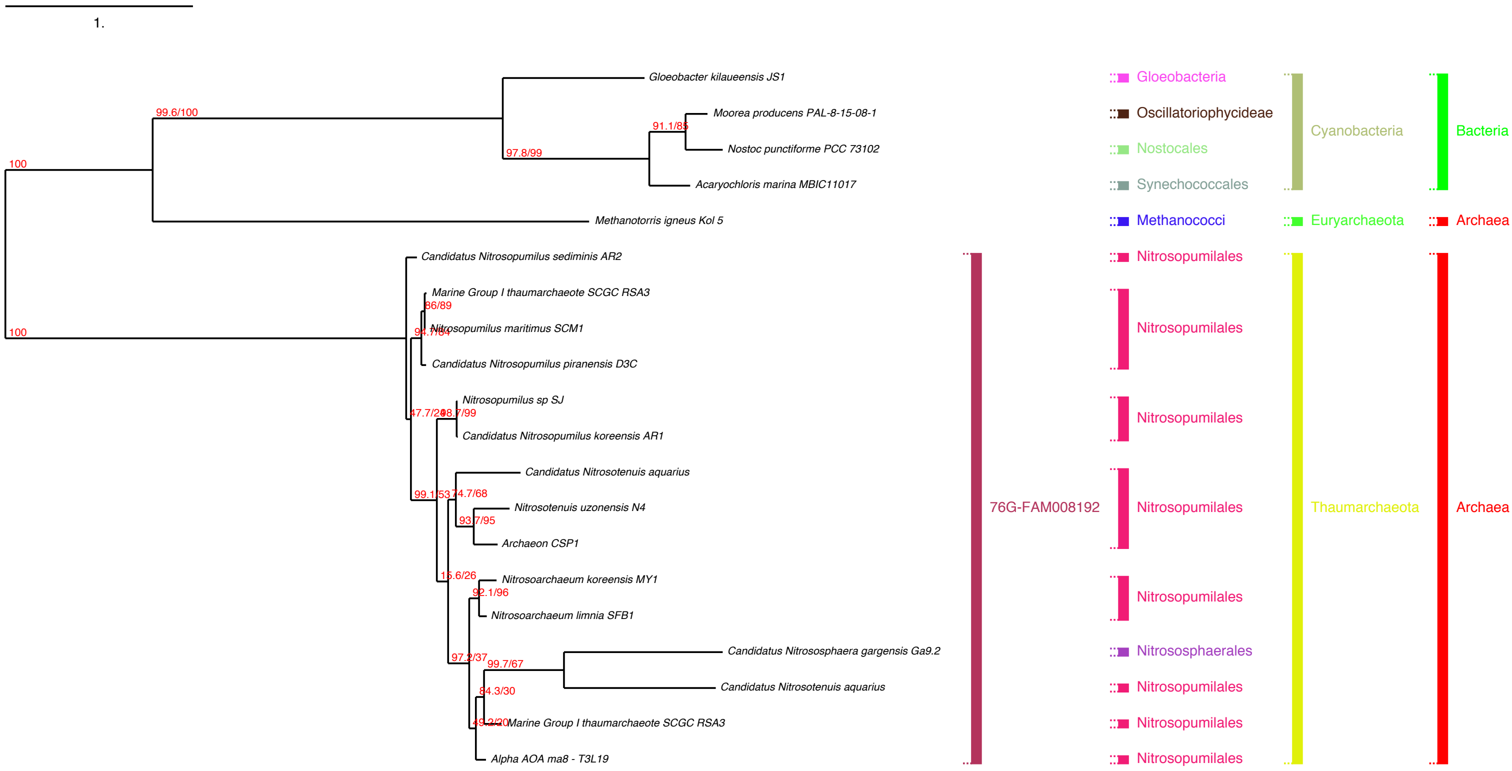

1.
Gloeobacter kilaueensis JS1
Moorea producens PAL-8-15-08-1
Nostoc punctiforme PCC 73102
Acaryochloris marina MBIC11017
Methanotorris igneus Kol 5
Candidatus Nitrosopumilus sediminis AR2
Marine Group I thaumarchaeote SCGC RSA3
Nitrosopumilus maritimus SCM1
Candidatus Nitrosopumilus piranensis D3C
Nitrosopumilus sp SJ
Candidatus Nitrosopumilus koreensis AR1
Candidatus Nitrosotenuis aquarius
Nitrosotenuis uzonensis N4
Archaeon CSP1
Nitrosoarchaeum koreensis MY1
Nitrosoarchaeum limnia SFB1
Candidatus Nitrososphaera gargensis Ga9.2
Candidatus Nitrosotenuis aquarius
Marine Group I thaumarchaeote SCGC RSA3
Alpha AOA ma8 - T3L19
100
99.6/100
97.8/99
91.1/85
100
86/89
94.7/84
47.7/20
98.7/99
99.1/53
74.7/68
93.7/95
15.6/26
92.1/96
97.2/37
99.7/67
84.3/30
49.2/20
76G-FAM008192
Gloeobacteria
Oscillatoriophycideae
Nostocales
Synechococcales
Methanococci
Nitrosopumilales
Nitrosopumilales
Nitrosopumilales
Nitrosopumilales
Nitrosopumilales
Nitrososphaerales
Nitrosopumilales
Nitrosopumilales
Nitrosopumilales
Cyanobacteria
Euryarchaeota
Thaumarchaeota
Bacteria
Archaea
Archaea

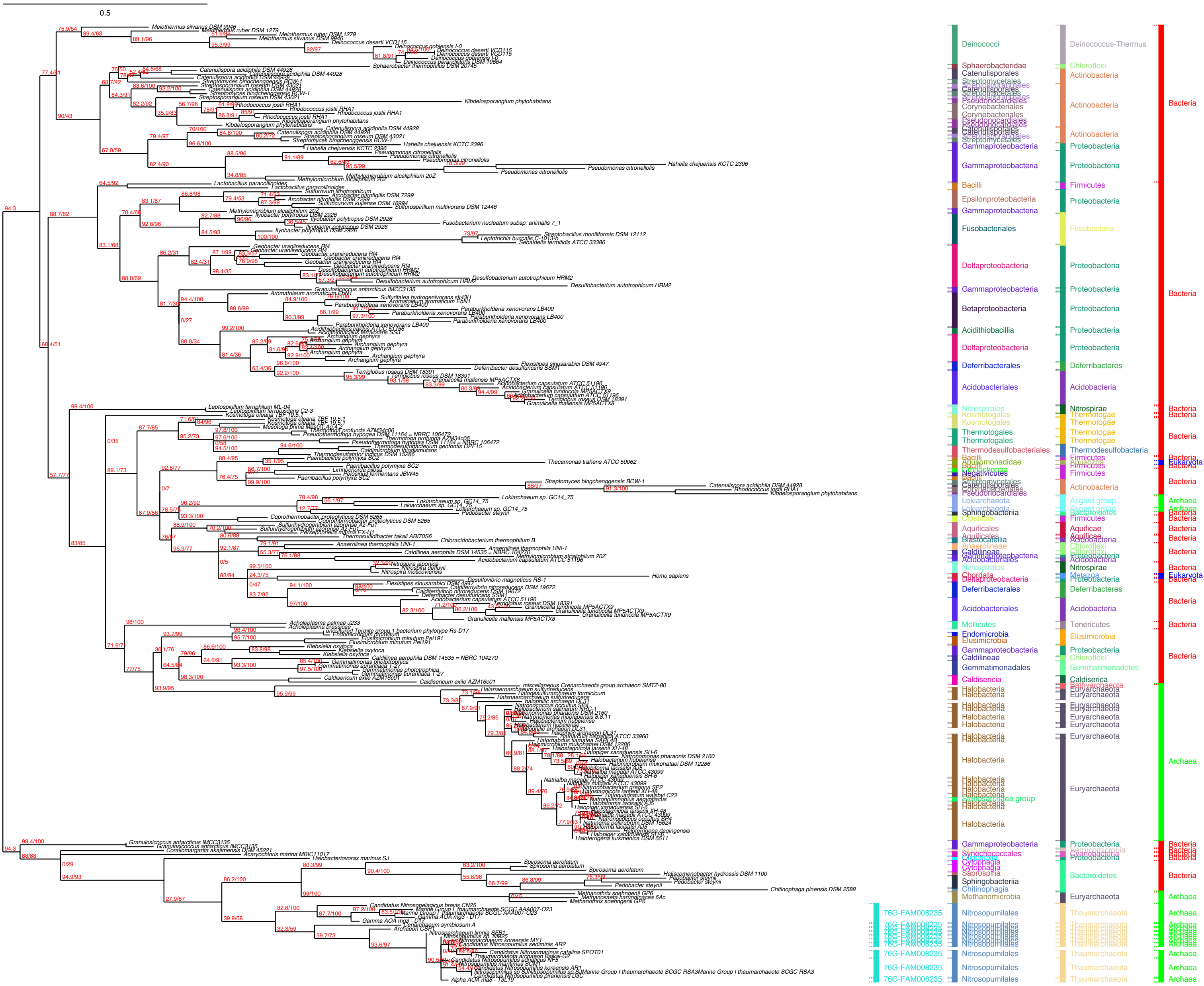

0.5
Deinococci
Sphaerobacteridae
Catenulisporales
Streptomycetales
Pseudonocardiales
Corynebacteriales
Gammaproteobacteria
Bacilli
Epsilonproteobacteria
Fusobacteriales
Deltaproteobacteria
Betaproteobacteria
Acidithiobacillia
Deferribacterales
Acidobacteriales
Nitrospirales
Kosmotogales
Thermotogales
Thermodesulfobacteriales
Lokiarchaeota
Sphingobacteriia
Clostridia
Aquificales
Blastocatellia
Anaerolineae
Caldilineae
Chordata
Mollicutes
Endomicrobia
Elusimicrobia
Gemmatimonadales
Caldisericia
Halobacteria
Synechococcales
Cytophagia
Saprospiria
Chitinophagia
Methanomicrobia
76G-FAM008235
Nitrosopumilales
Deinococcus-Thermus
Chloroflexi
Actinobacteria
Proteobacteria
Firmicutes
Fusobacteria
Deferribacteres
Acidobacteria
Nitrospirae
Thermotogae
Thermodesulfobacteria
Asgard group
Bacteroidetes
Aquificae
Metazoa
Tenericutes
Gemmatimonadetes
Caldiserica
Euryarchaeota
Cyanobacteria
Thaumarchaeota
Bacteria
Eukaryota
Archaea

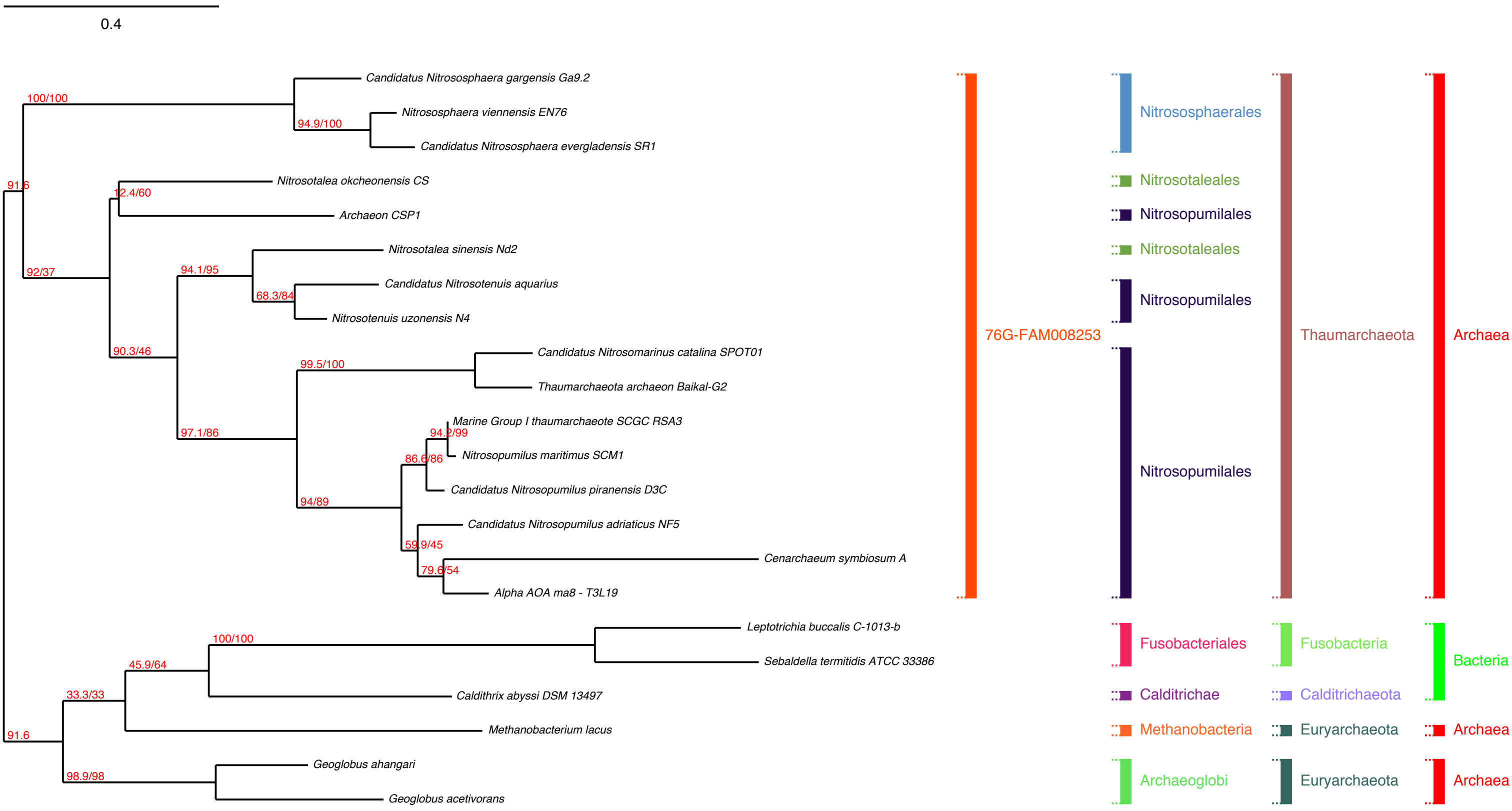
0.4
Candidatus Nitrososphaera gargensis Ga9.2
Nitrososphaera viennensis EN76
Candidatus Nitrososphaera evergladensis SR1
Nitrosotalea okcheonensis CS
Archaeon CSP1
Nitrosotalea sinensis Nd2
Candidatus Nitrosotenuis aquarius
Nitrosotenuis uzonensis N4
Candidatus Nitrosomarinus catalina SPOT01
Thaumarchaeota archaeon Baikal-G2
Marine Group I thaumarchaeote SCGC RSA3
Nitrosopumilus maritimus SCM1
Candidatus Nitrosopumilus piranensis D3C
Candidatus Nitrosopumilus adriaticus NF5
Cenarchaeum symbiosum A
Alpha AOA ma8 - T3L19
Leptotrichia buccalis C-1013-b
Sebaldella termitidis ATCC 33386
Caldithrix abyssi DSM 13497
Methanobacterium lacus
Geoglobus ahangari
Geoglobus acetivorans
100/100
94.9/100
91.6
12.4/60
92/37
94.1/95
68.3/84
90.3/46
99.5/100
97.1/86
94.2/99
86.6/86
94/89
59.9/45
79.6/54
100/100
45.9/64
33.3/33
91.6
98.9/98
76G-FAM008253
Nitrososphaerales
Nitrosotaleales
Nitrosopumilales
Nitrosotaleales
Nitrosopumilales
Nitrosopumilales
Fusobacteriales
Calditrichae
Methanobacteria
Archaeoglobi
Thaumarchaeota
Fusobacteria
Calditrichaeota
Euryarchaeota
Euryarchaeota
Archaea
Bacteria
Archaea
Archaea

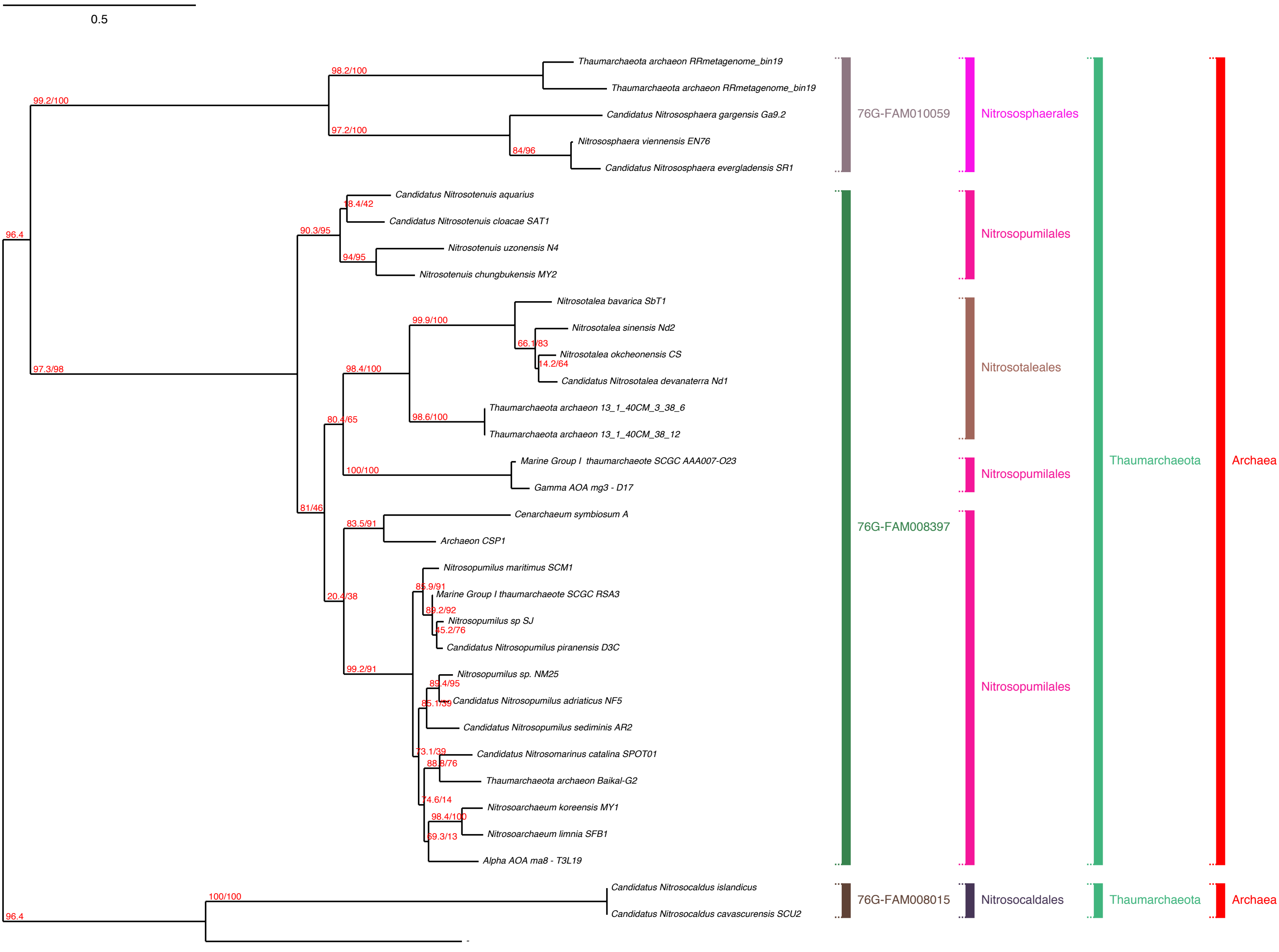
0.5
Thaumarchaeota archaeon RRmetagenome_bin19
Thaumarchaeota archaeon RRmetagenome_bin19
Candidatus Nitrososphaera gargensis Ga9.2
Nitrososphaera viennensis EN76
Candidatus Nitrososphaera evergladensis SR1
Candidatus Nitrosotenuis aquarius
Candidatus Nitrosotenuis cloacae SAT1
Nitrosotenuis uzonensis N4
Nitrosotenuis chungbukensis MY2
Nitrosotalea bavarica SbT1
Nitrosotalea sinensis Nd2
Nitrosotalea okcheonensis CS
Candidatus Nitrosotalea devanaterra Nd1
Thaumarchaeota archaeon 13_1_40CM_3_38_6
Thaumarchaeota archaeon 13_1_40CM_38_12
Marine Group I thaumarchaeote SCGC AAA007-O23
Gamma AOA mg3 - D17
Cenarchaeum symbiosum A
Archaeon CSP1
Nitrosopumilus maritimus SCM1
Marine Group I thaumarchaeote SCGC RSA3
Nitrosopumilus sp SJ
Candidatus Nitrosopumilus piranensis D3C
Nitrosopumilus sp. NM25
Candidatus Nitrosopumilus adriaticus NF5
Candidatus Nitrosopumilus sediminis AR2
Candidatus Nitrosomarinus catalina SPOT01
Thaumarchaeota archaeon Baikal-G2
Nitrosoarchaeum koreensis MY1
Nitrosoarchaeum limnia SFB1
Alpha AOA ma8 - T3L19
Candidatus Nitrosocaldus islandicus
Candidatus Nitrosocaldus cavascurensis SCU2
76G-FAM010059
Nitrososphaerales
Nitrosopumilales
Nitrosotaleales
Nitrosopumilales
76G-FAM008397
Nitrosopumilales
Thaumarchaeota
Archaea
76G-FAM008015
Nitrosocaldales
Thaumarchaeota
Archaea
99.2/100
98.2/100
97.2/100
84/96
96.4
18.4/42
90.3/95
94/95
99.9/100
66.1/83
14.2/64
97.3/98
98.4/100
98.6/100
80.4/65
100/100
81/46
83.5/91
85.9/91
20.4/38
89.2/92
45.2/76
99.2/91
89.4/95
85.1/39
73.1/39
88.8/76
74.6/14
98.4/100
69.3/13
100/100
96.4

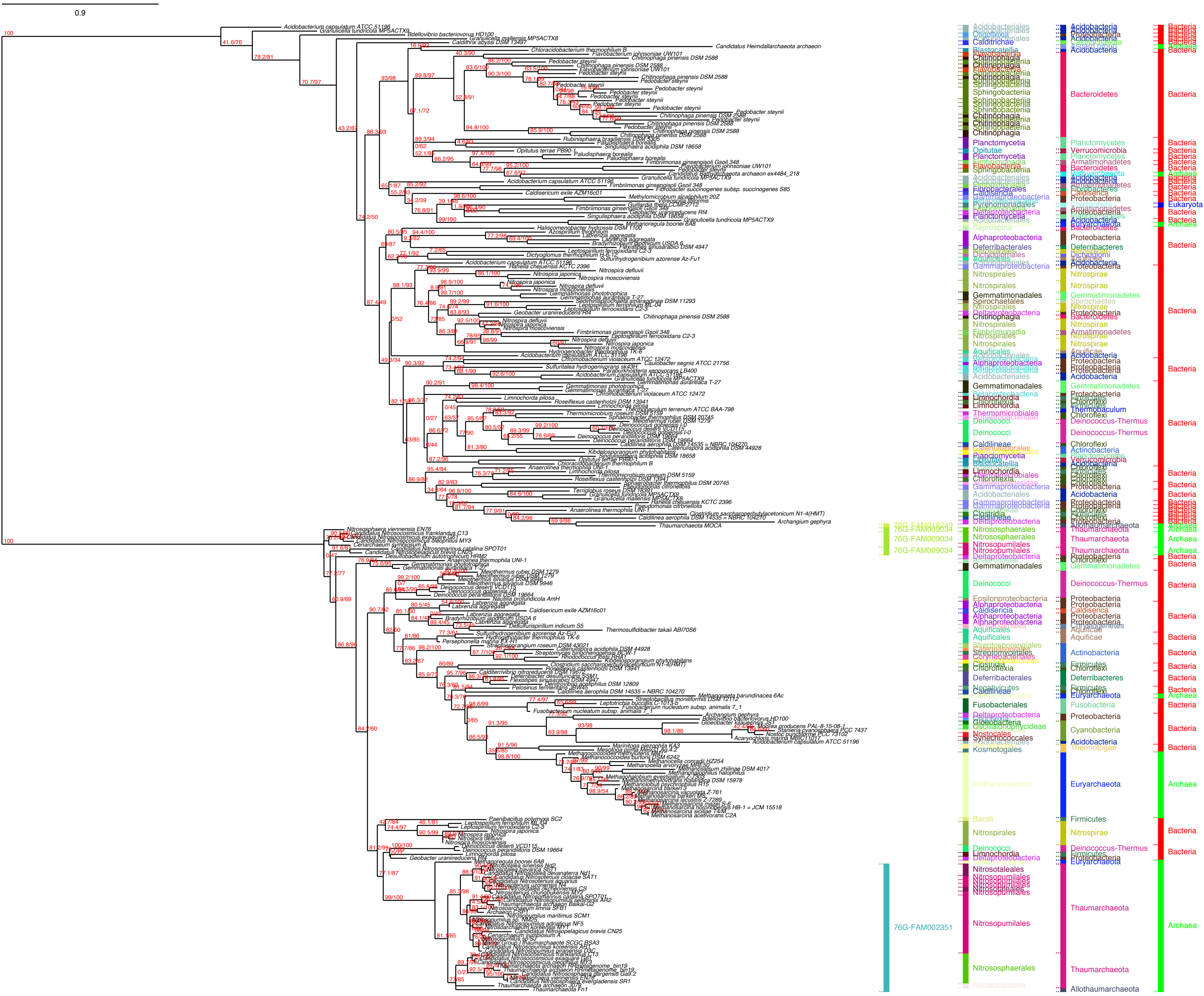

0.9
76G-FAM009034
76G-FAM002351
Bacteroidetes
Bacteria
Archaea
Eukaryota
Thaumarchaeota
Euryarchaeota
Deinococcus-Thermus
Nitrosopumilales
Nitrososphaerales
Methanomicrobia
Firmicutes
Nitrospirae
Proteobacteria
Fusobacteria
Cyanobacteria
Actinobacteria
Deferribacteres
Aquificae
Allothaumarchaeota

2.

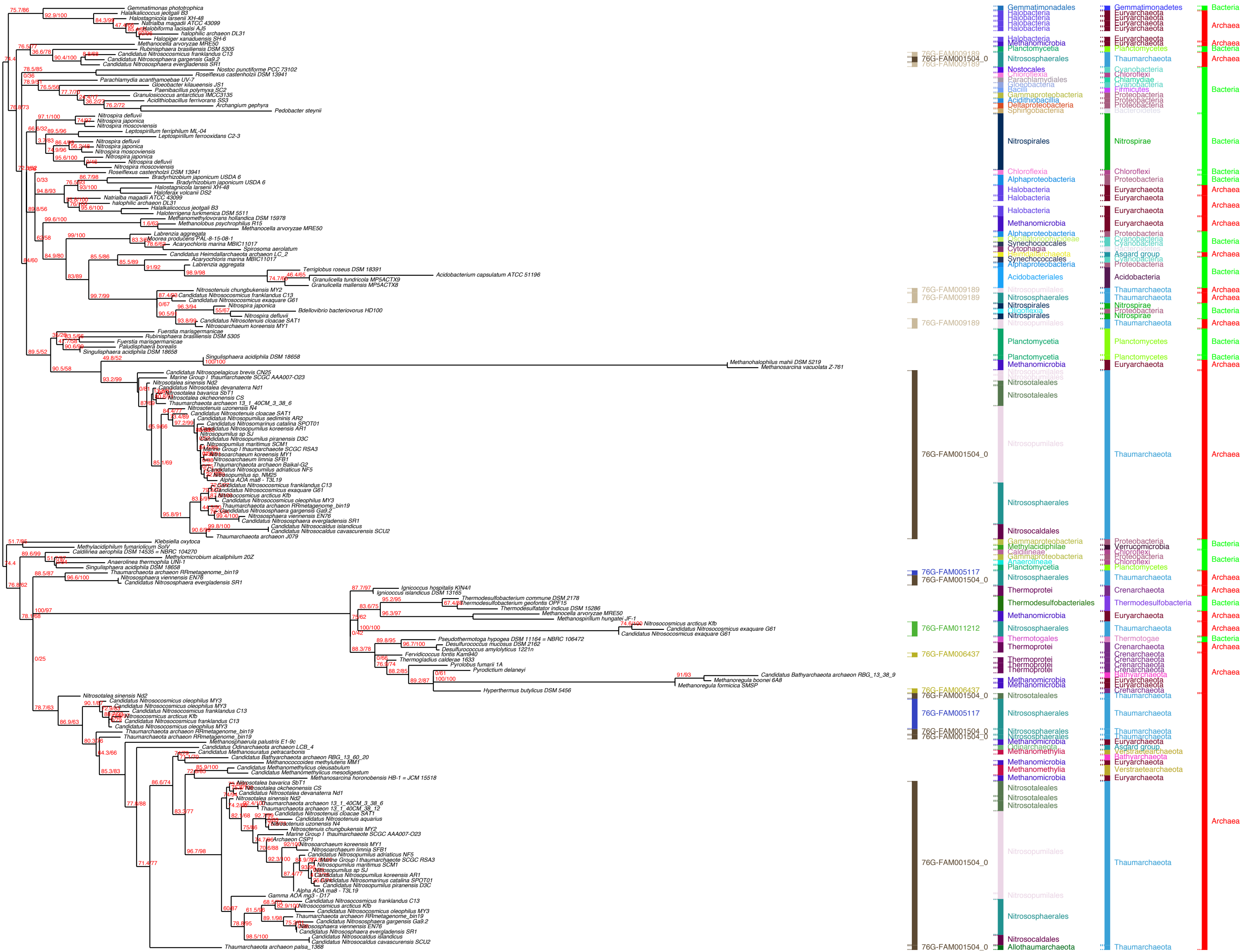

76G-FAM009189
76G-FAM001504_0
76G-FAM009189
76G-FAM009189
76G-FAM009189
76G-FAM001504_0
76G-FAM005117
76G-FAM001504_0
76G-FAM011212
76G-FAM006437
76G-FAM006437
76G-FAM001504_0
76G-FAM005117
76G-FAM001504_0
76G-FAM001504_0
76G-FAM001504_0
76G-FAM001504_0
Gemmatimonadales
Halobacteria
Methanomicrobia
Planctomycetia
Nitrososphaerales
Nostocales
Chloroflexia
Parachlamydiales
Gloeobacteria
Bacilli
Gammaproteobacteria
Acidithiobacillia
Deltaproteobacteria
Sphingobacteriia
Nitrospirales
Alphaproteobacteria
Oscillatoriophycideae
Synechococcales
Cytophagia
Heimdallarchaeota
Acidobacteriales
Oligoflexia
Nitrosopumilales
Nitrosotaleales
Nitrosocaldales
Methylacidiphilae
Caldilineae
Anaerolineae
Thermoprotei
Thermodesulfobacteriales
Thermotogales
Odinarchaeota
Methanomethylia
Allothaumarchaeota
Gemmatimonadetes
Euryarchaeota
Planctomycetes
Thaumarchaeota
Cyanobacteria
Chloroflexi
Chlamydiae
Firmicutes
Proteobacteria
Bacteroidetes
Nitrospirae
Asgard group
Acidobacteria
Verrucomicrobia
Crenarchaeota
Thermodesulfobacteria
Thermotogae
Bathyarchaeota
Verstraetearchaeota
Bacteria
Archaea

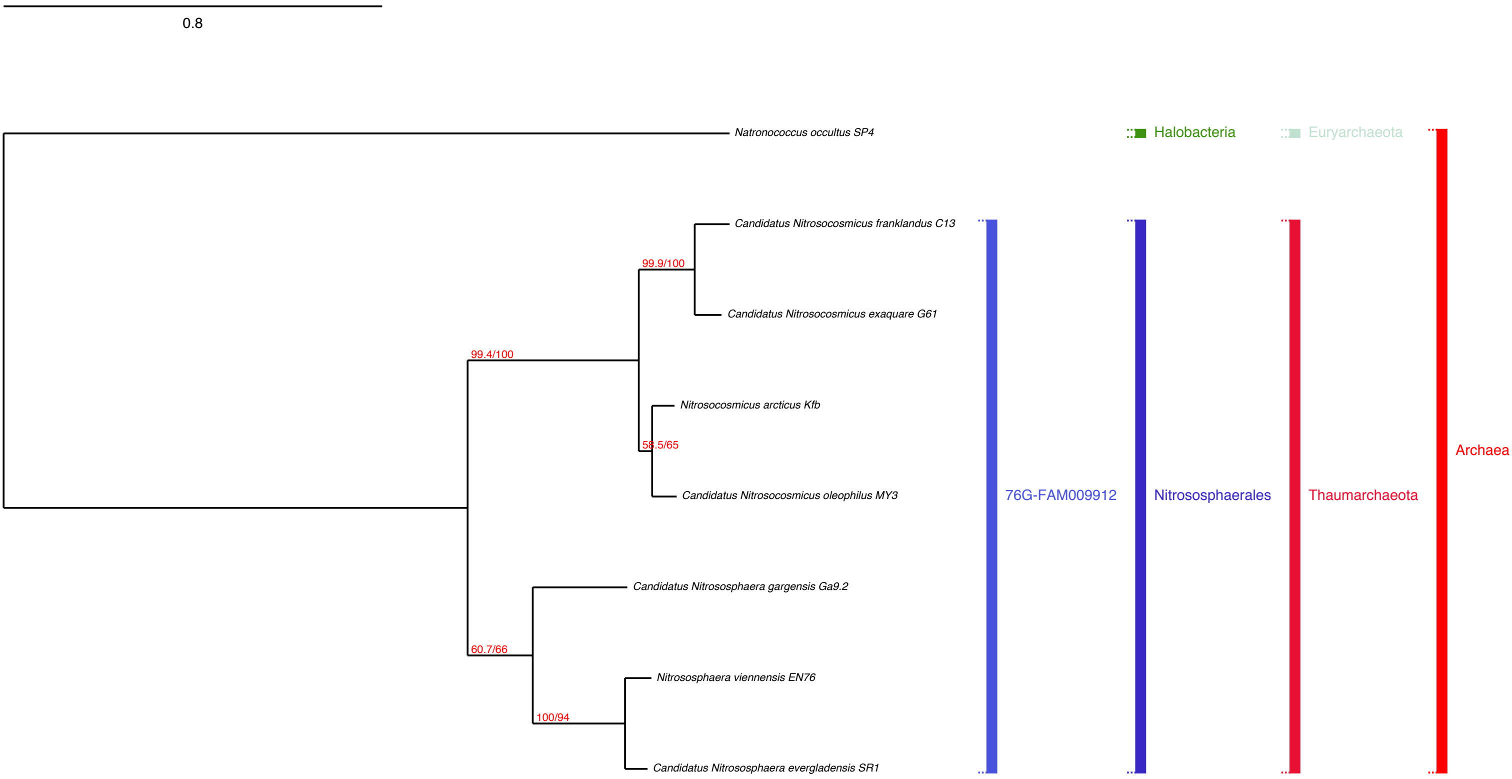
0.8
Natronococcus occultus SP4
Halobacteria
Euryarchaeota
Candidatus Nitrosocosmicus franklandus C13
99.9/100
Candidatus Nitrosocosmicus exaquare G61
99.4/100
Nitrosocosmicus arcticus Kfb
58.5/65
Candidatus Nitrosocosmicus oleophilus MY3
Candidatus Nitrososphaera gargensis Ga9.2
60.7/66
Nitrososphaera viennensis EN76
100/94
Candidatus Nitrososphaera evergladensis SR1
76G-FAM009912
Nitrososphaerales
Thaumarchaeota
Archaea

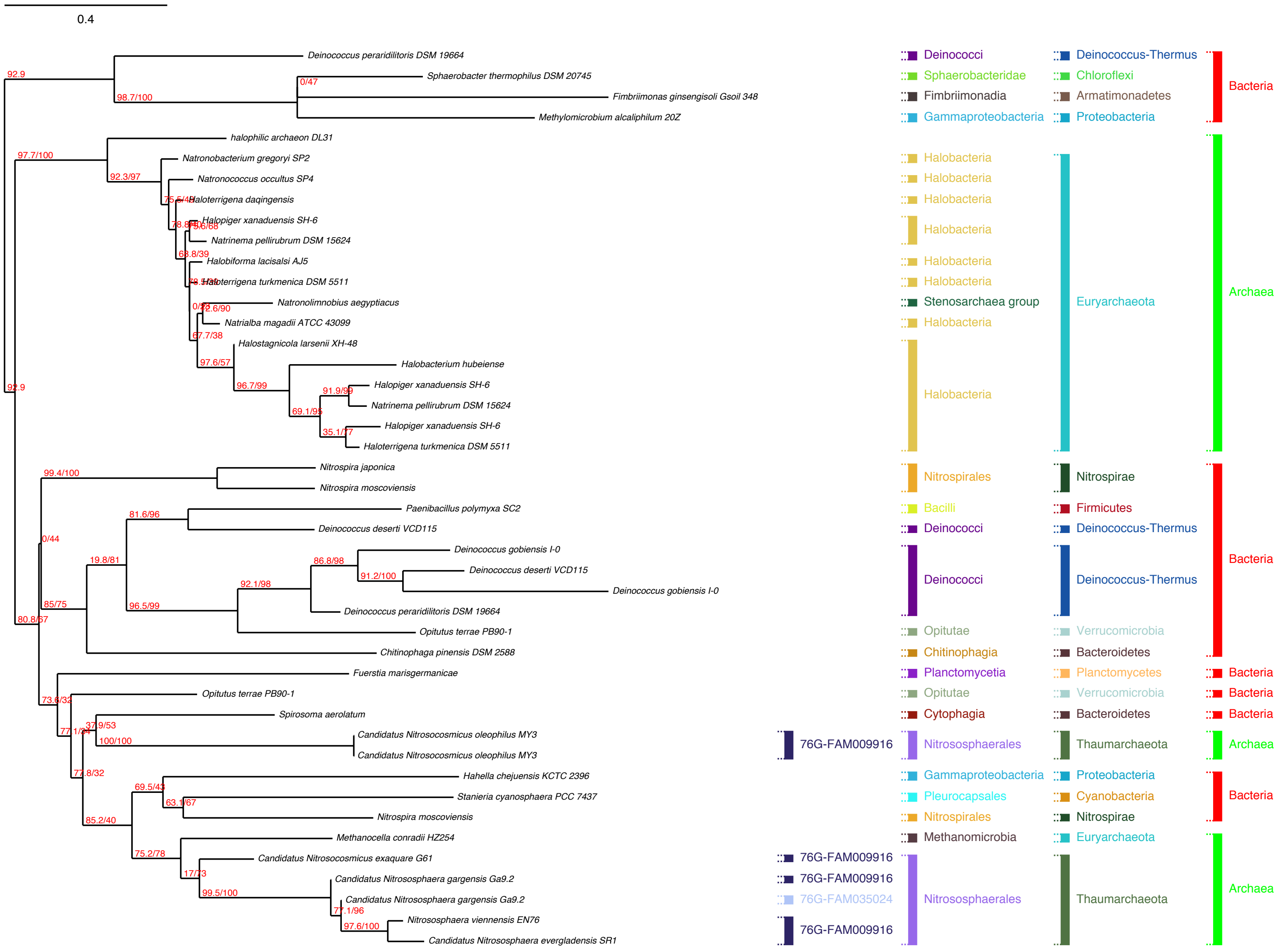
0.4
Deinococcus peraridilitoris DSM 19664
Sphaerobacter thermophilus DSM 20745
Fimbriimonas ginsengisoli Gsoil 348
Methylomicrobium alcaliphilum 20Z
halophilic archaeon DL31
Natronobacterium gregoryi SP2
Natronococcus occultus SP4
Haloterrigena daqingensis
Halopiger xanaduensis SH-6
Natrinema pellirubrum DSM 15624
Halobiforma lacisalsi AJ5
Haloterrigena turkmenica DSM 5511
Natronolimnobius aegyptiacus
Natrialba magadii ATCC 43099
Halostagnicola larsenii XH-48
Halobacterium hubeiense
Halopiger xanaduensis SH-6
Natrinema pellirubrum DSM 15624
Halopiger xanaduensis SH-6
Haloterrigena turkmenica DSM 5511
Nitrospira japonica
Nitrospira moscoviensis
Paenibacillus polymyxa SC2
Deinococcus deserti VCD115
Deinococcus gobiensis I-0
Deinococcus deserti VCD115
Deinococcus gobiensis I-0
Deinococcus peraridilitoris DSM 19664
Opitutus terrae PB90-1
Chitinophaga pinensis DSM 2588
Fuerstia marisgermanicae
Opitutus terrae PB90-1
Spirosoma aerolatum
Candidatus Nitrosocosmicus oleophilus MY3
Candidatus Nitrosocosmicus oleophilus MY3
Hahella chejuensis KCTC 2396
Stanieria cyanosphaera PCC 7437
Nitrospira moscoviensis
Methanocella conradii HZ254
Candidatus Nitrosocosmicus exaquare G61
Candidatus Nitrososphaera gargensis Ga9.2
Candidatus Nitrososphaera gargensis Ga9.2
Nitrososphaera viennensis EN76
Candidatus Nitrososphaera evergladensis SR1
92.9
98.7/100
0/47
97.7/100
92.3/97
92.9
99.4/100
81.6/96
0/44
19.8/81
86.8/98
91.2/100
92.1/98
85/75
96.5/99
80.8/67
73.6/32
37.9/53
100/100
77.8/32
69.5/43
63.1/67
85.2/40
75.2/78
17/73
99.5/100
77.1/96
97.6/100
97.6/57
96.7/99
91.9/99
69.1/95
67.7/38
Deinococci
Sphaerobacteridae
Fimbriimonadia
Gammaproteobacteria
Halobacteria
Stenosarchaea group
Nitrospirales
Bacilli
Opitutae
Chitinophagia
Planctomycetia
Cytophagia
Nitrososphaerales
Pleurocapsales
Methanomicrobia
Deinococcus-Thermus
Chloroflexi
Armatimonadetes
Proteobacteria
Euryarchaeota
Nitrospirae
Firmicutes
Verrucomicrobia
Bacteroidetes
Planctomycetes
Thaumarchaeota
Cyanobacteria
76G-FAM009916
76G-FAM035024
Bacteria
Archaea

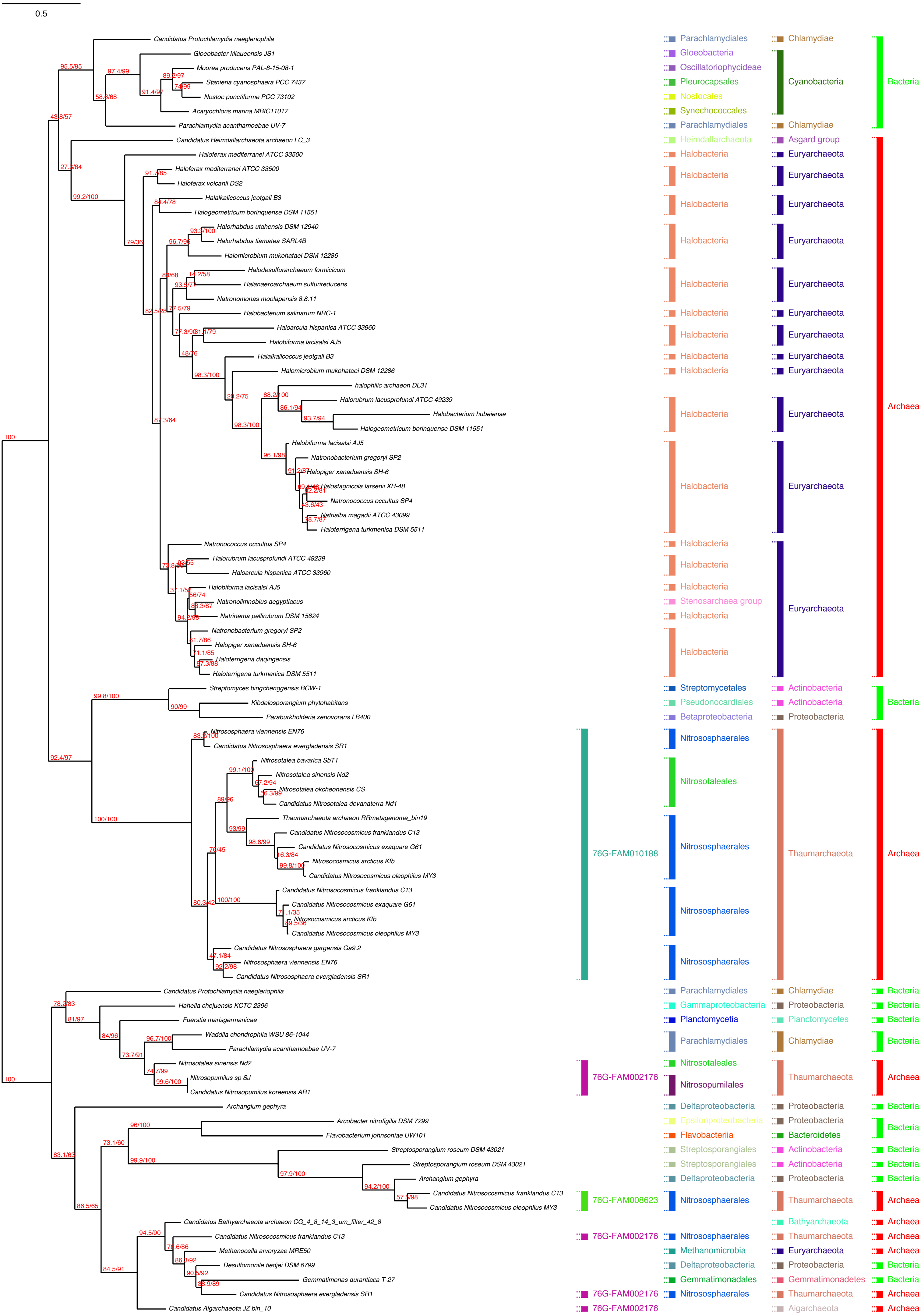
0.5
Candidatus Protochlamydia naegleriophila
Gloeobacter kilaueensis JS1
Moorea producens PAL-8-15-08-1
Stanieria cyanosphaera PCC 7437
Nostoc punctiforme PCC 73102
Acaryochloris marina MBIC11017
Parachlamydia acanthamoebae UV-7
Candidatus Heimdallarchaeota archaeon LC_3
Haloferax mediterranei ATCC 33500
Haloferax mediterranei ATCC 33500
Haloferax volcanii DS2
Halalkalicoccus jeotgali B3
Halogeometricum borinquense DSM 11551
Halorhabdus utahensis DSM 12940
Halorhabdus tiamatea SARL4B
Halomicrobium mukohataei DSM 12286
Halodesulfurarchaeum formicicum
Halanaeroarchaeum sulfurireducens
Natronomonas moolapensis 8.8.11
Halobacterium salinarum NRC-1
Haloarcula hispanica ATCC 33960
Halobiforma lacisalsi AJ5
Halalkalicoccus jeotgali B3
Halomicrobium mukohataei DSM 12286
halophilic archaeon DL31
Halorubrum lacusprofundi ATCC 49239
Halobacterium hubeiense
Halogeometricum borinquense DSM 11551
Halobiforma lacisalsi AJ5
Natronobacterium gregoryi SP2
Halopiger xanaduensis SH-6
Halostagnicola larsenii XH-48
Natronococcus occultus SP4
Natrialba magadii ATCC 43099
Haloterrigena turkmenica DSM 5511
Natronococcus occultus SP4
Halorubrum lacusprofundi ATCC 49239
Haloarcula hispanica ATCC 33960
Halobiforma lacisalsi AJ5
Natronolimnobius aegyptiacus
Natrinema pellirubrum DSM 15624
Natronobacterium gregoryi SP2
Halopiger xanaduensis SH-6
Haloterrigena daqingensis
Haloterrigena turkmenica DSM 5511
Streptomyces bingchenggensis BCW-1
Kibdelosporangium phytohabitans
Paraburkholderia xenovorans LB400
Nitrososphaera viennensis EN76
Candidatus Nitrososphaera evergladensis SR1
Nitrosotalea bavarica SbT1
Nitrosotalea sinensis Nd2
Nitrosotalea okcheonensis CS
Candidatus Nitrosotalea devanaterra Nd1
Thaumarchaeota archaeon RRmetagenome_bin19
Candidatus Nitrosocosmicus franklandus C13
Candidatus Nitrosocosmicus exaquare G61
Nitrosocosmicus arcticus Kfb
Candidatus Nitrosocosmicus oleophilus MY3
Candidatus Nitrosocosmicus franklandus C13
Candidatus Nitrosocosmicus exaquare G61
Nitrosocosmicus arcticus Kfb
Candidatus Nitrosocosmicus oleophilus MY3
Candidatus Nitrososphaera gargensis Ga9.2
Nitrososphaera viennensis EN76
Candidatus Nitrososphaera evergladensis SR1
Candidatus Protochlamydia naegleriophila
Hahella chejuensis KCTC 2396
Fuerstia marisgermanicae
Waddlia chondrophila WSU 86-1044
Parachlamydia acanthamoebae UV-7
Nitrosotalea sinensis Nd2
Nitrosopumilus sp SJ
Candidatus Nitrosopumilus koreensis AR1
Archangium gephyra
Arcobacter nitrofigilis DSM 7299
Flavobacterium johnsoniae UW101
Streptosporangium roseum DSM 43021
Streptosporangium roseum DSM 43021
Archangium gephyra
Candidatus Nitrosocosmicus franklandus C13
Candidatus Nitrosocosmicus oleophilus MY3
Candidatus Bathyarchaeota archaeon CG_4_8_14_3_um_filter_42_8
Candidatus Nitrosocosmicus franklandus C13
Methanocella arvoryzae MRE50
Desulfomonile tiedjei DSM 6799
Gemmatimonas aurantiaca T-27
Candidatus Nitrososphaera evergladensis SR1
Candidatus Aigarchaeota JZ bin_10
Parachlamydiales
Gloeobacteria
Oscillatoriophycideae
Pleurocapsales
Nostocales
Synechococcales
Parachlamydiales
Heimdallarchaeota
Halobacteria
Stenosarchaea group
Streptomycetales
Pseudonocardiales
Betaproteobacteria
Nitrososphaerales
Nitrosotaleales
Gammaproteobacteria
Planctomycetia
Nitrosopumilales
Deltaproteobacteria
Epsilonproteobacteria
Flavobacteriia
Streptosporangiales
Methanomicrobia
Gemmatimonadales
Chlamydiae
Cyanobacteria
Asgard group
Euryarchaeota
Actinobacteria
Proteobacteria
Thaumarchaeota
Planctomycetes
Bacteroidetes
Bathyarchaeota
Gemmatimonadetes
Aigarchaeota
Bacteria
Archaea
76G-FAM010188
76G-FAM002176
76G-FAM008623

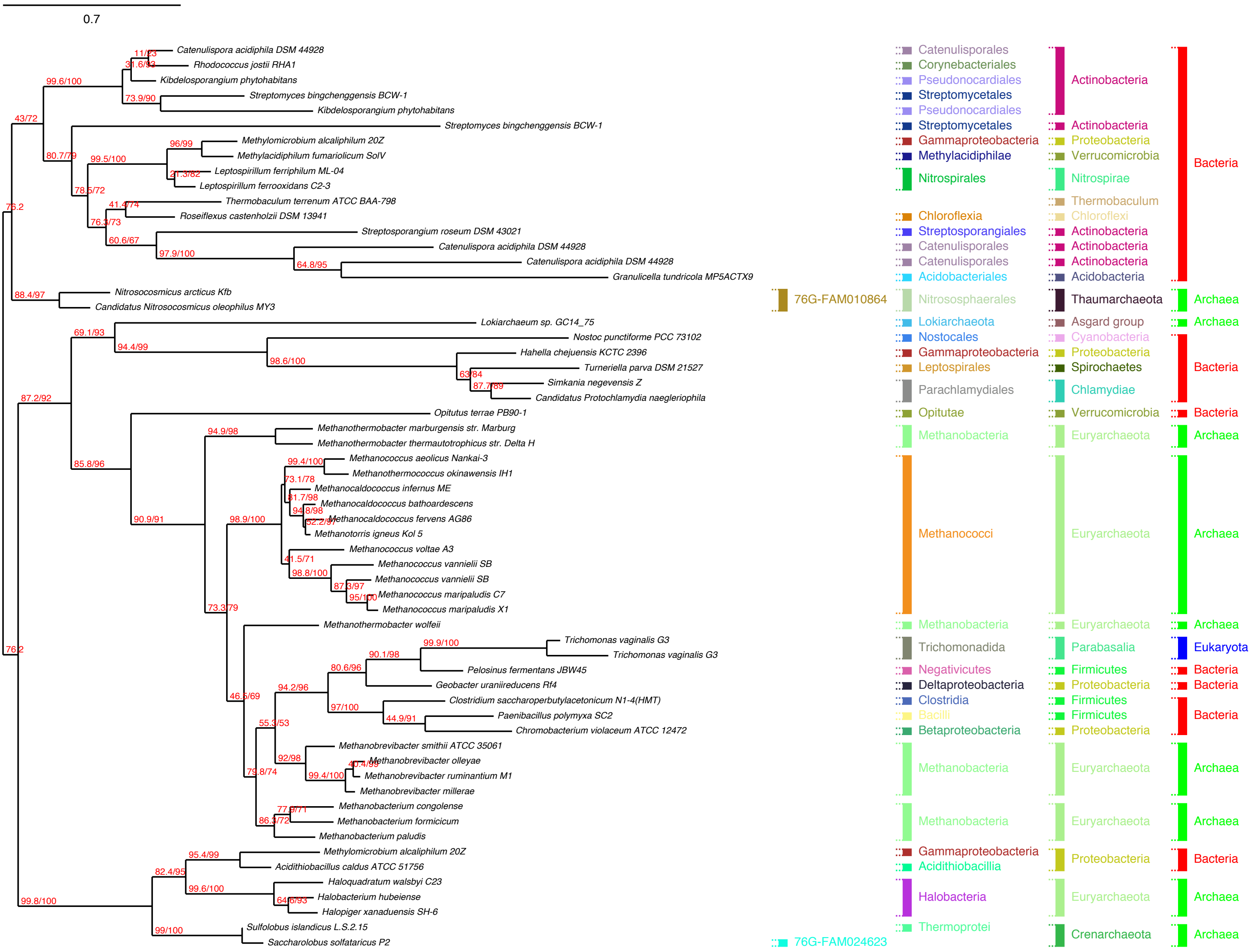

0.7
Catenulispora acidiphila DSM 44928
Rhodococcus jostii RHA1
Kibdelosporangium phytohabitans
Streptomyces bingchenggensis BCW-1
Kibdelosporangium phytohabitans
Streptomyces bingchenggensis BCW-1
Methylomicrobium alcaliphilum 20Z
Methylacidiphilum fumariolicum SolV
Leptospirillum ferriphilum ML-04
Leptospirillum ferrooxidans C2-3
Thermobaculum terrenum ATCC BAA-798
Roseiflexus castenholzii DSM 13941
Streptosporangium roseum DSM 43021
Catenulispora acidiphila DSM 44928
Catenulispora acidiphila DSM 44928
Granulicella tundricola MP5ACTX9
Nitrosocosmicus arcticus Kfb
Candidatus Nitrosocosmicus oleophilus MY3
Lokiarchaeum sp. GC14_75
Nostoc punctiforme PCC 73102
Hahella chejuensis KCTC 2396
Turneriella parva DSM 21527
Simkania negevensis Z
Candidatus Protochlamydia naegleriophila
Opitutus terrae PB90-1
Methanothermobacter marburgensis str. Marburg
Methanothermobacter thermautotrophicus str. Delta H
Methanococcus aeolicus Nankai-3
Methanothermococcus okinawensis IH1
Methanocaldococcus infernus ME
Methanocaldococcus bathoardescens
Methanocaldococcus fervens AG86
Methanotorris igneus Kol 5
Methanococcus voltae A3
Methanococcus vannielii SB
Methanococcus vannielii SB
Methanococcus maripaludis C7
Methanococcus maripaludis X1
Methanothermobacter wolfeii
Trichomonas vaginalis G3
Trichomonas vaginalis G3
Pelosinus fermentans JBW45
Geobacter uraniireducens Rf4
Clostridium saccharoperbutylacetonicum N1-4(HMT)
Paenibacillus polymyxa SC2
Chromobacterium violaceum ATCC 12472
Methanobrevibacter smithii ATCC 35061
Methanobrevibacter olleyae
Methanobrevibacter ruminantium M1
Methanobrevibacter millerae
Methanobacterium congolense
Methanobacterium formicicum
Methanobacterium paludis
Methylomicrobium alcaliphilum 20Z
Acidithiobacillus caldus ATCC 51756
Haloquadratum walsbyi C23
Halobacterium hubeiense
Halopiger xanaduensis SH-6
Sulfolobus islandicus L.S.2.15
Saccharolobus solfataricus P2
11/63
31.6/93
99.6/100
73.9/90
43/72
96/99
80.7/79
99.5/100
21.3/82
78.5/72
41.4/74
76.2
76.3/73
60.6/67
97.9/100
64.8/95
88.4/97
69.1/93
94.4/99
98.6/100
63/84
87.7/89
87.2/92
94.9/98
99.4/100
73.1/78
81.7/98
94.8/98
52.2/97
85.8/96
90.9/91
98.9/100
41.5/71
98.8/100
87.3/97
95/100
73.3/79
76.2
99.9/100
90.1/98
80.6/96
94.2/96
97/100
44.9/91
46.6/69
55.3/53
92/98
40.4/99
99.4/100
79.8/74
77.9/71
86.3/72
95.4/99
82.4/95
99.6/100
64.6/93
99.8/100
99/100
76G-FAM010864
76G-FAM024623
Catenulisporales
Corynebacteriales
Pseudonocardiales
Streptomycetales
Pseudonocardiales
Streptomycetales
Gammaproteobacteria
Methylacidiphilae
Nitrospirales
Chloroflexia
Streptosporangiales
Catenulisporales
Catenulisporales
Catenulisporales
Acidobacteriales
Nitrososphaerales
Lokiarchaeota
Nostocales
Gammaproteobacteria
Leptospirales
Parachlamydiales
Opitutae
Methanobacteria
Methanococci
Methanobacteria
Trichomonadida
Negativicutes
Deltaproteobacteria
Clostridia
Bacilli
Betaproteobacteria
Methanobacteria
Methanobacteria
Gammaproteobacteria
Acidithiobacillia
Halobacteria
Thermoprotei
Actinobacteria
Actinobacteria
Proteobacteria
Verrucomicrobia
Nitrospirae
Thermobaculum
Chloroflexi
Actinobacteria
Actinobacteria
Actinobacteria
Acidobacteria
Thaumarchaeota
Asgard group
Cyanobacteria
Proteobacteria
Spirochaetes
Chlamydiae
Verrucomicrobia
Euryarchaeota
Euryarchaeota
Euryarchaeota
Parabasalia
Firmicutes
Proteobacteria
Firmicutes
Firmicutes
Proteobacteria
Euryarchaeota
Euryarchaeota
Proteobacteria
Euryarchaeota
Crenarchaeota
Bacteria
Archaea
Archaea
Bacteria
Bacteria
Archaea
Archaea
Archaea
Eukaryota
Bacteria
Bacteria
Bacteria
Archaea
Archaea
Bacteria
Archaea
Archaea

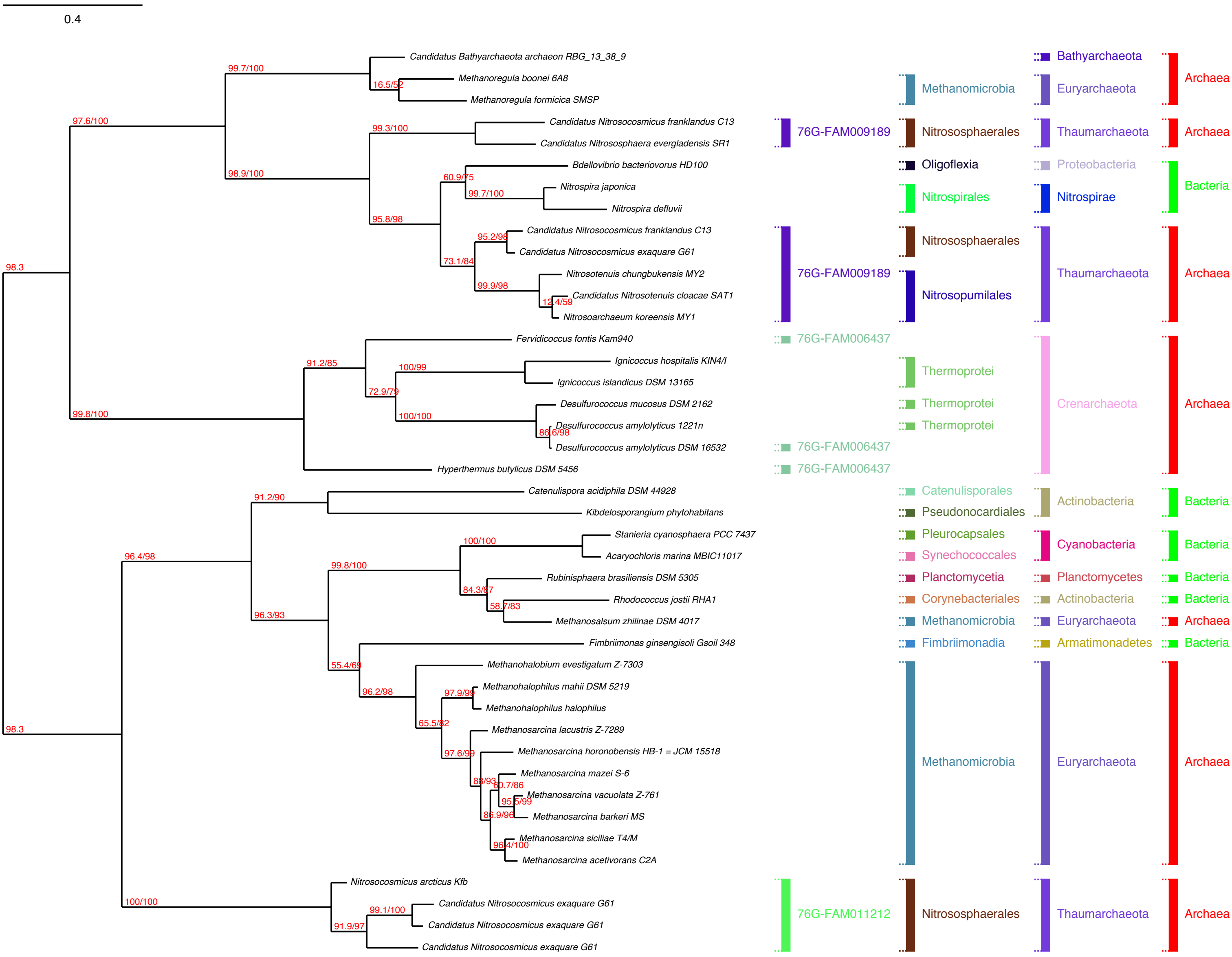
0.4
Candidatus Bathyarchaeota archaeon RBG_13_38_9
Methanoregula boonei 6A8
Methanoregula formicica SMSP
Candidatus Nitrosocosmicus franklandus C13
Candidatus Nitrososphaera evergladensis SR1
Bdellovibrio bacteriovorus HD100
Nitrospira japonica
Nitrospira defluvii
Candidatus Nitrosocosmicus franklandus C13
Candidatus Nitrosocosmicus exaquare G61
Nitrosotenuis chungbukensis MY2
Candidatus Nitrosotenuis cloacae SAT1
Nitrosoarchaeum koreensis MY1
Fervidicoccus fontis Kam940
Ignicoccus hospitalis KIN4/I
Ignicoccus islandicus DSM 13165
Desulfurococcus mucosus DSM 2162
Desulfurococcus amylolyticus 1221n
Desulfurococcus amylolyticus DSM 16532
Hyperthermus butylicus DSM 5456
Catenulispora acidiphila DSM 44928
Kibdelosporangium phytohabitans
Stanieria cyanosphaera PCC 7437
Acaryochloris marina MBIC11017
Rubinisphaera brasiliensis DSM 5305
Rhodococcus jostii RHA1
Methanosalsum zhilinae DSM 4017
Fimbriimonas ginsengisoli Gsoil 348
Methanohalobium evestigatum Z-7303
Methanohalophilus mahii DSM 5219
Methanohalophilus halophilus
Methanosarcina lacustris Z-7289
Methanosarcina horonobensis HB-1 = JCM 15518
Methanosarcina mazei S-6
Methanosarcina vacuolata Z-761
Methanosarcina barkeri MS
Methanosarcina siciliae T4/M
Methanosarcina acetivorans C2A
Nitrosocosmicus arcticus Kfb
Candidatus Nitrosocosmicus exaquare G61
Candidatus Nitrosocosmicus exaquare G61
Candidatus Nitrosocosmicus exaquare G61
98.3
97.6/100
99.7/100
16.5/52
98.9/100
99.3/100
95.8/98
60.9/75
99.7/100
73.1/84
95.2/98
99.9/98
12.4/59
99.8/100
91.2/85
72.9/79
100/99
100/100
86.6/98
98.3
96.4/98
91.2/90
96.3/93
99.8/100
100/100
84.3/87
58.7/83
55.4/69
96.2/98
65.5/82
97.9/99
97.6/99
88/93
60.7/86
95.5/99
86.9/96
96.4/100
100/100
91.9/97
99.1/100
76G-FAM009189
76G-FAM009189
76G-FAM006437
76G-FAM006437
76G-FAM006437
76G-FAM011212
Methanomicrobia
Nitrososphaerales
Oligoflexia
Nitrospirales
Nitrososphaerales
Nitrosopumilales
Thermoprotei
Thermoprotei
Thermoprotei
Catenulisporales
Pseudonocardiales
Pleurocapsales
Synechococcales
Planctomycetia
Corynebacteriales
Methanomicrobia
Fimbriimonadia
Methanomicrobia
Nitrososphaerales
Bathyarchaeota
Euryarchaeota
Thaumarchaeota
Proteobacteria
Nitrospirae
Thaumarchaeota
Crenarchaeota
Actinobacteria
Cyanobacteria
Planctomycetes
Actinobacteria
Euryarchaeota
Armatimonadetes
Euryarchaeota
Thaumarchaeota
Archaea
Archaea
Bacteria
Archaea
Archaea
Bacteria
Bacteria
Bacteria
Bacteria
Archaea
Bacteria
Archaea
Archaea

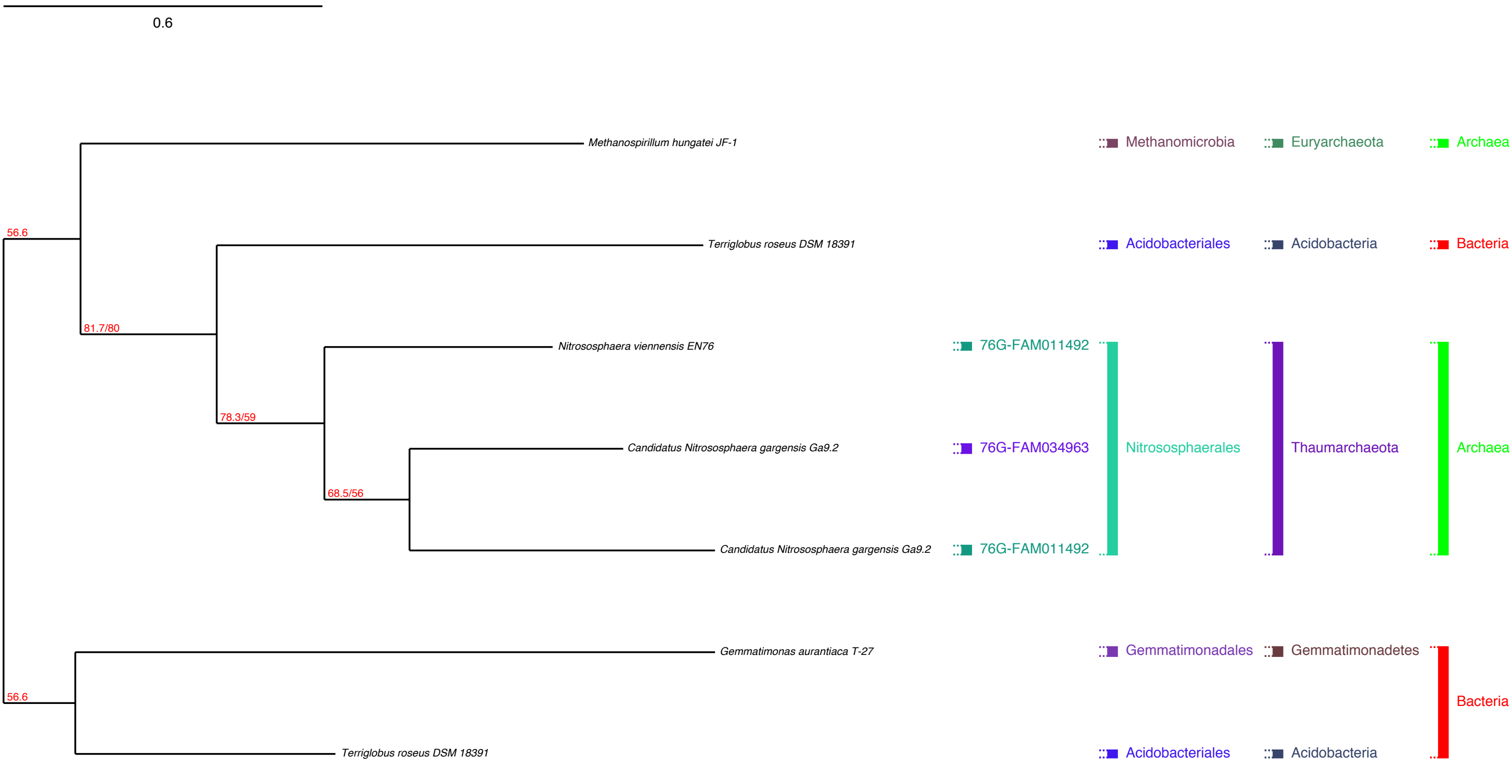
0.6
56.6
81.7/80
78.3/59
68.5/56
56.6
Methanospirillum hungatei JF-1
Terriglobus roseus DSM 18391
Nitrososphaera viennensis EN76
Candidatus Nitrososphaera gargensis Ga9.2
Candidatus Nitrososphaera gargensis Ga9.2
Gemmatimonas aurantiaca T-27
Terriglobus roseus DSM 18391
76G-FAM011492
76G-FAM034963
76G-FAM011492
Methanomicrobia
Acidobacteriales
Nitrososphaerales
Gemmatimonadales
Acidobacteriales
Euryarchaeota
Acidobacteria
Thaumarchaeota
Gemmatimonadetes
Acidobacteria
Archaea
Bacteria
Archaea
Bacteria

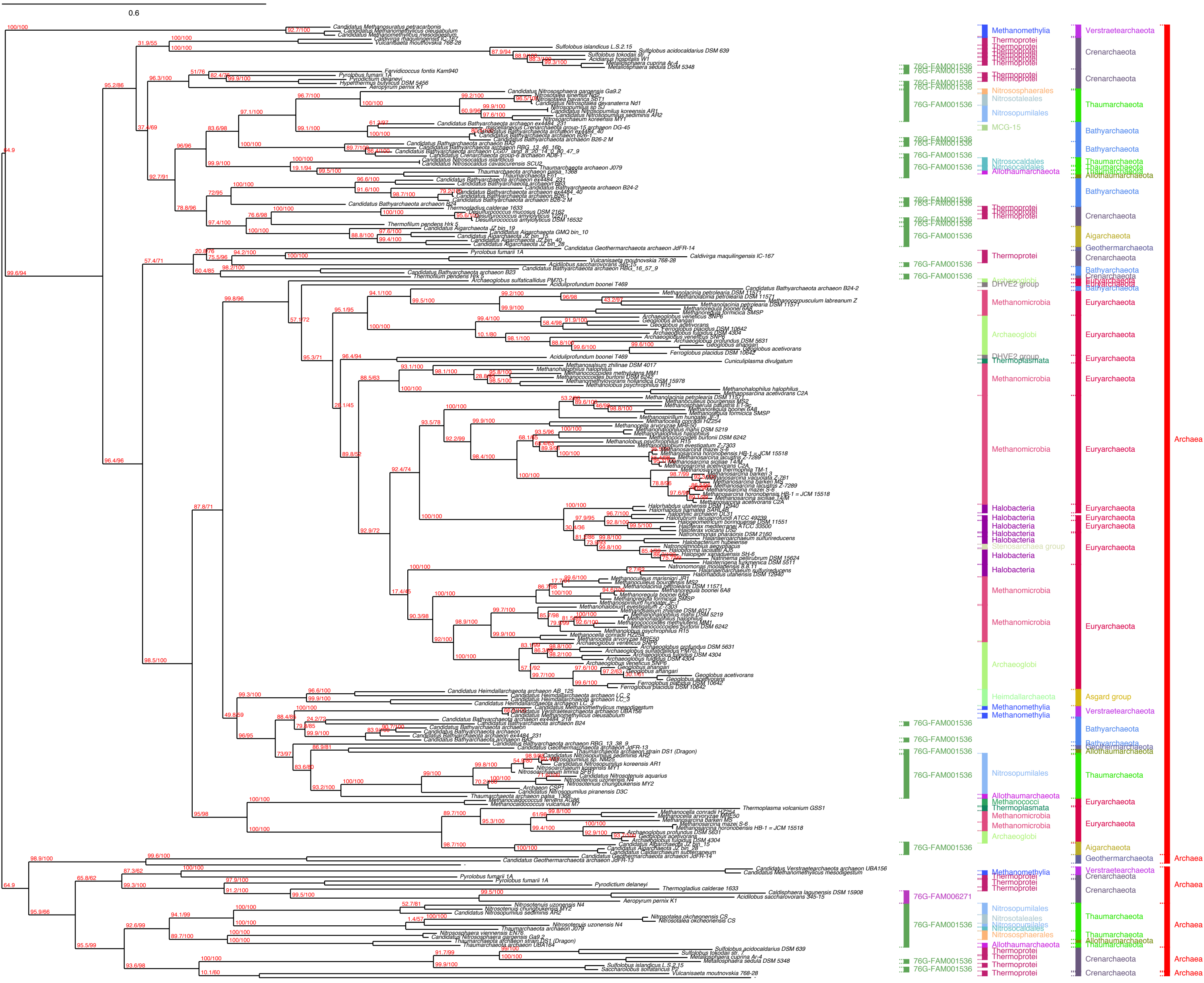

0.6
Methanomethylia
Thermoprotei
76G-FAM001536
Nitrososphaerales
Nitrosotaleales
Nitrosopumilales
MCG-15
Nitrosocaldales
Allothaumarchaeota
Archaeoglobi
DHVE2 group
Methanomicrobia
Thermoplasmata
Halobacteria
Stenosarchaea group
Heimdallarchaeota
Methanococci
76G-FAM006271
Verstraetearchaeota
Crenarchaeota
Thaumarchaeota
Bathyarchaeota
Aigarchaeota
Geothermarchaeota
Euryarchaeota
Asgard group
Archaea
